# Virtual metabolic human dynamic model for pathological analysis and therapy design for diabetes

**DOI:** 10.1101/2020.08.29.269399

**Authors:** Hiroyuki Kurata

## Abstract

A virtual metabolic human model is a valuable complement to experimental biology and clinical studies, because *in vivo* research involves serious ethical and technical problems. I have proposed a multi-organ and multi-scale kinetic model that formulates the reactions of enzymes and transporters with the regulation of enzyme activities and hormonal actions under prandial and rest conditions. The model consists of 202 ordinary differential equations for metabolites with 217 reaction rates and 1132 kinetic parameter constants. It is the most comprehensive, largest and highly predictive model of the whole-body metabolism. Use of the model revealed the mechanisms by which individual disorders, such as steatosis, β cell dysfunction and insulin resistance, were combined to cause type 2 diabetes. The model predicted a glycerol kinase inhibitor to be an effective medicine for type 2 diabetes, which not only decreased hepatic triglyceride but also reduced plasma glucose. The model also enabled us to rationally design combination therapy.

## Introduction

Virtual metabolic human is an attractive goal for synthetic biology, systems biology and bioinformatics, which greatly contributes to advances in medicine and life science. Many scientists have raised its concepts and have been developing computational frameworks on genome-scale gene networks and whole-body scale omics data (Thiele et al., 2020; Thiele et al., 2013; Viceconti and Hunter, 2016). In 2019, the virtual metabolic human database has been presented to facilitate computational modeling by linking genome-scale networks of human metabolism to diseases and nutrition (Noronha et al., 2019). On the other hand, multi-scale, large-scale dynamic models have been developed to understand human metabolism (Ashworth et al., 2016; Berndt et al., 2018a; Berndt and Holzhutter, 2018; Berndt et al., 2018b; Kim et al., 2007; Konig et al., 2012; Li et al., 2010; Sluka et al., 2016).

Metabolism plays a critical role in human health and diseases (Frayn, 2010). Perturbation of genetics and changes in lifestyle habitats, including excessive diet and inactivity, result in the development and progression of complex metabolic diseases, such as hyperglycemia, hyperlipidemia, obesity, non-alcoholic fatty liver disease (NAFLD) (Kitade et al., 2017; Xia et al., 2019) and diabetes(Ashcroft and Rorsman, 2012; Perry et al., 2014; Saltiel, 2001; Xia et al., 2019). A systems approach is necessary to elucidate the molecular mechanisms causing such metabolic dysfunctions and to propose the strategies for the prevention and treatment of them (Kitano, 2010). Since *in vivo* studies of human metabolism are hampered with serious ethical and technical problems, computational models are required to complement the *in vivo* studies (Eissing et al., 2011; Iyengar et al., 2012; Maldonado et al., 2018).

So far mathematical models have simulated human metabolism. Early models extensively investigated glucose-insulin metabolism to analyze the effect of insulin secretion on glucose homeostasis (Li et al., 2006; Sedaghat et al., 2002; Tolic et al., 2000). They used the compartment models that regarded particular metabolites as the representatives responsible for their associated functions such as glycolysis, gluconeogenesis, glycogenolysis, glycogenesis, and triglyceride (TG) synthesis/degradation. Compartment models simulated the glucagon/insulin-controlled glucose homeostasis by linking liver to other organ compartments (Hetherington et al., 2012; Pearson et al., 2014), and suggested the mechanisms by which changes in a ratio of carbohydrate to lipid alter hepatic TG synthesis through insulin action and generate different types of diabetes (Hassell Sweatman, 2020; Pratt et al., 2015).

Since those compartment models were the coarse-grained models, their applications were limited to an understanding of the specific functions. To overcome this limitation, biochemistry-based models were constructed that assigned a rate equation to each metabolic reaction within a cell of liver and skeletal muscle while considering allosteric effectors, enzyme activity regulation, and hormone-dependent reversible phosphorylation (Berndt et al., 2018a; Konig et al., 2012; Li et al., 2010). The kinetics were measured by means of *in vitro* assays. In 2019 Berndt et al developed a genome-scale, detailed kinetic model of hepatic cells that formulated thousands of enzymes, transporters, and hormone-dependent regulations (Berndt et al., 2018a).

Those biochemistry-based models pay attention to cells of liver and skeletal muscle, while it is important to consider the rest of the human body because organs are tightly connected with each other through blood (Ashworth et al., 2016; Sluka et al., 2016; Xu et al., 2011). Xu et al integrated hepatic glycogen regulation with extra-hepatic fuel metabolism under prandial and rest conditions in the whole-body context (Xu et al., 2011). Sluka et al. proposed a liver-centric model for acetaminophen pharmacology and metabolism (Sluka et al., 2016). They integrated three-scale modules of enzyme reactions within a cell, physiologically based pharmacokinetics of acetaminophen at organs, and its distribution at the whole-body level. Ashworth et al developed a spatial kinetic model of hepatic glucose and lipid metabolism and treated the sinusoidal tissue units instead of the single hepatocyte (Ashworth et al., 2016). They identified critical differences between periportal and pericentral cells, indicating high susceptibility to pericentral steatosis during the development of steatosis. Berndt et al. also presented a dynamic model of the sinusoidal tissue units to suggest that structural properties, enzymatic properties and regional blood flows are equally important for an understanding of liver functionality (Berndt and Holzhutter, 2018; Berndt et al., 2018b). Kim et al proposed a whole-body computational model to simulate hormonal control over plasma glucose and TG under a limited condition of physical exercise (Kim et al., 2007). They decomposed the whole body into several organs, assigned each major metabolite within organs to an ordinary differential equation. Palumbo et al. added details of subjects’ characteristics to the Kim’s model to simulate the effects on personal metabolic homeostasis during exercise (Palumbo et al., 2018).

Construction of whole-body metabolic models has just started, but the previous models are fit for a specific purpose and their application is limited due to lack of comprehensive, molecular mechanisms and due to excessive simplification. At present a virtual metabolic human model that integrates comprehensive molecular mechanisms for each organ is highly expected not only to reproduce a variety of physiological and metabolic functions, but also to analyze pathology and design therapy while considering side effects. To achieve these requirements, I have first proposed a virtual metabolic human dynamic model or the multi-organ and multi-scale kinetic model that accurately simulates the dynamics in key metabolites of glucose, lactate, pyruvate, glycerol, alanine, glycogen, free fatty acid (FFA) and TG under prandial and rest conditions. To enhance the accuracy and applicability, the model incorporated nucleotide cofactors that are critically responsible for global regulations and energy balance, while conserving essential reaction pathways of carbohydrates and lipids for each organ. Use of the model revealed the pathological mechanisms by which the individual disorders of steatosis, β cell dysfunction and insulin resistance (IR) are combined to cause diabetes. The model successfully predicted a glycerol kinase inhibitor to be a new medicine for type 2 diabetes and enabled us to rationally design combination therapy.

## Results

### Model validation by experimental data

A schematic diagram of the whole body is shown in **Figure 1**. I constructed the union of all the metabolic networks of the organs (**Figure 2**). Each organ metabolic network was built by selecting organ-specific reactions from the union network. Details of dynamic model construction is described in Methods. The proposed kinetic model simulated the time course of plasma glucose and insulin concentrations after an overnight fast and following a single meal of 100 g glucose and 33 g TG, as shown in **Figure 3AB**. The plasma glucose concentration increased to a peak around 60 min, then decreased to approximately 5 mM. An increase in plasma glucose triggered the insulin secretion from pancreas to enhance the uptake of plasma glucose by the organs of liver, skeletal muscle, gastrointestinal (GI), and adipose tissue. The plasma insulin concentration greatly increased to a peak, which was caused by an increase in plasma glucose, and then decreased with a decrease in plasma glucose. In liver and skeletal muscle, the glycogen concentration increased due to insulin action after a meal. At rest, liver glycogen was degraded into glucose (glyocogenolysis), which was released into blood. Skeletal muscle glycogen remained to be degraded, as suggested by (Jensen et al., 2011). Insulin dramatically changed the liver status between glucose utilization and production. Plasma lactate increased to a peak after the meal and then decreased. Skeletal muscle and adipose tissue converted a part of the utilized glucose into lactate and released it into blood. Plasma FFA decreased after the meal, showing a dump, and then gradually increased. In adipose tissue, insulin induced TG synthesis from FFA and glycerol-3-P (GRP) after the meal, which decreased the release of FFA from adipose tissue. On the other hand, insulin hardly affected the uptake of FFA by liver and skeletal muscle. Consequently, plasma FFA decreased soon after a meal. Plasma TG, ingredient of chylomicrons, slowly increased after a meal, and then decreased. Plasma TG was absorbed mainly by adipose tissue.

**Figure 1.**
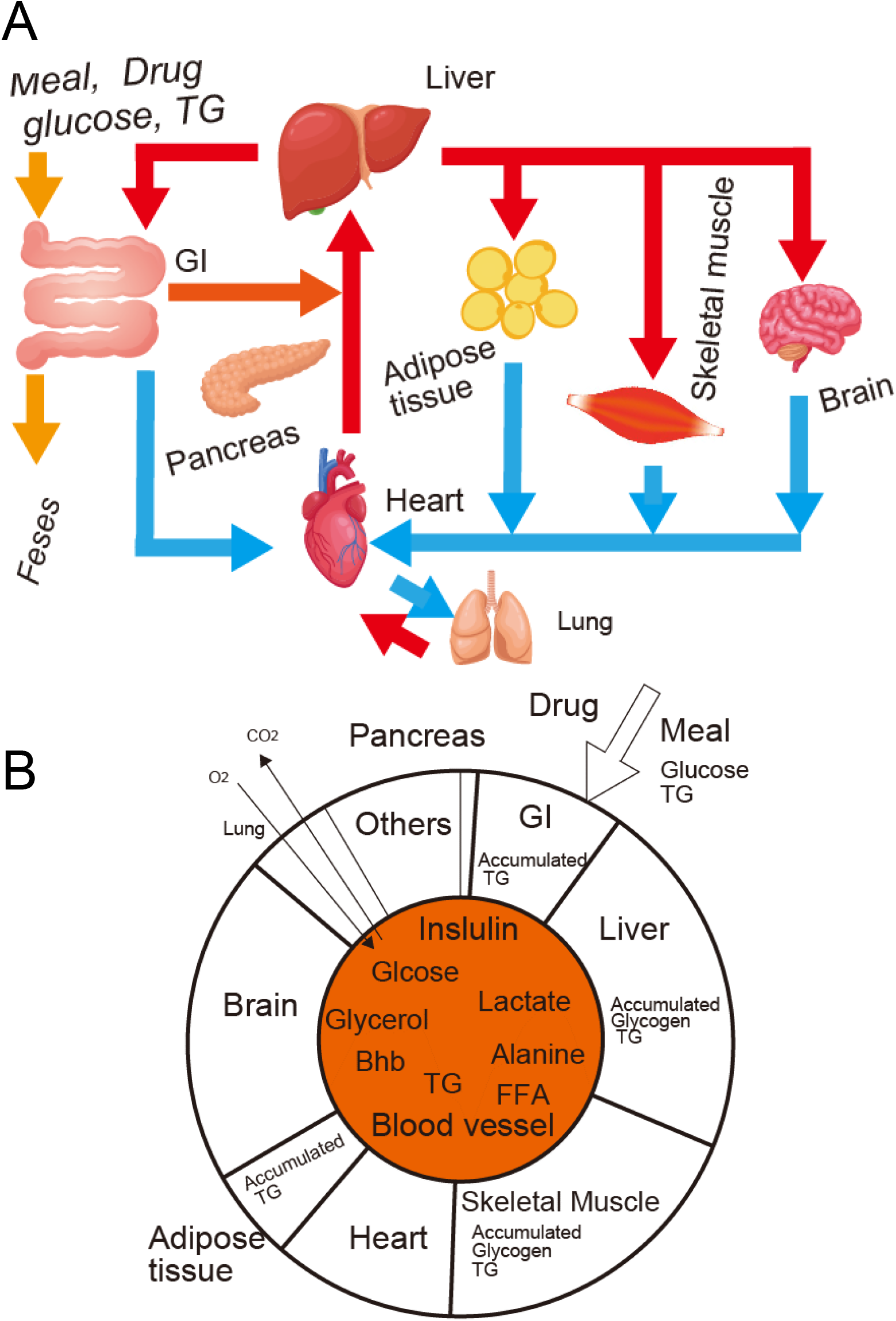
Schematic diagram of the human whole-body metabolism. (A) Realistic multi-organ model. (B) Perfect mixing model of the whole-body metabolism Oxygen and carbon dioxide concentrations are assumed to be constant in the whole body, thus the kinetic model does not include lung and them.

**Figure 2.**
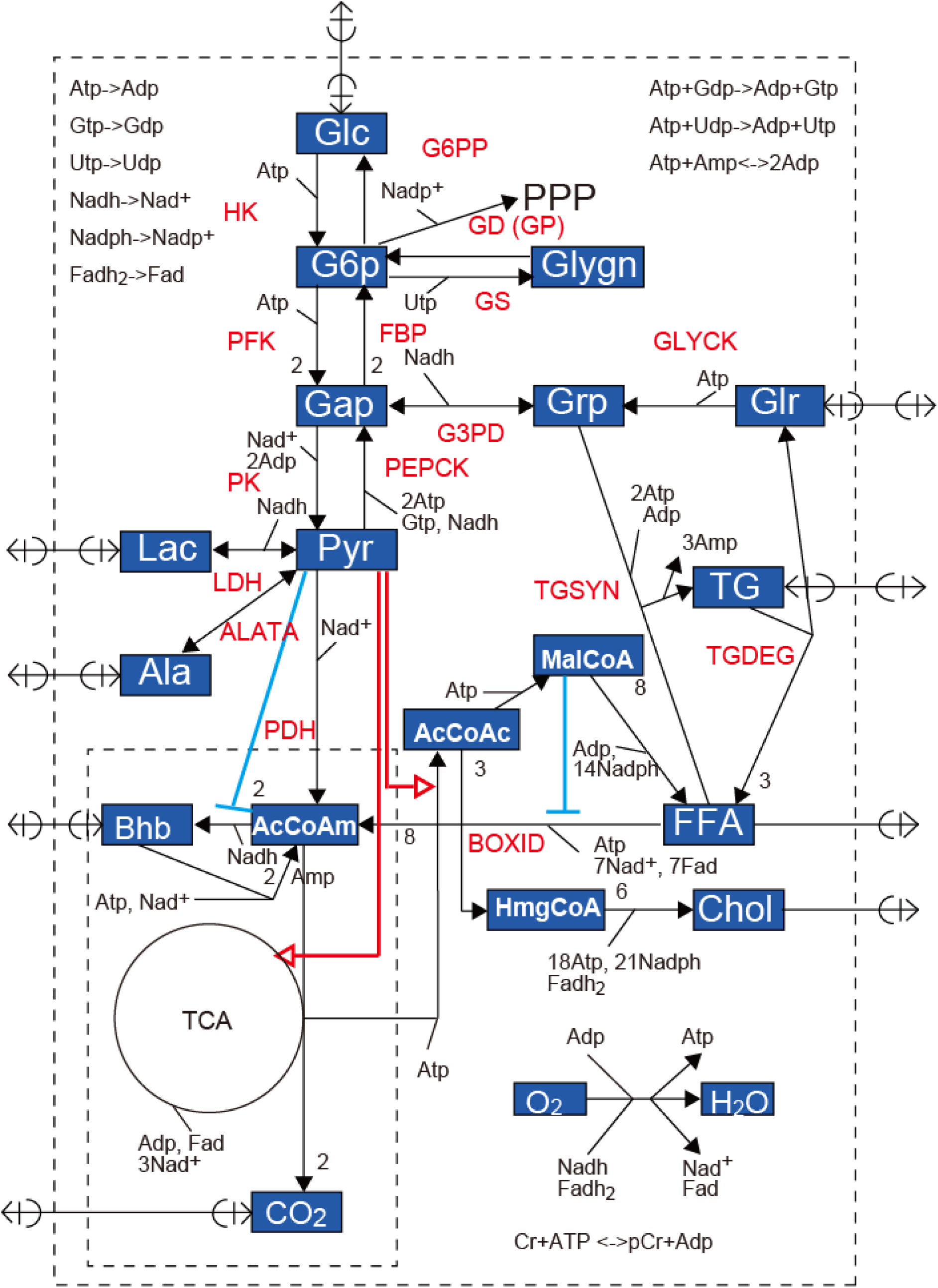
Union map of all organ metabolic networks

**Figure 3.**
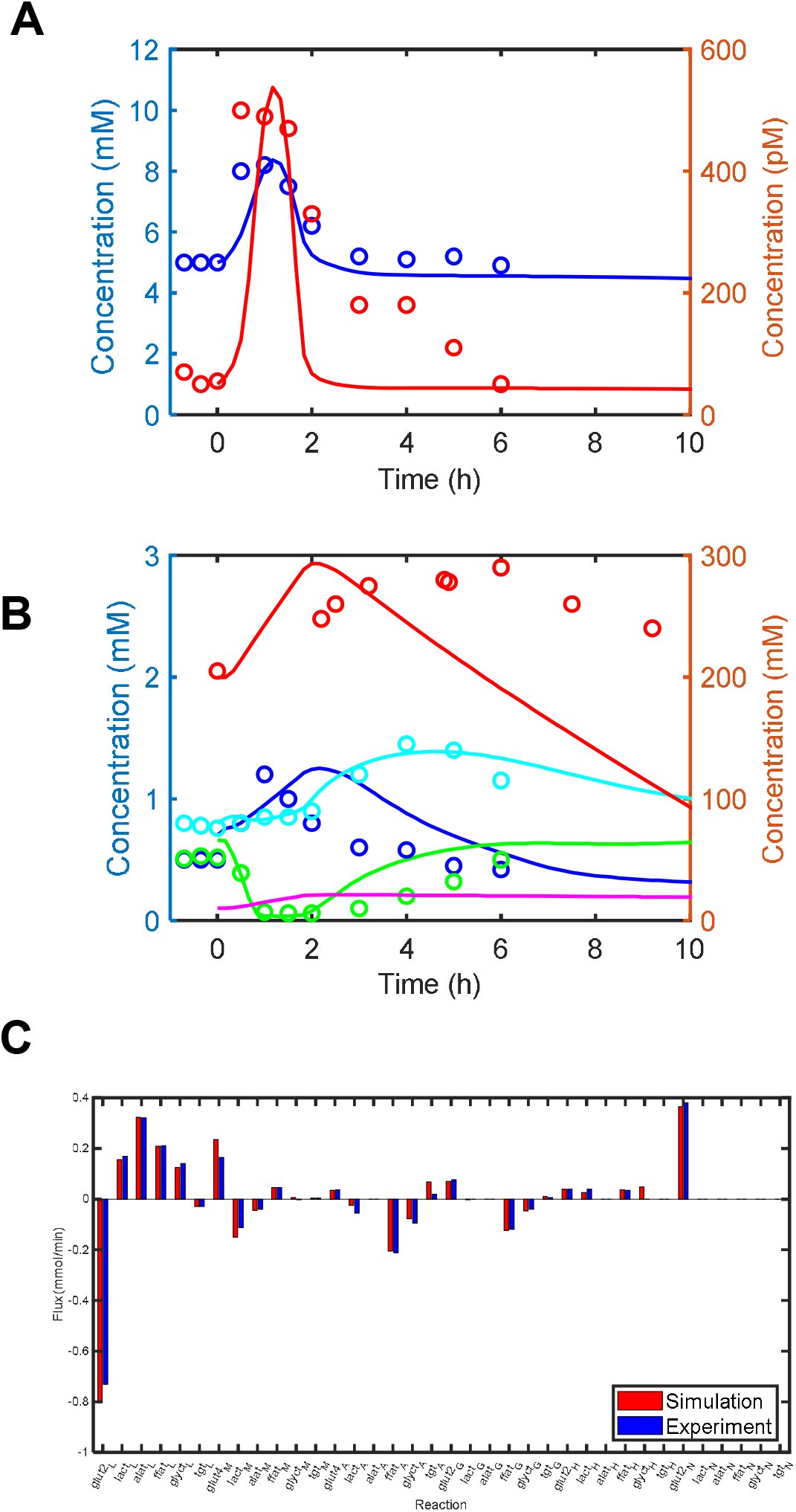
Experimental validation of the virtual metabolic human dynamic model. (A, B) Time course of key metabolites in blood, liver and skeletal muscle after an overnight fast and following a single meal of 100 g glucose and 33 g fat. (A) Circles and lines indicate the experimental and simulated concentrations of plasma glucose (blue) and insulin (red), respectively. (B) Circles and lines indicate the experimental and simulated concentrations of plasma lactate (blue), plasma FFA (green), plasma TG (cyan), hepatic glycogen (red) and skeletal muscle glycogen (magenta). (C) Transport/exchange fluxes at rest between blood and each organ. Red and blue bars indicate the simulated and experimental fluxes, respectively.

As shown in **Figure 3C**, the simulated exchange fluxes between each organ and blood at rest were consistent with the experimental data. At rest, skeletal muscle released alanine and lactate; adipose tissue released glycerol and lactate. Liver utilized the three carbon metabolites of lactate, glycerol and alanine to synthesize glucose and released it into blood, indicating that liver recycled the three metabolites into glucose through gluconeogenesis. I confirmed that utilized glycerol was converted to glycelaldehyde-3-phosphate (GAP) through GRP in liver (data not shown), indicating that plasma glycerol is used for gluconeogenesis. The model reproduced the glucose-lactate/glycerol/alanine cycles. Liver synthesized TG and cholesterol and then released them into blood. Heart utilized glucose, lactate, and FFA. GI utilized glucose and degraded TG storage into plasma FFA and glycerol. Adipose tissue utilized glucose and TG; it released lactate, FFA and glycerol. Brain exclusively utilized plasma glucose.

### Discrepancy between experimental data and simulation

I illustrated some discrepancies between experimental data and simulation (**Figure 3**). The simulated insulin pulse was sharper than the experimental pulse with a long tail. While the simulated glycogen concentration in liver increased to a peak 1 h after the glucose peak, the duration achieving the peak was shorter than that of the experimental data (4-6h). In other words, the simulated glycogen synthesis was faster than experimental data. In addition, the simulated glycogen decrease more rapidly than experimental data. While the simulated plasma lactate increased to a peak at 2 h following the glucose peak, the duration (2h) achieving the peak was longer than the experimental lactate data.

### Critically important features

The proposed model simulated the switching function between DNL and Bhb synthesis with respect to pyruvate in liver, as shown in **Figure 4A**. A high concentration of pyruvate increased DNL flux; a very low concentration of pyruvate induced Bhb synthesis flux. The β-oxidation flux gradually decreased with an increase in pyruvate, while the DNL flux and pyruvate dehydrogenase (PDH) flux increased. It corresponded to the observation of the reverse relationships between DNL and β-oxidation and between glycolysis and β-oxidation (Frayn, 2010).

**Figure 4.**
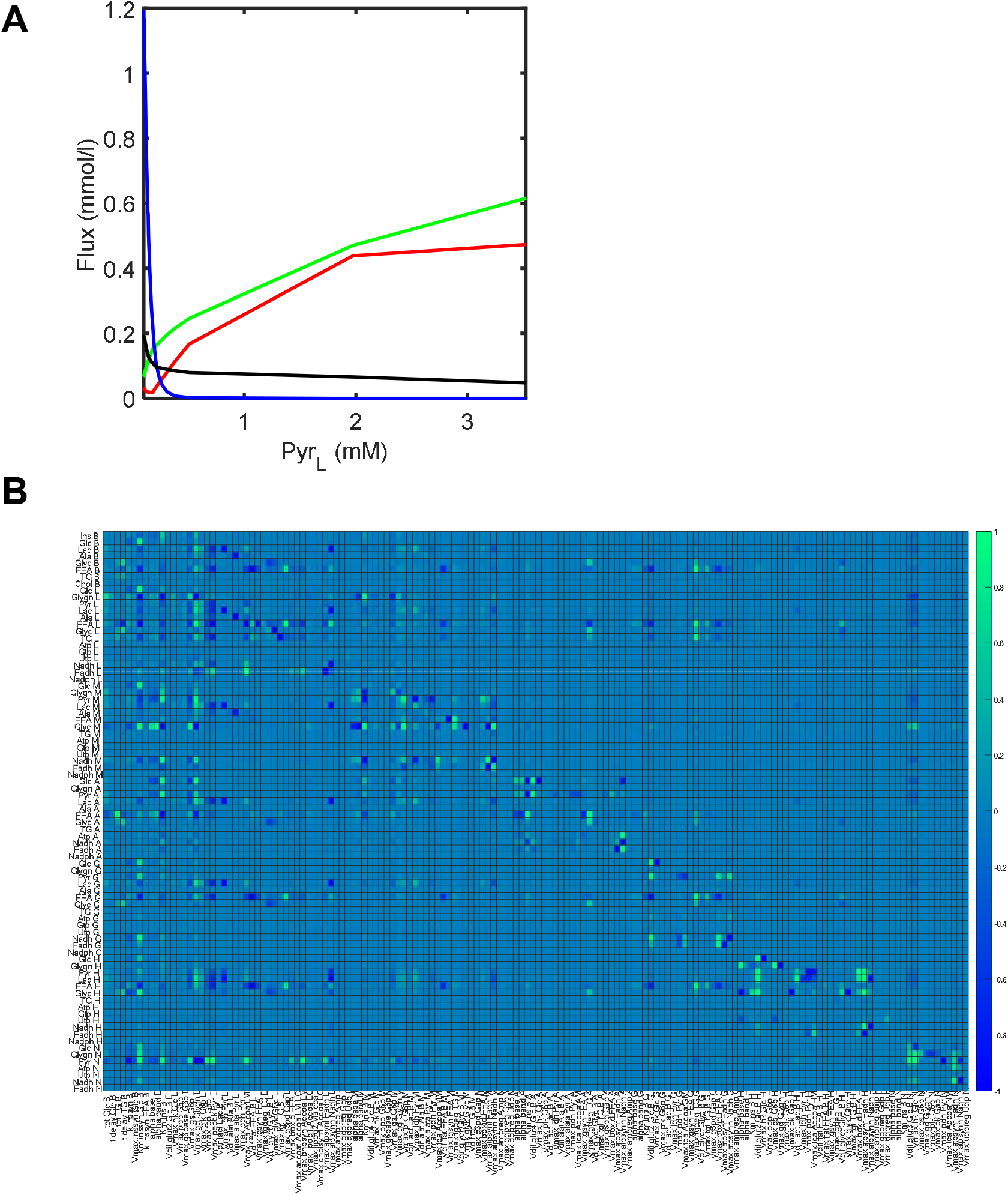
Critically important features of the virtual metabolic human dynamic model. The proposed model was simulated during 480 h after an overnight fast and following a single meal of 100 g glucose and 33 g fat. (A) Changes in the simulated fluxes of key reactions with respect to hepatic pyruvate. The four fluxes of the PDH reaction (red), DNL (green), Bhb synthesis (blue), and β-oxidation (black) were plotted with respect to pyruvate concentration in liver. (B) Dynamic sensitivity analysis of metabolite concentrations at rest with respect to a single kinetic constant. The dynamic sensitivity was calculated at 10 h, where the value of a single kinetic parameter was changed by 1.1-fold.

I performed dynamic sensitivity analysis of metabolite concentrations with respect to specific kinetic parameters, as shown in **Figure 4B**. The plasma glucose level was highly robust with respect to changes in the kinetic parameters except the insulin-related parameters including Km_inssyn_Glc_B, Km_Ins_B_L and Km_Ins_B_M. The set-point of plasma glucose was controlled dominantly by insulin. Km_inssyn_Glc_B, which determines glucose-stimulated insulin secretion, was the most effective parameter that altered the set-point of plasma glucose. Km_Ins_B_L and Km_Ins_B_M, which determine insulin-stimulated glucose uptake in liver and skeletal muscle, were the second and third critical parameters, respectively. Km_Ins_B_A hardly affected the set-point, because the glucose uptake rate of adipose tissue was much less than that by skeletal muscle (**Figure 3C**). In addition to the insulin-related parameters, nucleotide cofactor synthesis-related parameters greatly affected some metabolites, suggesting that incorporation of nucleotide cofactors induces remarkable changes. It is because nucleotide cofactors simultaneously affect multiple reactions. Hepatic FFA and TG concentrations indicated high sensitivities with respect to Vmax_tgdeg_TG_A and Vmax_tgdeg_TG_G that determine the TG degradation rate in adipose tissue and GI. It suggests that adipose tissue and GI are closely connected to liver through lipid.

### Prediction of hidden mechanisms

As shown in **Figure 5**, the hepatic FFA synthesis consisted of the two phases: the former phase was the malonyl-CoA increasing phase (lipog1) of 0.5-1.8 h, the latter was its decreasing phase (lipog2) of 1.8-3.5 h. They corresponded to the glucose utilization and production phases. During the former phase, an increase in pyruvate enhanced malonyl-CoA production (**Figure 2**), while a decrease in G6P reduced the pentose phosphate pathway flux and NADPH production. Since NADPH, which is required for the conversion from malonyl-CoA to FFA (lipog2), was decreased, malonyl-CoA was accumulated. During the latter phase, an increase in G6P enhanced the pentose phosphate pathway flux and NADPH production. The accumulated malonyl-CoA was converted into FFA using NADPH. As shown in **Figure S1**, the simulated Bhb concentration increased at the early phase of a fasted condition, as indicated by the experimental data (Owen et al., 1990), but it gradually decreased after 150 h with a decrease in plasma FFA, which was not consistent with the experimental data that gradually increased with time. The decrease in the simulated Bhb concentration during the late fasted phase reflected the simulation result that TG (a source of FFA) in GI was depleted and only adipose tissue released FFA. We need to uncover some mechanisms that increase plasma FFA under the fasted condition.

**Figure 5.**
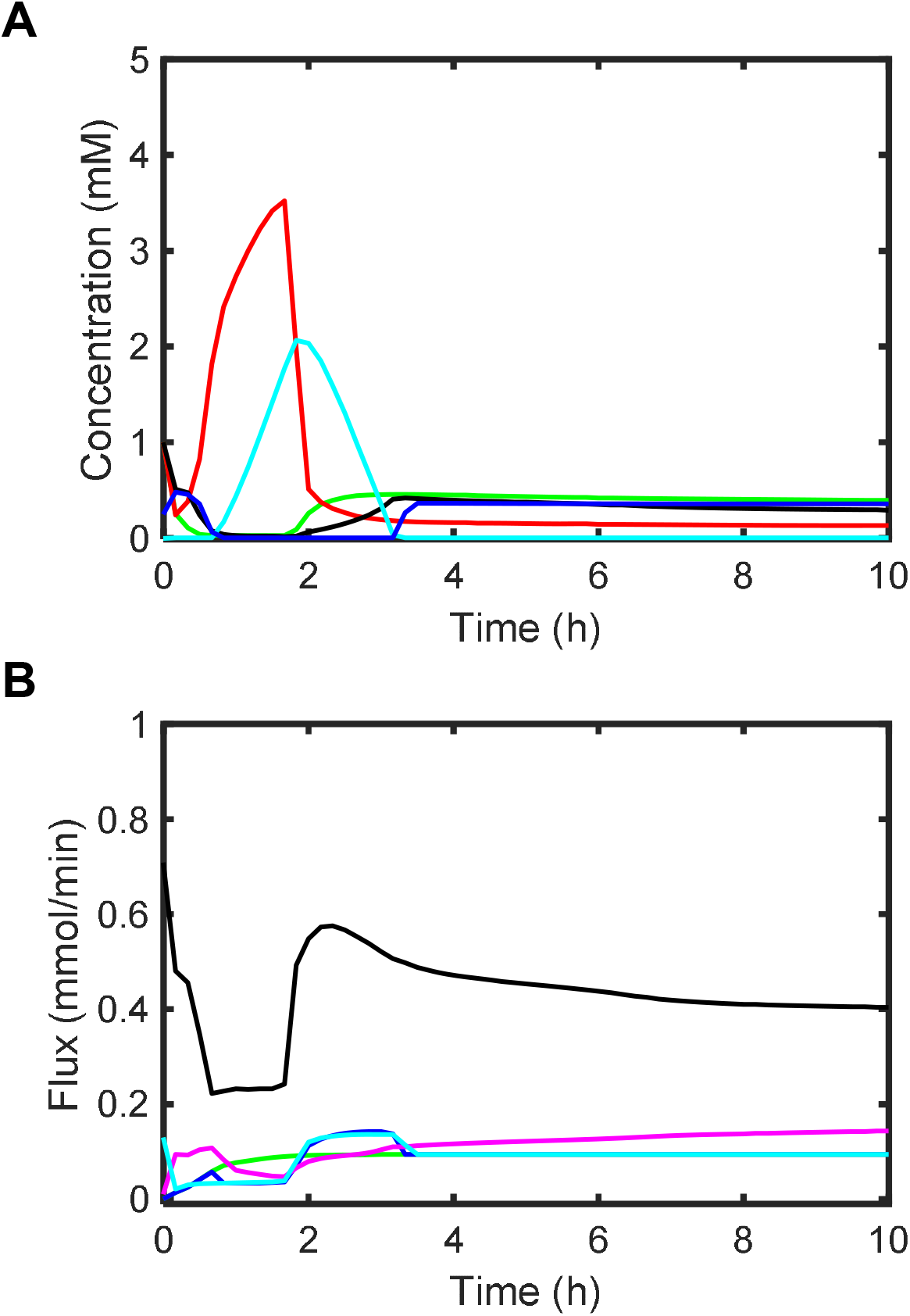
Prediction of a two-phase mechanisms of *de novo* lipogenesis in liver The proposed model was simulated for 10 h after an overnight fast and following a single meal of 100 g glucose and 33 g fat. (A) The time course of G6P (green), pyruvate (red), acetyl-CoA (black), malonyl-CoA (cyan) and NADPH (blue) concentrations in liver were simulated. (B) The time course of TCA cycle (black), lipog1(green), lipog2 (blue), β-oxidation (magenta) and pentose phosphate pathway (cyan) fluxes in liver were simulated. The lipog1 and lipog2 indicate the conversion from acetyl-CoA to malonyl-CoA and the conversion from malonyl-CoA to FFA, respectively.

### Pathological analysis

#### Individual disorders

The proposed model was used to perform pathological analysis of metabolic diseases. I simulated the effects of the individual disorders on marker metabolites such as plasma glucose, plasma insulin and hepatic TG, as shown in **Figure 6ABC**. In general, high plasma glucose and hepatic TG accumulation develop or exacerbate metabolic diseases (Ashcroft and Rorsman, 2012; Xia et al., 2019). In the steatosis model, hepatic TG increased due to acceleration of lipogenesis. Cholesterol synthesis also increased in liver (data not shown), while the dynamics of plasma insulin and plasma glucose were almost the same as that of the normal condition. In β cell dysfunction of pancreas, where glucose-stimulated insulin secretion function is impaired or Km_inssyn_Glc_B is increased, the plasma insulin concentration decreased; the peak and set-point of plasma glucose increased. Plasma glucose was less utilized by liver, skeletal muscle and adipose tissue due to the impaired insulin secretion. Interestingly, hepatic TG decreased, which conflicted with the fact that the β cell dysfunction exacerbates the diseases. Type 1 diabetes is an extreme case of the β cell dysfunction, where Km_inssyn_Glc_B approaches to infinity or insulin is hardly produced in pancreas.

**Figure 6.**
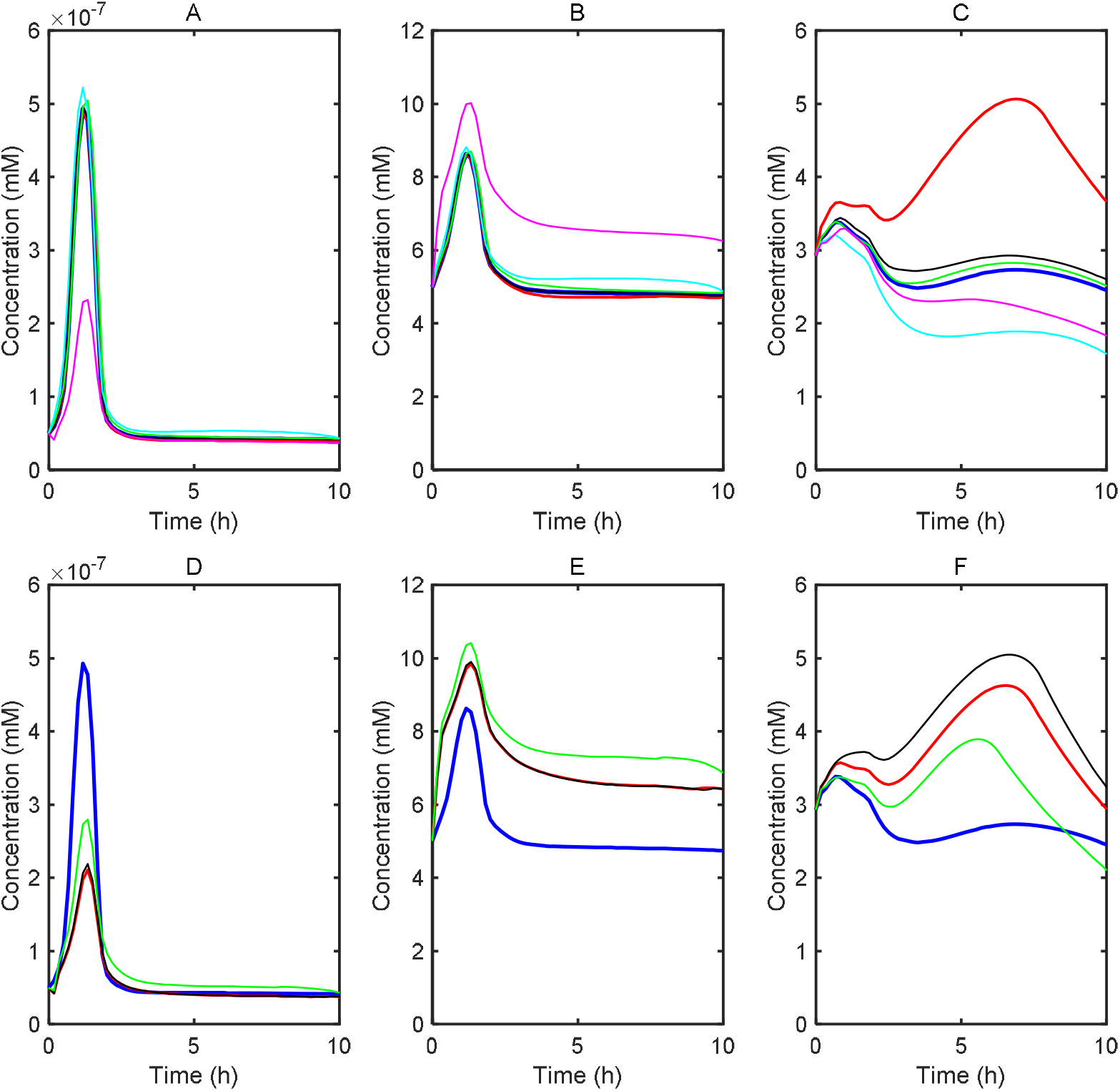
Pathological analysis using the virtual metabolic human model. (A-C) Single disorder. Time courses of plasma insulin (A), plasma glucose (B), and hepatic TG (C) for each disorder were simulated after an overnight fast and following a single meal of 100g glucose and 33 g TG. The lines indicate the normal condition (blue), steatosis (red), IR_A (black), IR_M (green), IR_L (cyan), and β-cell dysfunction (magenta). Specifically, steatosis is built by multiplying Vmax_accoat_Accoa_LM_LC, Vmax_lipog1_Accoa_LC, Vmax_lipog2_Malcoa_L, and Vmax_tgsyn_FFA_L, Vmax_cholsyn1_Accoa_LC by 2. β-cell dysfunction is built by multiplying Km_inssyn_Glc_B by 1.5. IR_A is built by multiplying Km_Ins_B_A by 1.5. IR_M is built by multiplying Km_Ins_B_M by 1.5. IR_L is built by multiplying Km_Ins_B_L by 1.5. (D-F) Combined disorders. Time courses of plasma insulin (D), plasma glucose (E), and hepatic TG (F) for each combined disorder model were simulated after an overnight fast and following a single meal of 100g glucose and 33 g TG. The lines indicate normal condition (blue), the combination of steatosis, β-cell dysfunction and IR_M (red), the combination of steatosis, β-cell dysfunction, IR_M, and IR_A (black), and the combination of steatosis, β-cell dysfunction, IR_M, IR_A, and IR_L (green).

IR_L increased the peak and set-point of plasma glucose. Hyperinsulinemia, which is a compensatory response to IR (DeFronzo and Tripathy, 2009; Petersen et al., 2007), was observed, as indicated by the experimental data (DeFronzo and Tripathy, 2009). Plasma glucose dropped after 8 h due to reduced glycogenesis. It is because IR_L reduces the hepatic glycogen synthesis, resulting in insufficient accumulation of glycogen after the meal (data not shown). Interestingly IR_L decreased hepatic TG, which presents a paradox that IR_L recovers hepatic steatosis of a major cause of IR_L (Perry et al., 2014). IR_M increased the peak and set-point of plasma glucose slightly. The plasma insulin slightly increased due to hyperglycemia. IR_M increased TG and DNL (data not shown) in liver. While skeletal muscle decreased the uptake of plasma glucose due to IR_M, liver utilized glucose for DNL in response to hyperinsulinemia and hyperglycemia. It indicates that plasma glucose is diverted away from muscle glycogen storage to hepatic TG. IR_A hardly increased the set-point of plasma glucose, while increasing hepatic TG and DNL (data not shown). It is because IR_A decreases the insulin-induced TG synthesis in adipose tissue, increasing FFA. The FFA is secreted into blood and enters liver to synthesize TG.

#### Combined disorders

The proposed model was applied to pathological analysis for type 2 diabetes. Particularly, I analyzed the effect of IR on progression of type 2 diabetes. I added the single disorders of IR_M, IR_A, and IR_L one by one to the base model consisting of steatosis and β cell dysfunction, as shown in **Figure 6DEF**. Addition of IR_M to the base model decreased plasma insulin and increased the peak and set-point of plasma glucose more than that of the normal condition, indicating hyperglycemia. IR_M decreased the glucose uptake by skeletal muscle, increasing hepatic TG. When IR_A was added to the steatosis, β cell dysfunction and IR_M model, the plasma insulin and glucose hardly changed; hepatic TG increased due to the enhanced release of FFA from adipose tissue. Addition of IR_L increased the set-point of plasma glucose and plasma insulin. Interestingly, it decreased hepatic TG.

### Medication analysis

The proposed model was employed to investigate how three medicines of widely prescribed type 2 diabetes of sulfonylurea, metformin and thiazolidinedione and a glycerol kinase inhibitor recover type 2 diabetes, as shown in **Figure 7ABC**. Sulfonylurea administration successfully decreased plasma glucose with an increase in plasma insulin, but increased hepatic TG as a side effect. Metformin not only act on LDH but also β-oxidation through AMPK. The LDH inhibition by metformin decreased plasma glucose and hepatic TG, while increasing plasma lactate (data not shown). Activation of β-oxidation by metformin decreased hepatic TG without changing plasma glucose. Thiazolidinedione activated TG synthesis in adipose tissue, which decreased hepatic TG and hardly changed plasma glucose. Administration of a glycerol kinase inhibitor decreased hepatic TG accumulation. Interestingly, it successfully decreased plasma glucose. The glycerol kinase inhibitor was referred to as a drug candidate for type 2 diabetes.

**Figure 7.**
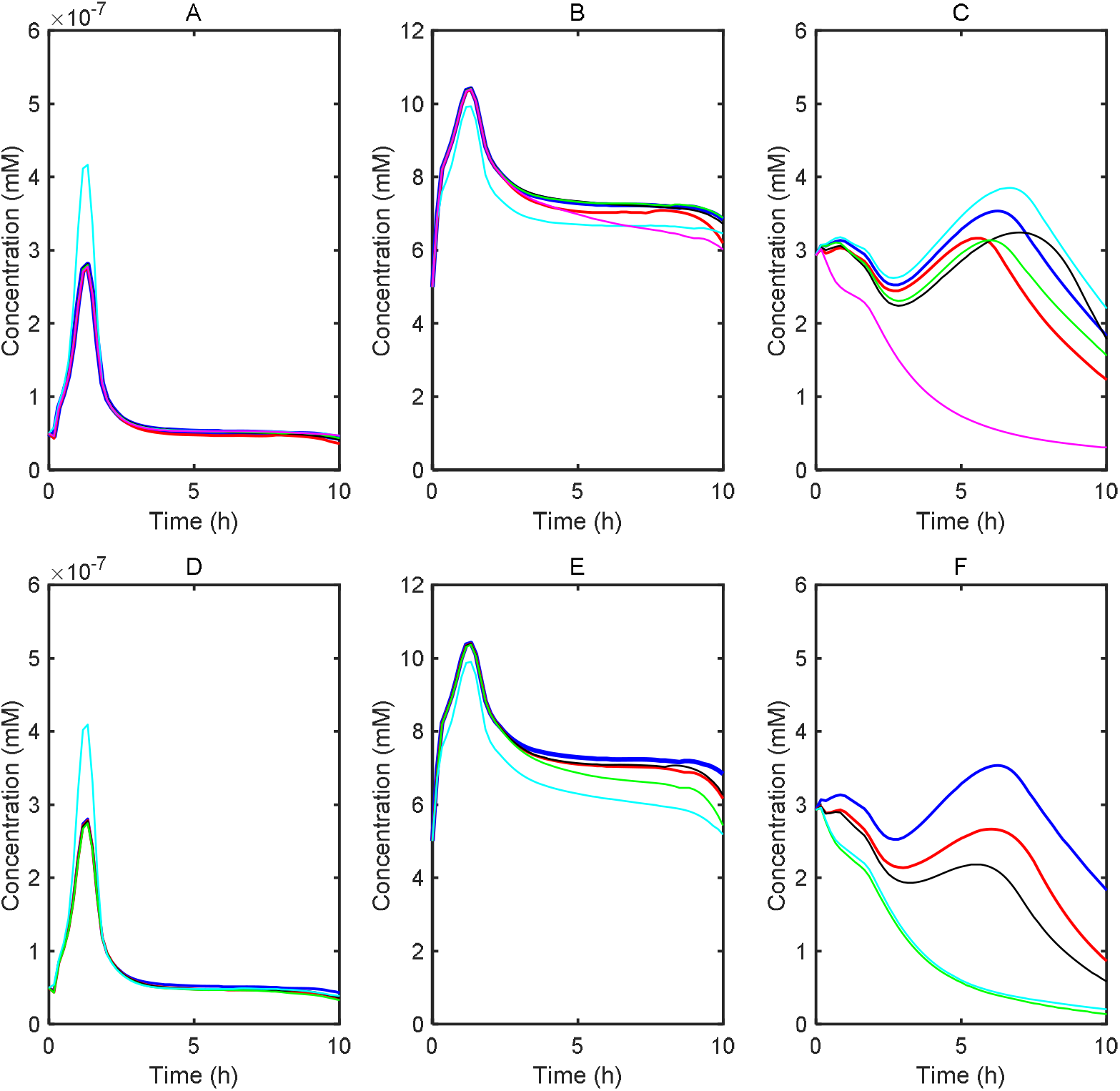
Single medication analysis and prediction of combination therapy for type 2 diabetes by using the virtual metabolic human model. (A-C) Single medication: Time courses of the concentration of plasma insulin (A), plasma glucose (B), and hepatic TG (C) for type 2 diabetes were simulated after an overnight fast and following a single meal of 100g glucose and 33 g TG. The lines indicate the type 2 diabetes model without any medication (blue), LDH inhibition by metformin (red), β-oxidation activation by metformin (black), thiazolidinedione (green), sulfonylurea (cyan), and glycerol kinase inhibitor (magenta). The type 2 diabetes model is the same as the green line of Figure 8DEF. Specifically, metformin reduces Vmax_ldh_Pyr_L to zero or multiplies Vmax_boxid_FFA_L by 2. Thiazolidinedione multiplies Vmax_tgsyn_FFA_A by 2. Sulfonylurea multiples Vmax_inssyn_Glc_B by 2. Glycerol kinase inhibitor reduces Vmax_glyk_Glyc_L to zero. (D-F) Combination therapy: Time courses of the concentration of plasma insulin (D), plasma glucose (E), and hepatic TG (F) for type 2 diabetes were simulated after an overnight fast and following a single meal of 100g glucose and 33 g TG. The lines indicate no medication (blue), metformin therapy (red), the combination therapy of metformin and thiazolidinedione (black), the combination therapy of metformin, thiazolidinedione and glycerol kinase inhibitor (green), and the combination therapy of metformin, thiazolidinedione, glycerol kinase inhibitor, and sulfonylurea (cyan).

### Design of combination therapy

To investigate the feasibility of combination therapy, I employed three medicines (metformin, thiazolidinedione and sulfonylurea) and a glycerol kinase inhibitor. **Figure 7DEF** shows the simulation results of three combination therapies. Expectedly metformin decreased plasma glucose and hepatic TG. The combination of metformin and thiazolidinedione additively decreased hepatic TG; it hardly changed the insulin and glucose dynamics. Addition of the glycerol kinase inhibitor to the above two medicines further decreased plasma glucose and hepatic TG; it hardly changed the insulin concentration. The addition of sulfonylurea to the above three compounds increased plasma insulin to further decrease plasma glucose. The combination of the four compounds hardly changed hepatic TG compared to that of the three compounds, because sulfonylurea has no capability to decrease TG. These simulation results revealed that the combination therapy is effective in reducing plasma glucose and hepatic TG. The combination therapy presented explicitly additive effects, suggesting that the four medicine acted on different reactions.

## Discussion

### Modular model construction

I have constructed a dynamic model that integrated eight organ modules and formulated the whole-body metabolism at the three scales of body, organ, and molecule. The resultant model consists of 202 ordinary differential equations with 217 reaction rates and 1132 kinetic parameter constants. It is the most comprehensive, largest and highly predictive kinetic model of the whole-body metabolism. The divide and conquer strategy at the organ scale allows us to run the individual organs separately and to exchange organs without any extensive rework at other scales. This modular dynamic model accurately reproduced the dynamics of experimental data of insulin, glucose, lactate, FFA, and TG concentrations and transport/exchange fluxes between blood and other organs after a meal and at rest. This model captured key features of human-whole body metabolism: lactate-glucose, alanine-glucose, glycerol-glucose cycles between liver and other organs. The model reproduced the two typical features of DNL suppressing β-oxidation and of glycolysis or PDH reaction decreasing β-oxidation. The former is effective in avoiding their futile cycles; the latter is effective in adjusting the energy balance between glycolysis and β-oxidation. In addition, the model presented a critical switching function between DNL and Bhb synthesis with respect to pyruvate in liver (**Figure 4)**. Pyruvate played a critical role in determining the entry of acetyl-CoA into TCA cycle, DNL or synthesis of FFA from citrate (CIT), and Bhb synthesis. Since the entry reaction of the TCA cycle in mitochondria is presented by oxaloacetate (OAA)+acetyl-CoA->CIT, OAA that comes from pyruvate is essential to drive the TCA reaction. Under a fasted condition a low glucose concentration causes shortage of pyruvate, suppressing the entry reaction of the TCA cycle. In this case, β-oxidation-produced acetyl-CoA is not consumed by the TCA cycle, but utilized for the synthesis of Bhb. On the other hand, at high plasma glucose, abundant pyruvate activates the TCA cycle flux, enhancing CIT production in mitochondria. CIT is transported into cytoplasm, converted into acetyl-CoA to synthesize FFA. In a word, high plasma glucose enhances DNL.

I discuss some discrepancies between experimental data and simulation. This model reproduced the experimental plasma glucose pulse, but did not exactly reproduce the experimental insulin pulse. It was inevitable because the insulin dynamics was tightly coupled with the glucose pulse (**Equation S4**). The insulin pulse was tailed probably due to some hidden mechanisms. The simulated glycogen concentration in liver increased to a peak 2-3 h before experimental peak, and it decreased more rapidly than experimental data. The fast glycogen synthesis may result from the simplification that the model lumps many reactions involved in glycogen synthesis. The fast degradation may be caused by the fact that the model does not include kidney. Kidney is known to release glucose after the meal, which facilitates substantial liver glycogen repletion or suppression of glycogenolysis (Meyer et al., 2002).

### Model validity

The employed experimental data were a collection of different references, i.e., the experimental data were not obtained under the exactly same conditions. For example, the time course data of plasma insulin, plasma glucose, and plasma lactate derived from (Frayn et al., 1993), while the experimental data of their transport fluxes came from (Kim et al., 2007). The proposed model did not insist on the exact fitting, but pursued capturing critical metabolic features. In human metabolism studies, direct measurements of *in vivo* kinetics are very hard and *in vivo* experiments are seriously limited due to ethical issues. Since it is difficult to obtain a sufficient amount of data for kinetic modeling at present and in the near future, we have to manage fragmental, heterogeneous quantitative data, qualitative data, and biological knowledge to build a mathematical model (Maeda et al., 2013; Maeda et al., 2019). The important thing is not to precisely measure or evaluate the exact values of kinetic parameters, but to capture essential functions underlying metabolic networks without insisting on the exact values *in vivo* nor on complexity in detailed biochemistry (Kurata et al., 2005; Kurata et al., 2003).

I derived the rate equations from precise chemical reaction equations (Swainston et al., 2016) when the experimentally validated rate equations are not available. Due to shortage of measured data, I did not uniquely determine the kinetic parameter values. However, existing experimental data including *in vitro* measured kinetic constants, qualitative data and biological information rather constrained the parameter space of the kinetic model. They were feasible enough for constructing kinetic models, for understanding the mechanisms by which the metabolic networks generate physiological functions and robustness, and for performing pathological and medication analysis. The resolution of the model would be appropriate enough for the given experimental data and biological information. It may be no use to build much more detailed mathematical equations, if there is little experimental data to validate them.

### Robustness

Generally, high sensitivity-providing parameters point out critical mechanisms such as ultrasensitivity, positive feedback loop, and bottleneck reactions. The insulin-related kinetic parameters of Km_inssyn_Glc_B, Km_Ins_B_L and Km_Ins_B_M, which were employed by Hill equations with ultrasensitivity (Konig et al., 2012), showed high sensitivity. Such ultrasensitivity makes it possible for insulin to induce the distinct shift between glucose utilization and production. Coenzyme (ATP, GTP, UTP, NADH, NADPH, and FADH2) synthesis-related parameters greatly affected the major pathways of β-oxidation, TCA cycle, and PDH reaction, while previous whole-body models hardly considered nucleotide cofactors (Kim et al., 2007; Palumbo et al., 2018; Xu et al., 2011). NAD^+^ affects the reactions of TCA cycle, β-oxidation, LDH, and glyceraldehyde-3-phosphate dehydrogenase (G3PD). G3PD is the critically responsible enzyme for linking glycolysis to TG synthesis. NADH, which is produced through glycolysis, β-oxidation, and TCA cycle, is essential to generate ATP through oxidative phosphorylation. NADPH, which is produced in the pentose phosphate pathway, is utilized for DNL. UTP, which is essential for glycogen synthesis, is regenerated by nucleotide diphosphate kinase reaction of ATP +UDP-> ADP + UTP. GTP, which is necessary for gluconeogenesis (**Figure 2**), is also regenerated by nucleotide diphosphate kinase reaction of ATP +GDP-> ADP + GTP. Those nucleotide cofactors conjugate multiple cofactor-coupled reactions to form global feedback loops, which is exemplified by glycolysis pathways with TCA cycle and oxidative phosphorylation (Kurata, 2019; Teusink et al., 1998). Glycolysis requires ATP at the initial step (hexokinase (HK), phosphofructokinase (PFK)) (**Figure 2**). An increase in glycolysis flux enhances the TCA cycle with oxidative phosphorylation to further produce ATP, which activates the initial step of glycolysis in a positive feedback manner. This ATP amplification is called turbo-design or self-replenishment cycle (Kurata, 2019; Teusink et al., 1998). It accelerates the glucose utilization (HK, PFK) to reduce plasma glucose. β-oxidation in each organ and Bhb degradation in brain also employ the similar positive feedback loops. Thus, incorporation of nucleotide cofactors is effective in understanding the mechanisms of global metabolic changes.

While the insulin-related parameters including Km_inssyn_Glc_B, Km_Ins_B_L and Km_Ins_B_M played a critical role in changing a set-point (5mM) of plasma glucose concentration, the set-point provided a highly robust property against changes in the other kinetic parameters. The set-point of plasma glucose is dominantly controlled by insulin. To confirm the robustness of the set-point, we boldly set Vdif_glut3_Glc_B_N = 0 to stop the glucose uptake by brain. Surprisingly, the set-point was held (data not shown) by the insulin-based feedback switching between glucose utilization and production in liver. In addition, nucleotide cofactors conjugate multiple cofactor-coupled reactions to generate global negative feedback loops. The negative loops control the balances of NADH redox energy and ATP energy. For example, an increase in NADH suppresses the β-oxidation and PDH reaction due to the decreased NAD^+^, which deactivates the NADH synthesizing-TCA cycle. Excess ATP would promote the synthesis of TG and cholesterol, which decrease ATP.

### Prediction of hidden mechanism

The proposed model suggested a two-phase mechanism of FFA synthesis in liver, which consists of the former malonyl-CoA accumulation phase; the latter malonyl-CoA decreasing phase (**Figure 5**). This separation is caused by the fact that NADPH synthesis in the pentose phosphate pathways is coupled with DNL (**Figure 2**). Malonyl-CoA is a key intermediate metabolite for the FFA synthesis. During the former phase, G6P in liver decreases, which decreases the pentose phosphate pathway flux and NADPH. Malonyl-CoA is accumulated due to shortage of NADPH. During the latter phase, G6P increases, which enhances the pentose phosphate pathway flux and NADPH. The accumulated malonyl-CoA is converted with NADPH into FFA. The two phases correspond to the glucose production and utilization phases. Consequently, the DNL occurs shortly after the glucose pulse.

### Pathological analysis

A computer model has an advantage in analysis of the individual disorders and in understanding some mechanisms by which the individual ones are combined to cause complex pathologies. Metabolic diseases take a matter of years to develop, but short-term simulation is effective in considering some signs of such diseases and in capturing typical pathological features. NAFLD, one of the most common chronic liver disorders worldwide, is the excessive build-up of TG in liver cells. It covers a wide range of liver dysfunctions, ranging from simple steatosis to non-alcoholic steatohepatitis (NASH), liver cirrhosis, and hepatocellular carcinoma (Ashworth et al., 2016; Kitade et al., 2017). NAFLD has strong association with type 2 diabetes, 90% of obese patients with type 2 diabetes have NAFLD. NAFLD with IR progression leads to type 2 diabetes (Perry et al., 2014). I considered that NAFLD is hepatic steatosis with some IRs, while I did not simulate NAFLD due to its broad definition.

The proposed kinetic model was applied to pathological analysis for diabetes (Ashworth et al., 2016; Utzschneider and Kahn, 2006). There are two types of diabetes: type 1 and type 2. In type 1 diabetes, little or no insulin is produced by pancreas. Type 2 diabetes is characterized by steatosis, IR and β cell dysfunction. Of course steatosis is the underlying disease. I investigated the effect of IR and β cell dysfunction on diabetes progression. IR_A alone hardly increased the plasma glucose set-point. The set-point increased in the order of IR_M, IR_L, and β cell dysfunction. The β cell dysfunction is the first major contribution; the IR_L is the second major one; IR_M is the third major one. β cell dysfunction and IR_L induced serious progression. The small difference in the plasma glucose set-point between IR_A and IR_M reflects that skeletal muscle accounts for 80% of the whole insulin-mediated glucose uptake (Saltiel and Kahn, 2001; Stump et al., 2006; Thong et al., 2005). Quantitative understanding of the mechanisms responsible for insulin-controlled glucose uptake in each disorder is useful to evaluate the contribution of each disorder to type 2 diabetes.

The simulation results supported the widely-recognized experimental data that IR_M and IR_A promote hepatic TG synthesis. However, **Figure 6** suggests a paradox or conflict that β cell dysfunction and IR_L, which are known to exacerbate diabetes, recover the steatosis of a major cause of IR. I solve the paradox as follows. When IR occurs, hepatic TG accumulation may be no longer controlled by insulin (Doege et al., 2008; Kim et al., 2001), but be transformed into a chronic or irreversible state through oxidation, ER stress, and inflammation (Ferre and Foufelle, 2010).

### Medication and combination therapy

Medication is a very popular way to treat type 2 diabetes. Out of many medications, I employed the three medicines of metformin, thiazolidinedione, and sulfonylurea and a glycerol kinase inhibitor that act on different reactions. Curative remedies are to reduce plasma glucose and to decrease ectopic TG accumulation. The three compounds of metformin, thiazolidinedione, glycerol kinase inhibitor were predicted to be effective in reduction of plasma glucose or hepatic TG. The model demonstrated a side effect of metformin that increased plasma lactate as lactic acidosis (simulation data not shown), as described elsewhere (Molavi et al., 2007; Pernicova and Korbonits, 2014). Interestingly, the proposed model uncovered that the glycerol kinase inhibitor decreased not only hepatic TG but also plasma glucose. The reduced plasma glucose results from the shortage of GRP. Specifically, the shortage of GRP decreases the backward reaction from GRP to GAP, which results in reduced gluconeogenesis. The glycerol kinase inhibitor is found to be a promising medicine for type 2 diabetes. Out of several medications, activation of β-oxidation, which burns FFA, can be a powerful remedy to cure type 2 diabetes, because it has an advantage in removing not only TG but also its precursors. Metformin acts on β-oxidation through AMPK activation, while its action is marginal. Thus, new medicines should be developed that effectively activate β-oxidation. Sulfonylurea successfully reduced plasma glucose, but increased hepatic TG, suggesting a limitation or side effect of sulfonylurea. The model not only accurately predicted the essential results of medications but also achieved a rational design of combination therapy. The combination therapy of metformin, thiazolidinedione, glycerol kinase inhibitor, and sulfonylurea was found to be a promising therapy to reduce plasma glucose and hepatic TG. The successful combination results from the fact that the four compounds act on different reactions.

### Model limitation

IR can be caused by ectopic TG accumulation. Specifically, a lipid metabolite of diacylglycerol activates protein kinase C isoforms, impairing insulin signaling in organs (Cerf, 2013; Perry et al., 2014). In addition, the ratio of saturated FFA to monounsaturated FFA plays a major role in disease progression (Alkhouri et al., 2009). Saturated FFA leads to inflammation, ER stress and apoptosis. Since the proposed model is neither able to consider DAG nor saturated FFA, more detailed models are necessary that consider the IR-inducing mechanisms. The present model seems a mono-stable model (I have not found any bistability yet in the model), while real disorders are often chronical and irreversible. Actually, accumulated TG may be aggregated and denatured due to inflammation, oxidation and ER stress to cause chronic and irreversible dysfunctions (Perry et al., 2014). Such irreversible disorders should be considered in the next model.

## Methods

### Mathematical model

#### Model overview

A schematic diagram of the whole body is shown in **Figure 1**. The union of all the metabolic networks of the organs is shown in **Figure 2**, where I lumped associated enzyme reactions into chemical reaction equations based on Recon2.2 (Swainston et al., 2016). Each organ network was built by selecting organ-specific reactions from the union network. Details of biochemical reactions in the whole body are summarized in **Text S1**. The whole-body model formulates the metabolic enzyme and transporter reactions with regulation of enzyme activities and hormonal actions for all the organs. A meal of glucose and TG is inputted into blood through the GI tract. The model focuses on insulin hormone (Yugi et al., 2014), assuming that glucagon effect is approximately opposite to insulin. Note that detailed kinetics of glucagon *in vivo* remains to be measured (Kawamori et al., 2009). The model takes account for glucose, lactate, glycerol, alanine, Bhb, TG, FFA and insulin in blood, assuming the perfect mixing that there is no spatial gradient of metabolites, oxygen and carbon dioxide in each organ. In this study alanine represents the gluconeogenic amino acids and Bhb represents ketone bodies. Notably, the nucleotide cofactors of ATP, GTP, UTP, NADH, NADPH, and FADH_2_ are incorporated to reflect realistic metabolic changes. The resultant model consists of 202 ordinary differential equations for metabolites with 217 reaction rates and 1132 kinetic parameter constants, as shown in **Equation S1-S419**, **Table S1**, and **Table S2**. Abbreviations of the kinetic parameter constants are defined in **Table S3**. The differential equations are integrated with ode15s (MATLAB R2019a, The MathWorks, Inc.) to simulate their dynamics.

#### Module decomposition

A divide and conquer strategy is employed for model construction (Karr et al., 2012; Kurata et al., 2007). The whole body is decomposed into blood (B) and eight distinct tissue/organ modules (**Figure 1**): liver (L), skeletal muscle (M) adipose tissue (A), GI tract (G), heart (H), brain (N), pancreas (P), and other tissues (T). Blood acts as the principal transport/exchange medium for metabolites between the different organs. Glucose and TG are inputted to blood through the GI tract, following ingestion of a meal (**Equations S1**, **S2**). Insulin secretion from pancreas is controlled by plasma glucose and FFA concentrations (Yaney and Corkey, 2003) (**Equations S4**, **S5**). Pancreas is simplified as an insulin controller. In liver, many rate equations are derived from several previous works (Ashworth et al., 2016; Berndt et al., 2018a; Berndt et al., 2018b; Kim et al., 2007; Pearson et al., 2014). I lump multiple associated enzyme reactions and simplify signal transduction pathways including glucose-controlled insulin secretion, phosphorylation of enzymes, and gene regulations by carbohydrate responsive element binding protein (ChREBP) and sterol-regulatory element binding protein 1c (SREBP-1c). When no experimental rate equation is available, I derive plain Michaelis-Menten type equations based on chemical reaction equations and then estimate the value of Vmax of rate equations. The parameter estimation is carried by genetic algorithms (Maeda et al., 2013; Maeda et al., 2019) so that the model can reproduce the experimental transport/exchange fluxes at rest between blood and each organ (Kim et al., 2007) without changing the experimental values of the Michaelis and dissociation constants. In the other organs, where little measured kinetic data is available, I estimate Vmax of rate equations so as to reproduce the experimental transport fluxes at rest (Kim et al., 2007), while fixing the experimental values the Michaelis and dissociation constants. I assume that gene expression profiles depend on organs but the employed enzymes are common to all the organs. The other tissue has no specific regulation.

#### Module assembly

I combine all the organ modules. To connect blood to each organ, I define *α_X_* and *β_X_* (*X* = *L,M*, *A, G, H*) as the insulin and glucagon factors for each organ, respectively, which alters the activity of insulin- and glucagon-controlled enzymes (**Equations S7-S16**). Organs of B, N, P, and T do not have the insulin/glucagon factors. I connect the liver module to the blood module. Subsequently, I add the skeletal muscle, adipose tissue, GI, heart and brain modules. The values of the insulin-related kinetic parameters are estimated so that the model can reproduce the experimental time course data of plasma insulin, plasma glucose, plasma lactate (Frayn et al., 1993), plasma FFA, hepatic glycogen (Taylor et al., 1996), and plasma TG (Karpe et al., 1992).

### Dynamic sensitivity analysis

To investigate the robustness of the model, the relative change of metabolite concentrations with respect to a change in a kinetic constant were evaluated. Sensitivity analysis explores some mechanisms by which a system of interest generates robustness and remarkable changes (Masunaga et al., 2017). The dynamic sensitivity of target parameter *y* with respect to a change in specific constant parameter *p* is given by:

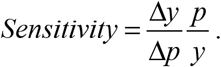

### Pathological analysis

A long-term excess supply of glucose develops steatosis, lipotoxicity, or ectopic TG accumulation by increasing synthesis of ChREBP, a transcription factor whose expression is more exclusively regulated by sugars than insulin (Mandarino et al., 1993, 1996). It also leads to the synthesis of SREBP-1c, which is stimulated by insulin under metabolically normal conditions. In steatosis, the activities of enzymes regarding *de novo* lipogenesis (DNL) (synthesis from acetyl-CoA to TG) and cholesterol synthesis abnormally increase in liver. Steatosis or ectopic TG accumulation can cause metabolic diseases, such as hyperglycemia, hyperlipidemia, hyperinsulinemia, obesity, NAFLD and type 2 diabetes (Friedman, 2002; Guebre-Egziabher et al., 2013; Kitade et al., 2017; Molavi et al., 2007). Chronic exposure to high plasma glucose and obesity, leading to oxidative stress and inflammation, induce changes in the regulation of gene expression that converge on impaired glucose-stimulated insulin secretion (Gilbert and Liu, 2012) and IR. β cell dysfunction in pancreas suppresses glucose-stimulated insulin secretion.

#### Individual disorder decomposition

While complex mechanisms of metabolic diseases may not be exactly defined, I conveniently decomposed the diseases into individual disorders to perform pathological analysis. Diabetes was decomposed into steatosis, β cell dysfunction, and IR. Steatosis is the underlying disease. β Cell dysfunction of pancreas is a major cause of shifting a set-point of plasma glucose concentration (Ashcroft and Rorsman, 2012; Cerf, 2013). IR is further classified with respect to each organ of liver, skeletal muscle, and adipose tissue. IR of each organ is also a major cause. To build a steatosis model, I increased the values of kinetic parameters regarding the DNL and cholesterol synthesis in liver.

#### Diabetes reconstruction

Type 2 diabetes consists of steatosis, β cell dysfunction and IR (Ashcroft and Rorsman, 2012; Saltiel, 2001; Xia et al., 2019). I built three types of type 2 diabetes to analyze the effect of IR on disease progression. To build a type 2 diabetes model, IR_L, IR_M or IR_A was added to the steatosis and β cell dysfunction model. Type 1 diabetes, once known as juvenile diabetes or insulin-dependent diabetes, is the extreme case of β cell dysfunction in which pancreas produces little or no insulin. Differing from the type 2 diseases, type 1 diabetes results from genetic defects or some viruses and it has no cure.

### Medication analysis

I used three types of widely prescribed type 2 diabetes medicines: sulfonylurea, metformin and thiazolidinedione to perform medication analysis. In addition to the three medicines, a glycerol kinase inhibitor was used to inhibit TG synthesis (Seltzer et al., 1986). Sulfonylurea promotes insulin secretion by pancreas (Aquilante, 2010). Metformin is active in the suppression of hepatic gluconeogenesis. (DeFronzo et al 1991; Stumvoll et al 1995) (Molavi et al., 2007). Specifically, it suppresses LDH in liver, accompanied by lactic acidosis (Pernicova and Korbonits, 2014). It also activates AMP-activated protein kinase (AMPK) that regulates energy homeostasis (Fullerton et al., 2013; Pernicova and Korbonits, 2014) or activate β-oxidation. Thiazolidinediones are a family of drugs that have been used in the treatment of type 2 diabetes since the late 1990s (Molavi et al., 2007). The thiazolidinedione derivatives, pioglitazone and rosiglitazone, are synthetic ligands for peroxisome proliferative-activated receptor γ (PPARγ) that improve insulin sensitivity. PPARγ is mainly expressed in adipose tissue; PPARγ agonists promote adipocyte differentiation and promote the FFA uptake and storage in subcutaneous adipose tissue rather than visceral sites (Rasouli et al., 2005).

### Experimental data and subject

The model targets a healthy young adult man with 70kg body weight. A single meal consists of 100 g glucose and 33 g TG. The model employs the experimental transport/exchange fluxes at rest between blood and each organ (Kim et al., 2007) and the experimental time course data of plasma insulin, plasma glucose, plasma lactate (Frayn et al., 1993), plasma FFA, liver glycogen (Taylor et al., 1996), plasma TG (Karpe et al., 1992), and plasma ketone body (Owen et al., 1990).

### Program availability

The program is freely available upon request.

## Supporting information

Supplemental Equation

## Acknowledgments

This work was supported by the Grant-in-Aid for Scientific Research (B) (19H04208) from Japan Society for the Promotion of Science. I am grateful of K. Horimoto, K, Ogata, K. Nasu, H. Shikita and R. Yoshimori performing computer simulation and system analysis. The organ illustrations in Figure 1A are provided by freepik (https://jp.freepik.com) and ac-illust (https://www.ac-illust.com/).

## Author contribution

The first author did everything.

