## Supplemental TextS1 for "Virtual metabolic human dynamic model for pathological analysis and therapy design for diabetes"

### Supplemental Text

#### Methods

##### Homeostasis of metabolites in blood

Glucose and hormone

Plasma glucose in human is controlled with a set-point of 5 mM by hormones of insulin and glucagon [1]. Insulin is the only hormone that decreases the plasma glucose concentration, while multiple glucose increasing hormones are known. Glucagon is a counter partner of insulin. The plasma concentrations of insulin and glucagon directly respond to changes in plasma glucose. The plasma glucose concentration is maintained in a narrow range between a minimum value of 3 mM after prolonged fasting or exercise and a maximum value of 9 mM after a meal [2]. Glucose enters blood in three ways: absorption from the intestine, glycogenolysis in liver, and gluconeogenesis in liver and kidney. After an overnight fast, 95% of glucose production comes from liver [3]. Liver produces glucose through glycogenolysis and gluconeogenesis with almost equal contribution at rest. Lactate, pyruvate, alanine and glycerol are the major gluconeogenic precursors. 50% of glucose at rest is utilized by brain, while skeletal muscle uses 20%. The gastrointestinal (GI) tract consumes only 10% of glucose. The organs except brain use free fatty acid (FFA) as metabolic fuels to save glucose.

Lactate, pyruvate and alanine

Liver and heart primarily consume plasma lactate, while skeletal muscle, adipose tissue and other tissues, including inactive upper body muscles and red blood cells, produce lactate. Pyruvate exchange occurs primarily between skeletal muscle and other tissues. Plasma pyruvate concentration is very small or negligible. Only liver consumes amino acids, especially alanine, for gluconeogenesis, while skeletal muscle is the main source of alanine and the inactive muscle in other tissues is an additional source.

FFA, glycerol and triglyceride (TG)

FFA and glycerol are mainly produced from lipolysis of TG in adipose tissue. Liver uptakes FFA from blood and utilizes FFA as a main fuel. A half of the liver-taken FFA is oxidized; the half is re-esterified into TG [1, 3]. Since adipose tissue lacks glycerol phosphorylase, lipolysis-produced glycerol is not utilized for TG synthesis in adipose tissue. Liver uptakes the glycerol released from adipose tissue and utilize it as a gluconeogenic precursor, i.e., a substrate for TG synthesis.

Ketone body

Ketone bodies including β-hydroxybutyrate (Bhb) are synthesized from acetyl-CoA produced through β-oxidation. Synthesis of ketone bodies are stimulated mainly by glucagon in liver under a fasted condition. Ketone bodies are utilized exclusively by brain.

##### Metabolic reactions of each organ

Liver and pancreas

Liver plays a central role in buffering or controlling plasma glucose. Switching between the glucose utilization (glycolysis and glycogenesis) and glucose production (gluconeogenesis and glycogenolysis) is dependent on the plasma glucose level. The glucose utilization occurs at glucose concentration exceeding a critical threshold value; the glucose production occurs below the critical concentration. Insulin alters the phosphorylation state of multiple key interconvertible enzymes of hexokinase (HK), glycogen synthase (GS), glycogen phosphorylase (GP), phosphofructokinase (PFK), fructose-1,6-bisphosphatase (FBP), pyruvate kinase (PK) and pyruvate dehydrogenase (PDH) to shift a remarkable metabolic state. Liver temporally stores substantial amounts of glucose as glycogen, synthesizes glucose from small carbohydrates, including lactate, pyruvate, glycerol and alanine, and converts excess glucose into FFA. It also synthesizes TG and cholesterol and secretes them into blood. Under a fasted condition, liver synthesizes Bhb from acetyl-CoA as a metabolic fuel for brain. In pancreas β cells serve as a controller of insulin synthesis and release in response to a plasma glucose concentration.

Skeletal muscle and heart

Insulin activates glucose transporter 4 (GLUT4) in skeletal muscle to uptake glucose and to accumulate glycogen. To control substantially glucose uptake rates, a few key enzymes of HK, GS, and PDH are activated [4, 5]. In this study skeletal muscle represents the lean muscles in the lower extremity. Skeletal muscle uptakes FFA as fuels and releases lactate and alanine into blood. Heart consists of specialized muscle cells (cardiomyocytes) and constantly uptakes metabolic fuels, including glucose, lactate, and FFA to generate ATP to maintain contractile function without any fatigue. In contrast to skeletal muscle, GLUT1, which is not controlled by insulin, is dominant. The major metabolic fuel for the heart is FFA.

Adipose tissue and GI tract

Adipose tissues are producers and reservoirs of TG. Plasma TG is degraded by lipase on the adipose tissue surface into FFA and glycerol. FFA enters the adipose cells; glycerol returns to blood. Within adipose tissue, FFA and glycerol-3-P are synthesized into TG. Since adipose tissue lacks glycerol kinase, glycerol-3-P comes just from glucose-derived glycelaldehyde-3-phosphate (GAP). Insulin activates the TG synthesis and lipase reaction, facilitating TG accumulation. GI tract includes the splanchnic region (stomach, spleen, intestines) except liver. It utilizes glucose, accumulates TG, and releases FFA and glycerol into blood.

Brain and other tissues

Brain constantly takes only glucose and Bhb as metabolic fuels, neither utilizes FFA nor TG, because the blood-brain barrier prevents such large-size molecules from entering brain. It has Bhb degradation pathways to degrade Bhb into acetyl-CoA for energy. The other tissue compartment includes kidney, upper extremity muscles, and the rest of tissues.
