## Supplemental Equation for "Virtual metabolic human dynamic model for pathological analysis and therapy design for diabetes"

### Supplemental Equations

#### Dietary input flux

$$v_{meal}^{Glc_B} = Glc_B^{meal} \frac{t}{T_{delay}^{Glc_B\,2}} \exp(-\frac{t^2}{2T_{delay}^{Glc_B\,2}}) \quad \text{(S1)}$$

$$v_{meal}^{TG_B} = TG_B^{meal} \frac{t}{T_{delay}^{TG_B\,2}} \exp(-\frac{t^2}{2T_{delay}^{TG_B\,2}}) \quad \text{(S2)}$$

where $t$ is time. [1]

#### Insulin flux

The insulin synthesis rate is described by Hill equation for plasma glucose [1, 2].

$$v_{inssyn}^{B} = k_{inssyn}^{B} \quad \text{(S3)}$$

$$v_{inssyn}^{Glc_B} = Vmax_{inssyn}^{Glc_B} \frac{Glc_B^{\,n_{inssyn}^{Glc_B}}}{K_{minssyn}^{Glc_B}{}^{n_{inssyn}^{Glc_B}} + Glc_B^{\,n_{inssyn}^{Glc_B}}} \quad \text{(S4)}$$

$$v_{inssyn}^{FFA_B} = k_{inssyn}^{FFA_B} FFA_B \quad \text{(S5)}$$

$$v_{insdeg}^{Ins_B} = k_{insdeg}^{Ins_B} Ins_B \quad \text{(S6)}$$

#### Insulin and glucagon activities in organs

Insulin-regulated enzyme activity factor in liver is defined as $\alpha_L$, which is determined by plasma insulin concentration, depending on each organ.

$$\alpha_L = \alpha_L^{Base} + \alpha_L^{Band} \frac{Ins_B^{\,n^{Ins_{BL}}}}{Km^{Ins_{BL}\,n^{Ins_{BL}}} + Ins_B^{\,n^{Ins_{BL}}}} \quad \text{(S7)}$$

$$\alpha_M = \alpha_M^{Base} + \alpha_M^{Band} \frac{Ins_B^{\,n^{Ins_{BM}}}}{Km^{Ins_{BM}\,n^{Ins_{BM}}} + Ins_B^{\,n^{Ins_{BM}}}} \quad \text{(S8)}$$

$$\alpha_A = \alpha_A^{Base} + \alpha_A^{Band} \frac{Ins_B^{\,n^{Ins_{BA}}}}{Km^{Ins_{BA}\,n^{Ins_{BA}}} + Ins_B^{\,n^{Ins_{BA}}}} \quad \text{(S9)}$$

$$\alpha_G = \alpha_G^{Base} + \alpha_G^{Band} \frac{Ins_B^{\,n^{Ins_{BG}}}}{Km^{Ins_{BG}\,n^{Ins_{BG}}} + Ins_B^{\,n^{Ins_{BG}}}} \quad \text{(S10)}$$

$$\alpha_H = \alpha_H^{Base} + \alpha_H^{Band} \frac{Ins_B^{\,n^{Ins_{BH}}}}{Km^{Ins_{BH}\,n^{Ins_{BH}}} + Ins_B^{\,n^{Ins_{BH}}}} \quad \text{(S11)}$$

Glucagon-regulated enzyme activity factor is given by

$$\beta_L = 1 - \alpha_L \quad \text{(S12)}$$

$$\beta_M = 1 - \alpha_M \quad \text{(S13)}$$

$$\beta_A = 1 - \alpha_A \quad \text{(S14)}$$

$$\beta_G = 1 - \alpha_G \quad \text{(S15)}$$

$$\beta_H = 1 - \alpha_H \quad \text{(S16)}$$

Subscripts: *L*, *M*, *A*, *G*, *H*, *N* and *T* indicate liver, skeletal muscle, adipose tissue, GI, heart, brain, and other tissue, respectively.

**Flux in liver**

Glucose transporter (GLUT2) [3-5]

$$v_{glut2}^{Glc_{BL}} = Vdif_{glut2}^{Glc_{BL}} \frac{Glc_B - Glc_L}{1 + \frac{Glc_B}{Kdif_{glut2}^{Glc_{BL}}} + \frac{Glc_L}{Kdif_{glut2}^{Glc_L}}} \quad \text{(S17)}$$

Hexokinase (HK) [3]

$$v_{hk}^{Glc_L} = \alpha_L \cdot Vmax_{hk}^{Glc_L} \frac{Glc_L}{Glc_L + Km_{hk}^{Glc_L}(1 + \frac{G6p_L}{Ki_{hk}^{G6p_L}})} \cdot \frac{Atp_L}{Atp_L + Km_{hk}^{Atp_L}(1 + \frac{G6p_L}{Ki_{hk}^{Atp_L}})} \quad \text{(S18)}$$

Pentose phosphate pathway (PPP)

$$v_{ppp}^{G6p_X} = Vmax_{g6pase}^{G6p_L} \frac{G6p_L}{G6p_L + Km_{ppp}^{G6p_L}} \frac{Nadp_L}{Nadp_L + Km_{ppp}^{Nadp_L}} \quad \text{(S19)}$$

G6Pase [4]

$$v_{g6pase}^{G6p_L} = Vmax_{g6pase}^{G6p_L} \frac{G6p_L}{G6p_L + Km_{g6pase}^{G6p_L}} \quad \text{(S20)}$$

Glycogen synthase (GS) [4]

$$v_{gs}^{G6p_X} = \alpha_L \cdot Vmax_{gs}^{G6p_L} \frac{G6p_L^{n_{gs}^{G6p_L}}}{G6p_X^{n_{gs}^{G6p_X}} + Km_{gs}^{G6p_X\, n_{gs}^{G6p_X}}} \cdot \frac{(Glygn_L^{\max} - Glygn_L)}{(Glygn_L^{\max} - Glygn_L) + Km_{gs}^{Glygn_L}} \cdot \frac{Utp_L}{Utp_L + Km_{gs}^{Utp_L}} \quad \text{(S21)}$$

$Glygn_L^{\max}$ denotes the maximum amount of glycogen that liver can store [1].

Glycogen degradation (GD) by glycogen phosphorylase (GP) [4]

$$v_{gd}^{Glygn_L} = \beta_L \cdot Vmax_{gd}^{Glygn_L} \frac{Glygn_L}{Glygn_L + Km_{gd}^{Glygn_L}} \frac{Phos_L}{Phos_L + Km_{gd}^{Phos_L}} \quad \text{(S22)}$$

Phosphofructokinase (PFK) [4]

$$v_{pfk}^{G6p_L} = \alpha_L \cdot Vmax_{pfk}^{G6p_L} \frac{G6p_L}{G6p_L + Km_{pfk}^{G6p_L}} \cdot \frac{Atp_L}{Atp_L + Km_{pfk}^{Atp_L}} \cdot \frac{Ki_{pfk}^{Atp_L}}{Atp_L + Ki_{pfk}^{Atp_L}} \cdot \frac{Adp_L}{Adp_L + Km_{pfk}^{Adp_L}} \cdot (1 - b_{pfk}^{Gap_L} \frac{Gap_L}{Gap_L + Ki_{pfk}^{Gap_L}}) \quad \text{(S23)}$$

Fructose-1,6-bisphosphatase (FBP) [4]

$$v_{fbp}^{Gap_L} = \beta_L \cdot Vmax_{fbp}^{Gap_L} \frac{Gap_L}{Gap_L + Km_{fbp}^{Gap_L}} \quad \text{(S24)}$$

Pyruvate kinase (PK) [4]

$$v_{pk}^{Gap_L} = \alpha_L \cdot Vmax_{pk}^{Gap_L} \cdot \frac{Gap_L}{Gap_L + Km_{pk}^{Gap_L}} \cdot (1 - b_{pk}^{Accoa_{LM}} \frac{Accoa_{LM}}{Accoa_{LM} + Ki_{pk}^{Accoa_{LM}}}) \cdot \frac{Adp_L}{Adp_L + Km_{pk}^{Adp_L}} \quad \text{(S25)}$$

Phosphoenolpyruvate carboxykinase (PEPCK) [4]

$$v_{pepck}^{Pyr_L} = \beta_L \cdot Vmax_{pepck}^{Pyr_L} \cdot \frac{Pyr_L}{Pyr_L + Km_{pepck}^{Pyr_X}} \cdot \frac{Atp_L}{Atp_L + Km_{pepck}^{Atp_L}} \cdot \frac{Gtp_L}{Gtp_L + Km_{pepck}^{Gtp_L}} \quad \text{(S26)}$$

Pyruvate transporter (PYRT) [3]

$$v_{pyrt}^{Pyr_{BL}} = Vdif_{pyrt}^{Pyr_{BL}} \frac{Pyr_B - Pyr_L}{1 + \frac{Pyr_B}{Kdif_{pyrt}^{Pyr_{BL}}} + \frac{Pyr_L}{Kdif_{pyrt}^{Pyr_L}}} \quad \text{(S27)}$$

Lactate transporter (LACT) [3]

$$v_{lact}^{Lac_{BL}} = Vdif_{lact}^{Lac_{BL}} \frac{Lac_B - Lac_L}{1 + \frac{Lac_B}{Kdif_{lact}^{Lac_{BL}}} + \frac{Lac_L}{Kdif_{lact}^{Lac_L}}} \quad \text{(S28)}$$

Lactate dehydrogenase (LDH) [3]

$$v_{ldh}^{Pyr_L} = Vmax_{ldh}^{Pyr_L} \cdot \frac{Pyr_L \cdot Nadh_L - Lac_L \cdot Nad_L / Keq_{ldh}^{Lac_L}}{(1 + \frac{Pyr_L}{Km_{ldh}^{Pyr_L}}) \cdot (1 + \frac{Nadh_L}{Km_{ldh}^{Nadh_L}}) + (1 + \frac{Lac_L}{Km_{ldh}^{Lac_L}}) \cdot (1 + \frac{Nad_L}{Km_{ldh}^{Nad_L}}) - 1} \quad \text{(S29)}$$

Alanine transporter (ALAT)

$$v_{alat}^{Ala_{BL}} = Vdif_{alat}^{Ala_{BL}} \frac{Ala_B - Ala_L}{1 + \frac{Ala_B}{Kdif_{alat}^{Ala_B}} + \frac{Ala_L}{Kdif_{alat}^{Ala_L}}} \quad \text{(S30)}$$

Alanine transaminase (ALAT)

$$v_{alata}^{Pyr_L} = Vmax_{alata}^{Pyr_L} \cdot \frac{Pyr_L - Ala_L / Keq_{alata}^{Ala_L}}{(1 + \frac{Pyr_L}{Km_{alata}^{Pyr_L}}) + (1 + \frac{Ala_L}{Km_{alata}^{Ala_L}}) - 1} \quad \text{(S31)}$$

Pyruvate dehydrogenase (PDH) [4]

$$v_{pdh}^{Pyr_L} = \alpha_L \cdot Vmax_{pdh}^{Pyr_L} \cdot \frac{Pyr_L}{Pyr_L + Km_{pdh}^{Pyr_L}} \cdot \frac{Nad_L}{Nad_L + Km_{pdh}^{Nad_L}} \cdot \frac{Km_{pdh}^{Accoa_{LM}}}{Accoa_{LM} + Km_{pdh}^{Accoa_{LM}}} \quad (S32)$$

TCA cycle (TCA)

$$v_{tca}^{Accoa_{LM}} = Vmax_{tca}^{Accoa_{LM}} \cdot \frac{Accoa_{LM}}{Accoa_{LM} + Km_{tca}^{Accoa_{LM}}} \cdot \frac{Adp_L}{Adp_L + Km_{tca}^{Adp_L}} \cdot \frac{Nad_L}{Nad_L + Km_{tca}^{Nad_L}} \cdot \frac{Fad_L}{Fad_L + Km_{tca}^{Fad_L}} \cdot \frac{Phos_L}{Phos_L + Km_{tca}^{Phos_L}} \cdot \frac{Pyr_L}{Pyr_L + Km_{tca}^{Pyr_L}} \quad (S33)$$

FFA transporter (FFAT) [4]

$$v_{ffat}^{FFA_{BL}} = v_{ffat\_d}^{FFA_{BL}} + v_{ffat\_a}^{FFA_{BL}} \quad (S34)$$

$$v_{ffat\_d}^{FFA_{BL}} = Vdif_{ffat}^{FFA_{BL}} \frac{FFA_B - FFA_L}{1 + \frac{FFA_B}{Kdif_{ffat}^{FFA_{BL}}} + \frac{FFA_L}{Kdif_{ffat}^{FFA_L}}} \quad (S35)$$

$$v_{ffat\_a}^{FFA_{BL}} = Vmax_{ffat}^{FFA_{BL}} \frac{FFA_B}{FFA_B + Km_{ffat}^{FFA_{BL}}} \quad (S36)$$

TG synthesis (TGSYN) [4]

$$v_{tgsyn}^{FFA_L} = \alpha_L \cdot Vmax_{tgsyn}^{FFA_L} \frac{FFA_L}{FFA_L + Km_{tgsyn}^{FFA_L}} \cdot \frac{Glycp_L}{Glycp_L + Km_{tgsyn}^{Glycp_L}} \cdot \frac{Atp_L}{Atp_L + Km_{tgsyn}^{Atp_L}} \quad (S37)$$

TG degradation (TGDEG) [4]

$$v_{tgdeg}^{TG_L} = \beta_L Vmax_{tgdeg}^{TG_L} \frac{TG_L}{TG_L + Km_{tgdeg}^{TG_L}} \quad (S38)$$

Glycerol transporter (GLYCT) [4]

$$v_{glyct}^{Glyc_{BL}} = Vdif_{glyct}^{Glyc_{BL}} \frac{Glyc_B - Glyc_L}{1 + \frac{Glyc_B}{Kdif_{glyct}^{Glyc_{BL}}} + \frac{Glyc_L}{Kdif_{glyct}^{Glyc_L}}} \quad (S39)$$

Glycerol kinase (GLYK) [6]

$$v_{glyk}^{Glyc_L} = Vmax_{glyk}^{Glyc_L} \frac{Glyc_L}{Glyc_L + Km_{glyk}^{Glyc_L}(1 + \frac{Gap_L}{Ki_{glyk}^{Gap_L}})} \frac{Atp_L}{Atp_L + Km_{glyk}^{Atp_L}} \quad (S40)$$

TG transporter (TGT) [4]

$$v_{tgt}^{TG_{BL}} = v_{tgt\_a}^{TG_L} + v_{tgt\_d}^{TG_L} \quad (S41)$$

$$v_{tgt\_d}^{TG_{BL}} = Vdif_{tgt}^{TG_{BL}} \frac{TG_B - TG_L \big/ Keq_{tgt}^{TG_L}}{TG_B + Kdif_{tgt}^{TG_{BL}} + \frac{TG_L}{Keq_{tgt}^{TG_L}}} \tag{S42}$$

$$v_{tgt\_a}^{TG_{BL}} = -\beta_L \cdot Vmax_{tgt}^{TG_L} \frac{TG_L}{TG_L + Km_{tgt}^{TG_L}} \tag{S43}$$

Glycerol-3-phosphate dehydrogenase (G3PD) [6]

$$v_{g3pd}^{Gap_L} = Vmax_{g3pd}^{Gap_L} \frac{Gap_L Nadh_L - \frac{Glycp_L}{Keq_{g3pd}^{Glycp_L}} Nad_L}{(1+\frac{Gap_L}{Km_{g3pd}^{Gap_L}})(1+\frac{Nadh_L}{Km_{g3pd}^{Nadh_L}}) + (1+\frac{Glycp_L}{Km_{g3pd}^{Glycp_L}})(1+\frac{Nad_L}{Km_{g3pd}^{Nad_L}}) - 1} \tag{S44}$$

β-oxidation (BOXID) [4]

$$\begin{aligned} v_{boxid}^{FFA_L} = Vmax_{boxid}^{FFA_L} & \frac{FFA_L}{FFA_L + Km_{boxid}^{FFA_L}} \cdot \frac{Atp_L}{Atp_L + Km_{boxid}^{Atp_L}} \cdot \frac{Nad_L}{Nad_L + Km_{boxid}^{Nad_L}} \cdot \frac{Fad_L}{Fad_L + Km_{boxid}^{Fad_L}} \\ & \cdot \frac{Ki_{boxid}^{Accoa_{LM}}}{Accoa_{LM} + Ki_{boxid}^{Accoa_{LM}}} \cdot \frac{Ki_{boxid}^{Malcoa_L}}{Malcoa_L + Ki_{boxid}^{Malcoa_L}} \end{aligned} \tag{S45}$$

AcCoA release into cytoplasm for FFA synthesis (ACCOAT)

$$v_{accoat}^{Accoa_{LMLC}} = Vmax_{accoat}^{Accoa_{LMLC}} \frac{Accoa_{LM}}{Accoa_{LM} + Km_{accoat}^{Accoa_{LM}}} \cdot \frac{Atp_L}{Atp_L + Km_{accoat}^{Atp_L}} \cdot \frac{Pyr_L}{Pyr_L + Km_{accoat}^{Pyr_L}} \tag{S46}$$

β-hydroxybutyrate synthesis (BHBSYN)

$$v_{bhbsyn}^{Accoa_{LM}} = \beta_L \cdot Vmax_{bhbsyn}^{Accoa_{LM}} \frac{Accoa_{LM}}{Accoa_{LM} + Km_{bhbsyn}^{Accoa_{LM}}} \frac{Nadh_L}{Nadh_L + Km_{bhbsyn}^{Nadh_L}} \cdot \frac{{Ki_{bhbsyn}^{Pyr_L}}^{n_{bhbsyn}^{Pyr_L}}}{{Pyr_L}^{n_{bhbsyn}^{Pyr_L}} + {Ki_{bhbsyn}^{Pyr_L}}^{n_{bhbsyn}^{Pyr_L}}} \tag{S47}$$

β-hydroxybutyrate transporter (BHBT)

$$v_{bhbt}^{Bhb_{BL}} = Vdif_{bhbt}^{Bhb_{BL}} \frac{Bhb_B - Bhb_L}{1 + \frac{Bhb_B}{Kdif_{bhbt}^{Bhb_{BL}}} + \frac{Bhb_L}{Kdif_{bhbt}^{Bhb_L}}} \tag{S48}$$

Lipogenesis 1 (LIPOG1)

$$v_{lipog1}^{Accoa_{LC}} = Vmax_{lipog1}^{Accoa_{LC}} \frac{Accoa_{LC}}{Accoa_{LC} + Km_{lipog1}^{Accoa_{LC}}} \cdot \frac{Atp_L}{Atp_L + Km_{lipog1}^{Atp_L}} \tag{S49}$$

Lipogenesis 2 (LIPOG2)

$$v_{lipog2}^{Malcoa_L} = Vmax_{lipog2}^{Malcoa_L} \frac{Malcoa_L}{Malcoa_L + Km_{lipog2}^{Malcoa_L}} \cdot \frac{Adp_L}{Adp_L + Km_{lipog2}^{Adp_L}} \cdot \frac{Nadph_L}{Nadph_L + Km_{lipog2}^{Nadph_L}} \quad \text{(S50)}$$

Cholesterol synthesis 1 (CHOLSYN1)

$$v_{cholsyn1}^{Accoa_{LC}} = Vmax_{cholsyn1}^{Accoa_{LC}} \frac{Accoa_{LC}}{Accoa_{LC} + Km_{cholsyn1}^{Accoa_{LC}}} \quad \text{(S51)}$$

Cholesterol synthesis 2 (CHOLSYN2)

$$v_{cholsyn2}^{Hmgcoa_L} = Vmax_{cholsyn2}^{Hmgcoa_L} \frac{Hmgcoa_L}{Hmgcoa_L + Km_{cholsyn2}^{Hmgcoa_L}} \cdot \frac{Atp_L}{Atp_L + Km_{cholsyn2}^{Atp_L}} \cdot \frac{Fadh_L}{Fadh_L + Km_{cholsyn2}^{Fadh_L}} \cdot \frac{Nadph_L}{Nadph_L + Km_{cholsyn2}^{Nadph_L}} \quad \text{(S52)}$$

Cholesterol transporter (CHOLT)

$$v_{cholt}^{Chol_{BL}} = -Vmax_{cholt}^{Chol_{BL}} \frac{Chol_L}{Chol_L + Km_{cholt}^{Chol_L}} \quad \text{(S53)}$$

ATP synthesis from FADH (ATPSYNF)

$$v_{atpsynf}^{Fadh_L} = Vmax_{atpsynf}^{Fadh_L} \frac{Fadh_L}{Fadh_L + Km_{atpsynf}^{Fadh_L}} \cdot \frac{Adp_L}{Adp_L + Km_{atpsynf}^{Adp_L}} \quad \text{(S54)}$$

ATP synthesis from NADH (ATPSYNN)

$$v_{atpsynn}^{Nadh_L} = Vmax_{atpsynn}^{Nadh_L} \frac{Nadh_L}{Nadh_L + Km_{atpsynn}^{Nadh_L}} \cdot \frac{Adp_L}{Adp_L + Km_{atpsynn}^{Adp_L}} \quad \text{(S55)}$$

ATP utilization (ATPUSE)

$$v_{atpuse}^{Atp_L} = Vmax_{atpuse}^{Atp_L} \frac{Atp_L}{Atp_L + Km_{atpuse}^{Atp_L}} \quad \text{(S56)}$$

Adenosine kinase (AMP regeneration into ADP (AMPREG)) [4]

$$v_{ampreg}^{Amp_L} = Vmax_{ampreg}^{Amp_L} \left( \frac{Amp_L}{Amp_L + Km_{ampreg}^{Amp_L}} \frac{Atp_L}{Atp_L + Km_{ampreg}^{Atp_L}} - \frac{Adp_L}{Adp_L + Km_{ampreg}^{Adp_L}} \frac{Adp_L}{Adp_L + Km_{ampreg}^{Adp_L}} \right) \quad \text{(S57)}$$

Guanosine diphosphate kinase (GDP regeneration to GTP (GDPREG) [3, 6]

$$v_{gdpreg}^{Gdp_X} = Vmax_{gdpreg}^{Gdp_L} \frac{Gdp_L Atp_L - \frac{Gtp_L Adp_L}{Keq_{gdpreg}^{Gtp_L}}}{(1+\frac{Gdp_L}{Km_{gdpreg}^{Gdp_L}})(1+\frac{Atp_L}{Km_{gdpreg}^{Atp_L}}) + (1+\frac{Gtp_L}{Km_{gdpreg}^{Gtp_L}})(1+\frac{Adp_L}{Km_{gdpreg}^{Adp_L}}) - 1} \quad \text{(S58)}$$

Uridine diphosphate kinase (UDP regeneration to UDP (UDPREG)) Berndt, 2018 #28}[6]

$$v_{udpreg}^{Udp_X} = Vmax_{udpreg}^{Udp_X} \frac{Udp_L Atp_L - \frac{Utp_L Adp_L}{Keq_{udpreg}^{Utp_L}}}{(1+\frac{Udp_L}{Km_{udpreg}^{Udp_L}})(1+\frac{Atp_L}{Km_{udpreg}^{Atp_L}}) + (1+\frac{Utp_L}{Km_{udpreg}^{Utp_L}})(1+\frac{Adp_L}{Km_{udpreg}^{Adp_L}}) - 1} \quad \text{(S59)}$$

NADH kinase (NADHK)

$$v_{nadhk}^{Nadh_L} = Vmax_{nadhk}^{Nadh_L} \frac{Nadh_L}{Nadh_L + Km_{nadhk}^{Nadh_L}} \frac{Atp_L}{Atp_L + Km_{nadhk}^{Atp_L}} \quad \text{(S60)}$$

UTP utilization (UTPUSE)

$$v_{utpuse}^{Utp_L} = Vmax_{utpuse}^{Utp_L} \frac{Utp_L}{Utp_L + Km_{utpuse}^{Utp_L}} \quad \text{(S61)}$$

GTP utilization (GTPUSE)

$$v_{gtpuse}^{Gtp_L} = Vmax_{gtpuse}^{Gtp_L} \frac{Gtp_L}{Gtp_L + Km_{gtpuse}^{Gtp_L}} \quad \text{(S62)}$$

NADH utilization (NADHUSE)

$$v_{nadhuse}^{Nadh_L} = Vmax_{nadhuse}^{Nadh_L} \frac{Nadh_L}{Nadh_L + Km_{nadhuse}^{Nadh_L}} \quad \text{(S63)}$$

NADPH utilization (NADPHUSE)

$$v_{nadphuse}^{Nadph_L} = Vmax_{nadphuse}^{Nadph_L} \frac{Nadph_L}{Nadph_L + Km_{nadphuse}^{Nadph_L}} \quad \text{(S64)}$$

FADH utilization (FADHUSE)

$$v_{fadhuse}^{Fadh_L} = Vmax_{fadhuse}^{Fadh_L} \frac{Fadh_L}{Fadh_L + Km_{fadhuse}^{Fadh_L}} \quad \text{(S65)}$$

Creatine kinase (CK) [7]

$$v_{ck}^{Cre_L} = Vmax_{ck}^{Cre_L} \cdot \frac{Cre_L \cdot Atp_L - Crep_L \cdot Adp_L / Keq_{ck}^{Crep_L}}{(1+\frac{Cre_L}{Km_{ck}^{Cre_L}})\cdot(1+\frac{Atp_L}{Km_{ck}^{Atp_L}})+(1+\frac{Crep_L}{Km_{ck}^{Crep_L}})\cdot(1+\frac{Adp_L}{Km_{ck}^{Adp_L}})-1} \tag{S66}$$

**Flux in skeletal muscle**

Glucose transporter (GLUT4)

$$v_{glut4}^{Glc_{BM}} = \alpha_M \cdot Vdif_{glut4}^{Glc_{BM}} \frac{Glc_B - Glc_M}{1+\frac{Glc_B}{Kdif_{glut4}^{Glc_{BM}}}+\frac{Glc_M}{Kdif_{glut4}^{Glc_M}}} \tag{S67}$$

Hexokinase (HK)

$$v_{hk}^{Glc_M} = \alpha_M \cdot Vmax_{hk}^{Glc_M} \frac{Glc_M}{Glc_M + Km_{hk}^{Glc_M}(1+\frac{G6p_M}{Ki_{hk}^{G6p_M}})} \frac{Atp_M}{Atp_M + Km_{hk}^{Atp_M}(1+\frac{G6p_M}{Ki_{hk}^{Atp_M}})} \tag{S68}$$

Glycogen synthase (GS)

$$v_{gs}^{G6p_M} = \alpha_M \cdot Vmax_{gs}^{G6p_M} \frac{G6p_M^{\;n_{gs}^{G6p_M}}}{G6p_M^{\;n_{gs}^{G6p_M}} + Km_{gs}^{G6p_M\; n_{gs}^{G6p_M}}} \frac{(Glygn_M^{\max} - Glygn_M)}{(Glygn_M^{\max} - Glygn_M) + Km_{gs}^{Glygn_L}} \frac{Utp_M}{Utp_M + Km_{gs}^{Utp_M}} \tag{S69}$$

$Glygn_M^{\max}$ denotes the maximum amount of glycogen that skeletal muscle can store.

Glycogen degradation (GD) by glycogen phosphorylase (GP)

$$v_{gd}^{Glygn_M} = Vmax_{gd}^{Glygn_M} \frac{Glygn_M}{Glygn_M + Km_{gd}^{Glygn_M}} \frac{Phos_M}{Phos_M + Km_{gd}^{Phos_M}} \tag{S70}$$

Phosphofructokinase (PFK)

$$v_{pfk}^{G6p_M} = \alpha_M \cdot Vmax_{pfk}^{G6p_M} \frac{G6p_M}{G6p_M + Km_{pfk}^{G6p_M}} \frac{Atp_M}{Atp_M + Km_{pfk}^{Atp_M}} \cdot \frac{Ki_{pfk}^{Atp_M}}{Atp_M + Ki_{pfk}^{Atp_M}} \frac{Adp_M}{Adp_M + Km_{pfk}^{Adp_M}} (1 - b_{pfk}^{Gap_M} \frac{Gap_M}{Gap_M + Ki_{pfk}^{Gap_M}}) \tag{S71}$$

Pyruvate kinase (PK)

$$v_{pk}^{Gap_M} = \alpha_M \cdot Vmax_{pk}^{Gap_M} \frac{Gap_M}{Gap_M + Km_{pk}^{Gap_M}} (1 - b_{pk}^{Accoa_{MM}} \frac{Accoa_{MM}}{Accoa_{MM} + Ki_{pk}^{Accoa_{MM}}}) \frac{Adp_M}{Adp_M + Km_{pk}^{Adp_M}} \tag{S72}$$

Pyruvate transporter (PYRT)

$$v_{pyrt}^{Pyr_{BM}} = Vdif_{pyrt}^{Pyr_{BM}} \frac{Pyr_B - Pyr_M}{1+\frac{Pyr_B}{Kdif_{pyrt}^{Pyr_{BM}}}+\frac{Pyr_M}{Kdif_{pyrt}^{Pyr_M}}} \tag{S73}$$

Lactate transporter (LACT)

$$v_{lact}^{Lac_{BM}} = Vdif_{lact}^{Lac_{BM}} \frac{Lac_B - Lac_M}{1 + \frac{Lac_B}{Kdif_{lact}^{Lac_{BM}}} + \frac{Lac_M}{Kdif_{lact}^{Lac_M}}} \tag{S74}$$

Lactate dehydrogenase (LDH)

$$v_{ldh}^{Pyr_M} = Vmax_{ldh}^{Pyr_M} \frac{Pyr_M \cdot Nadh_M - Lac_M \cdot Nad_M / Keq_{ldh}^{Lac_M}}{(1 + \frac{Pyr_M}{Km_{ldh}^{Pyr_M}})(1 + \frac{Nadh_M}{Km_{ldh}^{Nadh_M}}) + (1 + \frac{Lac_M}{Km_{ldh}^{Lac_M}})(1 + \frac{Nad_M}{Km_{ldh}^{Nad_M}}) - 1} \tag{S75}$$

Alanine transporter (ALAT)

$$v_{alat}^{Ala_{BM}} = Vdif_{alat}^{Ala_{BM}} \frac{Ala_B - Ala_M}{1 + \frac{Ala_B}{Kdif_{alat}^{Ala_{BM}}} + \frac{Ala_M}{Kdif_{alat}^{Ala_M}}} \tag{S76}$$

Alanine transaminase (ALAT)

$$v_{alata}^{Pyr_M} = Vmax_{alata}^{Pyr_M} \cdot \frac{Pyr_M - Ala_M / Keq_{alata}^{Ala_M}}{(1 + \frac{Pyr_M}{Km_{alata}^{Pyr_M}}) + (1 + \frac{Ala_M}{Km_{alata}^{Ala_M}}) - 1} \tag{S77}$$

Pyruvate dehydrogenase (PDH)

$$v_{pdh}^{Pyr_M} = \alpha_M \cdot Vmax_{pdh}^{Pyr_M} \frac{Pyr_M}{Pyr_M + Km_{pdh}^{Pyr_M}} \frac{Nad_M}{Nad_M + Km_{pdh}^{Nad_M}} \frac{Km_{pdh}^{Accoa_{MM}}}{Accoa_{MM} + Km_{pdh}^{Accoa_{MM}}} \tag{S78}$$

TCA cycle (TCA)

$$v_{tca}^{Accoa_{MM}} = Vmax_{tca}^{Accoa_{MM}} \frac{Accoa_{MM}}{Accoa_{MM} + Km_{tca}^{Accoa_{MM}}} \frac{Adp_M}{Adp_M + Km_{tca}^{Adp_M}} \frac{Nad_M}{Nad_M + Km_{tca}^{Nad_M}} \frac{Fad_M}{Fad_M + Km_{tca}^{Fad_M}} \frac{Phos_M}{Phos_M + Km_{tca}^{Phos_M}} \frac{Pyr_M}{Pyr_M + Km_{tca}^{Pyr_M}} \tag{S79}$$

FFA transporter (FFAT)

$$v_{ffat}^{FFA_{BM}} = v_{ffat\_d}^{FFA_{BM}} + v_{ffat\_a}^{FFA_{BM}} \tag{S80}$$

$$v_{ffat\_d}^{FFA_{BM}} = Vdif_{ffat}^{FFA_{BM}} \frac{FFA_B - FFA_M}{1 + \frac{FFA_B}{Kdif_{ffat}^{FFA_{BM}}} + \frac{FFA_M}{Kdif_{ffat}^{FFA_M}}} \tag{S81}$$

$$v_{ffat\_a}^{FFA_{BM}} = Vmax_{ffat}^{FFA_{BM}} \frac{FFA_B}{FFA_B + Km_{ffat}^{FFA_{BM}}} \tag{S82}$$

TG synthesis (TGSYN)

$$v_{tgsyn}^{FFA_M} = Vmax_{tgsyn}^{FFA_M} \frac{FFA_M}{FFA_M + Km_{tgsyn}^{FFA_M}} \frac{Glycp_M}{Glycp_M + Km_{tgsyn}^{Glycp_M}} \frac{TG_M^{max} - TG_M}{(TG_M^{max} - TG_M) + Km_{tgsyn}^{TG_M}} \frac{Atp_M}{Atp_M + Km_{tgsyn}^{Atp_M}} \quad \text{(S83)}$$

TG degradation (TGDEG)

$$v_{tgdeg}^{TG_L} = \beta_L Vmax_{tgdeg}^{TG_L} \frac{TG_L}{TG_L + Km_{tgdeg}^{TG_L}} \quad \text{(S84)}$$

Glycerol transporter (GLYCT)

$$v_{glyct}^{Glyc_{BM}} = Vdif_{glyct}^{Glyc_{BM}} \frac{Glyc_B - Glyc_M}{1 + \frac{Glyc_B}{Kdif_{glyct}^{Glyc_{BM}}} + \frac{Glyc_M}{Kdif_{glyct}^{Glyc_M}}} \quad \text{(S85)}$$

Glycerol kinase (GLYK)

$$v_{glyk}^{Glyc_M} = Vmax_{glyk}^{Glyc_M} \frac{Glyc_M}{Glyc_M + Km_{glyk}^{Glyc_M}(1 + \frac{Gap_M}{Ki_{glyk}^{Gap_M}})} \frac{Atp_M}{Atp_M + Km_{glyk}^{Atp_M}} \quad \text{(S86)}$$

TG transporter (TGT)

$$v_{tgt}^{TG_{BM}} = v_{tgt\_a}^{TG_{BM}} + v_{tgt\_d}^{TG_{BM}} \quad \text{(S80)}$$

$$v_{tgt\_d}^{TG_{BM}} = Vdif_{tgt}^{TG_{BM}} \frac{TG_B - TG_M / Keq_{tgt}^{TG_M}}{TG_B + Kdif_{tgt}^{TG_{BM}} + \frac{TG_M}{Keq_{tgt}^{TG_M}}} \quad \text{(S87)}$$

$$v_{tgt\_a}^{TG_{BM}} = Vmax_{tgt}^{TG_{BM}} \frac{TG_B}{TG_B + Km_{tgt}^{TG_M}} \quad \text{(S88)}$$

Glycerol-3-phosphate dehydrogenase (G3PD)

$$v_{g3pd}^{Gap_M} = Vmax_{g3pd}^{Gap_M} \frac{Gap_M Nadh_M - \frac{Glycp_M}{Keq_{g3pd}^{Glycp_M}} Nad_M}{(1 + \frac{Gap_M}{Km_{g3pd}^{Gap_M}})(1 + \frac{Nadh_M}{Km_{g3pd}^{Nadh_M}}) + (1 + \frac{Glycp_M}{Km_{g3pd}^{Glycp_M}})(1 + \frac{Nad_M}{Km_{g3pd}^{Nad_M}}) - 1} \quad \text{(S89)}$$

β-oxidation (BOXID)

$$v_{boxid}^{FFA_M} = Vmax_{boxid}^{FFA_M} \frac{FFA_M}{FFA_M + Km_{boxid}^{FFA_M}} \frac{Atp_M}{Atp_M + Km_{boxid}^{Atp_M}} \frac{Nad_M}{Nad_M + Km_{boxid}^{Nad_M}} \frac{Fad_M}{Fad_M + Km_{boxid}^{Fad_M}}$$
$$\cdot \frac{Ki_{boxid}^{Accoa_{MM}}}{Accoa_{MM} + Ki_{boxid}^{Accoa_{MM}}} \frac{Ki_{boxid}^{Malcoa_M}}{Malcoa_M + Ki_{boxid}^{Malcoa_M}} \quad \text{(S90)}$$

ATP synthesis from FADH (ATPSYNF)

$$v_{atpsynf}^{Fadh_M} = Vmax_{atpsynf}^{Fadh_M} \frac{Fadh_M}{Fadh_M + Km_{atpsynf}^{Fadh_M}} \frac{Adp_M}{Adp_M + Km_{atpsynf}^{Adp_M}} \quad \text{(S91)}$$

ATP synthesis from NADH (ATPSYNN)

$$v_{atpsynn}^{Nadh_M} = Vmax_{atpsynn}^{Nadh_M} \frac{Nadh_M}{Nadh_M + Km_{atpsynn}^{Nadh_M}} \frac{Adp_M}{Adp_M + Km_{atpsynn}^{Adp_M}} \quad \text{(S92)}$$

ATP utilization (ATPUSE)

$$v_{atpuse}^{Atp_M} = Vmax_{atpuse}^{Atp_M} \frac{Atp_M}{Atp_M + Km_{atpuse}^{Atp_M}} \quad \text{(S93)}$$

Adenosine kinase (AMP regeneration into ADP (AMPREG))

$$v_{ampreg}^{Amp_M} = Vmax_{ampreg}^{Amp_M} \left( \frac{Amp_M}{Amp_M + Km_{ampreg}^{Amp_M}} \frac{Atp_M}{Atp_M + Km_{ampreg}^{Atp_M}} - \frac{Adp_M}{Adp_M + Km_{ampreg}^{Adp_M}} \frac{Adp_M}{Adp_M + Km_{ampreg}^{Adp_M}} \right) \quad \text{(S94)}$$

Uridine diphosphate kinase (UDP regeneration to UDP (UDPREG))

$$v_{udpreg}^{Udp_M} = Vmax_{udpreg}^{Udp_M} \frac{Udp_M\, Atp_M - \frac{Utp_M\, Adp_M}{Keq_{udpreg}^{Utp_M}}}{(1+\frac{Udp_M}{Km_{udpreg}^{Udp_M}})(1+\frac{Atp_M}{Km_{udpreg}^{Atp_M}}) + (1+\frac{Utp_M}{Km_{udpreg}^{Utp_M}})(1+\frac{Adp_M}{Km_{udpreg}^{Adp_M}}) - 1} \quad \text{(S95)}$$

UTP utilization (UTPUSE)

$$v_{utpuse}^{Utp_M} = Vmax_{utpuse}^{Utp_M} \frac{Utp_M}{Utp_M + Km_{utpuse}^{Utp_M}} \quad \text{(S96)}$$

NADH utilization (NADHUSE)

$$v_{nadhuse}^{Nadh_M} = Vmax_{nadhuse}^{Nadh_M} \frac{Nadh_M}{Nadh_M + Km_{nadhuse}^{Nadh_M}} \quad \text{(S97)}$$

NADPH utilization (NADPHUSE)

$$v_{nadphuse}^{Nadph_L} = Vmax_{nadphuse}^{Nadph_L} \frac{Nadph_L}{Nadph_L + Km_{nadphuse}^{Nadph_L}} \tag{S98}$$

FADH utilization (FADHUSE)

$$v_{fadhuse}^{Fadh_M} = Vmax_{fadhuse}^{Fadh_M} \frac{Fadh_M}{Fadh_M + Km_{fadhuse}^{Fadh_M}} \tag{S99}$$

Creatine kinase (CK)

$$v_{ck}^{Cre_M} = Vmax_{ck}^{Cre_M} \cdot \frac{Cre_M \cdot Atp_M - Crep_M \cdot Adp_M / Keq_{ck}^{Crep_M}}{(1+\frac{Cre_M}{Km_{ck}^{Cre_M}})(1+\frac{Atp_M}{Km_{ck}^{Atp_M}}) + (1+\frac{Crep_M}{Km_{ck}^{Crep_M}})(1+\frac{Adp_M}{Km_{ck}^{Adp_M}}) - 1} \tag{S100}$$

**Flux in adipose tissue**

Glucose transporter (GLUT4)

$$v_{glut4}^{Glc_{BA}} = \alpha_A \cdot Vdif_{glut4}^{Glc_{BA}} \frac{Glc_B - Glc_A}{1 + \frac{Glc_B}{Kdif_{glut4}^{Glc_{BA}}} + \frac{Glc_A}{Kdif_{glut4}^{Glc_A}}} \tag{S101}$$

Hexokinase (HK)

$$v_{hk}^{Glc_A} = \alpha_A \cdot Vmax_{hk}^{Glc_A} \frac{Glc_A}{Glc_A + Km_{hk}^{Glc_A}(1+\frac{G6p_A}{Ki_{hk}^{G6p_A}})} \frac{Atp_A}{Atp_A + Km_{hk}^{Atp_A}(1+\frac{G6p_A}{Ki_{hk}^{Atp_A}})} \tag{S102}$$

Phosphofructokinase (PFK)

$$v_{pfk}^{G6p_A} = \alpha_A \cdot Vmax_{pfk}^{G6p_A} \frac{G6p_A}{G6p_A + Km_{pfk}^{G6p_A}} \frac{Atp_A}{Atp_A + Km_{pfk}^{Atp_A}} \cdot \frac{Ki_{pfk}^{Atp_A}}{Atp_A + Ki_{pfk}^{Atp_A}} \frac{Adp_A}{Adp_A + Km_{pfk}^{Adp_A}} (1 - b_{pfk}^{Gap_A} \frac{Gap_A}{Gap_A + Ki_{pfk}^{Gap_A}}) \tag{S103}$$

Pyruvate kinase (PK)

$$v_{pk}^{Gap_A} = \alpha_A \cdot Vmax_{pk}^{Gap_A} \frac{Gap_A}{Gap_A + Km_{pk}^{Gap_A}} (1 - b_{pk}^{Accoa_{AM}} \frac{Accoa_{AM}}{Accoa_{AM} + Ki_{pk}^{Accoa_{AM}}}) \frac{Adp_A}{Adp_A + Km_{pk}^{Adp_A}} \tag{S104}$$

Pyruvate transporter (PYRT)

$$v_{pyrt}^{Pyr_{BA}} = Vdif_{pyrt}^{Pyr_{BA}} \frac{Pyr_B - Pyr_A}{1 + \frac{Pyr_B}{Kdif_{pyrt}^{Pyr_{BM}}} + \frac{Pyr_A}{Kdif_{pyrt}^{Pyr_A}}} \tag{S105}$$

Lactate transporter (LACT)

$$v_{lact}^{Lac_{BA}} = Vdif_{lact}^{Lac_{BA}} \frac{Lac_B - Lac_A}{1 + \frac{Lac_B}{Kdif_{lact}^{Lac_{BM}}} + \frac{Lac_A}{Kdif_{lact}^{Lac_A}}} \tag{S106}$$

Lactate dehydrogenase (LDH)

$$v_{ldh}^{Pyr_A} = Vmax_{ldh}^{Pyr_A} \frac{Pyr_A \cdot Nadh_A - Lac_A \cdot Nad_A / Keq_{ldh}^{Lac_A}}{(1 + \frac{Pyr_A}{Km_{ldh}^{Pyr_A}})(1 + \frac{Nadh_A}{Km_{ldh}^{Nadh_A}}) + (1 + \frac{Lac_A}{Km_{ldh}^{Lac_A}})(1 + \frac{Nad_A}{Km_{ldh}^{Nad_A}}) - 1} \tag{S107}$$

Alanine transporter (ALAT)

$$v_{alat}^{Ala_{BA}} = Vdif_{alat}^{Ala_{BA}} \frac{Ala_B - Ala_A}{1 + \frac{Ala_B}{Kdif_{alat}^{Ala_{BA}}} + \frac{Ala_A}{Kdif_{alat}^{Ala_A}}} \tag{S108}$$

Alanine transaminase (ALAT)

$$v_{alata}^{Pyr_A} = Vmax_{alata}^{Pyr_A} \cdot \frac{Pyr_A - Ala_A / Keq_{alata}^{Ala_A}}{(1 + \frac{Pyr_A}{Km_{alata}^{Pyr_A}}) + (1 + \frac{Ala_A}{Km_{alata}^{Ala_A}}) - 1} \tag{S109}$$

Pyruvate dehydrogenase (PDH)

$$v_{pdh}^{Pyr_A} = \alpha_A \cdot Vmax_{pdh}^{Pyr_A} \frac{Pyr_A}{Pyr_A + Km_{pdh}^{Pyr_A}} \frac{Nad_A}{Nad_A + Km_{pdh}^{Nad_A}} \frac{Km_{pdh}^{Accoa_{AM}}}{Accoa_{AM} + Km_{pdh}^{Accoa_{AM}}} \tag{S110}$$

TCA cycle (TCA)

$$v_{tca}^{Accoa_{AM}} = Vmax_{tca}^{Accoa_{AM}} \frac{Accoa_{AM}}{Accoa_{AM} + Km_{tca}^{Accoa_{AM}}} \frac{Adp_A}{Adp_A + Km_{tca}^{Adp_A}} \frac{Nad_A}{Nad_A + Km_{tca}^{Nad_A}} \frac{Fad_A}{Fad_A + Km_{tca}^{Fad_A}} \frac{Phos_A}{Phos_A + Km_{tca}^{Phos_A}} \frac{Pyr_A}{Pyr_A + Km_{tca}^{Pyr_A}} \tag{S111}$$

FFA transporter (FFAT)

$$v_{ffat}^{FFA_{BA}} = v_{ffat\_d}^{FFA_{BA}} + v_{ffat\_a}^{FFA_{BA}} \tag{S112}$$

$$v_{ffat\_d}^{FFA_{BA}} = Vdif_{ffat}^{FFA_{BA}} \frac{FFA_B - FFA_A}{1 + \frac{FFA_B}{Kdif_{ffat}^{FFA_{BA}}} + \frac{FFA_M}{Kdif_{ffat}^{FFA_M}}} \tag{S113}$$

$$v_{ffat\_a}^{FFA_{BA}} = -Vmax_{ffat}^{FFA_{BA}} \frac{FFA_A}{FFA_A + Km_{ffat}^{FFA_{BA}}} \tag{S114}$$

TG synthesis (TGSYN)

$$v_{tgsyn}^{FFA_A} = Vmax_{tgsyn}^{FFA_A} \frac{FFA_A}{FFA_A + Km_{tgsyn}^{FFA_A}} \frac{Glycp_A}{Glycp_A + Km_{tgsyn}^{Glycp_A}} \frac{TG_A^{\max} - TG_A}{(TG_A^{\max} - TG_A) + Km_{tgsyn}^{TG_A}} \frac{Atp_A}{Atp_A + Km_{tgsyn}^{Atp_A}} \quad \text{(S115)}$$

TG degradation (TGDEG)

$$v_{tgdeg}^{TG_A} = \beta_A Vmax_{tgdeg}^{TG_A} \frac{TG_A}{TG_A + Km_{tgdeg}^{TG_A}} \quad \text{(S116)}$$

Glycerol transporter (GLYCT)

$$v_{glyct}^{Glyc_{BA}} = Vdif_{glyct}^{Glyc_{BA}} \frac{Glyc_B - Glyc_A}{1 + \frac{Glyc_B}{Kdif_{glyct}^{Glyc_{BA}}} + \frac{Glyc_A}{Kdif_{glyct}^{Glyc_A}}} \quad \text{(S117)}$$

Glycerol kinase (GLYK)

$$v_{glyk}^{Glyc_A} = Vmax_{glyk}^{Glyc_A} \frac{Glyc_A}{Glyc_A + Km_{glyk}^{Glyc_A}(1 + \frac{Gap_A}{Ki_{glyk}^{Gap_A}})} \frac{Atp_A}{Atp_A + Km_{glyk}^{Atp_A}} \quad \text{(S118)}$$

TG transporter (TGT)

$$v_{tgt}^{TG_{BM}} = v_{tgt\_a}^{TG_{BM}} + v_{tgt\_d}^{TG_{BM}} \quad \text{(S119)}$$

$$v_{tgt\_a}^{TG_{BA}} = \alpha_A \cdot Vmax_{tgt}^{TG_{BA}} \frac{TG_B}{TG_B + Km_{tgt}^{TG_A}} \quad \text{(S120)}$$

$$v_{tgt\_d}^{TG_{BA}} = Vdif_{tgt}^{TG_{BA}} \frac{TG_B - TG_A \Big/ Keq_{tgt}^{TG_A}}{TG_B + Kdif_{tgt}^{TG_{BA}} + \frac{TG_A}{Keq_{tgt}^{TG_A}}} \quad \text{(S121)}$$

Glycerol-3-phosphate dehydrogenase (G3PD)

$$v_{g3pd}^{Gap_A} = Vmax_{g3pd}^{Gap_A} \frac{Gap_A Nadh_A - \frac{Glycp_A}{Keq_{g3pd}^{Glycp_A}} Nad_A}{(1 + \frac{Gap_A}{Km_{g3pd}^{Gap_A}})(1 + \frac{Nadh_A}{Km_{g3pd}^{Nadh_A}}) + (1 + \frac{Glycp_A}{Km_{g3pd}^{Glycp_A}})(1 + \frac{Nad_A}{Km_{g3pd}^{Nad_A}}) - 1} \quad \text{(S122)}$$

β-oxidation (BOXID)

$$v_{boxid}^{FFA_A} = Vmax_{boxid}^{FFA_A} \frac{FFA_A}{FFA_A + Km_{boxid}^{FFA_A}} \frac{Atp_A}{Atp_A + Km_{boxid}^{Atp_A}} \frac{Nad_A}{Nad_A + Km_{boxid}^{Nad_A}} \frac{Fad_A}{Fad_A + Km_{boxid}^{Fad_A}} \cdot \frac{Ki_{boxid}^{Accoa_{AM}}}{Accoa_{AM} + Ki_{boxid}^{Accoa_{AM}}} \frac{Ki_{boxid}^{Malcoa_A}}{Malcoa_A + Ki_{boxid}^{Malcoa_A}} \quad \text{(S123)}$$

ATP synthesis from FADH (ATPSYNF)

$$v_{atpsynf}^{Fadh_A} = Vmax_{atpsynf}^{Fadh_A} \frac{Fadh_A}{Fadh_A + Km_{atpsynf}^{Fadh_A}} \frac{Adp_A}{Adp_A + Km_{atpsynf}^{Adp_A}} \quad \text{(S124)}$$

ATP synthesis from NADH (ATPSYNN)

$$v_{atpsynn}^{Nadh_A} = Vmax_{atpsynn}^{Nadh_A} \frac{Nadh_A}{Nadh_A + Km_{atpsynn}^{Nadh_A}} \frac{Adp_A}{Adp_A + Km_{atpsynn}^{Adp_A}} \quad \text{(S125)}$$

ATP utilization (ATPUSE)

$$v_{atpuse}^{Atp_A} = Vmax_{atpuse}^{Atp_A} \frac{Atp_A}{Atp_A + Km_{atpuse}^{Atp_A}} \quad \text{(S126)}$$

Adenosine kinase (AMP regeneration into ADP (AMPREG))

$$v_{ampreg}^{Amp_A} = Vmax_{ampreg}^{Amp_A} \left( \frac{Amp_A}{Amp_A + Km_{ampreg}^{Amp_A}} \frac{Atp_A}{Atp_A + Km_{ampreg}^{Atp_A}} - \frac{Adp_A}{Adp_A + Km_{ampreg}^{Adp_A}} \frac{Adp_A}{Adp_A + Km_{ampreg}^{Adp_A}} \right) \quad \text{(S127)}$$

NADH utilization (NADHUSE)

$$v_{nadhuse}^{Nadh_A} = Vmax_{nadhuse}^{Nadh_A} \frac{Nadh_A}{Nadh_A + Km_{nadhuse}^{Nadh_A}} \quad \text{(S128)}$$

FADH utilization (FADHUSE)

$$v_{fadhuse}^{Fadh_A} = Vmax_{fadhuse}^{Fadh_A} \frac{Fadh_A}{Fadh_A + Km_{fadhuse}^{Fadh_A}} \quad \text{(S129)}$$

Creatine kinase (CK)

$$v_{ck}^{Cre_A} = Vmax_{ck}^{Cre_A} \cdot \frac{Cre_A \cdot Atp_A - Crep_A \cdot Adp_A / Keq_{ck}^{Crep_A}}{(1 + \frac{Cre_A}{Km_{ck}^{Cre_A}})(1 + \frac{Atp_A}{Km_{ck}^{Atp_A}}) + (1 + \frac{Crep_A}{Km_{ck}^{Crep_A}})(1 + \frac{Adp_A}{Km_{ck}^{Adp_A}}) - 1} \quad \text{(S130)}$$

**Flux in GI tract**

Glucose transporter (GLUT2)

$$v_{glut2}^{Glc_{BG}} = Vdif_{glut2}^{Glc_{BG}} \frac{Glc_B - Glc_G}{1 + \frac{Glc_B}{Kdif_{glut2}^{Glc_{BG}}} + \frac{Glc_G}{Kdif_{glut2}^{Glc_G}}} \quad \text{(S131)}$$

Hexokinase (HK)

$$v_{hk}^{Glc_G} = \alpha_G \cdot Vmax_{hk}^{Glc_G} \frac{Glc_G}{Glc_G + Km_{hk}^{Glc_G}(1 + \frac{G6p_G}{Ki_{hk}^{G6p_G}})} \frac{Atp_G}{Atp_G + Km_{hk}^{Atp_G}(1 + \frac{G6p_G}{Ki_{hk}^{Atp_G}})} \quad \text{(S132)}$$

Phosphofructokinase (PFK)

$$v_{pfk}^{G6p_G} = \alpha_G \cdot Vmax_{pfk}^{G6p_G} \frac{G6p_G}{G6p_G + Km_{pfk}^{G6p_G}} \frac{Atp_G}{Atp_G + Km_{pfk}^{Atp_G}} \cdot \frac{Ki_{pfk}^{Atp_G}}{Atp_G + Ki_{pfk}^{Atp_G}} \frac{Adp_G}{Adp_G + Km_{pfk}^{Adp_G}} (1 - b_{pfk}^{Gap_G} \frac{Gap_G}{Gap_G + Ki_{pfk}^{Gap_G}}) \tag{S133}$$

Pyruvate kinase (PK)

$$v_{pk}^{Gap_G} = \alpha_G \cdot Vmax_{pk}^{Gap_G} \frac{Gap_G}{Gap_G + Km_{pk}^{Gap_G}} (1 - b_{pk}^{Accoa_{GM}} \frac{Accoa_{GM}}{Accoa_{GM} + Ki_{pk}^{Accoa_{GM}}}) \frac{Adp_G}{Adp_G + Km_{pk}^{Adp_G}} \tag{S134}$$

Pyruvate transporter (PYRT)

$$v_{pyrt}^{Pyr_{BG}} = Vdif_{pyrt}^{Pyr_{BG}} \frac{Pyr_B - Pyr_G}{1 + \frac{Pyr_B}{Kdif_{pyrt}^{Pyr_{BG}}} + \frac{Pyr_G}{Kdif_{pyrt}^{Pyr_G}}} \tag{S135}$$

Lactate transporter (LACT)

$$v_{lact}^{Lac_{BA}} = Vdif_{lact}^{Lac_{BA}} \frac{Lac_B - Lac_A}{1 + \frac{Lac_B}{Kdif_{lact}^{Lac_{BM}}} + \frac{Lac_A}{Kdif_{lact}^{Lac_A}}} \tag{S136}$$

Lactate dehydrogenase (LDH)

$$v_{ldh}^{Pyr_G} = Vmax_{ldh}^{Pyr_G} \frac{Pyr_G \cdot Nadh_G - Lac_G \cdot Nad_G / Keq_{ldh}^{Lac_G}}{(1 + \frac{Pyr_G}{Km_{ldh}^{Pyr_G}})(1 + \frac{Nadh_G}{Km_{ldh}^{Nadh_G}}) + (1 + \frac{Lac_G}{Km_{ldh}^{Lac_G}})(1 + \frac{Nad_G}{Km_{ldh}^{Nad_G}}) - 1} \tag{S137}$$

Alanine transporter (ALAT)

$$v_{alat}^{Ala_{BG}} = Vdif_{alat}^{Ala_{BG}} \frac{Ala_B - Ala_G}{1 + \frac{Ala_B}{Kdif_{alat}^{Ala_{BG}}} + \frac{Ala_G}{Kdif_{alat}^{Ala_G}}} \tag{S139}$$

Alanine transaminase (ALAT)

$$v_{alata}^{Pyr_G} = Vmax_{alata}^{Pyr_G} \cdot \frac{Pyr_G - Ala_G / Keq_{alata}^{Ala_G}}{(1 + \frac{Pyr_G}{Km_{alata}^{Pyr_G}}) + (1 + \frac{Ala_G}{Km_{alata}^{Ala_G}}) - 1} \tag{S139}$$

Pyruvate dehydrogenase (PDH)

$$v_{pdh}^{Pyr_G} = \alpha_G \cdot Vmax_{pdh}^{Pyr_G} \frac{Pyr_G}{Pyr_G + Km_{pdh}^{Pyr_G}} \frac{Nad_G}{Nad_G + Km_{pdh}^{Nad_G}} \frac{Km_{pdh}^{Accoa_{GM}}}{Accoa_{GM} + Km_{pdh}^{Accoa_{GM}}} \tag{S140}$$

TCA cycle (TCA)

$$v_{tca}^{Accoa_{GM}} = Vmax_{tca}^{Accoa_{GM}} \frac{Accoa_{GM}}{Accoa_{GM} + Km_{tca}^{Accoa_{GM}}} \frac{Adp_G}{Adp_G + Km_{tca}^{Adp_G}} \frac{Nad_G}{Nad_G + Km_{tca}^{Nad_G}} \frac{Fad_G}{Fad_G + Km_{tca}^{Fad_G}} \frac{Phos_G}{Phos_G + Km_{tca}^{Phos_G}} \frac{Pyr_G}{Pyr_G + Km_{tca}^{Pyr_G}} \quad \text{(S141)}$$

FFA transporter (FFAT)

$$v_{ffat}^{FFA_{BG}} = v_{ffat\_d}^{FFA_{BG}} + v_{ffat\_a}^{FFA_{BG}} \quad \text{(S142)}$$

$$v_{ffat\_d}^{FFA_{BG}} = Vdif_{ffat}^{FFA_{BG}} \frac{FFA_B - FFA_G}{1 + \frac{FFA_B}{Kdif_{ffat}^{FFA_{BG}}} + \frac{FFA_G}{Kdif_{ffat}^{FFA_G}}} \quad \text{(S143)}$$

$$v_{ffat\_a}^{FFA_{BG}} = -Vmax_{ffat}^{FFA_{BG}} \frac{FFA_G}{FFA_G + Km_{ffat}^{FFA_{BG}}} \quad \text{(S144)}$$

TG synthesis (TGSYN)

$$v_{tgsyn}^{FFA_G} = Vmax_{tgsyn}^{FFA_G} \frac{FFA_G}{FFA_G + Km_{tgsyn}^{FFA_G}} \frac{Glycp_G}{Glycp_G + Km_{tgsyn}^{Glycp_G}} \frac{TG_G^{\max} - TG_G}{(TG_G^{\max} - TG_G) + Km_{tgsyn}^{TG_G}} \frac{Atp_G}{Atp_G + Km_{tgsyn}^{Atp_G}} \quad \text{(S145)}$$

TG degradation (TGDEG)

$$v_{tgdeg}^{TG_G} = \beta_G Vmax_{tgdeg}^{TG_G} \frac{TG_G}{TG_G + Km_{tgdeg}^{TG_G}} \quad \text{(S146)}$$

Glycerol transporter (GLYCT)

$$v_{glyct}^{Glyc_{BG}} = Vdif_{glyct}^{Glyc_{BG}} \frac{Glyc_B - Glyc_G}{1 + \frac{Glyc_B}{Kdif_{glyct}^{Glyc_{BG}}} + \frac{Glyc_G}{Kdif_{glyct}^{Glyc_G}}} \quad \text{(S147)}$$

Glycerol kinase (GLYK)

$$v_{glyk}^{Glyc_G} = Vmax_{glyk}^{Glyc_G} \frac{Glyc_G}{Glyc_G + Km_{glyk}^{Glyc_G} (1 + \frac{Gap_G}{Ki_{glyk}^{Gap_G}})} \frac{Atp_G}{Atp_G + Km_{glyk}^{Atp_G}} \quad \text{(S147)}$$

TG transporter (TGT)

$$v_{tgt}^{TG_{BG}} = v_{tgt\_a}^{TG_{BG}} + v_{tgt\_d}^{TG_{BG}} \quad \text{(S148)}$$

$$v_{tgt\_a}^{TG_{BG}} = \alpha_G \cdot Vmax_{tgt}^{TG_{BG}} \frac{TG_B}{TG_B + Km_{tgt}^{TG_G}} \quad \text{(S149)}$$

$$v_{tgt\_d}^{TG_{BG}} = Vdif_{tgt}^{TG_{BG}} \frac{TG_B - TG_G \Big/ Keq_{tgt}^{TG_G}}{TG_B + Kdif_{tgt}^{TG_{BG}} + \dfrac{TG_G}{Keq_{tgt}^{TG_G}}} \tag{S150}$$

Glycerol-3-phosphate dehydrogenase (G3PD)

$$v_{g3pd}^{Gap_G} = Vmax_{g3pd}^{Gap_G} \frac{Gap_G Nadh_G - \dfrac{Glycp_G}{Keq_{g3pd}^{Glycp_G}} Nad_G}{(1+\dfrac{Gap_G}{Km_{g3pd}^{Gap_G}})(1+\dfrac{Nadh_G}{Km_{g3pd}^{Nadh_G}}) + (1+\dfrac{Glycp_G}{Km_{g3pd}^{Glycp_G}})(1+\dfrac{Nad_G}{Km_{g3pd}^{Nad_G}}) - 1} \tag{S151}$$

β-oxidation (BOXID)

$$v_{boxid}^{FFA_G} = Vmax_{boxid}^{FFA_G} \frac{FFA_G}{FFA_G + Km_{boxid}^{FFA_G}} \frac{Atp_G}{Atp_G + Km_{boxid}^{Atp_G}} \frac{Nad_G}{Nad_G + Km_{boxid}^{Nad_G}} \frac{Fad_G}{Fad_G + Km_{boxid}^{Fad_G}} \cdot \frac{Ki_{boxid}^{Accoa_{GM}}}{Accoa_{GM} + Ki_{boxid}^{Accoa_{GM}}} \frac{Ki_{boxid}^{Malcoa_G}}{Malcoa_G + Ki_{boxid}^{Malcoa_G}} \tag{S152}$$

ATP synthesis from FADH (ATPSYNF)

$$v_{atpsynf}^{Fadh_G} = Vmax_{atpsynf}^{Fadh_G} \frac{Fadh_G}{Fadh_G + Km_{atpsynf}^{Fadh_G}} \frac{Adp_G}{Adp_G + Km_{atpsynf}^{Adp_G}} \tag{S153}$$

ATP synthesis from NADH (ATPSYNN)

$$v_{atpsynn}^{Nadh_G} = Vmax_{atpsynn}^{Nadh_G} \frac{Nadh_G}{Nadh_G + Km_{atpsynn}^{Nadh_G}} \frac{Adp_G}{Adp_G + Km_{atpsynn}^{Adp_G}} \tag{S154}$$

ATP utilization (ATPUSE)

$$v_{atpuse}^{Atp_G} = Vmax_{atpuse}^{Atp_G} \frac{Atp_G}{Atp_G + Km_{atpuse}^{Atp_G}} \tag{S155}$$

Adenosine kinase (AMP regeneration into ADP (AMPREG))

$$v_{ampreg}^{Amp_G} = Vmax_{ampreg}^{Amp_G} \left( \frac{Amp_G}{Amp_G + Km_{ampreg}^{Amp_G}} \frac{Atp_G}{Atp_G + Km_{ampreg}^{Atp_G}} - \frac{Adp_G}{Adp_G + Km_{ampreg}^{Adp_G}} \frac{Adp_G}{Adp_G + Km_{ampreg}^{Adp_G}} \right) \tag{S156}$$

NADH utilization (NADHUSE)

$$v_{nadhuse}^{Nadh_G} = Vmax_{nadhuse}^{Nadh_G} \frac{Nadh_G}{Nadh_G + Km_{nadhuse}^{Nadh_G}} \tag{S157}$$

FADH utilization (FADHUSE)

$$v_{fadhuse}^{Fadh_G} = Vmax_{fadhuse}^{Fadh_G} \frac{Fadh_G}{Fadh_G + Km_{fadhuse}^{Fadh_G}} \quad \text{(S158)}$$

Creatine kinase (CK)

$$v_{ck}^{Cre_G} = Vmax_{ck}^{Cre_G} \cdot \frac{Cre_G \cdot Atp_G - Crep_G \cdot Adp_G / Keq_{ck}^{Crep_G}}{(1+\frac{Cre_G}{Km_{ck}^{Cre_G}})(1+\frac{Atp_G}{Km_{ck}^{Atp_G}}) + (1+\frac{Crep_G}{Km_{ck}^{Crep_G}})(1+\frac{Adp_G}{Km_{ck}^{Adp_G}}) - 1} \quad \text{(S159)}$$

**Flux in heart**

Glucose transporter (GLUT2)

$$v_{glut2}^{Glc_{BH}} = Vdif_{glut2}^{Glc_{BH}} \frac{Glc_B - Glc_H}{1 + \frac{Glc_B}{Kdif_{glut2}^{Glc_{BH}}} + \frac{Glc_H}{Kdif_{glut2}^{Glc_H}}} \quad \text{(S160)}$$

Hexokinase (HK)

$$v_{hk}^{Glc_H} = Vmax_{hk}^{Glc_H} \frac{Glc_H}{Glc_H + Km_{hk}^{Glc_H}(1+\frac{G6p_H}{Ki_{hk}^{G6p_H}})} \frac{Atp_H}{Atp_H + Km_{hk}^{Atp_H}(1+\frac{G6p_H}{Ki_{hk}^{Atp_H}})} \quad \text{(S161)}$$

Glycogen synthase (GS)

$$v_{gs}^{G6p_H} = \alpha_H \cdot Vmax_{gs}^{G6p_H} \frac{G6p_H^{n_{gs}^{G6pH}}}{G6p_H^{n_{gs}^{G6pH}} + Km_{gs}^{G6p_H \, n_{gs}^{G6pH}}} \frac{(Glygn_H^{\max} - Glygn_H)}{(Glygn_H^{\max} - Glygn_H) + Km_{gs}^{Glygn_H}} \frac{Utp_H}{Utp_H + Km_{gs}^{Utp_H}} \quad \text{(S162)}$$

$Glygn_H^{\max}$ denotes the maximum amount of glycogen that heart can store.

Glycogen degradation (GD) by glycogen phosphorylase (GP)

$$v_{gd}^{Glygn_H} = \beta_H \cdot Vmax_{gd}^{Glygn_H} \frac{Glygn_H}{Glygn_H + Km_{gd}^{Glygn_H}} \frac{Phos_H}{Phos_H + Km_{gd}^{Phos_H}} \quad \text{(S163)}$$

Phosphofructokinase (PFK)

$$v_{pfk}^{G6p_H} = Vmax_{pfk}^{G6p_H} \frac{G6p_H}{G6p_H + Km_{pfk}^{G6p_H}} \frac{Atp_H}{Atp_H + Km_{pfk}^{Atp_H}} \cdot \frac{Ki_{pfk}^{Atp_H}}{Atp_H + Ki_{pfk}^{Atp_H}} \frac{Adp_H}{Adp_H + Km_{pfk}^{Adp_H}} (1 - b_{pfk}^{Gap_H} \frac{Gap_H}{Gap_H + Ki_{pfk}^{Gap_H}}) \quad \text{(S164)}$$

Pyruvate kinase (PK)

$$v_{pk}^{Gap_H} = Vmax_{pk}^{Gap_H} \frac{Gap_H}{Gap_H + Km_{pk}^{Gap_H}} (1 - b_{pk}^{Accoa_{HM}} \frac{Accoa_{HM}}{Accoa_{HM} + Ki_{pk}^{Accoa_{HM}}}) \frac{Adp_H}{Adp_H + Km_{pk}^{Adp_H}} \quad \text{(S165)}$$

Pyruvate transporter (PYRT)

$$v_{pyrt}^{Pyr_{BH}} = Vdif_{pyrt}^{Pyr_{BH}} \frac{Pyr_B - Pyr_H}{1 + \frac{Pyr_B}{Kdif_{pyrt}^{Pyr_{BH}}} + \frac{Pyr_H}{Kdif_{pyrt}^{Pyr_H}}} \quad \text{(S166)}$$

Lactate transporter (LACT)

$$v_{lact}^{Lac_{BH}} = Vdif_{lact}^{Lac_{BH}} \frac{Lac_B - Lac_H}{1 + \frac{Lac_B}{Kdif_{lact}^{Lac_{BH}}} + \frac{Lac_H}{Kdif_{lact}^{Lac_H}}} \quad \text{(S167)}$$

Lactate dehydrogenase (LDH)

$$v_{ldh}^{Pyr_H} = Vmax_{ldh}^{Pyr_H} \frac{Pyr_H \cdot Nadh_H - Lac_H \cdot Nad_H / Keq_{ldh}^{Lac_H}}{(1 + \frac{Pyr_H}{Km_{ldh}^{Pyr_H}})(1 + \frac{Nadh_H}{Km_{ldh}^{Nadh_H}}) + (1 + \frac{Lac_H}{Km_{ldh}^{Lac_H}})(1 + \frac{Nad_H}{Km_{ldh}^{Nad_H}}) - 1} \quad \text{(S168)}$$

Alanine transporter (ALAT)

$$v_{alat}^{Ala_{BH}} = Vdif_{alat}^{Ala_{BH}} \frac{Ala_B - Ala_H}{1 + \frac{Ala_B}{Kdif_{alat}^{Ala_{BH}}} + \frac{Ala_H}{Kdif_{alat}^{Ala_H}}} \quad \text{(S169)}$$

Alanine transaminase (ALAT)

$$v_{alata}^{Pyr_H} = Vmax_{alata}^{Pyr_H} \cdot \frac{Pyr_H - Ala_H / Keq_{alata}^{Ala_H}}{(1 + \frac{Pyr_H}{Km_{alata}^{Pyr_H}}) + (1 + \frac{Ala_H}{Km_{alata}^{Ala_H}}) - 1} \quad \text{(S170)}$$

Pyruvate dehydrogenase (PDH)

$$v_{pdh}^{Pyr_H} = Vmax_{pdh}^{Pyr_H} \frac{Pyr_H}{Pyr_H + Km_{pdh}^{Pyr_H}} \frac{Nad_H}{Nad_H + Km_{pdh}^{Nad_H}} \frac{Km_{pdh}^{Accoa_{HM}}}{Accoa_{HM} + Km_{pdh}^{Accoa_{HM}}} \quad \text{(S171)}$$

TCA cycle (TCA)

$$v_{tca}^{Accoa_{HM}} = Vmax_{tca}^{Accoa_{HM}} \frac{Accoa_{HM}}{Accoa_{HM} + Km_{tca}^{Accoa_{HM}}} \frac{Adp_H}{Adp_H + Km_{tca}^{Adp_H}} \frac{Nad_H}{Nad_H + Km_{tca}^{Nad_H}} \frac{Fad_H}{Fad_H + Km_{tca}^{Fad_H}} \frac{Phos_H}{Phos_H + Km_{tca}^{Phos_H}} \frac{Pyr_H}{Pyr_H + Km_{tca}^{Pyr_H}} \quad \text{(S172)}$$

FFA transporter (FFAT)

$$v_{ffat}^{FFA_{BH}} = v_{ffat\_d}^{FFA_{BH}} + v_{ffat\_a}^{FFA_{BH}} \quad \text{(S173)}$$

$$v_{ffat\_d}^{FFA_{BH}} = Vdif_{ffat}^{FFA_{BH}} \frac{FFA_B - FFA_H}{1 + \frac{FFA_B}{Kdif_{ffat}^{FFA_{BH}}} + \frac{FFA_H}{Kdif_{ffat}^{FFA_H}}} \quad \text{(S174)}$$

$$v_{ffat\_a}^{FFA_{BH}} = Vmax_{ffat}^{FFA_{BH}} \frac{FFA_B}{FFA_B + Km_{ffat}^{FFA_{BH}}} \quad \text{(S175)}$$

TG synthesis (TGSYN)

$$v_{tgsyn}^{FFA_H} = Vmax_{tgsyn}^{FFA_H} \frac{FFA_H}{FFA_H + Km_{tgsyn}^{FFA_H}} \frac{Glycp_H}{Glycp_H + Km_{tgsyn}^{Glycp_H}} \frac{TG_H^{\max} - TG_H}{(TG_H^{\max} - TG_H) + Km_{tgsyn}^{TG_H}} \frac{Atp_H}{Atp_H + Km_{tgsyn}^{Atp_H}} \quad \text{(S176)}$$

TG degradation (TGDEG)

$$v_{tgdeg}^{TG_H} = Vmax_{tgdeg}^{TG_H} \frac{TG_H}{TG_H + Km_{tgdeg}^{TG_H}} \quad \text{(S177)}$$

Glycerol transporter (GLYCT)

$$v_{glyct}^{Glyc_{BH}} = Vdif_{glyct}^{Glyc_{BH}} \frac{Glyc_B - Glyc_H}{1 + \frac{Glyc_B}{Kdif_{glyct}^{Glyc_{BH}}} + \frac{Glyc_H}{Kdif_{glyct}^{Glyc_H}}} \quad \text{(S178)}$$

Glycerol kinase (GLYK)

$$v_{glyk}^{Glyc_H} = Vmax_{glyk}^{Glyc_H} \frac{Glyc_H}{Glyc_H + Km_{glyk}^{Glyc_H}(1 + \frac{Gap_H}{Ki_{glyk}^{Gap_H}})} \frac{Atp_H}{Atp_H + Km_{glyk}^{Atp_H}} \quad \text{(S179)}$$

TG transporter (TGT)

$$v_{tgt}^{TG_{BH}} = v_{tgt\_a}^{TG_{BH}} + v_{tgt\_d}^{TG_{BH}} \quad \text{(S180)}$$

$$v_{tgt\_d}^{TG_{BH}} = Vdif_{tgt}^{TG_{BH}} \frac{TG_B - TG_H \big/ Keq_{tgt}^{TG_H}}{TG_B + Kdif_{tgt}^{TG_{BH}} + \frac{TG_H}{Keq_{tgt}^{TG_H}}} \quad \text{(S181)}$$

$$v_{tgt\_a}^{TG_{BH}} = Vmax_{tgt}^{TG_{BH}} \frac{TG_B}{TG_B + Km_{tgt}^{TG_H}} \quad \text{(S182)}$$

Glycerol-3-phosphate dehydrogenase (G3PD)

$$v_{g3pd}^{Gap_H} = Vmax_{g3pd}^{Gap_H} \frac{Gap_H\, Nadh_H - \frac{Glycp_H}{Keq_{g3pd}^{Glycp_H}} Nad_H}{(1 + \frac{Gap_H}{Km_{g3pd}^{Gap_H}})(1 + \frac{Nadh_H}{Km_{g3pd}^{Nadh_H}}) + (1 + \frac{Glycp_H}{Km_{g3pd}^{Glycp_H}})(1 + \frac{Nad_H}{Km_{g3pd}^{Nad_H}}) - 1} \quad \text{(S183)}$$

β-oxidation (BOXID)

$$v_{boxid}^{FFA_H} = Vmax_{boxid}^{FFA_H} \frac{FFA_H}{FFA_H + Km_{boxid}^{FFA_H}} \frac{Atp_H}{Atp_H + Km_{boxid}^{Atp_H}} \frac{Nad_H}{Nad_H + Km_{boxid}^{Nad_H}} \frac{Fad_H}{Fad_H + Km_{boxid}^{Fad_H}} \cdot \frac{Ki_{boxid}^{Accoa_{HM}}}{Accoa_{HM} + Ki_{boxid}^{Accoa_{HM}}} \frac{Ki_{boxid}^{Malcoa_H}}{Malcoa_H + Ki_{boxid}^{Malcoa_H}} \quad \text{(S184)}$$

ATP synthesis from FADH (ATPSYNF)

$$v_{atpsynf}^{Fadh_H} = Vmax_{atpsynf}^{Fadh_H} \frac{Fadh_H}{Fadh_H + Km_{atpsynf}^{Fadh_H}} \frac{Adp_H}{Adp_H + Km_{atpsynf}^{Adp_H}} \quad \text{(S185)}$$

ATP synthesis from NADH (ATPSYNN)

$$v_{atpsynn}^{Nadh_H} = Vmax_{atpsynn}^{Nadh_H} \frac{Nadh_H}{Nadh_H + Km_{atpsynn}^{Nadh_H}} \frac{Adp_H}{Adp_H + Km_{atpsynn}^{Adp_H}} \quad \text{(S186)}$$

ATP utilization (ATPUSE)

$$v_{atpuse}^{Atp_H} = Vmax_{atpuse}^{Atp_H} \frac{Atp_H}{Atp_H + Km_{atpuse}^{Atp_H}} \quad \text{(S187)}$$

Adenosine kinase (AMP regeneration into ADP (AMPREG))

$$v_{ampreg}^{Amp_H} = Vmax_{ampreg}^{Amp_H} \left( \frac{Amp_H}{Amp_H + Km_{ampreg}^{Amp_H}} \frac{Atp_H}{Atp_H + Km_{ampreg}^{Atp_H}} - \frac{Adp_H}{Adp_H + Km_{ampreg}^{Adp_H}} \frac{Adp_H}{Adp_H + Km_{ampreg}^{Adp_H}} \right) \quad \text{(S188)}$$

Uridine diphosphate kinase (UDP regeneration to UDP (UDPREG))

$$v_{udpreg}^{Udp_H} = Vmax_{udpreg}^{Udp_H} \frac{Udp_H\, Atp_H - \dfrac{Utp_H\, Adp_H}{Keq_{udpreg}^{Utp_H}}}{(1 + \dfrac{Udp_H}{Km_{udpreg}^{Udp_H}})(1 + \dfrac{Atp_H}{Km_{udpreg}^{Atp_H}}) + (1 + \dfrac{Utp_H}{Km_{udpreg}^{Utp_H}})(1 + \dfrac{Adp_H}{Km_{udpreg}^{Adp_H}}) - 1} \quad \text{(S189)}$$

UTP utilization (UTPUSE)

$$v_{utpuse}^{Utp_H} = Vmax_{utpuse}^{Utp_H} \frac{Utp_H}{Utp_H + Km_{utpuse}^{Utp_H}} \quad \text{(S190)}$$

NADH utilization (NADHUSE)

$$v_{nadhuse}^{Nadh_H} = Vmax_{nadhuse}^{Nadh_H} \frac{Nadh_H}{Nadh_H + Km_{nadhuse}^{Nadh_H}} \quad \text{(S191)}$$

FADH utilization (FADHUSE)

$$v_{fadhuse}^{Fadh_H} = Vmax_{fadhuse}^{Fadh_H} \frac{Fadh_H}{Fadh_H + Km_{fadhuse}^{Fadh_H}} \tag{S192}$$

Creatine kinase (CK)

$$v_{ck}^{Cre_H} = Vmax_{ck}^{Cre_H} \cdot \frac{Cre_H \cdot Atp_H - Crep_H \cdot Adp_H / Keq_{ck}^{Crep_H}}{(1+\frac{Cre_H}{Km_{ck}^{Cre_M}})(1+\frac{Atp_H}{Km_{ck}^{Atp_H}}) + (1+\frac{Crep_H}{Km_{ck}^{Crep_H}})(1+\frac{Adp_H}{Km_{ck}^{Adp_H}}) - 1} \tag{S193}$$

**Flux in brain**

Glucose transporter (GLUT3)

$$v_{glut3}^{Glc_{BN}} = Vdif_{glut3}^{Glc_{BN}} \frac{Glc_B - Glc_N}{1+\frac{Glc_B}{Kdif_{glut3}^{Glc_{BN}}} + \frac{Glc_N}{Kdif_{glut3}^{Glc_N}}} \tag{S194}$$

Hexokinase (HK)

$$v_{hk}^{Glc_N} = Vmax_{hk}^{Glc_N} \frac{Glc_N}{Glc_N + Km_{hk}^{Glc_N}(1+\frac{G6p_N}{Ki_{hk}^{G6p_N}})} \frac{Atp_N}{Atp_N + Km_{hk}^{Atp_N}(1+\frac{G6p_N}{Ki_{hk}^{Atp_N}})} \tag{S195}$$

Glycogen synthase (GS)

$$v_{gs}^{G6p_N} = Vmax_{gs}^{G6p_N} \frac{G6p_N^{n_{gs}^{G6p_N}}}{G6p_N^{n_{gs}^{G6p_H}} + Km_{gs}^{G6p_N \, n_{gs}^{G6p_N}}} \frac{(Glygn_N^{\max} - Glygn_N)}{(Glygn_N^{\max} - Glygn_N) + Km_{gs}^{Glygn_N}} \frac{Utp_N}{Utp_N + Km_{gs}^{Utp_N}} \tag{S196}$$

$Glygn_N^{\max}$ denotes the maximum amount of glycogen that brain can store.

Glycogen degradation (GD) by glycogen phosphorylase (GP)

$$v_{gd}^{Glygn_N} = Vmax_{gd}^{Glygn_N} \frac{Glygn_N}{Glygn_N + Km_{gd}^{Glygn_N}} \frac{Phos_N}{Phos_N + Km_{gd}^{Phos_N}} \tag{S197}$$

Phosphofructokinase (PFK)

$$v_{pfk}^{G6p_N} = Vmax_{pfk}^{G6p_N} \frac{G6p_N}{G6p_N + Km_{pfk}^{G6p_N}} \frac{Atp_N}{Atp_N + Km_{pfk}^{Atp_N}} \cdot \frac{Ki_{pfk}^{Atp_N}}{Atp_N + Ki_{pfk}^{Atp_N}} \frac{Adp_N}{Adp_N + Km_{pfk}^{Adp_N}} (1 - b_{pfk}^{Gap_N} \frac{Gap_N}{Gap_N + Ki_{pfk}^{Gap_N}}) \tag{S198}$$

Pyruvate kinase (PK)

$$v_{pk}^{Gap_N} = Vmax_{pk}^{Gap_N} \frac{Gap_N}{Gap_N + Km_{pk}^{Gap_N}} (1 - b_{pk}^{Accoa_{NM}} \frac{Accoa_{NM}}{Accoa_{NM} + Ki_{pk}^{Accoa_{NM}}}) \frac{Adp_N}{Adp_N + Km_{pk}^{Adp_N}} \tag{S199}$$

Pyruvate transporter (PYRT)

$$v_{pyrt}^{Pyr_{BN}} = Vdif_{pyrt}^{Pyr_{BN}} \frac{Pyr_B - Pyr_N}{1 + \frac{Pyr_B}{Kdif_{pyrt}^{Pyr_{BN}}} + \frac{Pyr_N}{Kdif_{pyrt}^{Pyr_N}}} \tag{S200}$$

Lactate transporter (LACT)

$$v_{lact}^{Lac_{BN}} = Vdif_{lact}^{Lac_{BN}} \frac{Lac_B - Lac_N}{1 + \frac{Lac_B}{Kdif_{lact}^{Lac_{BN}}} + \frac{Lac_N}{Kdif_{lact}^{Lac_N}}} \tag{S201}$$

Lactate dehydrogenase (LDH)

$$v_{ldh}^{Pyr_N} = Vmax_{ldh}^{Pyr_N} \frac{Pyr_N \cdot Nadh_N - Lac_N \cdot Nad_N / Keq_{ldh}^{Lac_N}}{(1 + \frac{Pyr_N}{Km_{ldh}^{Pyr_N}})(1 + \frac{Nadh_N}{Km_{ldh}^{Nadh_N}}) + (1 + \frac{Lac_N}{Km_{ldh}^{Lac_N}})(1 + \frac{Nad_N}{Km_{ldh}^{Nad_N}}) - 1} \tag{S202}$$

Alanine transporter (ALAT)

$$v_{alat}^{Ala_{BN}} = Vdif_{alat}^{Ala_{BN}} \frac{Ala_B - Ala_N}{1 + \frac{Ala_B}{Kdif_{alat}^{Ala_{BN}}} + \frac{Ala_N}{Kdif_{alat}^{Ala_N}}} \tag{S203}$$

Alanine transaminase (ALAT)

$$v_{alata}^{Pyr_N} = Vmax_{alata}^{Pyr_N} \cdot \frac{Pyr_N - Ala_N / Keq_{alata}^{Ala_N}}{(1 + \frac{Pyr_N}{Km_{alata}^{Pyr_N}}) + (1 + \frac{Ala_N}{Km_{alata}^{Ala_N}}) - 1} \tag{S204}$$

Pyruvate dehydrogenase (PDH)

$$v_{pdh}^{Pyr_N} = Vmax_{pdh}^{Pyr_N} \frac{Pyr_N}{Pyr_N + Km_{pdh}^{Pyr_N}} \frac{Nad_N}{Nad_N + Km_{pdh}^{Nad_N}} \frac{Km_{pdh}^{Accoa_{NM}}}{Accoa_{NM} + Km_{pdh}^{Accoa_{NM}}} \tag{S205}$$

TCA cycle (TCA)

$$v_{tca}^{Accoa_{NM}} = Vmax_{tca}^{Accoa_{NM}} \frac{Accoa_{NM}}{Accoa_{NM} + Km_{tca}^{Accoa_{NM}}} \frac{Adp_N}{Adp_N + Km_{tca}^{Adp_N}} \frac{Nad_N}{Nad_N + Km_{tca}^{Nad_N}} \frac{Fad_N}{Fad_N + Km_{tca}^{Fad_N}} \frac{Phos_N}{Phos_N + Km_{tca}^{Phos_N}} \frac{Pyr_N}{Pyr_N + Km_{tca}^{Pyr_N}} \tag{S206}$$

β-hydroxybutyrate degradation (BHBDEG)

$$v_{bhb\deg}^{Bhb_N} = Vmax_{bhb\deg}^{Bhb_N} \frac{Bhb_N}{Bhb_N + Km_{bhb\deg}^{Bhb_N}} \frac{Nad_N}{Nad_N + Km_{bhb\deg}^{Nad_N}} \frac{Atp_N}{Atp_N + Km_{bhb\deg}^{Atp_N}} \tag{S207}$$

β-hydroxybutyrate transporter (BHBT)

$$v_{bhbt}^{Bhb_{BN}} = Vdif_{bhbt}^{Bhb_{BN}} \frac{Bhb_B - Bhb_N}{1 + \frac{Bhb_B}{Kdif_{bhbt}^{Bhb_{BN}}} + \frac{Bhb_N}{Kdif_{bhbt}^{Bhb_N}}} \quad \text{(S208)}$$

ATP synthesis from FADH (ATPSYNF)

$$v_{atpsynf}^{Fadh_N} = Vmax_{atpsynf}^{Fadh_N} \frac{Fadh_N}{Fadh_N + Km_{atpsynf}^{Fadh_N}} \frac{Adp_N}{Adp_N + Km_{atpsynf}^{Adp_N}} \quad \text{(S209)}$$

ATP synthesis from NADH (ATPSYNN)

$$v_{atpsynn}^{Nadh_N} = Vmax_{atpsynn}^{Nadh_N} \frac{Nadh_N}{Nadh_N + Km_{atpsynn}^{Nadh_N}} \frac{Adp_N}{Adp_N + Km_{atpsynn}^{Adp_N}} \quad \text{(S210)}$$

ATP utilization (ATPUSE)

$$v_{atpuse}^{Atp_N} = Vmax_{atpuse}^{Atp_N} \frac{Atp_N}{Atp_N + Km_{atpuse}^{Atp_N}} \quad \text{(S211)}$$

Uridine diphosphate kinase (UDP regeneration to UDP (UDPREG))

$$v_{udpreg}^{Udp_N} = Vmax_{udpreg}^{Udp_N} \frac{Udp_N Atp_N - \frac{Utp_N Adp_N}{Keq_{udpreg}^{Utp_N}}}{(1 + \frac{Udp_N}{Km_{udpreg}^{Udp_N}})(1 + \frac{Atp_N}{Km_{udpreg}^{Atp_N}}) + (1 + \frac{Utp_N}{Km_{udpreg}^{Utp_N}})(1 + \frac{Adp_N}{Km_{udpreg}^{Adp_N}}) - 1} \quad \text{(S212)}$$

UTP utilization (UTPUSE)

$$v_{utpuse}^{Utp_N} = Vmax_{utpuse}^{Utp_N} \frac{Utp_N}{Utp_N + Km_{utpuse}^{Utp_N}} \quad \text{(S213)}$$

NADH utilization (NADHUSE)

$$v_{nadhuse}^{Nadh_N} = Vmax_{nadhuse}^{Nadh_N} \frac{Nadh_N}{Nadh_N + Km_{nadhuse}^{Nadh_N}} \quad \text{(S214)}$$

FADH utilization (FADHUSE)

$$v_{fadhuse}^{Fadh_N} = Vmax_{fadhuse}^{Fadh_N} \frac{Fadh_N}{Fadh_N + Km_{fadhuse}^{Fadh_N}} \quad \text{(S215)}$$

Creatine kinase (CK)

$$v_{ck}^{Cre_N} = Vmax_{ck}^{Cre_N} \cdot \frac{Cre_N \cdot Atp_N - Crep_N \cdot Adp_N / Keq_{ck}^{Crep_N}}{(1 + \frac{Cre_N}{Km_{ck}^{Cre_N}})(1 + \frac{Atp_N}{Km_{ck}^{Atp_N}}) + (1 + \frac{Crep_N}{Km_{ck}^{Crep_N}})(1 + \frac{Adp_N}{Km_{ck}^{Adp_N}}) - 1} \quad \text{(S216)}$$

Cholesterol utilization (CHOLUSE)

$$v_{choluse}^{Chol_B} = k_{choluse}^{Chol_B} Chol_B \tag{S217}$$

**ODE in blood**

$$V_B \frac{dIns_B}{dt} = v_{inssyn}^{B} + v_{inssyn}^{Glc_B} + v_{inssyn}^{FFA_B} - v_{insdeg}^{Ins_B} \tag{S218}$$

$$V_B \frac{dGlc_B}{dt} = v_{meal}^{Glc_B} - v_{glut2}^{Glc_{BL}} - v_{glut4}^{Glc_{BM}} - v_{glut4}^{Glc_{BA}} - v_{glut2}^{Glc_{BG}} - v_{glut2}^{Glc_{BH}} - v_{glut3}^{Glc_{BN}} - v_{glutT}^{Glc_{BT}} \tag{S219}$$

$$V_B \frac{dPyr_B}{dt} = -v_{pyrt}^{Pyr_{BL}} - v_{pyrt}^{Pyr_{BM}} - v_{pyrt}^{Pyr_{BA}} - v_{pyrt}^{Pyr_{BG}} - v_{pyrt}^{Pyr_{BH}} - v_{pyrt}^{Pyr_{BN}} - v_{pyrt}^{Pyr_{BT}} \tag{S220}$$

$$V_B \frac{dLac_B}{dt} = -v_{lact}^{Lac_{BL}} - v_{lact}^{Lac_{BM}} - v_{lact}^{Lac_{BA}} - v_{lact}^{Lac_{BG}} - v_{lact}^{Lac_{BH}} - v_{lact}^{Lac_{BN}} - v_{lact}^{Lac_{BT}} \tag{S221}$$

$$V_B \frac{dAla_B}{dt} = -v_{alat}^{Ala_{BL}} - v_{alat}^{Ala_{BM}} - v_{alat}^{Ala_{BA}} - v_{alat}^{Ala_{BG}} - v_{alat}^{Ala_{BH}} - v_{alat}^{Ala_{BN}} - v_{alat}^{Ala_{BT}} \tag{S222}$$

$$\begin{aligned} V_B \frac{dGlyc_B}{dt} = & -v_{glyct}^{Glyc_{BL}} - v_{glyct}^{Glyc_{BM}} - v_{glyct}^{Glyc_{BA}} - v_{glyct}^{Glyc_{BG}} - v_{glyct}^{Glyc_{BH}} - v_{glyct}^{Glyc_{BN}} - v_{glyct}^{Glyc_{BT}} \\ & + v_{tgt}^{TG_{BL}} + v_{tgt}^{TG_{BM}} + v_{tgt}^{TG_{BA}} + v_{tgt}^{TG_{BG}} + v_{tgt}^{TG_{BH}} + v_{tgt}^{TG_{BN}} \end{aligned} \tag{S223}$$

$$V_B \frac{dTG_B}{dt} = v_{meal}^{TG_B} - v_{tgt}^{TG_{BL}} - v_{tgt}^{TG_{BM}} - v_{tgt}^{TG_{BA}} - v_{tgt}^{TG_{BG}} - v_{tgt}^{TG_{BH}} - v_{tgt}^{TG_{BN}} - v_{tgt}^{TG_{BT}} \tag{S224}$$

$$V_B \frac{dFFA_B}{dt} = -v_{ffat}^{FFA_{BL}} - v_{ffat}^{FFA_{BM}} - v_{ffat}^{FFA_{BA}} - v_{ffat}^{FFA_{BG}} - v_{ffat}^{FFA_{BH}} - v_{ffat}^{FFA_{BN}} - v_{ffat}^{FFA_{BT}} \tag{S225}$$

$$V_B \frac{dBhb_B}{dt} = -v_{bhbt}^{Bhb_{BL}} - v_{bhbt}^{Bhb_{BM}} - v_{bhbt}^{Bhb_{BA}} - v_{bhbt}^{Bhb_{BG}} - v_{bhbt}^{Bhb_{BH}} - v_{bhbt}^{Bhb_{BN}} \tag{S226}$$

$$V_B \frac{dChol_B}{dt} = -v_{cholt}^{Chol_{BL}} - v_{choluse}^{Chol_B} \tag{S227}$$

**ODE in liver**

$$V_L \frac{dGlc_L}{dt} = v_{glut2}^{Glc_{BL}} - v_{hk}^{Glc_L} + v_{g6pase}^{G6p_L} \tag{S228}$$

$$V_L \frac{dG6p_L}{dt} = v_{hk}^{Glc_L} - v_{g6pase}^{G6p_L} - v_{gs}^{G6p_L} + v_{gd}^{Glygn_L} - v_{pfk}^{G6p_L} + \frac{1}{2} v_{fbp}^{Gap_L} - v_{ppp}^{G6p_L} \tag{S229}$$

$$V_L \frac{dGlygn_L}{dt} = v_{gs}^{G6p_L} - v_{gd}^{Glygn_L} \tag{S230}$$

$$V_L \frac{dGap_L}{dt} = 2v_{pfk}^{G6p_L} - v_{fbp}^{Gap_L} - v_{pk}^{Gap_L} + v_{pepck}^{Pyr_L} - v_{g3pd}^{Gap_L} \tag{S231}$$

$$V_L \frac{dPyr_L}{dt} = v_{pk}^{Gap_L} - v_{pepck}^{Pyr_L} + v_{pryt}^{Pyr_{BL}} - v_{ldh}^{Pyr_L} - v_{alata}^{Pyr_L} - v_{pdh}^{Pyr_L} \tag{S232}$$

$$V_L \frac{dLac_L}{dt} = v_{lact}^{Pyr_{BL}} + v_{ldh}^{Pyr_L} \tag{S233}$$

$$V_L \frac{dAla_L}{dt} = v_{alat}^{Ala_{BL}} + v_{alata}^{Pyr_L} \tag{S234}$$

$$V_L \frac{dAccoa_{LM}}{dt} = v_{pdh}^{Pyr_L} - v_{tca}^{Accoa_{LM}} - v_{bhbsyn}^{Accoa_{LM}} + 8 \cdot v_{boxid}^{FFA_L} - v_{accoat}^{Accoa_{LMLC}} \tag{S235}$$

$$V_L \frac{dBhb_L}{dt} = \frac{1}{2} v_{bhbsyn}^{Accoa_{LM}} + v_{bhbt}^{Bhb_{BL}} \tag{S236}$$

$$V_L \frac{dAccoa_{LC}}{dt} = v_{accoat}^{Accoa_{LMLC}} - v_{lipog1}^{Accoa_{LC}} - v_{cholsyn1}^{Accoa_{LC}} \tag{S237}$$

$$V_L \frac{dMalcoa_L}{dt} = v_{lipog1}^{Accoa_{LC}} - v_{lipog2}^{Malcoa_L} \tag{S238}$$

$$V_L \frac{dHmgcoa_L}{dt} = \frac{1}{3} v_{cholsyn1}^{Accoa_{LC}} - v_{cholsyn2}^{Hmgcoa_L} \tag{S239}$$

$$V_L \frac{dChol_L}{dt} = \frac{1}{6} v_{cholsyn2}^{Hmgcoa_L} + v_{cholt}^{Chol_{BL}} \tag{S240}$$

$$V_L \frac{dFFA_L}{dt} = v_{ffat}^{FFA_{BL}} - 3 \cdot v_{tgsyn}^{FFA_L} + 3 \cdot v_{tgdeg}^{TG_L} - v_{boxid}^{FFA_L} + \frac{1}{8} v_{lipog2}^{Malcoa_L} \tag{S241}$$

$$V_L \frac{dGlyc_L}{dt} = v_{tgdeg}^{TG_L} + v_{glyct}^{Glyc_{BL}} - v_{glyck}^{Glyc_L} \tag{S242}$$

$$V_L \frac{dGlycp_L}{dt} = -v_{tgsyn}^{FFA_L} + v_{glyct}^{Glyc_L} + v_{g3pd}^{Gap_L} \tag{S243}$$

$$V_L \frac{dTG_L}{dt} = v_{tgsyn}^{FFA_L} - v_{tgdeg}^{TG_L} + v_{tgt}^{TG_L} \tag{S244}$$

$$\begin{aligned} V_L \frac{dAtp_L}{dt} = & -v_{hk}^{Glc_L} - v_{pfk}^{G6p_L} + 2v_{pk}^{Gap_L} - 2v_{pepck}^{Pyr_L} + v_{tca}^{Accoa_{LM}} - v_{accoat}^{Accoa_{LMLC}} - v_{glyk}^{Glyc_L} - v_{boxid}^{FFA_L} - v_{lipog1}^{Accoa_{LC}} \\ & + \frac{1}{8} v_{lipog2}^{Malcoa_{LC}} - 2v_{tgsyin}^{FFA_L} - 3v_{cholosyn2}^{Hmgcoa_L} + v_{atpsynf}^{Fadh_L} + 3v_{atpsynn}^{Nadh_L} - v_{ampreg}^{Amp_L} - v_{atpuse}^{Atp_L} - v_{nadhk}^{Nadh_L} \end{aligned} \tag{S245}$$

$$\begin{aligned} V_L \frac{dAdp_L}{dt} = & v_{hk}^{Glc_L} + v_{pfk}^{G6p_L} - 2 \cdot v_{pk}^{Gap_L} + 2 \cdot v_{pepck}^{Pyr_L} - v_{tca}^{Accoa_{LM}} + v_{accoat}^{Accoa_{LMLC}} + v_{glyk}^{Glyc_L} + v_{boxid}^{FFA_L} + v_{lipog1}^{Accoa_{LC}} \\ & - \frac{1}{8} v_{lipog2}^{Malcoa_{LC}} - v_{tgsyin}^{FFA_L} + 3 \cdot v_{cholosyn2}^{Hmgcoa_L} - v_{atpsynf}^{Fadh_L} - 3 \cdot v_{atpsynn}^{Nadh_L} + 2 \cdot v_{ampreg}^{Amp_L} + v_{atpuse}^{Atp_L} + v_{nadhk}^{Nadh_L} \end{aligned} \tag{S246}$$

$$V_L \frac{dAmp_L}{dt} = 3 \cdot v_{tgsyin}^{FFA_L} - v_{ampreg}^{Amp_L} \tag{S247}$$

$$V_L \frac{dNadh_L}{dt} = v_{pk}^{Gap_L} - v_{pepck}^{Pyr_L} - v_{ldh}^{Pyr_L} + v_{pdh}^{Pyr_L} + 3 \cdot v_{tca}^{Accoa_{LM}} - v_{g3pd}^{Gap_L} - \frac{1}{2} v_{bhbsyn}^{Accoa_{LM}} + 7v_{boxid}^{FFA_L} - v_{atpsynn}^{Nadh_L} - v_{nadhuse}^{Nadh_L} \tag{S248}$$

$$V_L \frac{dNad_L}{dt} = -v_{pk}^{Gap_L} + v_{pepck}^{Pyr_L} + {}_{ldh}^{Pyr_L} - v_{pdh}^{Pyr_L} - 3v_{tca}^{Accoa_{LM}} + v_{g3pd}^{Gap_L} + \frac{1}{2} v_{bhbsyn}^{Accoa_{LM}} - 7v_{boxid}^{FFA_L} + v_{atpsynn}^{Nadh_L} + v_{nadhuse}^{Nadh_L} \tag{S249}$$

$$V_L \frac{dFadh_L}{dt} = 7v_{boxid}^{FFA_L} + v_{tca}^{Accoa_{LM}} - \frac{1}{6} v_{cholsyn2}^{Hmgcoa_L} - v_{atpsynf}^{Fadh_L} - v_{fadhuse}^{Fadh_L} \tag{S250}$$

$$V_L \frac{dFad_L}{dt} = -7v_{boxid}^{FFA_L} - v_{tca}^{Accoa_{LM}} + \frac{1}{6} v_{cholsyn2}^{Hmgcoa_L} + v_{atpsynf}^{Fadh_L} + v_{fadhuse}^{Fadh_L} \tag{S251}$$

$$V_L \frac{dGtp_L}{dt} = -v_{pepck}^{Pyr_L} + v_{gdpreg}^{Gdp_L} \tag{S252}$$

$$V_L \frac{dGdp_L}{dt} = v_{pepck}^{Pyr_L} - v_{gdpreg}^{Gdp_L} \tag{S253}$$

$$V_L \frac{dUtp_L}{dt} = -v_{gs}^{G6p_L} + v_{udpreg}^{Udp_L} - v_{utpuse}^{Utp_L} \tag{S254}$$

$$V_L \frac{dUdp_L}{dt} = v_{gs}^{G6p_L} - v_{udpreg}^{Udp_L} + v_{utpuse}^{Utp_L} \tag{S255}$$

$$V_L \frac{dNadph_L}{dt} = -\frac{14}{8} v_{lipog2}^{Malcoa_L} - \frac{21}{6} v_{cholsyn2}^{Hmgcoa_L} + 2v_{ppp}^{G6p_L} + v_{nadhk}^{Nadh_L} - v_{nadphuse}^{Nadph_L} \tag{S256}$$

$$V_L \frac{dNadp_L}{dt} = \frac{14}{8} v_{lipog2}^{Malcoa_L} + \frac{21}{6} v_{cholsyn2}^{Hmgcoa_L} - 2v_{ppp}^{G6p_L} - v_{nadhk}^{Nadh_L} + v_{nadphuse}^{Nadph_L} \tag{S257}$$

$$V_L \frac{dCre_L}{dt} = -v_{ck}^{Cre_L} \tag{S258}$$

$$V_L \frac{dCrep_L}{dt} = v_{ck}^{Cre_L} \tag{S259}$$

**ODE in skeletal muscle**

$$V_M \frac{dGlc_M}{dt} = v_{glut4}^{Glc_{BM}} - v_{hk}^{Glc_M} \tag{S260}$$

$$V_M \frac{dG6p_M}{dt} = v_{hk}^{Glc_M} - v_{gs}^{G6p_M} + v_{gd}^{Glygn_M} - v_{pfk}^{G6p_M} \tag{S261}$$

$$V_M \frac{dGlygn_M}{dt} = v_{gs}^{G6p_M} - v_{gd}^{Glygn_M} \tag{S262}$$

$$V_M \frac{dGap_M}{dt} = 2v_{pfk}^{G6p_M} - v_{pk}^{Gap_M} - v_{g3pd}^{Gap_M} \tag{S263}$$

$$V_M \frac{dPyr_M}{dt} = v_{pk}^{Gap_M} + v_{pryt}^{Pyr_{BM}} - v_{ldh}^{Pyr_M} - v_{alata}^{Pyr_M} - v_{pdh}^{Pyr_M} \tag{S264}$$

$$V_M \frac{dLac_M}{dt} = v_{lact}^{Pyr_{BM}} + v_{ldh}^{Pyr_M} \tag{S265}$$

$$V_M \frac{dAla_M}{dt} = v_{alat}^{Ala_{BM}} + v_{alata}^{Pyr_M} \tag{S266}$$

$$V_M \frac{dAccoa_{MM}}{dt} = v_{pdh}^{Pyr_M} - v_{tca}^{Accoa_{MM}} + 8 \cdot v_{boxid}^{FFA_M} \tag{S267}$$

$$V_M \frac{dBhb_M}{dt} = 0 \tag{S268}$$

$$V_M \frac{dAccoa_{MC}}{dt} = v_{accoat}^{Accoa_{MMMC}} - v_{lipog1}^{Accoa_{MC}} \tag{S269}$$

$$V_M \frac{dMalcoa_M}{dt} = v_{lipog1}^{Accoa_{MC}} - v_{lipog2}^{Malcoa_M} \tag{S270}$$

$$V_M \frac{dHmgcoa_M}{dt} = 0 \tag{S271}$$

$$V_M \frac{dChol_M}{dt} = 0 \tag{S272}$$

$$V_M \frac{dFFA_M}{dt} = v^{FFA_{BM}}_{ffat} - 3 \cdot v^{FFA_M}_{tgsyn} + 3 \cdot v^{TG_M}_{tgdeg} - v^{FFA_M}_{boxid} \tag{S273}$$

$$V_M \frac{dGlyc_M}{dt} = v^{TG_M}_{tgdeg} + v^{Glyc_{BM}}_{glyct} - v^{Glyc_M}_{glyck} \tag{S274}$$

$$V_M \frac{dGlycp_M}{dt} = -v^{FFA_M}_{tgsyn} + v^{Glyc_M}_{glyct} + v^{Gap_M}_{g3pd} \tag{S275}$$

$$V_M \frac{dTG_M}{dt} = v^{FFA_M}_{tgsyn} - v^{TG_M}_{tgdeg} \tag{S276}$$

$$V_M \frac{dAtp_M}{dt} = -v^{Glc_M}_{hk} - v^{G6p_M}_{pfk} + 2v^{Gap_M}_{pk} + v^{Accoa_{MM}}_{tca} - v^{Glyc_M}_{glyk} - v^{FFA_M}_{boxid} - 2v^{FFA_M}_{tgsyin} + v^{Fadh_M}_{atpsynf} + 3v^{Nadh_M}_{atpsynn} - v^{Amp_M}_{ampreg} - v^{Atp_M}_{atpuse} \tag{S277}$$

$$V_M \frac{dAdp_M}{dt} = v^{Glc_M}_{hk} + v^{G6p_M}_{pfk} - 2 \cdot v^{Gap_M}_{pk} - v^{Accoa_{MM}}_{tca} + v^{Glyc_M}_{glyk} + v^{FFA_M}_{boxid} - v^{FFA_M}_{tgsyin} - v^{Fadh_M}_{atpsynf} - 3 \cdot v^{Nadh_M}_{atpsynn} + 2 \cdot v^{Amp_M}_{ampreg} + v^{Atp_M}_{atpuse} \tag{S278}$$

$$V_M \frac{dAmp_M}{dt} = 3 \cdot v^{FFA_M}_{tgsyin} - v^{Amp_M}_{ampreg} \tag{S279}$$

$$V_M \frac{dNadh_M}{dt} = v^{Gap_M}_{pk} - v^{Pyr_M}_{ldh} + v^{Pyr_M}_{pdh} + 3 \cdot v^{Accoa_{MM}}_{tca} - v^{Gap_M}_{g3pd} + 7v^{FFA_M}_{boxid} - v^{Nadh_M}_{atpsynn} - v^{Nadh_M}_{nadhuse} \tag{S280}$$

$$V_M \frac{dNad_M}{dt} = -v^{Gap_M}_{pk} +^{Pyr_M}_{ldh} - v^{Pyr_M}_{pdh} - 3v^{Accoa_{MM}}_{tca} + v^{Gap_M}_{g3pd} - 7v^{FFA_M}_{boxid} + v^{Nadh_M}_{atpsynn} + v^{Nadh_M}_{nadhuse} \tag{S281}$$

$$V_M \frac{dFadh_M}{dt} = 7v^{FFA_M}_{boxid} + v^{Accoa_{MM}}_{tca} - v^{Fadh_M}_{atpsynf} - v^{Fadh_M}_{fadhuse} \tag{S282}$$

$$V_M \frac{dFad_M}{dt} = -7v^{FFA_M}_{boxid} - v^{Accoa_{MM}}_{tca} + v^{Fadh_M}_{atpsynf} + v^{Fadh_M}_{fadhuse} \tag{S283}$$

$$V_M \frac{dGtp_M}{dt} = 0 \tag{S284}$$

$$V_M \frac{dGdp_M}{dt} = 0 \tag{S285}$$

$$V_M \frac{dUtp_M}{dt} = -v^{G6p_M}_{gs} + v^{Udp_M}_{udpreg} - v^{Utp_M}_{utpuse} \tag{S286}$$

$$V_M \frac{dUdp_M}{dt} = v^{G6p_M}_{gs} - v^{Udp_M}_{udpreg} + v^{Utp_M}_{utpuse} \tag{S297}$$

$$V_M \frac{dNadph_M}{dt} = 2v^{G6p_M}_{ppp} + v^{Nadh_M}_{nadhk} - v^{Nadph_M}_{nadphuse} \tag{S288}$$

$$V_M \frac{dNadp_M}{dt} = -2v^{G6p_M}_{ppp} - v^{Nadh_M}_{nadhk} + v^{Nadph_M}_{nadphuse} \tag{S289}$$

$$V_M \frac{dCre_M}{dt} = -v_{ck}^{Cre_M} \quad (S290)$$

$$V_M \frac{dCrep_M}{dt} = v_{ck}^{Cre_M} \quad (S291)$$

**ODE in adipose tissue**

$$V_A \frac{dGlc_A}{dt} = v_{glut4}^{Glc_{BA}} - v_{hk}^{Glc_A} \quad (S292)$$

$$V_A \frac{dG6p_A}{dt} = v_{hk}^{Glc_A} - v_{pfk}^{G6p_A} \quad (S293)$$

$$V_A \frac{dGlygn_A}{dt} = 0 \quad (S294)$$

$$V_A \frac{dGap_A}{dt} = 2v_{pfk}^{G6p_A} - v_{pk}^{Gap_A} - v_{g3pd}^{Gap_A} \quad (S295)$$

$$V_A \frac{dPyr_A}{dt} = v_{pk}^{Gap_A} + v_{pryt}^{Pyr_{BA}} - v_{ldh}^{Pyr_A} - v_{alata}^{Pyr_A} - v_{pdh}^{Pyr_A} \quad (S296)$$

$$V_A \frac{dLac_A}{dt} = v_{lact}^{Pyr_{BA}} + v_{ldh}^{Pyr_A} \quad (S297)$$

$$V_A \frac{dAla_A}{dt} = v_{alat}^{Ala_{BA}} + v_{alata}^{Pyr_A} \quad (S298)$$

$$V_A \frac{dAccoa_{AM}}{dt} = v_{pdh}^{Pyr_A} - v_{tca}^{Accoa_{AM}} + 8 \cdot v_{boxid}^{FFA_A} \quad (S299)$$

$$V_A \frac{dBhb_A}{dt} = 0 \quad (S300)$$

$$V_A \frac{dAccoa_{AC}}{dt} = v_{accoat}^{Accoa_{AMAC}} - v_{lipog1}^{Accoa_{AC}} \quad (S301)$$

$$V_A \frac{dMalcoa_A}{dt} = v_{lipog1}^{Accoa_{AC}} - v_{lipog2}^{Malcoa_A} \quad (S302)$$

$$V_A \frac{dHmgcoa_A}{dt} = 0 \quad (S303)$$

$$V_A \frac{dChol_A}{dt} = 0 \quad (S304)$$

$$V_A \frac{dFFA_A}{dt} = v_{ffat}^{FFA_{BA}} - 3 \cdot v_{tgsyn}^{FFA_A} + 3 \cdot v_{tgdeg}^{TG_A} - v_{boxid}^{FFA_A} \quad (S305)$$

$$V_A \frac{dGlyc_A}{dt} = v_{tgdeg}^{TG_A} + v_{glyct}^{Glyc_{BA}} - v_{glyck}^{Glyc_A} \quad (S306)$$

$$V_A \frac{dGlycp_A}{dt} = -v_{tgsyn}^{FFA_A} + v_{glyct}^{Glyc_A} + v_{g3pd}^{Gap_A} \quad (S307)$$

$$V_A \frac{dTG_A}{dt} = v_{tgsyn}^{FFA_A} - v_{tgdeg}^{TG_A} \tag{S308}$$

$$\begin{aligned} V_A \frac{dAtp_A}{dt} = & -v_{hk}^{Glc_A} - v_{pfk}^{G6p_A} + 2v_{pk}^{Gap_A} + v_{tca}^{Accoa_{AM}} - v_{glyk}^{Glyc_A} - v_{boxid}^{FFA_A} - 2v_{tgsyin}^{FFA_A} + v_{atpsynf}^{Fadh_A} \\ & +3v_{atpsynn}^{Nadh_A} - v_{ampreg}^{Amp_A} - v_{atpuse}^{Atp_A} \end{aligned} \tag{S309}$$

$$\begin{aligned} V_A \frac{dAdp_A}{dt} = & \; v_{hk}^{Glc_A} + v_{pfk}^{G6p_A} - 2 \cdot v_{pk}^{Gap_A} - v_{tca}^{Accoa_{AM}} + v_{glyk}^{Glyc_A} + v_{boxid}^{FFA_A} - v_{tgsyin}^{FFA_A} - v_{atpsynf}^{Fadh_A} \\ & -3 \cdot v_{atpsynn}^{Nadh_A} + 2 \cdot v_{ampreg}^{Amp_A} + v_{atpuse}^{Atp_A} \end{aligned} \tag{S310}$$

$$V_A \frac{dAmp_A}{dt} = 3 \cdot v_{tgsyin}^{FFA_A} - v_{ampreg}^{Amp_A} \tag{S311}$$

$$V_A \frac{dNadh_A}{dt} = v_{pk}^{Gap_A} - v_{ldh}^{Pyr_A} + v_{pdh}^{Pyr_A} + 3 \cdot v_{tca}^{Accoa_{AM}} - v_{g3pd}^{Gap_A} + 7v_{boxid}^{FFA_A} - v_{atpsynn}^{Nadh_A} - v_{nadhuse}^{Nadh_A} \tag{S312}$$

$$V_A \frac{dNad_A}{dt} = -v_{pk}^{Gap_A} +_{ldh}^{Pyr_A} - v_{pdh}^{Pyr_A} - 3v_{tca}^{Accoa_{AM}} + v_{g3pd}^{Gap_A} - 7v_{boxid}^{FFA_A} + v_{atpsynn}^{Nadh_A} + v_{nadhuse}^{Nadh_A} \tag{S313}$$

$$V_A \frac{dFadh_A}{dt} = 7v_{boxid}^{FFA_A} + v_{tca}^{Accoa_{AM}} - v_{atpsynf}^{Fadh_A} - v_{fadhuse}^{Fadh_A} \tag{S314}$$

$$V_A \frac{dFad_A}{dt} = -7v_{boxid}^{FFA_A} - v_{tca}^{Accoa_{AM}} + v_{atpsynf}^{Fadh_A} + v_{fadhuse}^{Fadh_A} \tag{S315}$$

$$V_A \frac{dGtp_A}{dt} = 0 \tag{S316}$$

$$V_A \frac{dGdp_A}{dt} = 0 \tag{S317}$$

$$V_A \frac{dUtp_A}{dt} = 0 \tag{S318}$$

$$V_A \frac{dUdp_A}{dt} = 0 \tag{S319}$$

$$V_A \frac{dNadph_A}{dt} = 2v_{ppp}^{G6p_A} + v_{nadhk}^{Nadh_A} - v_{nadphuse}^{Nadph_A} \tag{S320}$$

$$V_A \frac{dNadp_A}{dt} = -2v_{ppp}^{G6p_A} - v_{nadhk}^{Nadh_A} + v_{nadphuse}^{Nadph_A} \tag{S321}$$

$$V_A \frac{dCre_A}{dt} = -v_{ck}^{Cre_A} \tag{S322}$$

$$V_A \frac{dCrep_A}{dt} = v_{ck}^{Cre_A} \tag{S323}$$

#### ODE in GI tract

$$V_G \frac{dGlc_G}{dt} = v_{glut2}^{Glc_{BG}} - v_{hk}^{Glc_G} \tag{S324}$$

$$V_G \frac{dG6p_G}{dt} = v_{hk}^{Glc_G} - v_{pfk}^{G6p_G} \tag{S325}$$

$$V_G \frac{dGap_G}{dt} = 2v_{pfk}^{G6p_G} - v_{pk}^{Gap_G} - v_{g3pd}^{Gap_G} \quad \text{(S326)}$$

$$V_G \frac{dGlygn_G}{dt} = 0 \quad \text{(S327)}$$

$$V_G \frac{dPyr_G}{dt} = v_{pk}^{Gap_G} + v_{pryt}^{Pyr_{BG}} - v_{ldh}^{Pyr_G} - v_{alata}^{Pyr_G} - v_{pdh}^{Pyr_G} \quad \text{(S328)}$$

$$V_G \frac{dLac_G}{dt} = v_{lact}^{Pyr_{BG}} + v_{ldh}^{Pyr_G} \quad \text{(S329)}$$

$$V_G \frac{dAla_G}{dt} = v_{alat}^{Ala_{BG}} + v_{alata}^{Pyr_G} \quad \text{(S330)}$$

$$V_G \frac{dAccoa_{GM}}{dt} = v_{pdh}^{Pyr_G} - v_{tca}^{Accoa_{GM}} + 8 \cdot v_{boxid}^{FFA_G} \quad \text{(S331)}$$

$$V_A \frac{dBhb_A}{dt} = 0 \quad \text{(S332)}$$

$$V_A \frac{dAccoa_{AC}}{dt} = v_{accoat}^{Accoa_{AMAC}} - v_{lipog1}^{Accoa_{AC}} \quad \text{(S333)}$$

$$V_A \frac{dMalcoa_A}{dt} = v_{lipog1}^{Accoa_{AC}} - v_{lipog2}^{Malcoa_A} \quad \text{(S334)}$$

$$V_A \frac{dHmgcoa_A}{dt} = 0 \quad \text{(S335)}$$

$$V_A \frac{dChol_A}{dt} = 0 \quad \text{(S336)}$$

$$V_G \frac{dFFA_G}{dt} = v_{ffat}^{FFA_{BG}} - 3 \cdot v_{tgsyn}^{FFA_G} + 3 \cdot v_{tgdeg}^{TG_G} - v_{boxid}^{FFA_G} \quad \text{(S337)}$$

$$V_G \frac{dGlyc_G}{dt} = v_{tgdeg}^{TG_G} + v_{glyct}^{Glyc_{BG}} - v_{glyck}^{Glyc_G} \quad \text{(S338)}$$

$$V_G \frac{dGlycp_G}{dt} = -v_{tgsyn}^{FFA_G} + v_{glyct}^{Glyc_G} + v_{g3pd}^{Gap_G} \quad \text{(S339)}$$

$$V_G \frac{dTG_G}{dt} = v_{tgsyn}^{FFA_G} - v_{tgdeg}^{TG_G} \quad \text{(S340)}$$

$$V_G \frac{dAtp_G}{dt} = -v_{hk}^{Glc_G} - v_{pfk}^{G6p_G} + 2v_{pk}^{Gap_G} + v_{tca}^{Accoa_{GM}} - v_{glyk}^{Glyc_G} - v_{boxid}^{FFA_G} - 2v_{tgsyin}^{FFA_G} + v_{atpsynf}^{Fadh_G} + 3v_{atpsynn}^{Nadh_G} - v_{ampreg}^{Amp_G} - v_{atpuse}^{Atp_G} \quad \text{(S341)}$$

$$V_G \frac{dAdp_G}{dt} = v_{hk}^{Glc_G} + v_{pfk}^{G6p_G} - 2 \cdot v_{pk}^{Gap_G} - v_{tca}^{Accoa_{GM}} + v_{glyk}^{Glyc_G} + v_{boxid}^{FFA_G} - v_{tgsyin}^{FFA_G} - v_{atpsynf}^{Fadh_G} - 3 \cdot v_{atpsynn}^{Nadh_G} + 2 \cdot v_{ampreg}^{Amp_G} + v_{atpuse}^{Atp_G} \quad \text{(S342)}$$

$$V_G \frac{dAmp_G}{dt} = 3 \cdot v_{tgsyin}^{FFA_G} - v_{ampreg}^{Amp_G} \quad \text{(S343)}$$

$$V_G \frac{dNadh_G}{dt} = v_{pk}^{Gap_G} - v_{ldh}^{Pyr_G} + v_{pdh}^{Pyr_G} + 3 \cdot v_{tca}^{Accoa_{GM}} - v_{g3pd}^{Gap_G} + 7v_{boxid}^{FFA_G} - v_{atpsynn}^{Nadh_G} - v_{nadhuse}^{Nadh_G} \quad \text{(S344)}$$

$$V_G \frac{dNad_G}{dt} = -v_{pk}^{Gap_G} + {}_{ldh}^{Pyr_G} - v_{pdh}^{Pyr_G} - 3v_{tca}^{Accoa_{GM}} + v_{g3pd}^{Gap_G} - 7v_{boxid}^{FFA_G} + v_{atpsynn}^{Nadh_G} + v_{nadhuse}^{Nadh_G} \quad \text{(S345)}$$

$$V_G \frac{dFadh_G}{dt} = 7v_{boxid}^{FFA_G} + v_{tca}^{Accoa_{GM}} - v_{atpsynf}^{Fadh_G} - v_{fadhuse}^{Fadh_G} \quad \text{(S346)}$$

$$V_G \frac{dFad_G}{dt} = -7v_{boxid}^{FFA_G} - v_{tca}^{Accoa_{GM}} + v_{atpsynf}^{Fadh_G} + v_{fadhuse}^{Fadh_G} \quad \text{(S347)}$$

$$V_G \frac{dGtp_G}{dt} = 0 \quad \text{(S348)}$$

$$V_G \frac{dGdp_G}{dt} = 0 \quad \text{(S349)}$$

$$V_G \frac{dUtp_G}{dt} = 0 \quad \text{(S350)}$$

$$V_G \frac{dUdp_G}{dt} = 0 \quad \text{(S351)}$$

$$V_G \frac{dNadph_G}{dt} = 2v_{ppp}^{G6p_G} + v_{nadhk}^{Nadh_G} - v_{nadphuse}^{Nadph_G} \quad \text{(S352)}$$

$$V_G \frac{dNadp_G}{dt} = -2v_{ppp}^{G6p_G} - v_{nadhk}^{Nadh_G} + v_{nadphuse}^{Nadph_G} \quad \text{(S353)}$$

$$V_G \frac{dCre_G}{dt} = -v_{ck}^{Cre_G} \quad \text{(S354)}$$

$$V_G \frac{dCrep_G}{dt} = v_{ck}^{Cre_G} \quad \text{(S355)}$$

**ODE in heart**

$$V_H \frac{dGlc_H}{dt} = v_{glut2}^{Glc_{BH}} - v_{hk}^{Glc_H} \quad \text{(S356)}$$

$$V_H \frac{dG6p_H}{dt} = v_{hk}^{Glc_H} - v_{gs}^{G6p_H} + v_{gd}^{Glygn_H} - v_{pfk}^{G6p_H} \quad \text{(S357)}$$

$$V_H \frac{dGlygn_H}{dt} = v_{gs}^{G6p_H} - v_{gd}^{Glygn_H} \quad \text{(S358)}$$

$$V_H \frac{dGap_H}{dt} = 2v_{pfk}^{G6p_H} - v_{pk}^{Gap_H} - v_{g3pd}^{Gap_H} \quad \text{(S359)}$$

$$V_H \frac{dPyr_H}{dt} = v_{pk}^{Gap_H} + v_{pryt}^{Pyr_{BH}} - v_{ldh}^{Pyr_H} - v_{alata}^{Pyr_H} - v_{pdh}^{Pyr_H} \quad \text{(S360)}$$

$$V_H \frac{dLac_H}{dt} = v_{lact}^{Pyr_{BH}} + v_{ldh}^{Pyr_H} \quad \text{(S361)}$$

$$V_H \frac{dAla_H}{dt} = v_{alat}^{Ala_{BH}} + v_{alata}^{Pyr_H} \quad \text{(S362)}$$

$$V_H \frac{dAccoa_{HM}}{dt} = v_{pdh}^{Pyr_H} - v_{tca}^{Accoa_{HM}} + 8 \cdot v_{boxid}^{FFA_H} \quad \text{(S363)}$$

$$V_H \frac{dBhb_H}{dt} = 0 \quad \text{(S364)}$$

$$V_H \frac{dAccoa_{HC}}{dt} = v_{accoat}^{Accoa_{HMHC}} - v_{lipog1}^{Accoa_{HC}} \quad \text{(S365)}$$

$$V_H \frac{dMalcoa_H}{dt} = v_{lipog1}^{Accoa_{HC}} - v_{lipog2}^{Malcoa_H} \quad \text{(S366)}$$

$$V_H \frac{dHmgcoa_H}{dt} = 0 \quad \text{(S367)}$$

$$V_H \frac{dChol_H}{dt} = 0 \quad \text{(S368)}$$

$$V_H \frac{dFFA_H}{dt} = v_{ffat}^{FFA_{BH}} - 3 \cdot v_{tgsyn}^{FFA_H} + 3 \cdot v_{tgdeg}^{TG_H} - v_{boxid}^{FFA_H} \quad \text{(S369)}$$

$$V_H \frac{dGlyc_M}{dt} = v_{tgdeg}^{TG_M} + v_{glyct}^{Glyc_{BM}} - v_{glyck}^{Glyc_M} \quad \text{(S370)}$$

$$V_H \frac{dGlycp_H}{dt} = -v_{tgsyn}^{FFA_H} + v_{glyct}^{Glyc_H} + v_{g3pd}^{Gap_H} \quad \text{(S371)}$$

$$V_H \frac{dTG_H}{dt} = v_{tgsyn}^{FFA_H} - v_{tgdeg}^{TG_H} \quad \text{(S372)}$$

$$V_H \frac{dAtp_H}{dt} = -v_{hk}^{Glc_H} - v_{pfk}^{G6p_H} + 2v_{pk}^{Gap_H} + v_{tca}^{Accoa_{HM}} - v_{glyk}^{Glyc_H} - v_{boxid}^{FFA_H} - 2v_{tgsyin}^{FFA_H} + v_{atpsynf}^{Fadh_H} + 3v_{atpsynn}^{Nadh_H} - v_{ampreg}^{Amp_H} - v_{atpuse}^{Atp_H} \quad \text{(S373)}$$

$$V_H \frac{dAdp_H}{dt} = v_{hk}^{Glc_H} + v_{pfk}^{G6p_M} - 2 \cdot v_{pk}^{Gap_H} - v_{tca}^{Accoa_{HM}} + v_{glyk}^{Glyc_H} + v_{boxid}^{FFA_H} - v_{tgsyin}^{FFA_H} - v_{atpsynf}^{Fadh_H} - 3 \cdot v_{atpsynn}^{Nadh_H} + 2 \cdot v_{ampreg}^{Amp_H} + v_{atpuse}^{Atp_H} \quad \text{(S374)}$$

$$V_H \frac{dAmp_H}{dt} = 3 \cdot v_{tgsyin}^{FFA_H} - v_{ampreg}^{Amp_H} \quad \text{(S375)}$$

$$V_H \frac{dNadh_H}{dt} = v_{pk}^{Gap_H} - v_{ldh}^{Pyr_H} + v_{pdh}^{Pyr_H} + 3 \cdot v_{tca}^{Accoa_{HM}} - v_{g3pd}^{Gap_H} + 7v_{boxid}^{FFA_H} - v_{atpsynn}^{Nadh_H} - v_{nadhuse}^{Nadh_H} \quad \text{(S376)}$$

$$V_H \frac{dNad_H}{dt} = -v_{pk}^{Gap_H} + {}_{ldh}^{Pyr_H} - v_{pdh}^{Pyr_H} - 3v_{tca}^{Accoa_{HM}} + v_{g3pd}^{Gap_H} - 7v_{boxid}^{FFA_H} + v_{atpsynn}^{Nadh_H} + v_{nadhuse}^{Nadh_M} \quad \text{(S377)}$$

$$V_H \frac{dFadh_H}{dt} = 7v_{boxid}^{FFA_H} + v_{tca}^{Accoa_{MM}} - v_{atpsynf}^{Fadh_H} - v_{fadhuse}^{Fadh_H} \quad \text{(S378)}$$

$$V_H \frac{dFad_H}{dt} = -7v_{boxid}^{FFA_H} - v_{tca}^{Accoa_{HM}} + v_{atpsynf}^{Fadh_H} + v_{fadhuse}^{Fadh_H} \quad \text{(S379)}$$

$$V_H \frac{dGtp_H}{dt} = 0 \quad \text{(S380)}$$

$$V_H \frac{dGdp_H}{dt} = 0 \quad \text{(S381)}$$

$$V_H \frac{dUtp_H}{dt} = -v_{gs}^{G6p_H} + v_{udpreg}^{Udp_M} - v_{utpuse}^{Utp_H} \quad \text{(S382)}$$

$$V_H \frac{dUdp_H}{dt} = v_{gs}^{G6p_H} - v_{udpreg}^{Udp_H} + v_{utpuse}^{Utp_H} \quad \text{(S383)}$$

$$V_H \frac{dNadph_H}{dt} = 2v_{ppp}^{G6p_H} + v_{nadhk}^{Nadh_H} - v_{nadphuse}^{Nadph_H} \quad \text{(S384)}$$

$$V_H \frac{dNadp_H}{dt} = -2v_{ppp}^{G6p_H} - v_{nadhk}^{Nadh_H} + v_{nadphuse}^{Nadph_H} \quad \text{(S385)}$$

$$V_H \frac{dCre_H}{dt} = -v_{ck}^{Cre_H} \quad \text{(S386)}$$

$$V_H \frac{dCrep_H}{dt} = v_{ck}^{Cre_H} \quad \text{(S387)}$$

**ODE in brain**

$$V_N \frac{dGlc_N}{dt} = v_{glut3}^{Glc_{BN}} - v_{hk}^{Glc_N} \quad \text{(S388)}$$

$$V_N \frac{dG6p_N}{dt} = v_{hk}^{Glc_N} - v_{gs}^{G6p_N} + v_{gd}^{Glygn_N} - v_{pfk}^{G6p_N} \quad \text{(S389)}$$

$$V_N \frac{dGlygn_N}{dt} = v_{gs}^{G6p_N} - v_{gd}^{Glygn_N} \quad \text{(S390)}$$

$$V_N \frac{dGap_N}{dt} = 2v_{pfk}^{G6p_N} - v_{pk}^{Gap_N} \quad \text{(S391)}$$

$$V_N \frac{dPyr_N}{dt} = v_{pk}^{Gap_N} + v_{pryt}^{Pyr_{BN}} - v_{ldh}^{Pyr_N} - v_{alata}^{Pyr_N} - v_{pdh}^{Pyr_N} \quad \text{(S392)}$$

$$V_N \frac{dLac_N}{dt} = v_{lact}^{Pyr_{BN}} + v_{ldh}^{Pyr_N} \quad \text{(S393)}$$

$$V_N \frac{dAla_N}{dt} = v_{alat}^{Ala_{BN}} + v_{alata}^{Pyr_N} \quad \text{(S394)}$$

$$V_N \frac{dAccoa_{NM}}{dt} = v_{pdh}^{Pyr_N} - v_{tca}^{Accoa_{NM}} + 2v_{bhbdeg}^{Bhb_N} \quad \text{(S395)}$$

$$V_N \frac{dBhb_N}{dt} = v_{bhbt}^{Bhb_{BN}} - v_{bhbdeg}^{Bhb_N} \quad \text{(S396)}$$

$$V_N \frac{dAccoa_{NC}}{dt} = v_{accoat}^{Accoa_{NMNC}} - v_{lipog1}^{Accoa_{NC}} \quad \text{(S397)}$$

$$V_N \frac{dMalcoa_N}{dt} = v_{lipog1}^{Accoa_{NC}} - v_{lipog2}^{Malcoa_N} \quad \text{(S398)}$$

$$V_N \frac{dHmgcoa_N}{dt} = 0 \quad \text{(S399)}$$

$$V_N \frac{dChol_N}{dt} = 0 \tag{S400}$$

$$V_N \frac{dFFA_N}{dt} = 0 \tag{S401}$$

$$V_N \frac{dGlyc_N}{dt} = 0 \tag{S402}$$

$$V_N \frac{dGlycp_N}{dt} = 0 \tag{S403}$$

$$V_N \frac{dTG_N}{dt} = 0 \tag{S404}$$

$$V_N \frac{dAtp_N}{dt} = -v_{hk}^{Glc_N} - v_{pfk}^{G6p_N} + 2v_{pk}^{Gap_N} + v_{tca}^{Accoa_{NM}} - v_{bhb\deg}^{Bhb_N} + v_{atpsynf}^{Fadh_N} + 3v_{atpsynn}^{Nadh_N} - v_{atpuse}^{Atp_N} \tag{S405}$$

$$V_N \frac{dAdp_N}{dt} = v_{hk}^{Glc_N} + v_{pfk}^{G6p_N} - 2 \cdot v_{pk}^{Gap_N} - v_{tca}^{Accoa_{NM}} - v_{bhb\deg}^{Bhb_N} - v_{atpsynf}^{Fadh_N} - 3 \cdot v_{atpsynn}^{Nadh_N} + v_{atpuse}^{Atp_N} \tag{S406}$$

$$V_N \frac{dAmp_N}{dt} = 0 \tag{S407}$$

$$V_N \frac{dNadh_N}{dt} = v_{pk}^{Gap_N} - v_{ldh}^{Pyr_N} + v_{pdh}^{Pyr_N} + 3 \cdot v_{tca}^{Accoa_{NM}} + v_{bhbdeg}^{Bhb_N} - v_{atpsynn}^{Nadh_N} - v_{nadhuse}^{Nadh_N} \tag{S408}$$

$$V_N \frac{dNad_N}{dt} = -v_{pk}^{Gap_N} +_{ldh}^{Pyr_N} - v_{pdh}^{Pyr_N} - 3v_{tca}^{Accoa_{NM}} - v_{bhbdeg}^{Bhb_N} + v_{atpsynn}^{Nadh_N} + v_{nadhuse}^{Nadh_N} \tag{S409}$$

$$V_N \frac{dFadh_N}{dt} = v_{tca}^{Accoa_{NM}} - v_{atpsynf}^{Fadh_N} - v_{fadhuse}^{Fadh_N} \tag{S410}$$

$$V_N \frac{dFad_N}{dt} = -v_{tca}^{Accoa_{NM}} + v_{atpsynf}^{Fadh_N} + v_{fadhuse}^{Fadh_N} \tag{S411}$$

$$V_N \frac{dGtp_N}{dt} = 0 \tag{S412}$$

$$V_N \frac{dGdp_N}{dt} = 0 \tag{S413}$$

$$V_N \frac{dUtp_N}{dt} = -v_{gs}^{G6p_N} + v_{udpreg}^{Udp_N} - v_{utpuse}^{Utp_N} \tag{S414}$$

$$V_N \frac{dUdp_N}{dt} = v_{gs}^{G6p_N} - v_{udpreg}^{Udp_N} + v_{utpuse}^{Utp_N} \tag{S415}$$

$$V_N \frac{dNadph_N}{dt} = 2v_{ppp}^{G6p_N} + v_{nadhk}^{Nadh_N} - v_{nadphuse}^{Nadph_N} \tag{S416}$$

$$V_N \frac{dNadp_N}{dt} = -2v_{ppp}^{G6p_N} - v_{nadhk}^{Nadh_N} + v_{nadphuse}^{Nadph_N} \tag{S417}$$

$$V_N \frac{dCre_N}{dt} = -v_{ck}^{Cre_N} \tag{S418}$$

$$V_N \frac{dCrep_N}{dt} = v_{ck}^{Cre_N} \tag{S419}$$
