## Supplemental Table 1 for "Virtual metabolic human dynamic model for pathological analysis and therapy design for diabetes"

### Table S1 Kinetic parameter list

| Symbol | Definition | Value | Unit | Method | Reference |
|---|---|---|---|---|---|
| $V_B$ | blood volume | 5 | L | exp | [1] |
| $V_L$ | liver volume | 1.5 | L | exp | [1] |
| $V_M$ | blood volume | 20 | L | exp | [1] |
| $V_A$ | adipose tissue volume | 11 | L | exp | [1] |
| $V_G$ | GI tract volume | 0.3 | L | exp | [1] |
| $V_H$ | heart volume | 2 | L | exp | [1] |
| $V_N$ | brain volume | 1.5 | L | exp | [1] |
| Symbol | Variable name in program | Value | Unit | Method | Reference |
| $Glc_B^{meal}$ | tot_Glc_B | 4.10E+02 | mmol | exp | |
| $T_{delay}^{Glc_B}$ | t_delay_Glc_B | 5.00E+01 | min | est | |
| $TG_B^{meal}$ | tot_TG_B | 3.80E+01 | mmol | exp | |
| $T_{delay}^{TG_B}$ | t_delay_TG_B | 2.40E+02 | min | est | |
| $k_{inssyn}^{B}$ | k_inssyn_B | 6.97E-05 | mmol/min | est | [2] |
| $Vmax_{inssyn}^{Glc_B}$ | Vmax_inssyn_Glc_B | 1.80E-03 | mmol/min | opt | |
| $K_{minssyn}^{Glc_B}$ | Km_inssyn_Glc_B | 7.50E+00 | mmol | est | |
| $n_{inssyn}^{Glc_B}$ | n_inssyn_Glc_B | 8.00E+00 | - | est | |
| $k_{inssyn}^{FFA_B}$ | k_inssyn_FFA_B | 1.00E-05 | L/min | est | |
| $k_{insdeg}^{Ins_B}$ | k_insdeg_Ins_B | 2.50E+03 | L/min | est | |
| $k_{choluse}^{Chol_B}$ | k_choluse_Chol_B | 2.00E-04 | L/min | est | |
| $\alpha_L^{Base}$ | alpha_base_L | 5.00E-02 | - | opt | |
| $\alpha_L^{Band}$ | alpha_band_L | 8.00E-01 | - | opt | |
| $Km^{Ins_{BL}}$ | Km_Ins_B_L | 8.00E-08 | mM | est | |
| $n^{Ins_{BL}}$ | n_Ins_B_L | 4.00E+00 | - | est | |
| $Vdif_{glut2}^{Glc_{BL}}$ | Vdif_glut2_Glc_B_L | 1.20E+01 | L/min | opt | |
| $Kdif_{glut2}^{Glc_{BL}}$ | Kdif_glut2_Glc_B_L | 1.70E+01 | mM | exp | [3] |
| $Kdif_{glut2}^{Glc_L}$ | Kdif_glut2_Glc_L | 1.70E+01 | mM | exp | [3] |
| $Vmax_{hk}^{Glc_L}$ | Vmax_hk_Glc_L | 2.93E+00 | mmol/min | opt | |
| $Km_{hk}^{Glc_L}$ | Km_hk_Glc_L | 4.20E-01 | mM | exp | [4] |

| | | | | | |
|---|---|---|---|---|---|
| $Ki_{hk}^{G6p_L}$ | Ki_hk_G6p_L | 5.00E-01 | mM | exp | [4] |
| $Km_{hk}^{Atp_L}$ | Km_hk_Atp_L | 2.09E+00 | mM | exp | [4] |
| $Ki_{hk}^{Atp_L}$ | Ki_hk_Atp_L | 1.90E-01 | mM | exp | [4] |
| $Vmax_{g6pase}^{G6p_L}$ | Vmax_ppp_G6p_L | 1.00E+00 | mmol/min | est | |
| $Km_{ppp}^{G6p_L}$ | Km_ppp_G6p_L | 1.00E-01 | mM | est | |
| $Km_{ppp}^{Nadp_L}$ | Km_ppp_Nadp_L | 1.00E-01 | mM | est | |
| $Vmax_{g6pase}^{G6p_L}$ | Vmax_g6pase_G6p_L | 1.20E+01 | mmol/min | opt | |
| $Km_{g6pase}^{G6p_L}$ | Km_g6pase_G6p_L | 2.41E+00 | mM | exp | [5, 6] |
| $Vmax_{gs}^{G6p_L}$ | Vmax_gs_G6p_L | 1.10E+01 | mmol/min | opt | |
| $Km_{gs}^{G6p_X}$ | Km_gs_G6p_L | 5.00E-02 | mM | est | [7] |
| $n_{gs}^{G6p_L}$ | n_gs_G6p_L | 1.00E+00 | - | est | |
| $Km_{gs}^{Utp_L}$ | Km_gs_Utp_L | 4.80E-02 | mM | exp | [8] |
| $Glygn_L^{max}$ | Glygn_L_max | 3.75E+02 | mM | est | [9] |
| $Km_{gs}^{Glygn_L}$ | Km_gs_Glygn_L | 7.50E+01 | mM | est | |
| $Vmax_{gd}^{Glygn_L}$ | Vmax_gd_Glygn_L | 2.80E+00 | mmol/min | opt | |
| $Km_{gd}^{Glygn_L}$ | Km_gd_Glygn_L | 1.00E+01 | mM | est | [7] |
| $Km_{gd}^{Phos_L}$ | Km_gd_Phos_L | 4.00E+00 | mM | exp | [10] |
| $Vmax_{pfk}^{G6p_L}$ | Vmax_pfk_G6p_L | 1.26E+01 | mmol/min | opt | |
| $Km_{pfk}^{G6p_L}$ | Km_pfk_G6p_L | 5.00E-03 | mM | est | [7] |
| $Km_{pfk}^{Atp_L}$ | Km_pfk_Atp_L | 4.25E-02 | mM | exp | [11] |
| $Ki_{pfk}^{Atp_L}$ | Ki_pfk_Atp_L | 2.10E+00 | mM | exp | [12] |
| $Km_{pfk}^{Adp_L}$ | Km_pfk_Adp_L | 8.36E-02 | mM | exp | [11, 12] |
| $Ki_{pfk}^{Gap_L}$ | Ki_pfk_Gap_L | 2.07E-02 | mM | exp | [13] |
| $b_{pfk}^{Gap_L}$ | b_pfk_Gap_L | 7.50E-01 | - | est | [7] |
| $Vmax_{fbp}^{Gap_L}$ | Vmax_fbp_Gap_L | 8.00E+00 | mmol/min | opt | |
| $Km_{fbp}^{Gap_L}$ | Km_fbp_Gap_L | 2.50E-01 | mM | est | [7] |
| $Vmax_{pk}^{Gap_L}$ | Vmax_pk_Gap_L | 2.75E+02 | mmol/min | | |

| | | | | | |
|---|---|---|---|---|---|
| $Km_{pk}^{Gap_L}$ | Km_pk_Gap_L | 2.40E-01 | mM | est | [7] |
| $b_{pk}^{Accoa_{LM}}$ | b_pk_Accoa_LM | 8.00E-01 | - | exp | [14] |
| $Ki_{pk}^{Accoa_{LM}}$ | Ki_pk_Accoa_LM | 3.00E-02 | mM | exp | [14] |
| $Km_{pk}^{Adp_L}$ | Km_pk_Adp_L | 2.40E-01 | mM | exp | [15] |
| $Vmax_{pepck}^{Pyr_L}$ | Vmax_pepck_Pyr_L | 3.30E+01 | mM | opt | |
| $Km_{pepck}^{Pyr_X}$ | Km_pepck_Pyr_L | 2.44E+00 | mM | est | |
| $Km_{pepck}^{Atp_L}$ | Km_pepck_Atp_L | 1.00E-02 | mM | est | [7] |
| $Km_{pepck}^{Gtp_L}$ | Km_pepck_Gtp_L | 6.40E-02 | mM | exp | [16] |
| $Vdif_{pyrt}^{Pyr_{BL}}$ | Vdif_pyrt_Pyr_B_L | 0.00E+00 | L/min | est | |
| $Kdif_{pyrt}^{Pyr_{BL}}$ | Kdif_pyrt_Pyr_B_L | 1.00E+00 | mM | est | |
| $Kdif_{pyrt}^{Pyr_L}$ | Kdif_pyrt_Pyr_L | 1.00E+00 | mM | est | |
| $Vdif_{lact}^{Lac_{BL}}$ | Vdif_lact_Lac_B_L | 1.20E+01 | L/min | opt | |
| $Kdif_{lact}^{Lac_L}$ | Kdif_lact_Lac_L | 2.42E+00 | mM | exp | [17] |
| $Kdif_{lact}^{Lac_{BL}}$ | Kdif_lact_Lac_B_L | 2.42E+00 | mM | exp | [17] |
| $Vmax_{ldh}^{Pyr_L}$ | Vmax_ldh_Pyr_L | 1.00E+01 | $L^2$/mmol/min | opt | |
| $Km_{ldh}^{Pyr_L}$ | Km_ldh_Pyr_L | 1.50E-01 | mM | exp | [18] |
| $Km_{ldh}^{Nadh_L}$ | Km_ldh_Nadh_L | 1.50E-02 | mM | exp | [18] |
| $Km_{ldh}^{Lac_L}$ | Km_ldh_Lac_L | 3.60E+01 | mM | exp | [19] |
| $Km_{ldh}^{Nad_L}$ | Km_ldh_Nad_L | 1.10E-01 | mM | exp | [18] |
| $Keq_{ldh}^{Lac_L}$ | Keq_ldh_Lac_L | 6.00E-01 | - | est | |
| $Vdif_{alat}^{Ala_{BL}}$ | Vdif_alat_Ala_B_L | 1.20E+01 | L/min | est | |
| $Kdif_{alat}^{Ala_L}$ | Kdif_alat_Ala_L | 2.42E+00 | mM | est | |
| $Kdif_{alat}^{Ala_{BL}}$ | Kdif_alat_Ala_B_L | 2.42E+00 | mM | est | |
| $Vmax_{alata}^{Pyr_L}$ | Vmax_alata_Pyr_L | 4.00E+00 | L/min | opt | |
| $Km_{alata}^{Pyr_L}$ | Km_alata_Pyr_L | 1.50E-01 | mM | est | |
| $Km_{alata}^{Ala_L}$ | Km_alata_Ala_L | 3.60E+01 | mM | est | |
| $Keq_{alata}^{Ala_L}$ | Keq_alata_Ala_L | 1.00E+00 | - | est | |
| $Vmax_{pdh}^{Pyr_L}$ | Vmax_pdh_Pyr_L | 1.00E+00 | mmol/min | opt | |

| | | | | | |
|---|---|---|---|---|---|
| $Km_{pdh}^{Pyr_L}$ | Km_pdh_Pyr_L | 5.40E-01 | mM | est | [7] |
| $Km_{pdh}^{Accoa_{LM}}$ | Ki_pdh_Accoa_LM | 3.50E-02 | mM | exp | [20] |
| $Km_{pdh}^{Nad_L}$ | Km_pdh_Nad_L | 1.00E-02 | mM | opt | |
| $Vmax_{tca}^{Accoa_{LM}}$ | Vmax_tca_Accoa_LM | 1.38E+01 | mmol/min | opt | |
| $Km_{tca}^{Accoa_{LM}}$ | Km_tca_Accoa_LM | 4.00E-04 | mM | exp | [21] |
| $Km_{tca}^{Phos_L}$ | Km_tca_Phos_L | 1.00E+00 | mM | opt | |
| $Km_{tca}^{Adp_L}$ | Km_tca_Adp_L | 4.10E-01 | mM | exp | [22] |
| $Km_{tca}^{Pyr_L}$ | Km_tca_Pyr_L | 3.50E-01 | mM | opt | |
| $Km_{tca}^{Nad_L}$ | Km_tca_Nad_L | 1.00E-01 | mM | est | |
| $Km_{tca}^{Fad_L}$ | Km_tca_Fad_L | 1.00E-01 | mM | est | |
| $Vdif_{ffat}^{FFA_{BL}}$ | Vdif_ffat_FFA_B_L | 1.71E+00 | L/min | opt | |
| $Kdif_{ffat}^{FFA_{BL}}$ | Kdif_ffat_FFA_B_L | 2.00E-01 | mM | est | [7] |
| $Kdif_{ffat}^{FFA_L}$ | Kdif_ffat_FFA_L | 2.00E-01 | mM | est | [7] |
| $Vmax_{ffat}^{FFA_{BL}}$ | Vmax_ffat_FFA_B_L | 0.00E+00 | mmol/min | opt | |
| $Km_{ffat}^{FFA_{BL}}$ | Km_ffat_FFA_B_L | 2.00E-03 | mM | est | [7] |
| $Vmax_{tgsyn}^{FFA_L}$ | Vmax_tgsyn_FFA_L | 4.00E-01 | mmol/min | opt | |
| $Km_{tgsyn}^{FFA_L}$ | Km_tgsyn_FFA_L | 6.45E-01 | mM | est | [7] |
| $Km_{tgsyn}^{Glycp_L}$ | Km_tgsyn_Glycp_L | 4.60E-01 | mM | exp | [23] |
| $Km_{tgsyn}^{Atp_L}$ | Km_tgsyn_Atp_L | 1.00E-01 | mM | opt | |
| $TG_L^{max}$ | TG_L_max | 0.00E+00 | mM | est | |
| $Km_{tgsyn}^{TG_L}$ | Km_tgsyn_TG_L | 0.00E+00 | mM | est | |
| $Vmax_{tgdeg}^{TG_L}$ | Vmax_tgdeg_TG_L | 9.00E-02 | mmol/min | opt | |
| $Km_{tgdeg}^{TG_L}$ | Km_tgdeg_TG_L | 5.07E+01 | mM | est | [7] |
| $Vdif_{glyct}^{Glyc_{BL}}$ | Vdif_glyct_Glyc_B_L | 4.00E+00 | L/min | opt | |
| $Kdif_{glyct}^{Glyc_{BL}}$ | Kdif_glyct_Glyc_B_L | 2.70E-01 | mM | exp | [24] |
| $Kdif_{glyct}^{Glyc_L}$ | Kdif_glyct_Glyc_L | 2.70E-01 | mM | exp | [24] |
| $Vmax_{glyk}^{Glyc_L}$ | Vmax_glyk_Glyc_L | 5.00E-01 | mmol/min | opt | |
| $Km_{glyk}^{Glyc_L}$ | Km_glyk_Glyc_L | 4.00E-02 | mM | exp | [24] |

| | | | | | |
|---|---|---|---|---|---|
| $Km_{glyk}^{Atp_L}$ | Km_glyk_Atp_L | 5.80E-02 | mM | exp | [25] |
| $Ki_{glyk}^{Gap_L}$ | Ki_glyk_Gap_L | 5.80E-01 | mM | exp | [25] |
| $Vdif_{tgt}^{TG_{BL}}$ | Vdif_tgt_TG_B_L | 0.00E+00 | mmol/min | est | |
| $Kdif_{tgt}^{TG_{BL}}$ | Kdif_tgt_TG_B_L | 1.00E+00 | mM | est | [7] |
| $Keq_{tgt}^{TG_L}$ | Keq_tgt_TG_L | 3.38E+01 | - | est | [7] |
| $Vmax_{tgt}^{TG_{BL}}$ | Vmax_tgt_TG_B_L | 4.20E-01 | mmol/min | opt | |
| $Km_{tgt}^{TG_L}$ | Km_tgt_TG_L | 3.38E+01 | mM | est | [7] |
| $Vmax_{g3pd}^{Gap_L}$ | Vmax_g3pd_Gap_L | 5.00E+01 | $L^2$/mmol/min | opt | |
| $Km_{g3pd}^{Gap_L}$ | Km_g3pd_Gap_L | 1.60E-01 | mM | exp | [26] |
| $Km_{g3pd}^{Glycp_L}$ | Km_g3pd_Glycp_L | 2.20E-01 | mM | exp | [26] |
| $Km_{g3pd}^{Nadh_L}$ | Km_g3pd_Nadh_L | 8.00E-03 | mM | exp | [26] |
| $Km_{g3pd}^{Nad_L}$ | Km_g3pd_Nad_L | 1.30E-02 | mM | exp | [26] |
| $Keq_{g3pd}^{Glycp_L}$ | Keq_g3pd_Glycp_L | 1.00E+00 | - | opt | |
| $Vmax_{boxid}^{FFA_L}$ | Vmax_boxid_FFA_L | 8.88E-01 | mmol/min | opt | |
| $Km_{boxid}^{FFA_L}$ | Km_boxid_FFA_L | 5.00E-03 | mM | exp | [27-29] |
| $Km_{boxid}^{Atp_L}$ | Km_boxid_Atp_L | 8.70E-02 | mM | exp | [30] |
| $Ki_{boxid}^{Accoa_{LM}}$ | Ki_boxid_Accoa_LM | 1.20E-01 | mM | | [31] |
| $Ki_{boxid}^{Malcoa_L}$ | Ki_boxid_Malcoa_L | 1.00E+02 | mM | opt | |
| $Km_{boxid}^{Nad_L}$ | Km_boxid_Nad_L | 1.00E-01 | mM | est | |
| $Km_{boxid}^{Fad_L}$ | Km_boxid_Fad_L | 1.00E-01 | mM | est | |
| $Vmax_{accoat}^{Accoa_{LMLC}}$ | Vmax_accoat_Accoa_LM_LC | 1.00E+01 | mmol/min | est | |
| $Km_{accoat}^{Accoa_{LM}}$ | Km_accoat_Accoa_LM | 5.80E-02 | mM | opt | |
| $Km_{accoat}^{Atp_L}$ | Km_accoat_Atp_L | 1.00E-01 | mM | est | |
| $Km_{accoat}^{Pyr_L}$ | Km_accoat_Pyr_L | 1.00E+01 | mM | est | |
| $Vmax_{bhbsyn}^{Accoa_{LM}}$ | Vmax_bhbsyn_Accoa_LM | 1.00E+02 | mmol/min | opt | |
| $Km_{bhbsyn}^{Accoa_{LM}}$ | Km_bhbsyn_Accoa_LM | 1.00E-01 | mM | est | |
| $Km_{bhbsyn}^{Nadh_L}$ | Km_bhbsyn_Nadh_L | 1.00E-01 | mM | est | |
| $n_{bhbsyn}^{Pyr_L}$ | n_bhbsyn_Pyr_L | 4.00E+00 | - | est | |

| | | | | |
|---|---|---|---|---|
| $Ki_{bhbsyn}^{Pyr_L}$ | Ki_bhbsyn_Pyr_L | 5.00E-02 | mM | opt |
| $Vmax_{bhbdeg}^{Bhb_L}$ | Vmax_bhbdeg_Bhb_L | 0.00E+00 | mmol/min | est |
| $Km_{bhbdeg}^{Bhb_L}$ | Km_bhbdeg_Bhb_L | 1.00E-01 | mM | est |
| $Km_{bhbdeg}^{Atp_L}$ | Km_bhbdeg_Atp_L | 1.00E-01 | mM | est |
| $Km_{bhbdeg}^{Nad_L}$ | Km_bhbdeg_Nad_L | 1.00E-01 | mM | est |
| $Vdif_{bhbt}^{Bhb_{BL}}$ | Vdif_bhbt_Bhb_B_L | 1.00E+01 | L/min | est |
| $Kdif_{bhbt}^{Bhb_{BL}}$ | Kdif_bhbt_Bhb_B_L | 1.00E+00 | mM | est |
| $Kdif_{bhbt}^{Bhb_L}$ | Kdif_bhbt_Bhb_L | 1.00E+00 | mM | est |
| $Vmax_{lipog1}^{Accoa_{LC}}$ | Vmax_lipog1_Accoa_LC | 1.20E-01 | mmol/min | opt |
| $Km_{lipog1}^{Accoa_{LC}}$ | Km_lipog1_Accoa_LC | 1.00E+01 | mM | est |
| $Km_{lipog1}^{Atp_L}$ | Km_lipog1_Atp_L | 1.00E-01 | mM | est |
| $Vmax_{lipog2}^{Malcoa_L}$ | Vmax_lipog2_Malcoa_L | 1.00E+00 | mmol/min | est |
| $Km_{lipog2}^{Malcoa_L}$ | Km_lipog2_Malcoa_L | 1.00E-02 | mM | est |
| $Km_{lipog2}^{Adp_L}$ | Km_lipog2_Adp_L | 1.00E-01 | mM | est |
| $Km_{lipog2}^{Nadph_L}$ | Km_lipog2_Nadph_L | 1.00E-02 | mM | est |
| $Vmax_{cholsyn1}^{Accoa_{LC}}$ | Vmax_cholsyn1_Accoa_LC | 2.00E-02 | mmol/min | est |
| $Km_{cholsyn1}^{Accoa_{LC}}$ | Km_cholsyn1_Accoa_LC | 1.00E-01 | mM | est |
| $Vmax_{cholsyn2}^{Hmgcoa_L}$ | Vmax_cholsyn2_Hmgcoa_L | 9.00E-01 | mmol/min | est |
| $Km_{cholsyn2}^{Hmgcoa_L}$ | Km_cholsyn2_Hmgcoa_L | 1.00E-01 | mM | est |
| $Km_{cholsyn2}^{Atp_L}$ | Km_cholsyn2_Atp_L | 1.00E-01 | mM | est |
| $Km_{cholsyn2}^{Fadh_L}$ | Km_cholsyn2_Fadh_L | 1.00E-01 | mM | est |
| $Km_{cholsyn2}^{Nadph_L}$ | Km_cholsyn2_Nadph_L | 1.00E-01 | mM | est |
| $Vmax_{cholt}^{Chol_{BL}}$ | Vmax_cholt_Chol_L | 1.00E+00 | mmol/min | est |
| $Km_{cholt}^{Chol_L}$ | Km_cholt_Chol_L | 1.00E-01 | mM | est |
| $Vmax_{atpsynf}^{Fadh_L}$ | Vmax_atpsynf_Fadh_L | 3.94E+00 | mmol/min | opt |
| $Km_{atpsynf}^{Fadh_L}$ | Km_atpsynf_Fadh_L | 1.00E-01 | mM | est |
| $Km_{atpsynf}^{Adp_L}$ | Km_atpsynf_Adp_L | 1.00E-01 | mM | est |
| $Vmax_{atpsynn}^{Nadh_L}$ | Vmax_atpsynn_Nadh_L | 6.80E+00 | mmol/min | est |

| | | | | | |
|---|---|---|---|---|---|
| $Km^{Nadh_L}_{atpsynn}$ | Km_atpsynn_Nadh_L | 1.00E-01 | mM | est | |
| $Km^{Adp_L}_{atpsynn}$ | Km_atpsynn_Adp_L | 1.00E-01 | mM | est | |
| $Vmax^{Atp_L}_{atpuse}$ | Vmax_atpuse_Atp_L | 5.00E+00 | mmol/min | opt | |
| $Km^{Atp_L}_{atpuse}$ | Km_atpuse_Atp_L | 2.50E+00 | mM | opt | |
| $Vmax^{Amp_L}_{ampreg}$ | Vmax_ampreg_Amp_L | 1.00E+01 | mmol/min | est | |
| $Km^{Amp_L}_{ampreg}$ | Km_ampreg_Amp_L | 8.00E-02 | mM | exp | [32] |
| $Km^{Atp_L}_{ampreg}$ | Km_ampreg_Atp_L | 9.00E-02 | mM | exp | [32] |
| $Km^{Adp_L}_{ampreg}$ | Km_ampreg_Adp_L | 1.10E-01 | mM | exp | [32] |
| $Vmax^{Nadh_L}_{nadhk}$ | Vmax_nadhk_Nadh_L | 0.00E+00 | mmol/min | est | |
| $Km^{Nadh_L}_{nadhk}$ | Km_nadhk_Nadh_L | 1.00E-01 | mM | est | |
| $Km^{Atp_L}_{nadhk}$ | Km_nadhk_Atp_L | 0.00E+00 | mM | est | |
| $Vmax^{Nadh_L}_{nadhuse}$ | Vmax_nadhuse_Nadh_L | 4.00E-01 | mmol/min | est | |
| $Km^{Nadh_L}_{nadhuse}$ | Km_nadhuse_Nadh_L | 1.00E-01 | mM | est | |
| $Vmax^{Gdp_L}_{gdpreg}$ | Vmax_gdpreg_Gdp_L | 8.00E+02 | $L^2$/mmol/min | est | |
| $Km^{Gdp_L}_{gdpreg}$ | Km_gdpreg_Gdp_L | 3.10E-02 | mM | exp | [33] |
| $Km^{Atp_L}_{gdpreg}$ | Km_gdpreg_Atp_L | 1.33E+00 | mM | exp | [33] |
| $Km^{Gtp_L}_{gdpreg}$ | Km_gdpreg_Gtp_L | 1.50E-01 | mM | exp | [34] |
| $Km^{Adp_L}_{gdpreg}$ | Km_gdpreg_Adp_L | 4.20E-02 | mM | exp | [33] |
| $Keq^{Gtp_L}_{gdpreg}$ | Keq_gdpreg_Gtp_L | 1.00E+03 | - | opt | |
| $Vmax^{Gtp_L}_{gtpuse}$ | Vmax_gtpuse_Gtp_L | 0.00E+00 | mmol/min | est | |
| $Km^{Gtp_L}_{gtpuse}$ | Km_gtpuse_Gtp_L | 1.00E-01 | mM | est | |
| $Vmax^{Udp_X}_{udpreg}$ | Vmax_udpreg_Udp_L | 4.00E+01 | $L^2$/mmol/min | est | |
| $Km^{Udp_L}_{udpreg}$ | Km_udpreg_Udp_L | 1.90E-01 | mM | exp | [33] |
| $Km^{Atp_L}_{udpreg}$ | Km_udpreg_Atp_L | 1.33E+00 | mM | exp | [33] |
| $Km^{Utp_L}_{udpreg}$ | Km_udpreg_Utp_L | 1.60E+01 | mM | exp | [34] |
| $Km^{Adp_L}_{udpreg}$ | Km_udpreg_Adp_L | 4.20E-02 | mM | exp | [33] |
| $Keq^{Utp_L}_{udpreg}$ | Keq_udpreg_Utp_L | 1.00E+03 | - | est | |

| | | | | | |
|---|---|---|---|---|---|
| $Vmax_{utpuse}^{Utp_L}$ | Vmax_utpuse_Utp_L | 0.00E+00 | mmol/min | est | |
| $Km_{utpuse}^{Utp_L}$ | Km_utpuse_Utp_L | 1.00E-01 | mM | est | |
| $Vmax_{nadphuse}^{Nadph_L}$ | Vmax_nadphuse_Nadph_L | 0.00E+00 | mmol/min | est | |
| $Km_{nadphuse}^{Nadph_L}$ | Km_nadphuse_Nadph_L | 1.00E-01 | mM | est | |
| $Vmax_{fadhuse}^{Fadh_L}$ | Vmax_fadhuse_Fadh_L | 0.00E+00 | mmol/min | est | |
| $Km_{fadhuse}^{Fadh_L}$ | Km_fadhuse_Fadh_L | 1.00E-01 | mM | est | |
| $Vmax_{ck}^{Cre_L}$ | Vmax_ck_Cre_L | 0.00E+00 | $L^2$/mmol/min | est | |
| $Km_{ck}^{Cre_L}$ | Km_ck_Cre_L | 1.00E+00 | mM | est | |
| $Km_{ck}^{Atp_L}$ | Km_ck_Atp_L | 3.00E-02 | mM | est | |
| $Km_{ck}^{Crep_L}$ | Km_ck_Crep_L | 1.50E+01 | mM | est | |
| $Km_{ck}^{Adp_L}$ | Km_ck_Adp_L | 1.00E-01 | mM | est | |
| $Keq_{ck}^{Crep_L}$ | Keq_ck_Crep_L | 1.00E+00 | - | est | |
| $Phos_L$ | Phos_L | 5.00E+00 | mM | est | |
| $\alpha_M^{Base}$ | alpha_base_M | 5.00E-02 | - | opt | |
| $\alpha_M^{Band}$ | alpha_band_M | 8.50E-01 | - | opt | |
| $Km^{Ins_{BM}}$ | Km_Ins_B_M | 8.00E-08 | mM | est | |
| $n^{Ins_{BM}}$ | n_Ins_B_M | 4.00E+00 | - | est | |
| $Vdif_{glut4}^{Glc_{BM}}$ | Vdif_glut4_Glc_B_M | 1.20E+01 | L/min | opt | |
| $Kdif_{glut4}^{Glc_{BM}}$ | Kdif_glut4_Glc_B_M | 5.00E+00 | mM | exp | [3] |
| $Kdif_{glut4}^{Glc_M}$ | Kdif_glut4_Glc_M | 5.00E+00 | mM | exp | [3] |
| $Vmax_{hk}^{Glc_M}$ | Vmax_hk_Glc_M | 1.20E+01 | mmol/min | opt | |
| $Km_{hk}^{Glc_M}$ | Km_hk_Glc_M | 1.00E+00 | mM | exp | [4] |
| $Ki_{hk}^{G6p_M}$ | Ki_hk_G6p_M | 5.00E-01 | mM | exp | [4] |
| $Km_{hk}^{Atp_M}$ | Km_hk_Atp_M | 2.09E+00 | mM | exp | [4] |
| $Ki_{hk}^{Atp_M}$ | Ki_hk_Atp_M | 1.90E-01 | mM | exp | [4] |
| $Vmax_{g6pase}^{G6p_M}$ | Vmax_ppp_G6p_M | 0.00E+00 | mmol/min | est | |
| $Km_{ppp}^{G6p_M}$ | Km_ppp_G6p_M | 1.00E-01 | mM | est | |
| $Km_{ppp}^{Nadp_M}$ | Km_ppp_Nadp_M | 1.00E-01 | mM | est | |
| $Vmax_{g6pase}^{G6p_M}$ | Vmax_g6pase_G6p_M | 0.00E+00 | mmol/min | opt | |

| | | | | | |
|---|---|---|---|---|---|
| $Km_{g6pase}^{G6p_M}$ | Km_g6pase_G6p_M | 2.41E+00 | mM | exp | [5, 6] |
| $Vmax_{gs}^{G6p_M}$ | Vmax_gs_G6p_M | 5.00E+00 | mmol/min | opt | |
| $Km_{gs}^{G6p_M}$ | Km_gs_G6p_M | 5.00E-02 | mM | est | [7] |
| $n_{gs}^{G6p_M}$ | n_gs_G6p_M | 1.00E+00 | - | est | |
| $Km_{gs}^{Utp_M}$ | Km_gs_Utp_M | 4.80E-02 | mM | exp | [8] |
| $Glygn_M^{max}$ | Glygn_M_max | 4.60E+01 | mM | est | [9] |
| $Km_{gs}^{Glygn_M}$ | Km_gs_Glygn_M | 1.00E+01 | mM | est | |
| $Vmax_{gd}^{Glygn_M}$ | Vmax_gd_Glygn_M | 1.20E+00 | mmol/min | opt | |
| $Km_{gd}^{Glygn_M}$ | Km_gd_Glygn_M | 1.00E+01 | mM | est | [7] |
| $Km_{gd}^{Phos_M}$ | Km_gd_Phos_M | 4.00E+00 | mM | exp | [10] |
| $Vmax_{pfk}^{G6p_M}$ | Vmax_pfk_G6p_M | 1.05E+01 | mmol/min | opt | |
| $Km_{pfk}^{G6p_M}$ | Km_pfk_G6p_M | 5.00E-03 | mM | est | [7] |
| $Km_{pfk}^{Atp_M}$ | Km_pfk_Atp_M | 4.25E-02 | mM | exp | [11] |
| $Ki_{pfk}^{Atp_M}$ | Ki_pfk_Atp_M | 2.10E+00 | mM | exp | [12] |
| $Km_{pfk}^{Adp_M}$ | Km_pfk_Adp_M | 8.36E-02 | mM | exp | [11, 12] |
| $Ki_{pfk}^{Gap_M}$ | Ki_pfk_Gap_M | 2.07E-02 | mM | exp | [13] |
| $b_{pfk}^{Gap_M}$ | b_pfk_Gap_M | 7.50E-01 | - | est | [7] |
| $Vmax_{fbp}^{Gap_M}$ | Vmax_fbp_Gap_M | 0.00E+00 | mmol/min | opt | |
| $Km_{fbp}^{Gap_M}$ | Km_fbp_Gap_M | 2.50E-01 | mM | est | [7] |
| $Vmax_{pk}^{Gap_M}$ | Vmax_pk_Gap_M | 1.50E+02 | mmol/min | | |
| $Km_{pk}^{Gap_M}$ | Km_pk_Gap_M | 2.40E-01 | mM | est | [7] |
| $b_{pk}^{Accoa_{MM}}$ | b_pk_Accoa_MM | 8.00E-01 | - | exp | [14] |
| $Ki_{pk}^{Accoa_{MM}}$ | Ki_pk_Accoa_MM | 3.00E-02 | mM | exp | [14] |
| $Km_{pk}^{Adp_M}$ | Km_pk_Adp_M | 2.40E-01 | mM | exp | [15] |
| $Vmax_{pepck}^{Pyr_M}$ | Vmax_pepck_Pyr_M | 0.00E+00 | mM | opt | |
| $Km_{pepck}^{Pyr_M}$ | Km_pepck_Pyr_M | 1.14E+00 | mM | est | |
| $Km_{pepck}^{Atp_M}$ | Km_pepck_Atp_M | 1.00E-02 | mM | est | [7] |

| | | | | | |
|---|---|---|---|---|---|
| $Km_{pepck}^{Gtp_M}$ | Km_pepck_Gtp_M | 6.40E-02 | mM | exp | [16] |
| $Vdif_{pyrt}^{Pyr_{BM}}$ | Vdif_pyrt_Pyr_B_M | 1.40E-02 | L/min | est | |
| $Kdif_{pyrt}^{Pyr_{BM}}$ | Kdif_pyrt_Pyr_B_M | 1.00E+00 | mM | est | |
| $Kdif_{pyrt}^{Pyr_M}$ | Kdif_pyrt_Pyr_M | 1.00E+00 | mM | est | |
| $Vdif_{lact}^{Lac_{BM}}$ | Vdif_lact_Lac_B_M | 8.07E+00 | L/min | opt | |
| $Kdif_{lact}^{Lac_M}$ | Kdif_lact_Lac_M | 2.42E+00 | mM | exp | [17] |
| $Kdif_{lact}^{Lac_{BM}}$ | Kdif_lact_Lac_B_M | 2.42E+00 | mM | exp | [17] |
| $Vmax_{ldh}^{Pyr_M}$ | Vmax_ldh_Pyr_M | 1.00E+02 | $L^2$/mmol/min | opt | |
| $Km_{ldh}^{Pyr_M}$ | Km_ldh_Pyr_M | 1.50E-01 | mM | exp | [18] |
| $Km_{ldh}^{Nadh_M}$ | Km_ldh_Nadh_M | 1.50E-02 | mM | exp | [18] |
| $Km_{ldh}^{Lac_M}$ | Km_ldh_Lac_M | 3.60E+01 | mM | exp | [19] |
| $Km_{ldh}^{Nad_M}$ | Km_ldh_Nad_M | 1.10E-01 | mM | exp | [18] |
| $Keq_{ldh}^{Lac_M}$ | Keq_ldh_Lac_M | 1.00E+03 | - | est | |
| $Vdif_{alat}^{Ala_{BM}}$ | Vdif_alat_Ala_B_M | 7.20E+00 | L/min | est | |
| $Kdif_{alat}^{Ala_M}$ | Kdif_alat_Ala_M | 2.42E+00 | mM | est | |
| $Kdif_{alat}^{Ala_{BM}}$ | Kdif_alat_Ala_B_M | 2.42E+00 | mM | est | |
| $Vmax_{alata}^{Pyr_M}$ | Vmax_alata_Pyr_M | 4.67E-01 | L/min | opt | |
| $Km_{alata}^{Pyr_M}$ | Km_alata_Pyr_M | 1.50E-01 | mM | est | |
| $Km_{alata}^{Ala_M}$ | Km_alata_Ala_M | 3.60E+01 | mM | est | |
| $Keq_{alata}^{Ala_M}$ | Keq_alata_Ala_M | 1.00E+03 | - | est | |
| $Vmax_{pdh}^{Pyr_M}$ | Vmax_pdh_Pyr_M | 2.00E+01 | mmol/min | opt | |
| $Km_{pdh}^{Pyr_M}$ | Km_pdh_Pyr_M | 5.40E-01 | mM | est | [7] |
| $Km_{pdh}^{Accoa_{MM}}$ | Ki_pdh_Accoa_MM | 3.00E-02 | mM | exp | [20] |
| $Km_{pdh}^{Nad_M}$ | Km_pdh_Nad_M | 1.00E-01 | mM | opt | |
| $Vmax_{tca}^{Accoa_{MM}}$ | Vmax_tca_Accoa_MM | 4.82E+01 | mmol/min | opt | |
| $Km_{tca}^{Accoa_{MM}}$ | Km_tca_Accoa_MM | 4.00E-04 | mM | exp | [21] |
| $Km_{tca}^{Phos_M}$ | Km_tca_Phos_M | 1.00E+00 | mM | opt | |
| $Km_{tca}^{Adp_M}$ | Km_tca_Adp_M | 4.10E-01 | mM | exp | [22] |

| | | | | | |
|---|---|---|---|---|---|
| $Km_{tca}^{Pyr_M}$ | Km_tca_Pyr_M | 3.86E-01 | mM | opt | |
| $Km_{tca}^{Nad_M}$ | Km_tca_Nad_M | 1.00E-02 | mM | est | |
| $Km_{tca}^{Fad_M}$ | Km_tca_Fad_M | 1.00E-01 | mM | est | |
| $Vdif_{ffat}^{FFA_{BM}}$ | Vdif_ffat_FFA_B_M | 3.70E-01 | L/min | opt | |
| $Kdif_{ffat}^{FFA_{BM}}$ | Kdif_ffat_FFA_B_M | 2.00E-01 | mM | est | [7] |
| $Kdif_{ffat}^{FFA_M}$ | Kdif_ffat_FFA_M | 2.00E-01 | mM | est | [7] |
| $Vmax_{ffat}^{FFA_{BM}}$ | Vmax_ffat_FFA_B_M | 0.00E+00 | mmol/min | opt | |
| $Km_{ffat}^{FFA_{BM}}$ | Km_ffat_FFA_B_M | 2.00E-03 | mM | est | [7] |
| $Vmax_{tgsyn}^{FFA_M}$ | Vmax_tgsyn_FFA_M | 3.00E+00 | mmol/min | opt | |
| $Km_{tgsyn}^{FFA_M}$ | Km_tgsyn_FFA_M | 6.45E-01 | mM | est | [7] |
| $Km_{tgsyn}^{Glycp_M}$ | Km_tgsyn_Glycp_M | 4.60E-01 | mM | exp | [23] |
| $Km_{tgsyn}^{Atp_M}$ | Km_tgsyn_Atp_M | 1.00E-01 | mM | opt | |
| $TG_M^{max}$ | TG_M_max | 1.48E+01 | mM | est | |
| $Km_{tgsyn}^{TG_M}$ | Km_tgsyn_TG_M | 1.00E+01 | mM | est | |
| $Vmax_{tgdeg}^{TG_M}$ | Vmax_tgdeg_TG_M | 1.80E+00 | mmol/min | opt | |
| $Km_{tgdeg}^{TG_M}$ | Km_tgdeg_TG_M | 5.07E+01 | mM | est | [7] |
| $Vdif_{glyct}^{Glyc_{BM}}$ | Vdif_glyct_Glyc_B_M | 1.20E-01 | L/min | opt | |
| $Kdif_{glyct}^{Glyc_{BM}}$ | Kdif_glyct_Glyc_B_M | 2.70E-01 | mM | exp | [24] |
| $Kdif_{glyct}^{Glyc_M}$ | Kdif_glyct_Glyc_M | 2.70E-01 | mM | exp | [24] |
| $Vmax_{glyk}^{Glyc_M}$ | Vmax_glyk_Glyc_M | 4.00E+00 | mmol/min | opt | |
| $Km_{glyk}^{Glyc_M}$ | Km_glyk_Glyc_M | 1.00E-03 | mM | exp | [24] |
| $Km_{glyk}^{Atp_M}$ | Km_glyk_Atp_M | 1.00E-01 | mM | exp | [25] |
| $Ki_{glyk}^{Gap_M}$ | Ki_glyk_Gap_M | 1.00E-01 | mM | exp | [25] |
| $Vdif_{tgt}^{TG_{BM}}$ | Vdif_tgt_TG_B_M | 8.00E-03 | mmol/min | est | |
| $Kdif_{tgt}^{TG_{BM}}$ | Kdif_tgt_TG_B_M | 1.00E+00 | mM | est | [7] |
| $Keq_{tgt}^{TG_M}$ | Keq_tgt_TG_M | 3.38E+01 | - | est | [7] |
| $Vmax_{tgt}^{TG_{BM}}$ | Vmax_tgt_TG_B_M | 0.00E+00 | mmol/min | opt | |

| | | | | | |
|---|---|---|---|---|---|
| $Km_{tgt}^{TG_M}$ | Km_tgt_TG_M | 3.38E+01 | mM | est | [7] |
| $Vmax_{g3pd}^{Gap_M}$ | Vmax_g3pd_Gap_M | 1.00E+00 | $L^2$/mmol/min | opt | |
| $Km_{g3pd}^{Gap_M}$ | Km_g3pd_Gap_M | 1.00E+00 | mM | exp | [26] |
| $Km_{g3pd}^{Glycp_M}$ | Km_g3pd_Glycp_M | 1.00E+00 | mM | exp | [26] |
| $Km_{g3pd}^{Nadh_M}$ | Km_g3pd_Nadh_M | 1.00E-01 | mM | exp | [26] |
| $Km_{g3pd}^{Nad_M}$ | Km_g3pd_Nad_M | 1.00E-01 | mM | exp | [26] |
| $Keq_{g3pd}^{Glycp_M}$ | Keq_g3pd_Glycp_M | 1.00E+03 | - | opt | |
| $Vmax_{boxid}^{FFA_M}$ | Vmax_boxid_FFA_M | 8.00E-01 | mmol/min | opt | |
| $Km_{boxid}^{FFA_M}$ | Km_boxid_FFA_M | 5.00E-03 | mM | exp | [27-29] |
| $Km_{boxid}^{Atp_M}$ | Km_boxid_Atp_M | 8.70E-02 | mM | exp | [30] |
| $Ki_{boxid}^{Accoa_{MM}}$ | Ki_boxid_Accoa_MM | 1.20E-01 | mM | | [31] |
| $Ki_{boxid}^{Malcoa_M}$ | Ki_boxid_Malcoa_M | 6.00E-02 | mM | opt | |
| $Km_{boxid}^{Nad_M}$ | Km_boxid_Nad_M | 1.00E-01 | mM | est | |
| $Km_{boxid}^{Fad_M}$ | Km_boxid_Fad_M | 1.00E-02 | mM | est | |
| $Vmax_{accoat}^{Accoa_{MMMC}}$ | Vmax_accoat_Accoa_MM_MC | 0.00E+00 | mmol/min | est | |
| $Km_{accoat}^{Accoa_{MM}}$ | Km_accoat_Accoa_MM | 4.00E-04 | mM | opt | |
| $Km_{accoat}^{Atp_M}$ | Km_accoat_Atp_M | 1.00E+00 | mM | est | |
| $Km_{accoat}^{Pyr_M}$ | Km_accoat_Pyr_M | 1.00E-02 | mM | est | |
| $Vmax_{bhbsyn}^{Accoa_{MM}}$ | Vmax_bhbsyn_Accoa_MM | 0.00E+00 | mmol/min | opt | |
| $Km_{bhbsyn}^{Accoa_{MM}}$ | Km_bhbsyn_Accoa_MM | 1.00E-01 | mM | est | |
| $Km_{bhbsyn}^{Nadh_M}$ | Km_bhbsyn_Nadh_M | 1.00E-01 | mM | est | |
| $n_{bhbsyn}^{Pyr_M}$ | n_bhbsyn_Pyr_M | 4.00E+00 | - | est | |
| $Ki_{bhbsyn}^{Pyr_M}$ | Ki_bhbsyn_Pyr_M | 1.00E-02 | mM | opt | |
| $Vmax_{bhbdeg}^{Bhb_M}$ | Vmax_bhbdeg_Bhb_M | 0.00E+00 | mmol/min | est | |
| $Km_{bhbdeg}^{Bhb_M}$ | Km_bhbdeg_Bhb_M | 1.00E-01 | mM | est | |
| $Km_{bhbdeg}^{Atp_M}$ | Km_bhbdeg_Atp_M | 1.00E-01 | mM | est | |
| $Km_{bhbdeg}^{Nad_M}$ | Km_bhbdeg_Nad_M | 1.00E-01 | mM | est | |
| $Vdif_{bhbt}^{Bhb_{BM}}$ | Vdif_bhbt_Bhb_B_M | 0.00E+00 | L/min | est | |

| | | | | | |
|---|---|---|---|---|---|
| $Kdif_{bhbt}^{Bhb_{BM}}$ | Kdif_bhbt_Bhb_B_M | 1.00E+00 | mM | est | |
| $Kdif_{bhbt}^{Bhb_{M}}$ | Kdif_bhbt_Bhb_M | 1.00E+00 | mM | est | |
| $Vmax_{lipog1}^{Accoa_{MC}}$ | Vmax_lipog1_Accoa_MC | 0.00E+00 | mmol/min | opt | |
| $Km_{lipog1}^{Accoa_{MC}}$ | Km_lipog1_Accoa_MC | 1.00E-02 | mM | est | |
| $Km_{lipog1}^{Atp_{M}}$ | Km_lipog1_Atp_M | 1.00E-01 | mM | est | |
| $Vmax_{lipog2}^{Malcoa_{M}}$ | Vmax_lipog2_Malcoa_M | 0.00E+00 | mmol/min | est | |
| $Km_{lipog2}^{Malcoa_{M}}$ | Km_lipog2_Malcoa_M | 1.00E-02 | mM | est | |
| $Km_{lipog2}^{Adp_{M}}$ | Km_lipog2_Adp_M | 1.00E-01 | mM | est | |
| $Km_{lipog2}^{Nadph_{M}}$ | Km_lipog2_Nadph_M | 1.00E-02 | mM | est | |
| $Vmax_{cholsyn1}^{Accoa_{MC}}$ | Vmax_cholsyn1_Accoa_MC | 0.00E+00 | mmol/min | est | |
| $Km_{cholsyn1}^{Accoa_{MC}}$ | Km_cholsyn1_Accoa_MC | 1.00E-01 | mM | est | |
| $Vmax_{cholsyn2}^{Hmgcoa_{M}}$ | Vmax_cholsyn2_Hmgcoa_M | 0.00E+00 | mmol/min | est | |
| $Km_{cholsyn2}^{Hmgcoa_{M}}$ | Km_cholsyn2_Hmgcoa_M | 1.00E-01 | mM | est | |
| $Km_{cholsyn2}^{Atp_{M}}$ | Km_cholsyn2_Atp_M | 1.00E-01 | mM | est | |
| $Km_{cholsyn2}^{Fadh_{M}}$ | Km_cholsyn2_Fadh_M | 1.00E-01 | mM | est | |
| $Km_{cholsyn2}^{Nadph_{M}}$ | Km_cholsyn2_Nadph_M | 1.00E-01 | mM | est | |
| $Vmax_{cholt}^{Chol_{BM}}$ | Vmax_cholt_Chol_M | 0.00E+00 | mmol/min | est | |
| $Km_{cholt}^{Chol_{M}}$ | Km_cholt_Chol_M | 1.00E-01 | mM | est | |
| $Vmax_{atpsynf}^{Fadh_{M}}$ | Vmax_atpsynf_Fadh_M | 5.99E+00 | mmol/min | opt | |
| $Km_{atpsynf}^{Fadh_{M}}$ | Km_atpsynf_Fadh_M | 1.00E-01 | mM | est | |
| $Km_{atpsynf}^{Adp_{M}}$ | Km_atpsynf_Adp_M | 1.00E-01 | mM | est | |
| $Vmax_{atpsynn}^{Nadh_{M}}$ | Vmax_atpsynn_Nadh_M | 1.00E+01 | mmol/min | est | |
| $Km_{atpsynn}^{Nadh_{M}}$ | Km_atpsynn_Nadh_M | 1.00E-01 | mM | est | |
| $Km_{atpsynn}^{Adp_{M}}$ | Km_atpsynn_Adp_M | 1.00E-01 | mM | est | |
| $Vmax_{atpuse}^{Atp_{M}}$ | Vmax_atpuse_Atp_M | 9.09E+00 | mmol/min | opt | |
| $Km_{atpuse}^{Atp_{M}}$ | Km_atpuse_Atp_M | 2.50E+00 | mM | opt | |
| $Vmax_{ampreg}^{Amp_{M}}$ | Vmax_ampreg_Amp_M | 1.00E+01 | mmol/min | est | |
| $Km_{ampreg}^{Amp_{M}}$ | Km_ampreg_Amp_M | 8.00E-02 | mM | exp | [32] |

| | | | | | |
|---|---|---|---|---|---|
| $Km_{ampreg}^{Atp_M}$ | Km_ampreg_Atp_M | 9.00E-02 | mM | exp | [32] |
| $Km_{ampreg}^{Adp_M}$ | Km_ampreg_Adp_M | 1.10E-01 | mM | exp | [32] |
| $Vmax_{nadhk}^{Nadh_M}$ | Vmax_nadhk_Nadh_M | 0.00E+00 | mmol/min | est | |
| $Km_{nadhk}^{Nadh_M}$ | Km_nadhk_Nadh_M | 1.00E-01 | mM | est | |
| $Km_{nadhk}^{Atp_M}$ | Km_nadhk_Atp_M | 1.00E-01 | mM | est | |
| $Vmax_{nadhuse}^{Nadh_M}$ | Vmax_nadhuse_Nadh_M | 1.00E+00 | mmol/min | est | |
| $Km_{nadhuse}^{Nadh_M}$ | Km_nadhuse_Nadh_M | 1.00E-01 | mM | est | |
| $Vmax_{gdpreg}^{Gdp_M}$ | Vmax_gdpreg_Gdp_M | 1.73E+03 | $L^2$/mmol/min | est | |
| $Km_{gdpreg}^{Gdp_M}$ | Km_gdpreg_Gdp_M | 3.10E-02 | mM | exp | [33] |
| $Km_{gdpreg}^{Atp_M}$ | Km_gdpreg_Atp_M | 1.33E+00 | mM | exp | [33] |
| $Km_{gdpreg}^{Gtp_M}$ | Km_gdpreg_Gtp_M | 1.50E-01 | mM | exp | [34] |
| $Km_{gdpreg}^{Adp_M}$ | Km_gdpreg_Adp_M | 4.20E-02 | mM | exp | [33] |
| $Keq_{gdpreg}^{Gtp_M}$ | Keq_gdpreg_Gtp_M | 1.00E+03 | - | opt | |
| $Vmax_{gtpuse}^{Gtp_M}$ | Vmax_gtpuse_Gtp_M | 0.00E+00 | mmol/min | est | |
| $Km_{gtpuse}^{Gtp_M}$ | Km_gtpuse_Gtp_M | 1.00E-01 | mM | est | |
| $Vmax_{udpreg}^{Udp_M}$ | Vmax_udpreg_Udp_M | 9.11E+01 | $L^2$/mmol/min | est | |
| $Km_{udpreg}^{Udp_M}$ | Km_udpreg_Udp_M | 1.90E-01 | mM | exp | [33] |
| $Km_{udpreg}^{Atp_M}$ | Km_udpreg_Atp_M | 1.33E+00 | mM | exp | [33] |
| $Km_{udpreg}^{Utp_M}$ | Km_udpreg_Utp_M | 1.60E+01 | mM | exp | [34] |
| $Km_{udpreg}^{Adp_M}$ | Km_udpreg_Adp_M | 4.20E-02 | mM | exp | [33] |
| $Keq_{udpreg}^{Utp_M}$ | Keq_udpreg_Utp_M | 1.00E+03 | - | est | |
| $Vmax_{utpuse}^{Utp_M}$ | Vmax_utpuse_Utp_M | 0.00E+00 | mmol/min | est | |
| $Km_{utpuse}^{Utp_M}$ | Km_utpuse_Utp_M | 1.00E-01 | mM | est | |
| $Vmax_{nadphuse}^{Nadph_M}$ | Vmax_nadphuse_Nadph_M | 5.00E-01 | mmol/min | est | |
| $Km_{nadphuse}^{Nadph_M}$ | Km_nadphuse_Nadph_M | 1.00E-01 | mM | est | |
| $Vmax_{fadhuse}^{Fadh_M}$ | Vmax_fadhuse_Fadh_M | 0.00E+00 | mmol/min | est | |
| $Km_{fadhuse}^{Fadh_M}$ | Km_fadhuse_Fadh_M | 1.00E-01 | mM | est | |
| $Vmax_{ck}^{Cre_M}$ | Vmax_ck_Cre_M | 0.00E+00 | $L^2$/mmol/min | est | |

| | | | | | |
|---|---|---|---|---|---|
| $Km_{ck}^{Cre_M}$ | Km_ck_Cre_M | 1.00E+00 | mM | est | |
| $Km_{ck}^{Atp_M}$ | Km_ck_Atp_M | 3.00E-02 | mM | est | |
| $Km_{ck}^{Crep_M}$ | Km_ck_Crep_M | 1.50E+01 | mM | est | |
| $Km_{ck}^{Adp_M}$ | Km_ck_Adp_M | 1.00E-01 | mM | est | |
| $Keq_{ck}^{Crep_M}$ | Keq_ck_Crep_M | 1.00E+00 | - | est | |
| $Phos_M$ | Phos_M | 5.00E+00 | mM | est | |
| $\alpha_A^{Base}$ | alpha_base_A | 5.00E-02 | - | opt | |
| $\alpha_A^{Band}$ | alpha_band_A | 6.00E-01 | - | opt | |
| $Km^{Ins_{BA}}$ | Km_Ins_B_A | 8.00E-08 | mM | est | |
| $n^{Ins_{BA}}$ | n_Ins_B_A | 4.00E+00 | - | est | |
| $Vdif_{glut4}^{Glc_{BA}}$ | Vdif_glut4_Glc_B_A | 1.55E-01 | L/min | opt | |
| $Kdif_{glut4}^{Glc_{BA}}$ | Kdif_glut4_Glc_B_A | 5.00E+00 | mM | exp | [3] |
| $Kdif_{glut4}^{Glc_A}$ | Kdif_glut4_Glc_A | 1.00E+00 | mM | exp | [3] |
| $Vmax_{hk}^{Glc_A}$ | Vmax_hk_Glc_A | 7.83E-01 | mmol/min | opt | |
| $Km_{hk}^{Glc_A}$ | Km_hk_Glc_A | 4.20E-01 | mM | exp | [4] |
| $Ki_{hk}^{G6p_A}$ | Ki_hk_G6p_A | 5.00E-01 | mM | exp | [4] |
| $Km_{hk}^{Atp_A}$ | Km_hk_Atp_A | 2.09E+00 | mM | exp | [4] |
| $Ki_{hk}^{Atp_A}$ | Ki_hk_Atp_A | 1.90E-01 | mM | exp | [4] |
| $Vmax_{g6pase}^{G6p_A}$ | Vmax_ppp_G6p_A | 0.00E+00 | mmol/min | est | |
| $Km_{ppp}^{G6p_A}$ | Km_ppp_G6p_A | 8.65E-01 | mM | est | |
| $Km_{ppp}^{Nadp_A}$ | Km_ppp_Nadp_A | 1.55E-01 | mM | est | |
| $Vmax_{g6pase}^{G6p_A}$ | Vmax_g6pase_G6p_A | 0.00E+00 | mmol/min | opt | |
| $Km_{g6pase}^{G6p_A}$ | Km_g6pase_G6p_A | 1.00E+00 | mM | exp | [5, 6] |
| $Vmax_{gs}^{G6p_A}$ | Vmax_gs_G6p_A | 0.00E+00 | mmol/min | opt | |
| $Km_{gs}^{G6p_A}$ | Km_gs_G6p_A | 5.00E-02 | mM | est | [7] |
| $n_{gs}^{G6p_A}$ | n_gs_G6p_A | 1.00E+00 | - | est | |
| $Km_{gs}^{Utp_A}$ | Km_gs_Utp_A | 4.80E-02 | mM | exp | [8] |
| $Glygn_A^{max}$ | Glygn_A_max | 0.00E+00 | mM | est | [9] |

| | | | | | |
|---|---|---|---|---|---|
| $Km_{gs}^{Glygn_A}$ | Km_gs_Glygn_A | 9.20E+00 | mM | est | |
| $Vmax_{gd}^{Glygn_A}$ | Vmax_gd_Glygn_A | 0.00E+00 | mmol/min | opt | |
| $Km_{gd}^{Glygn_A}$ | Km_gd_Glygn_A | 1.00E+01 | mM | est | [7] |
| $Km_{gd}^{Phos_A}$ | Km_gd_Phos_A | 4.00E+00 | mM | exp | [10] |
| $Vmax_{pfk}^{G6p_A}$ | Vmax_pfk_G6p_A | 1.26E+02 | mmol/min | opt | |
| $Km_{pfk}^{G6p_A}$ | Km_pfk_G6p_A | 5.00E-03 | mM | est | [7] |
| $Km_{pfk}^{Atp_A}$ | Km_pfk_Atp_A | 4.25E-02 | mM | exp | [11] |
| $Ki_{pfk}^{Atp_A}$ | Ki_pfk_Atp_A | 2.10E+00 | mM | exp | [12] |
| $Km_{pfk}^{Adp_A}$ | Km_pfk_Adp_A | 8.36E-02 | mM | exp | [11, 12] |
| $Ki_{pfk}^{Gap_A}$ | Ki_pfk_Gap_A | 2.07E-02 | mM | exp | [13] |
| $b_{pfk}^{Gap_A}$ | b_pfk_Gap_A | 7.50E-01 | - | est | [7] |
| $Vmax_{fbp}^{Gap_A}$ | Vmax_fbp_Gap_A | 0.00E+00 | mmol/min | opt | |
| $Km_{fbp}^{Gap_A}$ | Km_fbp_Gap_A | 2.50E-01 | mM | est | [7] |
| $Vmax_{pk}^{Gap_A}$ | Vmax_pk_Gap_A | 1.30E+00 | mmol/min | | |
| $Km_{pk}^{Gap_A}$ | Km_pk_Gap_A | 2.40E-01 | mM | est | [7] |
| $b_{pk}^{Accoa_{AM}}$ | b_pk_Accoa_AM | 8.00E-01 | - | exp | [14] |
| $Ki_{pk}^{Accoa_{AM}}$ | Ki_pk_Accoa_AM | 3.00E-02 | mM | exp | [14] |
| $Km_{pk}^{Adp_A}$ | Km_pk_Adp_A | 2.40E-01 | mM | exp | [15] |
| $Vmax_{pepck}^{Pyr_A}$ | Vmax_pepck_Pyr_A | 0.00E+00 | mM | opt | |
| $Km_{pepck}^{Pyr_A}$ | Km_pepck_Pyr_A | 1.00E+00 | mM | est | |
| $Km_{pepck}^{Atp_A}$ | Km_pepck_Atp_A | 1.00E-02 | mM | est | [7] |
| $Km_{pepck}^{Gtp_A}$ | Km_pepck_Gtp_A | 6.40E-02 | mM | exp | [16] |
| $Vdif_{pyrt}^{Pyr_{BA}}$ | Vdif_pyrt_Pyr_B_A | 0.00E+00 | L/min | est | |
| $Kdif_{pyrt}^{Pyr_{BA}}$ | Kdif_pyrt_Pyr_B_A | 1.00E+00 | mM | est | |
| $Kdif_{pyrt}^{Pyr_A}$ | Kdif_pyrt_Pyr_A | 1.00E+00 | mM | est | |
| $Vdif_{lact}^{Lac_{BA}}$ | Vdif_lact_Lac_B_A | 1.20E+01 | L/min | opt | |
| $Kdif_{lact}^{Lac_A}$ | Kdif_lact_Lac_A | 2.42E+00 | mM | exp | [17] |

| | | | | | |
|---|---|---|---|---|---|
| $Kdif_{lact}^{Lac_{BA}}$ | Kdif_lact_Lac_B_A | 2.42E+00 | mM | exp | [17] |
| $Vmax_{ldh}^{Pyr_A}$ | Vmax_ldh_Pyr_A | 1.00E+02 | $L^2$/mmol/min | opt | |
| $Km_{ldh}^{Pyr_A}$ | Km_ldh_Pyr_A | 1.50E-01 | mM | exp | [18] |
| $Km_{ldh}^{Nadh_A}$ | Km_ldh_Nadh_A | 1.50E-02 | mM | exp | [18] |
| $Km_{ldh}^{Lac_A}$ | Km_ldh_Lac_A | 3.60E+01 | mM | exp | [19] |
| $Km_{ldh}^{Nad_A}$ | Km_ldh_Nad_A | 1.10E-01 | mM | exp | [18] |
| $Keq_{ldh}^{Lac_A}$ | Keq_ldh_Lac_A | 1.00E+03 | - | est | |
| $Vdif_{alat}^{Ala_{BA}}$ | Vdif_alat_Ala_B_A | 0.00E+00 | L/min | est | |
| $Kdif_{alat}^{Ala_A}$ | Kdif_alat_Ala_A | 2.42E+00 | mM | est | |
| $Kdif_{alat}^{Ala_{BA}}$ | Kdif_alat_Ala_B_A | 2.42E+00 | mM | est | |
| $Vmax_{alata}^{Pyr_A}$ | Vmax_alata_Pyr_A | 0.00E+00 | L/min | opt | |
| $Km_{alata}^{Pyr_A}$ | Km_alata_Pyr_A | 1.50E-01 | mM | est | |
| $Km_{alata}^{Ala_A}$ | Km_alata_Ala_A | 3.60E+01 | mM | est | |
| $Keq_{alata}^{Ala_A}$ | Keq_alata_Ala_A | 1.00E+03 | - | est | |
| $Vmax_{pdh}^{Pyr_A}$ | Vmax_pdh_Pyr_A | 5.00E+01 | mmol/min | opt | |
| $Km_{pdh}^{Pyr_A}$ | Km_pdh_Pyr_A | 5.40E-01 | mM | est | [7] |
| $Km_{pdh}^{Accoa_{AM}}$ | Ki_pdh_Accoa_AM | 3.50E-02 | mM | exp | [20] |
| $Km_{pdh}^{Nad_A}$ | Km_pdh_Nad_A | 1.00E-02 | mM | opt | |
| $Vmax_{tca}^{Accoa_{AM}}$ | Vmax_tca_Accoa_AM | 7.05E+00 | mmol/min | opt | |
| $Km_{tca}^{Accoa_{AM}}$ | Km_tca_Accoa_AM | 4.00E-04 | mM | exp | [21] |
| $Km_{tca}^{Phos_A}$ | Km_tca_Phos_A | 1.00E+00 | mM | opt | |
| $Km_{tca}^{Adp_A}$ | Km_tca_Adp_A | 4.10E-01 | mM | exp | [22] |
| $Km_{tca}^{Pyr_A}$ | Km_tca_Pyr_A | 1.12E-01 | mM | opt | |
| $Km_{tca}^{Nad_A}$ | Km_tca_Nad_A | 1.00E-01 | mM | est | |
| $Km_{tca}^{Fad_A}$ | Km_tca_Fad_A | 1.00E-01 | mM | est | |
| $Vdif_{ffat}^{FFA_{BA}}$ | Vdif_ffat_FFA_B_A | 0.00E+00 | L/min | opt | |
| $Kdif_{ffat}^{FFA_{BA}}$ | Kdif_ffat_FFA_B_A | 2.00E-01 | mM | est | [7] |
| $Kdif_{ffat}^{FFA_A}$ | Kdif_ffat_FFA_A | 2.00E-01 | mM | est | [7] |

| | | | | | |
|---|---|---|---|---|---|
| $Vmax_{ffat}^{FFA_{BA}}$ | Vmax_ffat_FFA_B_A | 2.20E-01 | mmol/min | opt | |
| $Km_{ffat}^{FFA_{BA}}$ | Km_ffat_FFA_B_A | 4.00E-01 | mM | est | [7] |
| $Vmax_{tgsyn}^{FFA_A}$ | Vmax_tgsyn_FFA_A | 5.50E-01 | mmol/min | opt | |
| $Km_{tgsyn}^{FFA_A}$ | Km_tgsyn_FFA_A | 2.00E-01 | mM | est | [7] |
| $Km_{tgsyn}^{Glycp_A}$ | Km_tgsyn_Glycp_A | 4.60E-01 | mM | exp | [23] |
| $Km_{tgsyn}^{Atp_A}$ | Km_tgsyn_Atp_A | 1.00E-01 | mM | opt | |
| $TG_A^{max}$ | TG_A_max | 9.00E+02 | mM | est | |
| $Km_{tgsyn}^{TG_A}$ | Km_tgsyn_TG_A | 1.00E+01 | mM | est | |
| $Vmax_{tgdeg}^{TG_A}$ | Vmax_tgdeg_TG_A | 9.20E-02 | mmol/min | opt | |
| $Km_{tgdeg}^{TG_A}$ | Km_tgdeg_TG_A | 5.07E+01 | mM | est | [7] |
| $Vdif_{glyct}^{Glyc_{BA}}$ | Vdif_glyct_Glyc_B_A | 1.00E+01 | L/min | opt | |
| $Kdif_{glyct}^{Glyc_{BA}}$ | Kdif_glyct_Glyc_B_A | 2.70E-01 | mM | exp | [24] |
| $Kdif_{glyct}^{Glyc_A}$ | Kdif_glyct_Glyc_A | 2.70E-01 | mM | exp | [24] |
| $Vmax_{glyk}^{Glyc_A}$ | Vmax_glyk_Glyc_A | 0.00E+00 | mmol/min | opt | |
| $Km_{glyk}^{Glyc_A}$ | Km_glyk_Glyc_A | 1.00E-03 | mM | exp | [24] |
| $Km_{glyk}^{Atp_A}$ | Km_glyk_Atp_A | 1.00E+00 | mM | exp | [25] |
| $Ki_{glyk}^{Gap_A}$ | Ki_glyk_Gap_A | 1.00E+00 | mM | exp | [25] |
| $Vdif_{tgt}^{TG_{BA}}$ | Vdif_tgt_TG_B_A | 6.00E-01 | mmol/min | est | |
| $Kdif_{tgt}^{TG_{BA}}$ | Kdif_tgt_TG_B_A | 1.00E+00 | mM | est | [7] |
| $Keq_{tgt}^{TG_A}$ | Keq_tgt_TG_A | 6.00E+02 | - | est | [7] |
| $Vmax_{tgt}^{TG_{BA}}$ | Vmax_tgt_TG_B_A | 0.00E+00 | mmol/min | opt | |
| $Km_{tgt}^{TG_A}$ | Km_tgt_TG_A | 3.38E+01 | mM | est | [7] |
| $Vmax_{g3pd}^{Gap_A}$ | Vmax_g3pd_Gap_A | 3.00E+00 | $L^2$/mmol/min | opt | |
| $Km_{g3pd}^{Gap_A}$ | Km_g3pd_Gap_A | 1.00E+00 | mM | exp | [26] |
| $Km_{g3pd}^{Glycp_A}$ | Km_g3pd_Glycp_A | 1.00E+00 | mM | exp | [26] |
| $Km_{g3pd}^{Nadh_A}$ | Km_g3pd_Nadh_A | 1.00E+00 | mM | exp | [26] |
| $Km_{g3pd}^{Nad_A}$ | Km_g3pd_Nad_A | 1.00E+00 | mM | exp | [26] |

| | | | | | |
|---|---|---|---|---|---|
| $Keq_{g3pd}^{Glycp_A}$ | Keq_g3pd_Glycp_A | 1.00E+03 | - | opt | |
| $Vmax_{boxid}^{FFA_A}$ | Vmax_boxid_FFA_A | 1.50E-01 | mmol/min | opt | |
| $Km_{boxid}^{FFA_A}$ | Km_boxid_FFA_A | 5.00E-03 | mM | exp | [27-29] |
| $Km_{boxid}^{Atp_A}$ | Km_boxid_Atp_A | 8.70E-02 | mM | exp | [30] |
| $Ki_{boxid}^{Accoa_{AM}}$ | Ki_boxid_Accoa_AM | 1.20E-01 | mM | | [31] |
| $Ki_{boxid}^{Malcoa_A}$ | Ki_boxid_Malcoa_A | 1.00E+00 | mM | opt | |
| $Km_{boxid}^{Nad_A}$ | Km_boxid_Nad_A | 1.00E-01 | mM | est | |
| $Km_{boxid}^{Fad_A}$ | Km_boxid_Fad_A | 1.00E-01 | mM | est | |
| $Vmax_{accoat}^{Accoa_{AMAC}}$ | Vmax_accoat_Accoa_AM_AC | 0.00E+00 | mmol/min | est | |
| $Km_{accoat}^{Accoa_{AM}}$ | Km_accoat_Accoa_AM | 1.00E-01 | mM | opt | |
| $Km_{accoat}^{Atp_A}$ | Km_accoat_Atp_A | 1.00E-02 | mM | est | |
| $Km_{accoat}^{Pyr_A}$ | Km_accoat_Pyr_A | 1.00E+00 | mM | est | |
| $Vmax_{bhbsyn}^{Accoa_{AM}}$ | Vmax_bhbsyn_Accoa_AM | 0.00E+00 | mmol/min | opt | |
| $Km_{bhbsyn}^{Accoa_{AM}}$ | Km_bhbsyn_Accoa_AM | 1.00E-01 | mM | est | |
| $Km_{bhbsyn}^{Nadh_A}$ | Km_bhbsyn_Nadh_A | 1.00E-01 | mM | est | |
| $n_{bhbsyn}^{Pyr_A}$ | n_bhbsyn_Pyr_A | 4.00E+00 | - | est | |
| $Ki_{bhbsyn}^{Pyr_A}$ | Ki_bhbsyn_Pyr_A | 1.00E-02 | mM | opt | |
| $Vmax_{bhbdeg}^{Bhb_A}$ | Vmax_bhbdeg_Bhb_A | 0.00E+00 | mmol/min | est | |
| $Km_{bhbdeg}^{Bhb_A}$ | Km_bhbdeg_Bhb_A | 1.00E-01 | mM | est | |
| $Km_{bhbdeg}^{Atp_A}$ | Km_bhbdeg_Atp_A | 1.00E-01 | mM | est | |
| $Km_{bhbdeg}^{Nad_A}$ | Km_bhbdeg_Nad_A | 1.00E-01 | mM | est | |
| $Vdif_{bhbt}^{Bhb_{BA}}$ | Vdif_bhbt_Bhb_B_A | 1.00E+01 | L/min | est | |
| $Kdif_{bhbt}^{Bhb_{BA}}$ | Kdif_bhbt_Bhb_B_A | 1.00E+00 | mM | est | |
| $Kdif_{bhbt}^{Bhb_A}$ | Kdif_bhbt_Bhb_A | 1.00E+00 | mM | est | |
| $Vmax_{lipog1}^{Accoa_{AC}}$ | Vmax_lipog1_Accoa_AC | 0.00E+00 | mmol/min | opt | |
| $Km_{lipog1}^{Accoa_{AC}}$ | Km_lipog1_Accoa_AC | 1.00E-02 | mM | est | |
| $Km_{lipog1}^{Atp_A}$ | Km_lipog1_Atp_A | 1.00E-01 | mM | est | |
| $Vmax_{lipog2}^{Malcoa_A}$ | Vmax_lipog2_Malcoa_A | 0.00E+00 | mmol/min | est | |

| | | | | | |
|---|---|---|---|---|---|
| $Km_{lipog2}^{Malcoa_A}$ | Km_lipog2_Malcoa_A | 1.00E-01 | mM | est | |
| $Km_{lipog2}^{Adp_A}$ | Km_lipog2_Adp_A | 1.00E-01 | mM | est | |
| $Km_{lipog2}^{Nadph_A}$ | Km_lipog2_Nadph_A | 1.00E-02 | mM | est | |
| $Vmax_{cholsyn1}^{Accoa_{AC}}$ | Vmax_cholsyn1_Accoa_AC | 0.00E+00 | mmol/min | est | |
| $Km_{cholsyn1}^{Accoa_{AC}}$ | Km_cholsyn1_Accoa_AC | 1.00E-01 | mM | est | |
| $Vmax_{cholsyn2}^{Hmgcoa_A}$ | Vmax_cholsyn2_Hmgcoa_A | 0.00E+00 | mmol/min | est | |
| $Km_{cholsyn2}^{Hmgcoa_A}$ | Km_cholsyn2_Hmgcoa_A | 1.00E-01 | mM | est | |
| $Km_{cholsyn2}^{Atp_A}$ | Km_cholsyn2_Atp_A | 1.00E-01 | mM | est | |
| $Km_{cholsyn2}^{Fadh_A}$ | Km_cholsyn2_Fadh_A | 1.00E-01 | mM | est | |
| $Km_{cholsyn2}^{Nadph_A}$ | Km_cholsyn2_Nadph_A | 1.00E-01 | mM | est | |
| $Vmax_{cholt}^{Chol_{BA}}$ | Vmax_cholt_Chol_A | 0.00E+00 | mmol/min | est | |
| $Km_{cholt}^{Chol_A}$ | Km_cholt_Chol_A | 1.00E-01 | mM | est | |
| $Vmax_{atpsynf}^{Fadh_A}$ | Vmax_atpsynf_Fadh_A | 1.00E+00 | mmol/min | opt | |
| $Km_{atpsynf}^{Fadh_A}$ | Km_atpsynf_Fadh_A | 1.15E+00 | mM | est | |
| $Km_{atpsynf}^{Adp_A}$ | Km_atpsynf_Adp_A | 4.31E-01 | mM | est | |
| $Vmax_{atpsynn}^{Nadh_A}$ | Vmax_atpsynn_Nadh_A | 5.00E-01 | mmol/min | est | |
| $Km_{atpsynn}^{Nadh_A}$ | Km_atpsynn_Nadh_A | 1.00E-01 | mM | est | |
| $Km_{atpsynn}^{Adp_A}$ | Km_atpsynn_Adp_A | 1.00E-01 | mM | est | |
| $Vmax_{atpuse}^{Atp_A}$ | Vmax_atpuse_Atp_A | 2.75E+00 | mmol/min | opt | |
| $Km_{atpuse}^{Atp_A}$ | Km_atpuse_Atp_A | 2.50E+00 | mM | opt | |
| $Vmax_{ampreg}^{Amp_A}$ | Vmax_ampreg_Amp_A | 1.00E+01 | mmol/min | est | |
| $Km_{ampreg}^{Amp_A}$ | Km_ampreg_Amp_A | 8.00E-02 | mM | exp | [32] |
| $Km_{ampreg}^{Atp_A}$ | Km_ampreg_Atp_A | 9.00E-02 | mM | exp | [32] |
| $Km_{ampreg}^{Adp_A}$ | Km_ampreg_Adp_A | 1.10E-01 | mM | exp | [32] |
| $Vmax_{nadhk}^{Nadh_A}$ | Vmax_nadhk_Nadh_A | 0.00E+00 | mmol/min | est | |
| $Km_{nadhk}^{Nadh_A}$ | Km_nadhk_Nadh_A | 1.00E-01 | mM | est | |
| $Km_{nadhk}^{Atp_A}$ | Km_nadhk_Atp_A | 1.00E-01 | mM | est | |
| $Vmax_{nadhuse}^{Nadh_A}$ | Vmax_nadhuse_Nadh_A | 9.26E-02 | mmol/min | est | |

| | | | | | |
|---|---|---|---|---|---|
| $Km_{nadhuse}^{Nadh_A}$ | Km_nadhuse_Nadh_A | 1.00E-01 | mM | est | |
| $Vmax_{gdpreg}^{Gdp_A}$ | Vmax_gdpreg_Gdp_A | 0.00E+00 | $L^2$/mmol/min | est | |
| $Km_{gdpreg}^{Gdp_A}$ | Km_gdpreg_Gdp_A | 3.10E-02 | mM | exp | [33] |
| $Km_{gdpreg}^{Atp_A}$ | Km_gdpreg_Atp_A | 1.33E+00 | mM | exp | [33] |
| $Km_{gdpreg}^{Gtp_A}$ | Km_gdpreg_Gtp_A | 1.50E-01 | mM | exp | [34] |
| $Km_{gdpreg}^{Adp_A}$ | Km_gdpreg_Adp_A | 4.20E-02 | mM | exp | [33] |
| $Keq_{gdpreg}^{Gtp_A}$ | Keq_gdpreg_Gtp_A | 1.00E+03 | - | opt | |
| $Vmax_{gtpuse}^{Gtp_A}$ | Vmax_gtpuse_Gtp_A | 0.00E+00 | mmol/min | est | |
| $Km_{gtpuse}^{Gtp_A}$ | Km_gtpuse_Gtp_A | 1.00E-01 | mM | est | |
| $Vmax_{udpreg}^{Udp_A}$ | Vmax_udpreg_Udp_A | 0.00E+00 | $L^2$/mmol/min | est | |
| $Km_{udpreg}^{Udp_A}$ | Km_udpreg_Udp_A | 1.90E-01 | mM | exp | [33] |
| $Km_{udpreg}^{Atp_A}$ | Km_udpreg_Atp_A | 1.33E+00 | mM | exp | [33] |
| $Km_{udpreg}^{Utp_A}$ | Km_udpreg_Utp_A | 1.60E+01 | mM | exp | [34] |
| $Km_{udpreg}^{Adp_A}$ | Km_udpreg_Adp_A | 4.20E-02 | mM | exp | [33] |
| $Keq_{udpreg}^{Utp_A}$ | Keq_udpreg_Utp_A | 1.00E+03 | - | est | |
| $Vmax_{utpuse}^{Utp_A}$ | Vmax_utpuse_Utp_A | 0.00E+00 | mmol/min | est | |
| $Km_{utpuse}^{Utp_A}$ | Km_utpuse_Utp_A | 1.00E-01 | mM | est | |
| $Vmax_{nadphuse}^{Nadph_A}$ | Vmax_nadphuse_Nadph_A | 0.00E+00 | mmol/min | est | |
| $Km_{nadphuse}^{Nadph_A}$ | Km_nadphuse_Nadph_A | 1.00E-01 | mM | est | |
| $Vmax_{fadhuse}^{Fadh_A}$ | Vmax_fadhuse_Fadh_A | 0.00E+00 | mmol/min | est | |
| $Km_{fadhuse}^{Fadh_A}$ | Km_fadhuse_Fadh_A | 1.00E-01 | mM | est | |
| $Vmax_{ck}^{Cre_A}$ | Vmax_ck_Cre_A | 0.00E+00 | $L^2$/mmol/min | est | |
| $Km_{ck}^{Cre_A}$ | Km_ck_Cre_A | 1.00E+00 | mM | est | |
| $Km_{ck}^{Atp_A}$ | Km_ck_Atp_A | 3.00E-02 | mM | est | |
| $Km_{ck}^{Crep_A}$ | Km_ck_Crep_A | 1.50E+01 | mM | est | |
| $Km_{ck}^{Adp_A}$ | Km_ck_Adp_A | 1.00E-01 | mM | est | |
| $Keq_{ck}^{Crep_A}$ | Keq_ck_Crep_A | 1.00E+00 | - | est | |
| $Phos_A$ | Phos_A | 5.00E+00 | mM | est | |

| | | | | | |
|---|---|---|---|---|---|
| $\alpha_G^{Base}$ | alpha_base_G | 1.00E-01 | - | opt | |
| $\alpha_G^{Band}$ | alpha_band_G | 6.00E-01 | - | opt | |
| $Km^{Ins_{BG}}$ | Km_Ins_B_G | 8.00E-08 | mM | est | |
| $n^{Ins_{BG}}$ | n_Ins_B_G | 4.00E+00 | - | est | |
| $Vdif_{glut4}^{Glc_{BG}}$ | Vdif_glut2_Glc_B_G | 2.00E-02 | L/min | opt | |
| $Kdif_{glut4}^{Glc_{BG}}$ | Kdif_glut2_Glc_B_G | 1.70E+01 | mM | exp | [3] |
| $Kdif_{glut4}^{Glc_G}$ | Kdif_glut2_Glc_G | 1.70E+01 | mM | exp | [3] |
| $Vmax_{hk}^{Glc_G}$ | Vmax_hk_Glc_G | 1.00E+00 | mmol/min | opt | |
| $Km_{hk}^{Glc_G}$ | Km_hk_Glc_G | 4.20E-01 | mM | exp | [4] |
| $Ki_{hk}^{G6p_G}$ | Ki_hk_G6p_G | 5.00E-01 | mM | exp | [4] |
| $Km_{hk}^{Atp_G}$ | Km_hk_Atp_G | 2.09E+00 | mM | exp | [4] |
| $Ki_{hk}^{Atp_G}$ | Ki_hk_Atp_G | 1.90E-01 | mM | exp | [4] |
| $Vmax_{g6pase}^{G6p_G}$ | Vmax_ppp_G6p_G | 0.00E+00 | mmol/min | est | |
| $Km_{ppp}^{G6p_G}$ | Km_ppp_G6p_G | 1.00E-01 | mM | est | |
| $Km_{ppp}^{Nadp_G}$ | Km_ppp_Nadp_G | 1.00E-01 | mM | est | |
| $Vmax_{g6pase}^{G6p_G}$ | Vmax_g6pase_G6p_G | 0.00E+00 | mmol/min | opt | |
| $Km_{g6pase}^{G6p_G}$ | Km_g6pase_G6p_G | 2.41E+00 | mM | exp | [5, 6] |
| $Vmax_{gs}^{G6p_G}$ | Vmax_gs_G6p_G | 0.00E+00 | mmol/min | opt | |
| $Km_{gs}^{G6p_G}$ | Km_gs_G6p_G | 5.00E-02 | mM | est | [7] |
| $n_{gs}^{G6p_G}$ | n_gs_G6p_G | 1.00E+00 | - | est | |
| $Km_{gs}^{Utp_G}$ | Km_gs_Utp_G | 4.80E-02 | mM | exp | [8] |
| $Glygn_G^{max}$ | Glygn_G_max | 3.30E+01 | mM | est | [9] |
| $Km_{gs}^{Glygn_G}$ | Km_gs_Glygn_G | 9.20E+00 | mM | est | |
| $Vmax_{gd}^{Glygn_G}$ | Vmax_gd_Glygn_G | 0.00E+00 | mmol/min | opt | |
| $Km_{gd}^{Glygn_G}$ | Km_gd_Glygn_G | 1.00E+01 | mM | est | [7] |
| $Km_{gd}^{Phos_G}$ | Km_gd_Phos_G | 4.00E+00 | mM | exp | [10] |
| $Vmax_{pfk}^{G6p_G}$ | Vmax_pfk_G6p_G | 2.50E+01 | mmol/min | opt | |
| $Km_{pfk}^{G6p_G}$ | Km_pfk_G6p_G | 5.00E-03 | mM | est | [7] |

| | | | | | |
|---|---|---|---|---|---|
| $Km_{pfk}^{Atp_G}$ | Km_pfk_Atp_G | 4.25E-02 | mM | exp | [11] |
| $Ki_{pfk}^{Atp_G}$ | Ki_pfk_Atp_G | 2.10E+00 | mM | exp | [12] |
| $Km_{pfk}^{Adp_G}$ | Km_pfk_Adp_G | 8.36E-02 | mM | exp | [11, 12] |
| $Ki_{pfk}^{Gap_G}$ | Ki_pfk_Gap_G | 2.07E-02 | mM | exp | [13] |
| $b_{pfk}^{Gap_G}$ | b_pfk_Gap_G | 7.50E-01 | - | est | [7] |
| $Vmax_{fbp}^{Gap_G}$ | Vmax_fbp_Gap_G | 0.00E+00 | mmol/min | opt | |
| $Km_{fbp}^{Gap_G}$ | Km_fbp_Gap_G | 2.50E-01 | mM | est | [7] |
| $Vmax_{pk}^{Gap_G}$ | Vmax_pk_Gap_G | 4.00E+00 | mmol/min | | |
| $Km_{pk}^{Gap_G}$ | Km_pk_Gap_G | 2.40E-01 | mM | est | [7] |
| $b_{pk}^{Accoa_{GM}}$ | b_pk_Accoa_GM | 8.00E-01 | - | exp | [14] |
| $Ki_{pk}^{Accoa_{GM}}$ | Ki_pk_Accoa_GM | 3.00E-02 | mM | exp | [14] |
| $Km_{pk}^{Adp_G}$ | Km_pk_Adp_G | 2.40E-01 | mM | exp | [15] |
| $Vmax_{pepck}^{Pyr_G}$ | Vmax_pepck_Pyr_G | 0.00E+00 | mM | opt | |
| $Km_{pepck}^{Pyr_G}$ | Km_pepck_Pyr_G | 1.00E+00 | mM | est | |
| $Km_{pepck}^{Atp_G}$ | Km_pepck_Atp_G | 1.00E-02 | mM | est | [7] |
| $Km_{pepck}^{Gtp_G}$ | Km_pepck_Gtp_G | 6.40E-02 | mM | exp | [16] |
| $Vdif_{pyrt}^{Pyr_{BG}}$ | Vdif_pyrt_Pyr_B_G | 0.00E+00 | L/min | est | |
| $Kdif_{pyrt}^{Pyr_{BG}}$ | Kdif_pyrt_Pyr_B_G | 1.00E+00 | mM | est | |
| $Kdif_{pyrt}^{Pyr_G}$ | Kdif_pyrt_Pyr_G | 1.00E+00 | mM | est | |
| $Vdif_{lact}^{Lac_{BG}}$ | Vdif_lact_Lac_B_G | 1.00E+01 | L/min | opt | |
| $Kdif_{lact}^{Lac_G}$ | Kdif_lact_Lac_G | 2.42E+00 | mM | exp | [17] |
| $Kdif_{lact}^{Lac_{GM}}$ | Kdif_lact_Lac_B_G | 2.42E+00 | mM | exp | [17] |
| $Vmax_{ldh}^{Pyr_G}$ | Vmax_ldh_Pyr_G | 1.00E+00 | $L^2$/mmol/min | opt | |
| $Km_{ldh}^{Pyr_G}$ | Km_ldh_Pyr_G | 1.50E-01 | mM | exp | [18] |
| $Km_{ldh}^{Nadh_G}$ | Km_ldh_Nadh_G | 1.50E-02 | mM | exp | [18] |
| $Km_{ldh}^{Lac_G}$ | Km_ldh_Lac_G | 3.60E+01 | mM | exp | [19] |
| $Km_{ldh}^{Nad_G}$ | Km_ldh_Nad_G | 1.10E-01 | mM | exp | [18] |

| | | | | | |
|---|---|---|---|---|---|
| $Keq_{ldh}^{Lac_G}$ | Keq_ldh_Lac_G | 1.00E+03 | - | est | |
| $Vdif_{alat}^{Ala_{BG}}$ | Vdif_alat_Ala_B_G | 0.00E+00 | L/min | est | |
| $Kdif_{alat}^{Ala_G}$ | Kdif_alat_Ala_G | 2.42E+00 | mM | est | |
| $Kdif_{alat}^{Ala_{BG}}$ | Kdif_alat_Ala_B_G | 2.42E+00 | mM | est | |
| $Vmax_{alata}^{Pyr_G}$ | Vmax_alata_Pyr_G | 0.00E+00 | L/min | opt | |
| $Km_{alata}^{Pyr_G}$ | Km_alata_Pyr_G | 1.50E-01 | mM | est | |
| $Km_{alata}^{Ala_G}$ | Km_alata_Ala_G | 3.60E+01 | mM | est | |
| $Keq_{alata}^{Ala_G}$ | Keq_alata_Ala_G | 1.00E+03 | - | est | |
| $Vmax_{pdh}^{Pyr_G}$ | Vmax_pdh_Pyr_G | 1.00E+02 | mmol/min | opt | |
| $Km_{pdh}^{Pyr_G}$ | Km_pdh_Pyr_G | 5.40E-01 | mM | est | [7] |
| $Km_{pdh}^{Accoa_{GM}}$ | Ki_pdh_Accoa_GM | 3.50E-02 | mM | exp | [20] |
| $Km_{pdh}^{Nad_G}$ | Km_pdh_Nad_G | 1.00E-02 | mM | opt | |
| $Vmax_{tca}^{Accoa_{GM}}$ | Vmax_tca_Accoa_GM | 2.00E+01 | mmol/min | opt | |
| $Km_{tca}^{Accoa_{GM}}$ | Km_tca_Accoa_GM | 4.00E-04 | mM | exp | [21] |
| $Km_{tca}^{Phos_G}$ | Km_tca_Phos_G | 1.00E+00 | mM | opt | |
| $Km_{tca}^{Adp_G}$ | Km_tca_Adp_G | 4.10E-01 | mM | exp | [22] |
| $Km_{tca}^{Pyr_G}$ | Km_tca_Pyr_G | 3.00E-02 | mM | opt | |
| $Km_{tca}^{Nad_G}$ | Km_tca_Nad_G | 1.00E-02 | mM | est | |
| $Km_{tca}^{Fad_G}$ | Km_tca_Fad_G | 1.00E-02 | mM | est | |
| $Vdif_{ffat}^{FFA_{BG}}$ | Vdif_ffat_FFA_B_G | 5.00E+00 | L/min | opt | |
| $Kdif_{ffat}^{FFA_{BG}}$ | Kdif_ffat_FFA_B_G | 2.00E-01 | mM | est | [7] |
| $Kdif_{ffat}^{FFA_G}$ | Kdif_ffat_FFA_G | 2.00E-01 | mM | est | [7] |
| $Vmax_{ffat}^{FFA_{BG}}$ | Vmax_ffat_FFA_B_G | 0.00E+00 | mmol/min | opt | |
| $Km_{ffat}^{FFA_{BG}}$ | Km_ffat_FFA_B_G | 2.00E-03 | mM | est | [7] |
| $Vmax_{tgsyn}^{FFA_G}$ | Vmax_tgsyn_FFA_G | 1.50E-01 | mmol/min | opt | |
| $Km_{tgsyn}^{FFA_G}$ | Km_tgsyn_FFA_G | 1.00E-01 | mM | est | [7] |
| $Km_{tgsyn}^{Glycp_G}$ | Km_tgsyn_Glycp_G | 4.60E-01 | mM | exp | [23] |
| $Km_{tgsyn}^{Atp_G}$ | Km_tgsyn_Atp_G | 1.00E-01 | mM | opt | |

| | | | | | |
|---|---|---|---|---|---|
| $TG_{G}^{max}$ | TG_G_max | 4.50E+02 | mM | est | |
| $Km_{tgsyn}^{TG_G}$ | Km_tgsyn_TG_G | 1.00E+01 | mM | est | |
| $Vmax_{tgdeg}^{TG_G}$ | Vmax_tgdeg_TG_G | 7.00E-02 | mmol/min | opt | |
| $Km_{tgdeg}^{TG_G}$ | Km_tgdeg_TG_G | 5.07E+01 | mM | est | [7] |
| $Vdif_{glyct}^{Glyc_{BG}}$ | Vdif_glyct_Glyc_B_G | 1.00E+01 | L/min | opt | |
| $Kdif_{glyct}^{Glyc_{BG}}$ | Kdif_glyct_Glyc_B_G | 2.70E-01 | mM | exp | [24] |
| $Kdif_{glyct}^{Glyc_G}$ | Kdif_glyct_Glyc_G | 2.70E-01 | mM | exp | [24] |
| $Vmax_{glyk}^{Glyc_G}$ | Vmax_glyk_Glyc_G | 0.00E+00 | mmol/min | opt | |
| $Km_{glyk}^{Glyc_G}$ | Km_glyk_Glyc_G | 4.00E-02 | mM | exp | [24] |
| $Km_{glyk}^{Atp_G}$ | Km_glyk_Atp_G | 5.80E-02 | mM | exp | [25] |
| $Ki_{glyk}^{Gap_G}$ | Ki_glyk_Gap_G | 5.80E-01 | mM | exp | [25] |
| $Vdif_{tgt}^{TG_{BG}}$ | Vdif_tgt_TG_B_G | 0.00E+00 | mmol/min | est | |
| $Kdif_{tgt}^{TG_{BG}}$ | Kdif_tgt_TG_B_G | 1.00E+00 | mM | est | [7] |
| $Keq_{tgt}^{TG_G}$ | Keq_tgt_TG_G | 3.38E+01 | - | est | [7] |
| $Vmax_{tgt}^{TG_{BG}}$ | Vmax_tgt_TG_B_G | 2.20E+00 | mmol/min | opt | |
| $Km_{tgt}^{TG_G}$ | Km_tgt_TG_G | 3.38E+01 | mM | est | [7] |
| $Vmax_{g3pd}^{Gap_G}$ | Vmax_g3pd_Gap_G | 2.00E+01 | $L^2$/mmol/min | opt | |
| $Km_{g3pd}^{Gap_G}$ | Km_g3pd_Gap_G | 1.60E-01 | mM | exp | [26] |
| $Km_{g3pd}^{Glycp_G}$ | Km_g3pd_Glycp_G | 2.20E-01 | mM | exp | [26] |
| $Km_{g3pd}^{Nadh_G}$ | Km_g3pd_Nadh_G | 8.00E-03 | mM | exp | [26] |
| $Km_{g3pd}^{Nad_G}$ | Km_g3pd_Nad_G | 1.30E-02 | mM | exp | [26] |
| $Keq_{g3pd}^{Glycp_G}$ | Keq_g3pd_Glycp_G | 1.00E+03 | - | opt | |
| $Vmax_{boxid}^{FFA_G}$ | Vmax_boxid_FFA_G | 5.00E-02 | mmol/min | opt | |
| $Km_{boxid}^{FFA_G}$ | Km_boxid_FFA_G | 5.00E-03 | mM | exp | [27-29] |
| $Km_{boxid}^{Atp_G}$ | Km_boxid_Atp_G | 8.70E-02 | mM | exp | [30] |
| $Ki_{boxid}^{Accoa_{GM}}$ | Ki_boxid_Accoa_GM | 1.20E-01 | mM | | [31] |
| $Ki_{boxid}^{Malcoa_G}$ | Ki_boxid_Malcoa_G | 1.00E+00 | mM | opt | |

| | | | | |
|---|---|---|---|---|
| $Km_{boxid}^{Nad_G}$ | Km_boxid_Nad_G | 1.00E-01 | mM | est |
| $Km_{boxid}^{Fad_G}$ | Km_boxid_Fad_G | 1.00E-01 | mM | est |
| $Vmax_{accoat}^{Accoa_{GMGC}}$ | Vmax_accoat_Accoa_GM_GC | 0.00E+00 | mmol/min | est |
| $Km_{accoat}^{Accoa_{GM}}$ | Km_accoat_Accoa_GM | 5.80E-02 | mM | opt |
| $Km_{accoat}^{Atp_G}$ | Km_accoat_Atp_G | 1.00E-01 | mM | est |
| $Km_{accoat}^{Pyr_G}$ | Km_accoat_Pyr_G | 1.00E+00 | mM | est |
| $Vmax_{bhbsyn}^{Accoa_{GM}}$ | Vmax_bhbsyn_Accoa_GM | 0.00E+00 | mmol/min | opt |
| $Km_{bhbsyn}^{Accoa_{GM}}$ | Km_bhbsyn_Accoa_GM | 1.00E-01 | mM | est |
| $Km_{bhbsyn}^{Nadh_G}$ | Km_bhbsyn_Nadh_G | 1.00E-01 | mM | est |
| $n_{bhbsyn}^{Pyr_G}$ | n_bhbsyn_Pyr_G | 4.00E+00 | - | est |
| $Ki_{bhbsyn}^{Pyr_G}$ | Ki_bhbsyn_Pyr_G | 1.00E-02 | mM | opt |
| $Vmax_{bhbdeg}^{Bhb_G}$ | Vmax_bhbdeg_Bhb_G | 0.00E+00 | mmol/min | est |
| $Km_{bhbdeg}^{Bhb_G}$ | Km_bhbdeg_Bhb_G | 1.00E-01 | mM | est |
| $Km_{bhbdeg}^{Atp_G}$ | Km_bhbdeg_Atp_G | 1.00E-01 | mM | est |
| $Km_{bhbdeg}^{Nad_G}$ | Km_bhbdeg_Nad_G | 1.00E-01 | mM | est |
| $Vdif_{bhbt}^{Bhb_{BG}}$ | Vdif_bhbt_Bhb_B_G | 0.00E+00 | L/min | est |
| $Kdif_{bhbt}^{Bhb_{BG}}$ | Kdif_bhbt_Bhb_B_G | 1.00E+00 | mM | est |
| $Kdif_{bhbt}^{Bhb_G}$ | Kdif_bhbt_Bhb_G | 1.00E+00 | mM | est |
| $Vmax_{lipog1}^{Accoa_{GC}}$ | Vmax_lipog1_Accoa_GC | 0.00E+00 | mmol/min | opt |
| $Km_{lipog1}^{Accoa_{GC}}$ | Km_lipog1_Accoa_GC | 1.00E-02 | mM | est |
| $Km_{lipog1}^{Atp_G}$ | Km_lipog1_Atp_G | 1.00E-01 | mM | est |
| $Vmax_{lipog2}^{Malcoa_G}$ | Vmax_lipog2_Malcoa_G | 0.00E+00 | mmol/min | est |
| $Km_{lipog2}^{Malcoa_G}$ | Km_lipog2_Malcoa_G | 1.00E-02 | mM | est |
| $Km_{lipog2}^{Adp_G}$ | Km_lipog2_Adp_G | 1.00E-01 | mM | est |
| $Km_{lipog2}^{Nadph_G}$ | Km_lipog2_Nadph_G | 1.00E-02 | mM | est |
| $Vmax_{cholsyn1}^{Accoa_{GC}}$ | Vmax_cholsyn1_Accoa_GC | 0.00E+00 | mmol/min | est |
| $Km_{cholsyn1}^{Accoa_{GC}}$ | Km_cholsyn1_Accoa_GC | 1.00E-01 | mM | est |
| $Vmax_{cholsyn2}^{Hmgcoa_G}$ | Vmax_cholsyn2_Hmgcoa_G | 0.00E+00 | mmol/min | est |

| | | | | | |
|---|---|---|---|---|---|
| $Km_{cholsyn2}^{Hmgcoa_G}$ | Km_cholsyn2_Hmgcoa_G | 1.00E-01 | mM | est | |
| $Km_{cholsyn2}^{Atp_G}$ | Km_cholsyn2_Atp_G | 1.00E-01 | mM | est | |
| $Km_{cholsyn2}^{Fadh_G}$ | Km_cholsyn2_Fadh_G | 1.00E-01 | mM | est | |
| $Km_{cholsyn2}^{Nadph_G}$ | Km_cholsyn2_Nadph_G | 1.00E-01 | mM | est | |
| $Vmax_{cholt}^{Chol_{BG}}$ | Vmax_cholt_Chol_G | 0.00E+00 | mmol/min | est | |
| $Km_{cholt}^{Chol_G}$ | Km_cholt_Chol_G | 1.00E-01 | mM | est | |
| $Vmax_{atpsynf}^{Fadh_G}$ | Vmax_atpsynf_Fadh_G | 4.30E-01 | mmol/min | opt | |
| $Km_{atpsynf}^{Fadh_G}$ | Km_atpsynf_Fadh_G | 1.00E-01 | mM | est | |
| $Km_{atpsynf}^{Adp_G}$ | Km_atpsynf_Adp_G | 1.00E-01 | mM | est | |
| $Vmax_{atpsynn}^{Nadh_G}$ | Vmax_atpsynn_Nadh_G | 1.00E+00 | mmol/min | est | |
| $Km_{atpsynn}^{Nadh_G}$ | Km_atpsynn_Nadh_G | 1.00E-01 | mM | est | |
| $Km_{atpsynn}^{Adp_G}$ | Km_atpsynn_Adp_G | 1.00E-01 | mM | est | |
| $Vmax_{atpuse}^{Atp_G}$ | Vmax_atpuse_Atp_G | 3.90E+00 | mmol/min | opt | |
| $Km_{atpuse}^{Atp_G}$ | Km_atpuse_Atp_G | 2.50E+00 | mM | opt | |
| $Vmax_{ampreg}^{Amp_G}$ | Vmax_ampreg_Amp_G | 1.00E+01 | mmol/min | est | |
| $Km_{ampreg}^{Amp_G}$ | Km_ampreg_Amp_G | 8.00E-02 | mM | exp | [32] |
| $Km_{ampreg}^{Atp_G}$ | Km_ampreg_Atp_G | 9.00E-02 | mM | exp | [32] |
| $Km_{ampreg}^{Adp_G}$ | Km_ampreg_Adp_G | 1.10E-01 | mM | exp | [32] |
| $Vmax_{nadhk}^{Nadh_G}$ | Vmax_nadhk_Nadh_G | 0.00E+00 | mmol/min | est | |
| $Km_{nadhk}^{Nadh_G}$ | Km_nadhk_Nadh_G | 1.00E-01 | mM | est | |
| $Km_{nadhk}^{Atp_G}$ | Km_nadhk_Atp_A | 1.00E-01 | mM | est | |
| $Vmax_{nadhuse}^{Nadh_G}$ | Vmax_nadhuse_Nadh_G | 1.20E+00 | mmol/min | est | |
| $Km_{nadhuse}^{Nadh_G}$ | Km_nadhuse_Nadh_G | 1.00E-01 | mM | est | |
| $Vmax_{gdpreg}^{Gdp_G}$ | Vmax_gdpreg_Gdp_G | 0.00E+00 | $L^2$/mmol/min | est | |
| $Km_{gdpreg}^{Gdp_G}$ | Km_gdpreg_Gdp_G | 3.10E-02 | mM | exp | [33] |
| $Km_{gdpreg}^{Atp_G}$ | Km_gdpreg_Atp_G | 1.33E+00 | mM | exp | [33] |
| $Km_{gdpreg}^{Gtp_G}$ | Km_gdpreg_Gtp_G | 1.50E-01 | mM | exp | [34] |
| $Km_{gdpreg}^{Adp_G}$ | Km_gdpreg_Adp_G | 4.20E-02 | mM | exp | [33] |

| | | | | | |
|---|---|---|---|---|---|
| $Keq_{gdpreg}^{Gtp_G}$ | Keq_gdpreg_Gtp_G | 1.00E+03 | - | opt | |
| $Vmax_{gtpuse}^{Gtp_G}$ | Vmax_gtpuse_Gtp_G | 0.00E+00 | mmol/min | est | |
| $Km_{gtpuse}^{Gtp_G}$ | Km_gtpuse_Gtp_G | 1.00E-01 | mM | est | |
| $Vmax_{udpreg}^{Udp_G}$ | Vmax_udpreg_Udp_G | 1.00E+02 | $L^2$/mmol/min | est | |
| $Km_{udpreg}^{Udp_G}$ | Km_udpreg_Udp_G | 1.90E-01 | mM | exp | [33] |
| $Km_{udpreg}^{Atp_G}$ | Km_udpreg_Atp_G | 1.33E+00 | mM | exp | [33] |
| $Km_{udpreg}^{Utp_G}$ | Km_udpreg_Utp_G | 1.60E+01 | mM | exp | [34] |
| $Km_{udpreg}^{Adp_G}$ | Km_udpreg_Adp_G | 4.20E-02 | mM | exp | [33] |
| $Keq_{udpreg}^{Utp_G}$ | Keq_udpreg_Utp_G | 1.00E+03 | - | est | |
| $Vmax_{utpuse}^{Utp_G}$ | Vmax_utpuse_Utp_G | 0.00E+00 | mmol/min | est | |
| $Km_{utpuse}^{Utp_G}$ | Km_utpuse_Utp_G | 1.00E-01 | mM | est | |
| $Vmax_{nadphuse}^{Nadph_G}$ | Vmax_nadphuse_Nadph_G | 0.00E+00 | mmol/min | est | |
| $Km_{nadphuse}^{Nadph_G}$ | Km_nadphuse_Nadph_G | 1.00E-01 | mM | est | |
| $Vmax_{fadhuse}^{Fadh_G}$ | Vmax_fadhuse_Fadh_G | 5.00E-01 | mmol/min | est | |
| $Km_{fadhuse}^{Fadh_G}$ | Km_fadhuse_Fadh_G | 1.00E-01 | mM | est | |
| $Vmax_{ck}^{Cre_G}$ | Vmax_ck_Cre_G | 8.60E+01 | $L^2$/mmol/min | est | |
| $Km_{ck}^{Cre_G}$ | Km_ck_Cre_G | 1.00E+00 | mM | est | |
| $Km_{ck}^{Atp_G}$ | Km_ck_Atp_G | 3.00E-02 | mM | est | |
| $Km_{ck}^{Crep_G}$ | Km_ck_Crep_G | 1.50E+01 | mM | est | |
| $Km_{ck}^{Adp_G}$ | Km_ck_Adp_G | 1.00E-01 | mM | est | |
| $Keq_{ck}^{Crep_G}$ | Keq_ck_Crep_G | 1.00E+00 | - | est | |
| $Phos_G$ | Phos_G | 5.00E+00 | mM | est | |
| $\alpha_H^{Base}$ | alpha_base_H | 1.00E-01 | - | opt | |
| $\alpha_H^{Band}$ | alpha_band_H | 1.00E-01 | - | opt | |
| $Km^{Ins_{BH}}$ | Km_Ins_B_H | 8.00E-08 | mM | est | |
| $n^{Ins_{BH}}$ | n_Ins_B_H | 4.00E+00 | - | est | |
| $Vdif_{glut4}^{Glc_{BH}}$ | Vdif_glut2_Glc_B_H | 1.61E-02 | L/min | opt | |
| $Kdif_{glut4}^{Glc_{BH}}$ | Kdif_glut2_Glc_B_H | 5.00E+00 | mM | exp | [3] |

| | | | | | |
|---|---|---|---|---|---|
| $Kdif_{glut4}^{Glc_H}$ | Kdif_glut2_Glc_H | 5.00E+00 | mM | exp | [3] |
| $Vmax_{hk}^{Glc_H}$ | Vmax_hk_Glc_H | 5.03E+00 | mmol/min | opt | |
| $Km_{hk}^{Glc_H}$ | Km_hk_Glc_H | 4.20E-01 | mM | exp | [4] |
| $Ki_{hk}^{G6p_H}$ | Ki_hk_G6p_H | 5.00E-01 | mM | exp | [4] |
| $Km_{hk}^{Atp_H}$ | Km_hk_Atp_H | 2.09E+00 | mM | exp | [4] |
| $Ki_{hk}^{Atp_H}$ | Ki_hk_Atp_H | 1.90E-01 | mM | exp | [4] |
| $Vmax_{g6pase}^{G6p_H}$ | Vmax_ppp_G6p_H | 2.00E-01 | mmol/min | est | |
| $Km_{ppp}^{G6p_H}$ | Km_ppp_G6p_H | 1.00E-01 | mM | est | |
| $Km_{ppp}^{Nadp_H}$ | Km_ppp_Nadp_H | 1.00E-01 | mM | est | |
| $Vmax_{g6pase}^{G6p_H}$ | Vmax_g6pase_G6p_H | 0.00E+00 | mmol/min | opt | |
| $Km_{g6pase}^{G6p_H}$ | Km_g6pase_G6p_H | 2.41E+00 | mM | exp | [5, 6] |
| $Vmax_{gs}^{G6p_H}$ | Vmax_gs_G6p_H | 2.79E+00 | mmol/min | opt | |
| $Km_{gs}^{G6p_H}$ | Km_gs_G6p_H | 5.00E-02 | mM | est | [7] |
| $n_{gs}^{G6p_H}$ | n_gs_G6p_H | 4.21E-02 | - | est | |
| $Km_{gs}^{Utp_H}$ | Km_gs_Utp_H | 4.80E-02 | mM | exp | [8] |
| $Glygn_{H}^{max}$ | Glygn_H_max | 3.75E+02 | mM | est | [9] |
| $Km_{gs}^{Glygn_H}$ | Km_gs_Glygn_H | 7.50E+01 | mM | est | |
| $Vmax_{gd}^{Glygn_H}$ | Vmax_gd_Glygn_H | 8.58E+00 | mmol/min | opt | |
| $Km_{gd}^{Glygn_H}$ | Km_gd_Glygn_H | 1.00E+01 | mM | est | [7] |
| $Km_{gd}^{Phos_H}$ | Km_gd_Phos_H | 4.00E+00 | mM | exp | [10] |
| $Vmax_{pfk}^{G6p_H}$ | Vmax_pfk_G6p_H | 2.00E+01 | mmol/min | opt | |
| $Km_{pfk}^{G6p_H}$ | Km_pfk_G6p_H | 5.00E-03 | mM | est | [7] |
| $Km_{pfk}^{Atp_H}$ | Km_pfk_Atp_H | 4.25E-02 | mM | exp | [11] |
| $Ki_{pfk}^{Atp_H}$ | Ki_pfk_Atp_H | 2.10E+00 | mM | exp | [12] |
| $Km_{pfk}^{Adp_H}$ | Km_pfk_Adp_H | 8.36E-02 | mM | exp | [11, 12] |
| $Ki_{pfk}^{Gap_H}$ | Ki_pfk_Gap_H | 2.07E-02 | mM | exp | [13] |
| $b_{pfk}^{Gap_H}$ | b_pfk_Gap_H | 7.50E-01 | - | est | [7] |

| | | | | | |
|---|---|---|---|---|---|
| $Vmax_{fbp}^{Gap_H}$ | Vmax_fbp_Gap_H | 0.00E+00 | mmol/min | opt | |
| $Km_{fbp}^{Gap_H}$ | Km_fbp_Gap_H | 2.50E-01 | mM | est | [7] |
| $Vmax_{pk}^{Gap_H}$ | Vmax_pk_Gap_H | 5.00E+01 | mmol/min | | |
| $Km_{pk}^{Gap_H}$ | Km_pk_Gap_H | 2.40E-01 | mM | est | [7] |
| $b_{pk}^{Accoa_{HM}}$ | b_pk_Accoa_HM | 8.00E-01 | - | exp | [14] |
| $Ki_{pk}^{Accoa_{HM}}$ | Ki_pk_Accoa_HM | 3.00E-02 | mM | exp | [14] |
| $Km_{pk}^{Adp_H}$ | Km_pk_Adp_H | 2.40E-01 | mM | exp | [15] |
| $Vmax_{pepck}^{Pyr_H}$ | Vmax_pepck_Pyr_H | 0.00E+00 | mM | opt | |
| $Km_{pepck}^{Pyr_H}$ | Km_pepck_Pyr_H | 6.29E-01 | mM | est | |
| $Km_{pepck}^{Atp_H}$ | Km_pepck_Atp_H | 1.00E-02 | mM | est | [7] |
| $Km_{pepck}^{Gtp_H}$ | Km_pepck_Gtp_H | 6.40E-02 | mM | exp | [16] |
| $Vdif_{pyrt}^{Pyr_{BH}}$ | Vdif_pyrt_Pyr_B_H | 0.00E+00 | L/min | est | |
| $Kdif_{pyrt}^{Pyr_{BH}}$ | Kdif_pyrt_Pyr_B_H | 1.00E+00 | mM | est | |
| $Kdif_{pyrt}^{Pyr_H}$ | Kdif_pyrt_Pyr_H | 1.00E+00 | mM | est | |
| $Vdif_{lact}^{Lac_{BH}}$ | Vdif_lact_Lac_B_H | 8.48E-02 | L/min | opt | |
| $Kdif_{lact}^{Lac_H}$ | Kdif_lact_Lac_H | 2.42E+00 | mM | exp | [17] |
| $Kdif_{lact}^{Lac_{BH}}$ | Kdif_lact_Lac_B_H | 2.42E+00 | mM | exp | [17] |
| $Vmax_{ldh}^{Pyr_H}$ | Vmax_ldh_Pyr_H | 2.00E+01 | $L^2$/mmol/min | opt | |
| $Km_{ldh}^{Pyr_H}$ | Km_ldh_Pyr_H | 1.50E-01 | mM | exp | [18] |
| $Km_{ldh}^{Nadh_H}$ | Km_ldh_Nadh_H | 1.50E-02 | mM | exp | [18] |
| $Km_{ldh}^{Lac_H}$ | Km_ldh_Lac_H | 3.60E+01 | mM | exp | [19] |
| $Km_{ldh}^{Nad_H}$ | Km_ldh_Nad_H | 1.10E-01 | mM | exp | [18] |
| $Keq_{ldh}^{Lac_H}$ | Keq_ldh_Lac_H | 5.00E-01 | - | est | |
| $Vdif_{alat}^{Ala_{BH}}$ | Vdif_alat_Ala_B_H | 0.00E+00 | L/min | est | |
| $Kdif_{alat}^{Ala_H}$ | Kdif_alat_Ala_H | 2.42E+00 | mM | est | |
| $Kdif_{alat}^{Ala_{BH}}$ | Kdif_alat_Ala_B_H | 2.42E+00 | mM | est | |
| $Vmax_{alata}^{Pyr_H}$ | Vmax_alata_Pyr_H | 0.00E+00 | L/min | opt | |
| $Km_{alata}^{Pyr_H}$ | Km_alata_Pyr_H | 1.50E-01 | mM | est | |

| | | | | | |
|---|---|---|---|---|---|
| $Km_{alata}^{Ala_H}$ | Km_alata_Ala_H | 3.60E+01 | mM | est | |
| $Keq_{alata}^{Ala_H}$ | Keq_alata_Ala_H | 1.00E+00 | - | est | |
| $Vmax_{pdh}^{Pyr_H}$ | Vmax_pdh_Pyr_H | 1.00E+01 | mmol/min | opt | |
| $Km_{pdh}^{Pyr_H}$ | Km_pdh_Pyr_H | 5.40E-01 | mM | est | [7] |
| $Km_{pdh}^{Accoa_{HM}}$ | Ki_pdh_Accoa_HM | 3.50E-02 | mM | exp | [20] |
| $Km_{pdh}^{Nad_H}$ | Km_pdh_Nad_H | 1.00E-02 | mM | opt | |
| $Vmax_{tca}^{Accoa_{HM}}$ | Vmax_tca_Accoa_HM | 2.49E+01 | mmol/min | opt | |
| $Km_{tca}^{Accoa_{HM}}$ | Km_tca_Accoa_HM | 4.00E-04 | mM | exp | [21] |
| $Km_{tca}^{Phos_H}$ | Km_tca_Phos_H | 8.05E-02 | mM | opt | |
| $Km_{tca}^{Adp_H}$ | Km_tca_Adp_H | 4.10E-01 | mM | exp | [22] |
| $Km_{tca}^{Pyr_H}$ | Km_tca_Pyr_H | 1.11E-01 | mM | opt | |
| $Km_{tca}^{Nad_H}$ | Km_tca_Nad_H | 1.00E-01 | mM | est | |
| $Km_{tca}^{Fad_H}$ | Km_tca_Fad_H | 1.00E-01 | mM | est | |
| $Vdif_{ffat}^{FFA_{BH}}$ | Vdif_ffat_FFA_B_H | 2.48E+00 | L/min | opt | |
| $Kdif_{ffat}^{FFA_{BH}}$ | Kdif_ffat_FFA_B_H | 2.00E-01 | mM | est | [7] |
| $Kdif_{ffat}^{FFA_H}$ | Kdif_ffat_FFA_H | 2.00E-01 | mM | est | [7] |
| $Vmax_{ffat}^{FFA_{BH}}$ | Vmax_ffat_FFA_B_H | 4.41E-02 | mmol/min | opt | |
| $Km_{ffat}^{FFA_{BH}}$ | Km_ffat_FFA_B_H | 2.00E-03 | mM | est | [7] |
| $Vmax_{tgsyn}^{FFA_H}$ | Vmax_tgsyn_FFA_H | 1.00E-04 | mmol/min | opt | |
| $Km_{tgsyn}^{FFA_H}$ | Km_tgsyn_FFA_H | 6.45E-01 | mM | est | [7] |
| $Km_{tgsyn}^{Glycp_H}$ | Km_tgsyn_Glycp_H | 4.60E-01 | mM | exp | [23] |
| $Km_{tgsyn}^{Atp_H}$ | Km_tgsyn_Atp_H | 1.00E-01 | mM | opt | |
| $TG_H^{max}$ | TG_H_max | 3.12E+00 | mM | est | |
| $Km_{tgsyn}^{TG_H}$ | Km_tgsyn_TG_H | 1.00E+01 | mM | est | |
| $Vmax_{tgdeg}^{TG_H}$ | Vmax_tgdeg_TG_H | 2.69E-03 | mmol/min | opt | |
| $Km_{tgdeg}^{TG_H}$ | Km_tgdeg_TG_H | 5.07E+01 | mM | est | [7] |
| $Vdif_{glyct}^{Glyc_{BH}}$ | Vdif_glyct_Glyc_B_H | 1.00E+00 | L/min | opt | |
| $Kdif_{glyct}^{Glyc_{BH}}$ | Kdif_glyct_Glyc_B_H | 2.70E-01 | mM | exp | [24] |

| | | | | | |
|---|---|---|---|---|---|
| $Kdif_{glyct}^{Glyc_H}$ | Kdif_glyct_Glyc_H | 2.70E-01 | mM | exp | [24] |
| $Vmax_{glyk}^{Glyc_H}$ | Vmax_glyk_Glyc_H | 1.30E+00 | mmol/min | opt | |
| $Km_{glyk}^{Glyc_H}$ | Km_glyk_Glyc_H | 1.00E-03 | mM | exp | [24] |
| $Km_{glyk}^{Atp_H}$ | Km_glyk_Atp_H | 1.50E-02 | mM | exp | [25] |
| $Ki_{glyk}^{Gap_H}$ | Ki_glyk_Gap_H | 1.00E+00 | mM | exp | [25] |
| $Vdif_{tgt}^{TG_{BH}}$ | Vdif_tgt_TG_B_H | 0.00E+00 | mmol/min | est | |
| $Kdif_{tgt}^{TG_{BH}}$ | Kdif_tgt_TG_B_H | 1.00E+00 | mM | est | [7] |
| $Keq_{tgt}^{TG_H}$ | Keq_tgt_TG_H | 3.38E+01 | - | est | [7] |
| $Vmax_{tgt}^{TG_{BH}}$ | Vmax_tgt_TG_B_H | 0.00E+00 | mmol/min | opt | |
| $Km_{tgt}^{TG_H}$ | Km_tgt_TG_H | 3.38E+01 | mM | est | [7] |
| $Vmax_{g3pd}^{Gap_H}$ | Vmax_g3pd_Gap_H | 0.00E+00 | $L^2$/mmol/min | opt | |
| $Km_{g3pd}^{Gap_H}$ | Km_g3pd_Gap_H | 1.00E+00 | mM | exp | [26] |
| $Km_{g3pd}^{Glycp_H}$ | Km_g3pd_Glycp_H | 1.00E+00 | mM | exp | [26] |
| $Km_{g3pd}^{Nadh_H}$ | Km_g3pd_Nadh_H | 1.00E+00 | mM | exp | [26] |
| $Km_{g3pd}^{Nad_H}$ | Km_g3pd_Nad_H | 1.00E+00 | mM | exp | [26] |
| $Keq_{g3pd}^{Glycp_H}$ | Keq_g3pd_Glycp_H | 1.34E+03 | - | opt | |
| $Vmax_{boxid}^{FFA_H}$ | Vmax_boxid_FFA_H | 6.00E-01 | mmol/min | opt | |
| $Km_{boxid}^{FFA_H}$ | Km_boxid_FFA_H | 5.00E-03 | mM | exp | [27-29] |
| $Km_{boxid}^{Atp_H}$ | Km_boxid_Atp_H | 8.70E-02 | mM | exp | [30] |
| $Ki_{boxid}^{Accoa_{HM}}$ | Ki_boxid_Accoa_HM | 1.20E-01 | mM | | [31] |
| $Ki_{boxid}^{Malcoa_H}$ | Ki_boxid_Malcoa_H | 1.00E+00 | mM | opt | |
| $Km_{boxid}^{Nad_H}$ | Km_boxid_Nad_H | 1.00E-01 | mM | est | |
| $Km_{boxid}^{Fad_H}$ | Km_boxid_Fad_H | 1.00E-01 | mM | est | |
| $Vmax_{accoat}^{Accoa_{HMHC}}$ | Vmax_accoat_Accoa_HM_HC | 0.00E+00 | mmol/min | est | |
| $Km_{accoat}^{Accoa_{HM}}$ | Km_accoat_Accoa_HM | 5.80E-02 | mM | opt | |
| $Km_{accoat}^{Atp_H}$ | Km_accoat_Atp_H | 1.00E-01 | mM | est | |
| $Km_{accoat}^{Pyr_H}$ | Km_accoat_Pyr_H | 1.00E+00 | mM | est | |
| $Vmax_{bhbsyn}^{Accoa_{HM}}$ | Vmax_bhbsyn_Accoa_HM | 0.00E+00 | mmol/min | opt | |

| | | | | |
|---|---|---|---|---|
| $Km_{bhbsyn}^{Accoa_{HM}}$ | Km_bhbsyn_Accoa_HM | 1.00E-01 | mM | est |
| $Km_{bhbsyn}^{Nadh_{H}}$ | Km_bhbsyn_Nadh_H | 1.00E-01 | mM | est |
| $n_{bhbsyn}^{Pyr_{H}}$ | n_bhbsyn_Pyr_H | 4.00E+00 | - | est |
| $Ki_{bhbsyn}^{Pyr_{H}}$ | Ki_bhbsyn_Pyr_H | 1.00E-02 | mM | opt |
| $Vmax_{bhbdeg}^{Bhb_{H}}$ | Vmax_bhbdeg_Bhb_H | 0.00E+00 | mmol/min | est |
| $Km_{bhbdeg}^{Bhb_{H}}$ | Km_bhbdeg_Bhb_H | 1.00E-01 | mM | est |
| $Km_{bhbdeg}^{Atp_{H}}$ | Km_bhbdeg_Atp_H | 1.00E-01 | mM | est |
| $Km_{bhbdeg}^{Nad_{H}}$ | Km_bhbdeg_Nad_H | 1.00E-01 | mM | est |
| $Vdif_{bhbt}^{Bhb_{BH}}$ | Vdif_bhbt_Bhb_B_H | 0.00E+00 | L/min | est |
| $Kdif_{bhbt}^{Bhb_{BH}}$ | Kdif_bhbt_Bhb_B_H | 1.00E+00 | mM | est |
| $Kdif_{bhbt}^{Bhb_{H}}$ | Kdif_bhbt_Bhb_H | 1.00E+00 | mM | est |
| $Vmax_{lipog1}^{Accoa_{HC}}$ | Vmax_lipog1_Accoa_HC | 0.00E+00 | mmol/min | opt |
| $Km_{lipog1}^{Accoa_{HC}}$ | Km_lipog1_Accoa_HC | 1.00E-02 | mM | est |
| $Km_{lipog1}^{Atp_{H}}$ | Km_lipog1_Atp_H | 1.00E-01 | mM | est |
| $Vmax_{lipog2}^{Malcoa_{H}}$ | Vmax_lipog2_Malcoa_H | 0.00E+00 | mmol/min | est |
| $Km_{lipog2}^{Malcoa_{H}}$ | Km_lipog2_Malcoa_H | 1.00E-02 | mM | est |
| $Km_{lipog2}^{Adp_{H}}$ | Km_lipog2_Adp_H | 1.00E-01 | mM | est |
| $Km_{lipog2}^{Nadph_{H}}$ | Km_lipog2_Nadph_H | 1.00E-02 | mM | est |
| $Vmax_{cholsyn1}^{Accoa_{HC}}$ | Vmax_cholsyn1_Accoa_HC | 0.00E+00 | mmol/min | est |
| $Km_{cholsyn1}^{Accoa_{HC}}$ | Km_cholsyn1_Accoa_HC | 1.00E-01 | mM | est |
| $Vmax_{cholsyn2}^{Hmgcoa_{H}}$ | Vmax_cholsyn2_Hmgcoa_H | 0.00E+00 | mmol/min | est |
| $Km_{cholsyn2}^{Hmgcoa_{H}}$ | Km_cholsyn2_Hmgcoa_H | 1.00E-01 | mM | est |
| $Km_{cholsyn2}^{Atp_{H}}$ | Km_cholsyn2_Atp_H | 1.00E-01 | mM | est |
| $Km_{cholsyn2}^{Fadh_{H}}$ | Km_cholsyn2_Fadh_H | 1.00E-01 | mM | est |
| $Km_{cholsyn2}^{Nadph_{H}}$ | Km_cholsyn2_Nadph_H | 1.00E-01 | mM | est |
| $Vmax_{cholt}^{Chol_{BH}}$ | Vmax_cholt_Chol_H | 0.00E+00 | mmol/min | est |
| $Km_{cholt}^{Chol_{H}}$ | Km_cholt_Chol_H | 1.00E-01 | mM | est |

| | | | | | |
|---|---|---|---|---|---|
| $Vmax_{atpsynf}^{Fadh_H}$ | Vmax_atpsynf_Fadh_H | 5.00E+00 | mmol/min | opt | |
| $Km_{atpsynf}^{Fadh_H}$ | Km_atpsynf_Fadh_H | 1.00E-01 | mM | est | |
| $Km_{atpsynf}^{Adp_H}$ | Km_atpsynf_Adp_H | 1.00E-01 | mM | est | |
| $Vmax_{atpsynn}^{Nadh_H}$ | Vmax_atpsynn_Nadh_H | 2.85E+01 | mmol/min | est | |
| $Km_{atpsynn}^{Nadh_H}$ | Km_atpsynn_Nadh_H | 1.00E-01 | mM | est | |
| $Km_{atpsynn}^{Adp_H}$ | Km_atpsynn_Adp_H | 1.00E-01 | mM | est | |
| $Vmax_{atpuse}^{Atp_H}$ | Vmax_atpuse_Atp_H | 1.00E+01 | mmol/min | opt | |
| $Km_{atpuse}^{Atp_H}$ | Km_atpuse_Atp_H | 2.50E+00 | mM | opt | |
| $Vmax_{ampreg}^{Amp_H}$ | Vmax_ampreg_Amp_H | 1.00E+01 | mmol/min | est | |
| $Km_{ampreg}^{Amp_H}$ | Km_ampreg_Amp_H | 8.00E-02 | mM | exp | [32] |
| $Km_{ampreg}^{Atp_H}$ | Km_ampreg_Atp_H | 9.00E-02 | mM | exp | [32] |
| $Km_{ampreg}^{Adp_H}$ | Km_ampreg_Adp_H | 1.10E-01 | mM | exp | [32] |
| $Vmax_{nadhk}^{Nadh_H}$ | Vmax_nadhk_Nadh_H | 0.00E+00 | mmol/min | est | |
| $Km_{nadhk}^{Nadh_H}$ | Km_nadhk_Nadh_H | 1.00E-01 | mM | est | |
| $Km_{nadhk}^{Atp_H}$ | Km_nadhk_Atp_H | 1.00E-01 | mM | est | |
| $Vmax_{nadhuse}^{Nadh_H}$ | Vmax_nadhuse_Nadh_H | 2.56E-01 | mmol/min | est | |
| $Km_{nadhuse}^{Nadh_H}$ | Km_nadhuse_Nadh_H | 1.00E-01 | mM | est | |
| $Vmax_{gdpreg}^{Gdp_H}$ | Vmax_gdpreg_Gdp_H | 1.00E+01 | $L^2$/mmol/min | est | |
| $Km_{gdpreg}^{Gdp_H}$ | Km_gdpreg_Gdp_H | 3.35E-02 | mM | exp | [33] |
| $Km_{gdpreg}^{Atp_H}$ | Km_gdpreg_Atp_H | 2.90E-01 | mM | exp | [33] |
| $Km_{gdpreg}^{Gtp_H}$ | Km_gdpreg_Gtp_H | 1.20E-01 | mM | exp | [34] |
| $Km_{gdpreg}^{Adp_H}$ | Km_gdpreg_Adp_H | 2.40E-02 | mM | exp | [33] |
| $Keq_{gdpreg}^{Gtp_H}$ | Keq_gdpreg_Gtp_H | 1.00E+03 | - | opt | |
| $Vmax_{gtpuse}^{Gtp_H}$ | Vmax_gtpuse_Gtp_H | 0.00E+00 | mmol/min | est | |
| $Km_{gtpuse}^{Gtp_H}$ | Km_gtpuse_Gtp_H | 1.00E-01 | mM | est | |
| $Vmax_{udpreg}^{Udp_H}$ | Vmax_udpreg_Udp_H | 1.00E+01 | $L^2$/mmol/min | est | |
| $Km_{udpreg}^{Udp_H}$ | Km_udpreg_Udp_H | 1.75E-01 | mM | exp | [33] |
| $Km_{udpreg}^{Atp_H}$ | Km_udpreg_Atp_H | 2.90E-01 | mM | exp | [33] |

| | | | | | |
|---|---|---|---|---|---|
| $Km_{udpreg}^{Utp_H}$ | Km_udpreg_Utp_H | 2.15E-02 | mM | exp | [34] |
| $Km_{udpreg}^{Adp_H}$ | Km_udpreg_Adp_H | 2.40E-02 | mM | exp | [33] |
| $Keq_{udpreg}^{Utp_H}$ | Keq_udpreg_Utp_H | 1.00E+03 | - | est | |
| $Vmax_{utpuse}^{Utp_H}$ | Vmax_utpuse_Utp_H | 0.00E+00 | mmol/min | est | |
| $Km_{utpuse}^{Utp_H}$ | Km_utpuse_Utp_H | 1.00E-01 | mM | est | |
| $Vmax_{nadphuse}^{Nadph_H}$ | Vmax_nadphuse_Nadph_H | 0.00E+00 | mmol/min | est | |
| $Km_{nadphuse}^{Nadph_H}$ | Km_nadphuse_Nadph_H | 1.00E-01 | mM | est | |
| $Vmax_{fadhuse}^{Fadh_H}$ | Vmax_fadhuse_Fadh_H | 0.00E+00 | mmol/min | est | |
| $Km_{fadhuse}^{Fadh_H}$ | Km_fadhuse_Fadh_H | 1.00E-01 | mM | est | |
| $Vmax_{ck}^{Cre_H}$ | Vmax_ck_Cre_H | 0.00E+00 | $L^2$/mmol/min | est | |
| $Km_{ck}^{Cre_H}$ | Km_ck_Cre_H | 1.00E+00 | mM | est | |
| $Km_{ck}^{Atp_H}$ | Km_ck_Atp_H | 3.00E-02 | mM | est | |
| $Km_{ck}^{Crep_H}$ | Km_ck_Crep_H | 1.50E+01 | mM | est | |
| $Km_{ck}^{Adp_H}$ | Km_ck_Adp_H | 1.00E-01 | mM | est | |
| $Keq_{ck}^{Crep_H}$ | Keq_ck_Crep_H | 1.00E+00 | - | est | |
| $Phos_H$ | Phos_H | 5.00E+00 | mM | est | |
| $\alpha_N^{Base}$ | alpha_base_N | 1.00E-01 | - | opt | |
| $\alpha_N^{Band}$ | alpha_band_N | 1.00E-01 | - | opt | |
| $Km^{Ins_{BN}}$ | Km_Ins_B_N | 8.00E-08 | mM | est | |
| $n^{Ins_{BN}}$ | n_Ins_B_N | 4.00E+00 | - | est | |
| $Vdif_{glut3}^{Glc_{BN}}$ | Vdif_glut3_Glc_B_N | 5.00E-01 | L/min | opt | |
| $Kdif_{glut3}^{Glc_{BN}}$ | Kdif_glut3_Glc_B_N | 2.90E+00 | mM | exp | [3] |
| $Kdif_{glut3}^{Glc_N}$ | Kdif_glut3_Glc_N | 2.90E+00 | mM | exp | [3] |
| $Vmax_{hk}^{Glc_N}$ | Vmax_hk_Glc_N | 3.20E+00 | mmol/min | opt | |
| $Km_{hk}^{Glc_N}$ | Km_hk_Glc_N | 7.50E+00 | mM | exp | [4] |
| $Ki_{hk}^{G6p_N}$ | Ki_hk_G6p_N | 5.00E-01 | mM | exp | [4] |
| $Km_{hk}^{Atp_N}$ | Km_hk_Atp_N | 2.09E+00 | mM | exp | [4] |
| $Ki_{hk}^{Atp_N}$ | Ki_hk_Atp_N | 1.90E-01 | mM | exp | [4] |

| | | | | | |
|---|---|---|---|---|---|
| $Vmax^{G6p_N}_{g6pase}$ | Vmax_ppp_G6p_N | 0.00E+00 | mmol/min | est | |
| $Km^{G6p_N}_{ppp}$ | Km_ppp_G6p_N | 1.00E-01 | mM | est | |
| $Km^{Nadp_N}_{ppp}$ | Km_ppp_Nadp_N | 1.00E-01 | mM | est | |
| $Vmax^{G6p_N}_{g6pase}$ | Vmax_g6pase_G6p_N | 0.00E+00 | mmol/min | opt | |
| $Km^{G6p_N}_{g6pase}$ | Km_g6pase_G6p_N | 2.41E+00 | mM | exp | [5, 6] |
| $Vmax^{G6p_N}_{gs}$ | Vmax_gs_G6p_N | 1.60E-01 | mmol/min | opt | |
| $Km^{G6p_N}_{gs}$ | Km_gs_G6p_N | 5.00E-02 | mM | est | [7] |
| $n^{G6pN}_{gs}$ | n_gs_G6p_N | 1.00E+00 | - | est | |
| $Km^{Utp_N}_{gs}$ | Km_gs_Utp_N | 4.80E-02 | mM | exp | [8] |
| $Glygn^{max}_{N}$ | Glygn_N_max | 4.60E+01 | mM | est | [9] |
| $Km^{Glygn_N}_{gs}$ | Km_gs_Glygn_N | 9.20E+00 | mM | est | |
| $Vmax^{Glygn_N}_{gd}$ | Vmax_gd_Glygn_N | 3.00E-02 | mmol/min | opt | |
| $Km^{Glygn_N}_{gd}$ | Km_gd_Glygn_N | 1.00E+01 | mM | est | [7] |
| $Km^{Phos_N}_{gd}$ | Km_gd_Phos_N | 4.00E+00 | mM | exp | [10] |
| $Vmax^{G6p_N}_{pfk}$ | Vmax_pfk_G6p_N | 1.00E+01 | mmol/min | opt | |
| $Km^{G6p_N}_{pfk}$ | Km_pfk_G6p_N | 5.00E-03 | mM | est | [7] |
| $Km^{Atp_N}_{pfk}$ | Km_pfk_Atp_N | 4.25E-02 | mM | exp | [11] |
| $Ki^{Atp_N}_{pfk}$ | Ki_pfk_Atp_N | 2.10E+00 | mM | exp | [12] |
| $Km^{Adp_N}_{pfk}$ | Km_pfk_Adp_N | 8.36E-02 | mM | exp | [11, 12] |
| $Ki^{Gap_N}_{pfk}$ | Ki_pfk_Gap_N | 2.07E-02 | mM | exp | [13] |
| $b^{Gap_N}_{pfk}$ | b_pfk_Gap_N | 7.50E-01 | - | est | [7] |
| $Vmax^{Gap_N}_{fbp}$ | Vmax_fbp_Gap_N | 0.00E+00 | mmol/min | opt | |
| $Km^{Gap_N}_{fbp}$ | Km_fbp_Gap_N | 2.50E-01 | mM | est | [7] |
| $Vmax^{Gap_N}_{pk}$ | Vmax_pk_Gap_N | 6.58E+00 | mmol/min | | |
| $Km^{Gap_N}_{pk}$ | Km_pk_Gap_N | 2.40E-01 | mM | est | [7] |
| $b^{Accoa_{NM}}_{pk}$ | b_pk_Accoa_NM | 8.00E-01 | - | exp | [14] |
| $Ki^{Accoa_{NM}}_{pk}$ | Ki_pk_Accoa_NM | 3.00E-02 | mM | exp | [14] |

| | | | | | |
|---|---|---|---|---|---|
| $Km_{pk}^{Adp_N}$ | Km_pk_Adp_N | 2.40E-01 | mM | exp | [15] |
| $Vmax_{pepck}^{Pyr_N}$ | Vmax_pepck_Pyr_N | 0.00E+00 | mM | opt | |
| $Km_{pepck}^{Pyr_H}$ | Km_pepck_Pyr_N | 1.14E+00 | mM | est | |
| $Km_{pepck}^{Atp_H}$ | Km_pepck_Atp_N | 1.00E-02 | mM | est | [7] |
| $Km_{pepck}^{Gtp_N}$ | Km_pepck_Gtp_N | 6.40E-02 | mM | exp | [16] |
| $Vdif_{pyrt}^{Pyr_{BN}}$ | Vdif_pyrt_Pyr_B_N | 0.00E+00 | L/min | est | |
| $Kdif_{pyrt}^{Pyr_{BN}}$ | Kdif_pyrt_Pyr_B_N | 1.00E+00 | mM | est | |
| $Kdif_{pyrt}^{Pyr_N}$ | Kdif_pyrt_Pyr_N | 1.00E+00 | mM | est | |
| $Vdif_{lact}^{Lac_{BN}}$ | Vdif_lact_Lac_B_N | 0.00E+00 | L/min | opt | |
| $Kdif_{lact}^{Lac_N}$ | Kdif_lact_Lac_N | 2.42E+00 | mM | exp | [17] |
| $Kdif_{lact}^{Lac_{BN}}$ | Kdif_lact_Lac_B_N | 2.42E+00 | mM | exp | [17] |
| $Vmax_{ldh}^{Pyr_N}$ | Vmax_ldh_Pyr_N | 0.00E+00 | $L^2$/mmol/min | opt | |
| $Km_{ldh}^{Pyr_N}$ | Km_ldh_Pyr_N | 1.50E-01 | mM | exp | [18] |
| $Km_{ldh}^{Nadh_N}$ | Km_ldh_Nadh_N | 1.50E-02 | mM | exp | [18] |
| $Km_{ldh}^{Lac_N}$ | Km_ldh_Lac_N | 3.60E+01 | mM | exp | [19] |
| $Km_{ldh}^{Nad_N}$ | Km_ldh_Nad_N | 1.10E-01 | mM | exp | [18] |
| $Keq_{ldh}^{Lac_N}$ | Keq_ldh_Lac_N | 5.00E-01 | - | est | |
| $Vdif_{alat}^{Ala_{BN}}$ | Vdif_alat_Ala_B_N | 0.00E+00 | L/min | est | |
| $Kdif_{alat}^{Ala_N}$ | Kdif_alat_Ala_N | 2.42E+00 | mM | est | |
| $Kdif_{alat}^{Ala_{BN}}$ | Kdif_alat_Ala_B_N | 2.42E+00 | mM | est | |
| $Vmax_{alata}^{Pyr_N}$ | Vmax_alata_Pyr_N | 0.00E+00 | L/min | opt | |
| $Km_{alata}^{Pyr_N}$ | Km_alata_Pyr_N | 1.50E-01 | mM | est | |
| $Km_{alata}^{Ala_N}$ | Km_alata_Ala_N | 3.60E+01 | mM | est | |
| $Keq_{alata}^{Ala_N}$ | Keq_alata_Ala_N | 1.00E+00 | - | est | |
| $Vmax_{pdh}^{Pyr_N}$ | Vmax_pdh_Pyr_N | 2.00E+01 | mmol/min | opt | |
| $Km_{pdh}^{Pyr_N}$ | Km_pdh_Pyr_N | 5.40E-01 | mM | est | [7] |
| $Km_{pdh}^{Accoa_{NM}}$ | Ki_pdh_Accoa_NM | 3.50E-02 | mM | exp | [20] |
| $Km_{pdh}^{Nad_N}$ | Km_pdh_Nad_N | 1.00E-02 | mM | opt | |

| | | | | | |
|---|---|---|---|---|---|
| $Vmax_{tca}^{Accoa_{NM}}$ | Vmax_tca_Accoa_NM | 6.00E+01 | mmol/min | opt | |
| $Km_{tca}^{Accoa_{NM}}$ | Km_tca_Accoa_NM | 4.00E-04 | mM | exp | [21] |
| $Km_{tca}^{Phos_{N}}$ | Km_tca_Phos_N | 1.00E+00 | mM | opt | |
| $Km_{tca}^{Adp_{N}}$ | Km_tca_Adp_N | 4.10E-01 | mM | exp | [22] |
| $Km_{tca}^{Pyr_{N}}$ | Km_tca_Pyr_N | 0.00E+00 | mM | opt | |
| $Km_{tca}^{Nad_{N}}$ | Km_tca_Nad_N | 1.00E-02 | mM | est | |
| $Km_{tca}^{Fad_{N}}$ | Km_tca_Fad_N | 1.00E-02 | mM | est | |
| $Vdif_{ffat}^{FFA_{BN}}$ | Vdif_ffat_FFA_B_N | 0.00E+00 | L/min | opt | |
| $Kdif_{ffat}^{FFA_{BN}}$ | Kdif_ffat_FFA_B_N | 2.00E-01 | mM | est | [7] |
| $Kdif_{ffat}^{FFA_{N}}$ | Kdif_ffat_FFA_N | 2.00E-01 | mM | est | [7] |
| $Vmax_{ffat}^{FFA_{BN}}$ | Vmax_ffat_FFA_B_N | 0.00E+00 | mmol/min | opt | |
| $Km_{ffat}^{FFA_{BN}}$ | Km_ffat_FFA_B_N | 2.00E-03 | mM | est | [7] |
| $Vmax_{tgsyn}^{FFA_{N}}$ | Vmax_tgsyn_FFA_N | 0.00E+00 | mmol/min | opt | |
| $Km_{tgsyn}^{FFA_{N}}$ | Km_tgsyn_FFA_N | 6.45E-01 | mM | est | [7] |
| $Km_{tgsyn}^{Glycp_{N}}$ | Km_tgsyn_Glycp_N | 4.60E-01 | mM | exp | [23] |
| $Km_{tgsyn}^{Atp_{N}}$ | Km_tgsyn_Atp_N | 1.00E-01 | mM | opt | |
| $TG_{N}^{max}$ | TG_N_max | 9.99E+02 | mM | est | |
| $Km_{tgsyn}^{TG_{N}}$ | Km_tgsyn_TG_N | 9.99E+02 | mM | est | |
| $Vmax_{tgdeg}^{TG_{N}}$ | Vmax_tgdeg_TG_N | 0.00E+00 | mmol/min | opt | |
| $Km_{tgdeg}^{TG_{N}}$ | Km_tgdeg_TG_N | 5.07E+01 | mM | est | [7] |
| $Vdif_{glyct}^{Glyc_{BN}}$ | Vdif_glyct_Glyc_B_N | 4.00E+00 | L/min | opt | |
| $Kdif_{glyct}^{Glyc_{BN}}$ | Kdif_glyct_Glyc_B_N | 2.70E-01 | mM | exp | [24] |
| $Kdif_{glyct}^{Glyc_{N}}$ | Kdif_glyct_Glyc_N | 2.70E-01 | mM | exp | [24] |
| $Vmax_{glyk}^{Glyc_{N}}$ | Vmax_glyk_Glyc_N | 0.00E+00 | mmol/min | opt | |
| $Km_{glyk}^{Glyc_{N}}$ | Km_glyk_Glyc_N | 4.00E-02 | mM | exp | [24] |
| $Km_{glyk}^{Atp_{N}}$ | Km_glyk_Atp_N | 5.80E-02 | mM | exp | [25] |
| $Ki_{glyk}^{Gap_{N}}$ | Ki_glyk_Gap_N | 5.80E-01 | mM | exp | [25] |
| $Vdif_{tgt}^{TG_{BN}}$ | Vdif_tgt_TG_B_N | 0.00E+00 | mmol/min | est | |

| | | | | | |
|---|---|---|---|---|---|
| $Kdif_{tgt}^{TG_{BN}}$ | Kdif_tgt_TG_B_N | 1.00E+00 | mM | est | [7] |
| $Keq_{tgt}^{TG_N}$ | Keq_tgt_TG_N | 3.38E+01 | - | est | [7] |
| $Vmax_{tgt}^{TG_{BN}}$ | Vmax_tgt_TG_B_N | 0.00E+00 | mmol/min | opt | |
| $Km_{tgt}^{TG_N}$ | Km_tgt_TG_N | 3.38E+01 | mM | est | [7] |
| $Vmax_{g3pd}^{Gap_N}$ | Vmax_g3pd_Gap_N | 0.00E+00 | $L^2$/mmol/min | opt | |
| $Km_{g3pd}^{Gap_N}$ | Km_g3pd_Gap_N | 1.60E-01 | mM | exp | [26] |
| $Km_{g3pd}^{Glycp_N}$ | Km_g3pd_Glycp_N | 2.20E-01 | mM | exp | [26] |
| $Km_{g3pd}^{Nadh_N}$ | Km_g3pd_Nadh_N | 8.00E-03 | mM | exp | [26] |
| $Km_{g3pd}^{Nad_H}$ | Km_g3pd_Nad_N | 1.30E-02 | mM | exp | [26] |
| $Keq_{g3pd}^{Glycp_N}$ | Keq_g3pd_Glycp_N | 1.00E+03 | - | opt | |
| $Vmax_{boxid}^{FFA_N}$ | Vmax_boxid_FFA_N | 0.00E+00 | mmol/min | opt | |
| $Km_{boxid}^{FFA_N}$ | Km_boxid_FFA_N | 5.00E-03 | mM | exp | [27-29] |
| $Km_{boxid}^{Atp_N}$ | Km_boxid_Atp_N | 8.70E-02 | mM | exp | [30] |
| $Ki_{boxid}^{Accoa_{NM}}$ | Ki_boxid_Accoa_NM | 1.20E-01 | mM | | [31] |
| $Ki_{boxid}^{Malcoa_N}$ | Ki_boxid_Malcoa_N | 1.00E+00 | mM | opt | |
| $Km_{boxid}^{Nad_N}$ | Km_boxid_Nad_N | 1.00E-01 | mM | est | |
| $Km_{boxid}^{Fad_N}$ | Km_boxid_Fad_N | 1.00E-01 | mM | est | |
| $Vmax_{accoat}^{Accoa_{NMNC}}$ | Vmax_accoat_Accoa_NM_NC | 1.00E+00 | mmol/min | est | |
| $Km_{accoat}^{Accoa_{NM}}$ | Km_accoat_Accoa_NM | 5.80E-02 | mM | opt | |
| $Km_{accoat}^{Atp_N}$ | Km_accoat_Atp_N | 1.00E-01 | mM | est | |
| $Km_{accoat}^{Pyr_N}$ | Km_accoat_Pyr_N | 1.00E+00 | mM | est | |
| $Vmax_{bhbsyn}^{Accoa_{NM}}$ | Vmax_bhbsyn_Accoa_NM | 0.00E+00 | mmol/min | opt | |
| $Km_{bhbsyn}^{Accoa_{NM}}$ | Km_bhbsyn_Accoa_NM | 1.00E-01 | mM | est | |
| $Km_{bhbsyn}^{Nadh_N}$ | Km_bhbsyn_Nadh_N | 1.00E-01 | mM | est | |
| $n_{bhbsyn}^{Pyr_N}$ | n_bhbsyn_Pyr_N | 4.00E+00 | - | est | |
| $Ki_{bhbsyn}^{Pyr_N}$ | Ki_bhbsyn_Pyr_N | 1.00E-02 | mM | opt | |
| $Vmax_{bhbdeg}^{Bhb_N}$ | Vmax_bhbdeg_Bhb_N | 2.00E+02 | mmol/min | est | |
| $Km_{bhbdeg}^{Bhb_N}$ | Km_bhbdeg_Bhb_N | 1.00E-01 | mM | est | |

| | | | | |
|---|---|---|---|---|
| $Km_{bhbdeg}^{Atp_N}$ | Km_bhbdeg_Atp_N | 1.00E-01 | mM | est |
| $Km_{bhbdeg}^{Nad_N}$ | Km_bhbdeg_Nad_N | 1.00E-01 | mM | est |
| $Vdif_{bhbt}^{Bhb_{BN}}$ | Vdif_bhbt_Bhb_B_N | 1.00E+00 | L/min | est |
| $Kdif_{bhbt}^{Bhb_{BN}}$ | Kdif_bhbt_Bhb_B_N | 2.00E+00 | mM | est |
| $Kdif_{bhbt}^{Bhb_N}$ | Kdif_bhbt_Bhb_N | 2.00E+00 | mM | est |
| $Vmax_{lipog1}^{Accoa_{NC}}$ | Vmax_lipog1_Accoa_NC | 0.00E+00 | mmol/min | opt |
| $Km_{lipog1}^{Accoa_{NC}}$ | Km_lipog1_Accoa_NC | 1.00E-02 | mM | est |
| $Km_{lipog1}^{Atp_N}$ | Km_lipog1_Atp_N | 1.00E-01 | mM | est |
| $Vmax_{lipog2}^{Malcoa_N}$ | Vmax_lipog2_Malcoa_N | 0.00E+00 | mmol/min | est |
| $Km_{lipog2}^{Malcoa_N}$ | Km_lipog2_Malcoa_N | 1.00E-02 | mM | est |
| $Km_{lipog2}^{Adp_N}$ | Km_lipog2_Adp_N | 1.00E-01 | mM | est |
| $Km_{lipog2}^{Nadph_N}$ | Km_lipog2_Nadph_N | 1.00E-02 | mM | est |
| $Vmax_{cholsyn1}^{Accoa_{NC}}$ | Vmax_cholsyn1_Accoa_NC | 0.00E+00 | mmol/min | est |
| $Km_{cholsyn1}^{Accoa_{NC}}$ | Km_cholsyn1_Accoa_NC | 1.00E-01 | mM | est |
| $Vmax_{cholsyn2}^{Hmgcoa_N}$ | Vmax_cholsyn2_Hmgcoa_N | 0.00E+00 | mmol/min | est |
| $Km_{cholsyn2}^{Hmgcoa_N}$ | Km_cholsyn2_Hmgcoa_N | 1.00E-01 | mM | est |
| $Km_{cholsyn2}^{Atp_N}$ | Km_cholsyn2_Atp_N | 1.00E-01 | mM | est |
| $Km_{cholsyn2}^{Fadh_N}$ | Km_cholsyn2_Fadh_N | 1.00E-01 | mM | est |
| $Km_{cholsyn2}^{Nadph_N}$ | Km_cholsyn2_Nadph_N | 1.00E-01 | mM | est |
| $Vmax_{cholt}^{Chol_{BN}}$ | Vmax_cholt_Chol_N | 0.00E+00 | mmol/min | est |
| $Km_{cholt}^{Chol_N}$ | Km_cholt_Chol_N | 1.00E-01 | mM | est |
| $Vmax_{atpsynf}^{Fadh_N}$ | Vmax_atpsynf_Fadh_N | 1.00E+00 | mmol/min | opt |
| $Km_{atpsynf}^{Fadh_N}$ | Km_atpsynf_Fadh_N | 1.00E-01 | mM | est |
| $Km_{atpsynf}^{Adp_N}$ | Km_atpsynf_Adp_N | 1.00E-01 | mM | est |
| $Vmax_{atpsynn}^{Nadh_N}$ | Vmax_atpsynn_Nadh_N | 9.00E+00 | mmol/min | est |
| $Km_{atpsynn}^{Nadh_N}$ | Km_atpsynn_Nadh_N | 1.00E-01 | mM | est |
| $Km_{atpsynn}^{Adp_N}$ | Km_atpsynn_Adp_N | 1.00E-01 | mM | est |

| | | | | | |
|---|---|---|---|---|---|
| $Vmax_{atpuse}^{Atp_N}$ | Vmax_atpuse_Atp_N | 3.35E+01 | mmol/min | opt | |
| $Km_{atpuse}^{Atp_N}$ | Km_atpuse_Atp_N | 4.52E+00 | mM | opt | |
| $Vmax_{ampreg}^{Amp_N}$ | Vmax_ampreg_Amp_N | 0.00E+00 | mmol/min | est | |
| $Km_{ampreg}^{Amp_N}$ | Km_ampreg_Amp_N | 8.00E-02 | mM | exp | [32] |
| $Km_{ampreg}^{Atp_N}$ | Km_ampreg_Atp_N | 9.00E-02 | mM | exp | [32] |
| $Km_{ampreg}^{Adp_N}$ | Km_ampreg_Adp_N | 1.10E-01 | mM | exp | [32] |
| $Vmax_{nadhk}^{Nadh_N}$ | Vmax_nadhk_Nadh_N | 0.00E+00 | mmol/min | est | |
| $Km_{nadhk}^{Nadh_N}$ | Km_nadhk_Nadh_N | 1.00E-01 | mM | est | |
| $Km_{nadhk}^{Atp_N}$ | Km_nadhk_Atp_N | 1.00E-01 | mM | est | |
| $Vmax_{nadhuse}^{Nadh_N}$ | Vmax_nadhuse_Nadh_N | 2.00E+00 | mmol/min | est | |
| $Km_{nadhuse}^{Nadh_N}$ | Km_nadhuse_Nadh_N | 7.12E-02 | mM | est | |
| $Vmax_{gdpreg}^{Gdp_N}$ | Vmax_gdpreg_Gdp_N | 0.00E+00 | $L^2$/mmol/min | est | |
| $Km_{gdpreg}^{Gdp_N}$ | Km_gdpreg_Gdp_N | 3.10E-02 | mM | exp | [33] |
| $Km_{gdpreg}^{Atp_N}$ | Km_gdpreg_Atp_N | 1.33E+00 | mM | exp | [33] |
| $Km_{gdpreg}^{Gtp_N}$ | Km_gdpreg_Gtp_N | 1.50E-01 | mM | exp | [34] |
| $Km_{gdpreg}^{Adp_N}$ | Km_gdpreg_Adp_N | 4.20E-02 | mM | exp | [33] |
| $Keq_{gdpreg}^{Gtp_N}$ | Keq_gdpreg_Gtp_N | 1.00E+03 | - | opt | |
| $Vmax_{gtpuse}^{Gtp_N}$ | Vmax_gtpuse_Gtp_N | 0.00E+00 | mmol/min | est | |
| $Km_{gtpuse}^{Gtp_N}$ | Km_gtpuse_Gtp_N | 1.00E-01 | mM | est | |
| $Vmax_{udpreg}^{Udp_N}$ | Vmax_udpreg_Udp_N | 1.57E+01 | $L^2$/mmol/min | est | |
| $Km_{udpreg}^{Udp_N}$ | Km_udpreg_Udp_N | 1.90E-01 | mM | exp | [33] |
| $Km_{udpreg}^{Atp_N}$ | Km_udpreg_Atp_N | 1.33E+00 | mM | exp | [33] |
| $Km_{udpreg}^{Utp_N}$ | Km_udpreg_Utp_N | 1.60E+01 | mM | exp | [34] |
| $Km_{udpreg}^{Adp_N}$ | Km_udpreg_Adp_N | 4.20E-02 | mM | exp | [33] |
| $Keq_{udpreg}^{Utp_N}$ | Keq_udpreg_Utp_N | 1.00E+03 | - | est | |
| $Vmax_{utpuse}^{Utp_N}$ | Vmax_utpuse_Utp_N | 5.22E-01 | mmol/min | est | |
| $Km_{utpuse}^{Utp_N}$ | Km_utpuse_Utp_N | 1.86E-01 | mM | est | |
| $Vmax_{nadphuse}^{Nadph_N}$ | Vmax_nadphuse_Nadph_N | 0.00E+00 | mmol/min | est | |

| | | | | |
|---|---|---|---|---|
| $Km_{nadphuse}^{Nadph_N}$ | Km_nadphuse_Nadph_N | 1.00E-01 | mM | est |
| $Vmax_{fadhuse}^{Fadh_N}$ | Vmax_fadhuse_Fadh_N | 1.00E-01 | mmol/min | est |
| $Km_{fadhuse}^{Fadh_N}$ | Km_fadhuse_Fadh_N | 1.00E-01 | mM | est |
| $Vmax_{ck}^{Cre_N}$ | Vmax_ck_Cre_N | 6.75E+00 | $L^2$/mmol/min | est |
| $Km_{ck}^{Cre_N}$ | Km_ck_Cre_N | 4.91E+00 | mM | est |
| $Km_{ck}^{Atp_N}$ | Km_ck_Atp_N | 6.68E-01 | mM | est |
| $Km_{ck}^{Crep_N}$ | Km_ck_Crep_N | 7.33E+00 | mM | est |
| $Km_{ck}^{Adp_N}$ | Km_ck_Adp_N | 6.56E-01 | mM | est |
| $Keq_{ck}^{Crep_N}$ | Keq_ck_Crep_N | 7.33E+00 | - | est |
| $Phos_N$ | Phos_N | 5.00E+00 | mM | est |

Method: exp (experiment), est (estimation), opt(optimization)
