## Supplemental Table 2 for "Virtual metabolic human dynamic model for pathological analysis and therapy design for diabetes"

**Table S2 Metabolite list**

| Symbol | Symbol in program | Name | Concentration (mM) |
|---|---|---|---|
| $Ins_B$ | Ins_B | insulin in blood | 0.00000005 |
| $Glc_B$ | Glc_B | glucose in blood | 5 |
| $Pyr_B$ | Pyr_B | pyruvate in blood | 0.68 |
| $Lac_B$ | Lac_B | lactate in blood | 0.7 |
| $Ala_B$ | Ala_B | alanine in blood | 0.192 |
| $Glyc_B$ | Glyc_B | glycerol in blood | 0.07 |
| $Bhb_B$ | Bhb_B | β-hydroxybutyrate in blood | 0.01 |
| $FFA_B$ | FFA_B | free fatty acid in blood | 0.66 |
| $TG_B$ | TG_B | triglyceride in blood | 0.8 |
| $Chol_B$ | Chol_B | cholesterol in blood | 5 |
| $Glc_L$ | Glc_L | glucose in liver | 5 |
| $G6p_L$ | G6p_L | glucose-6-phosphate in liver | 1 |
| $Glygn_L$ | Glygn_L | glycogen in liver | 200 |
| $Gap_L$ | Gap_L | glyceraldehyde 3-phosphate in liver | 1 |
| $Pyr_L$ | Pyr_L | pyruvate in liver | 1 |
| $Lac_L$ | Lac_L | lactate in liver | 1 |
| $Ala_L$ | Ala_L | alanine in liver | 1 |
| $Accoa_{LM}$ | Accoa_LM | acetyl-CoA in liver | 1 |
| $Bhb_L$ | Bhb_L | β-hydroxybutyrate in liver | 0 |
| $Accoa_{LC}$ | Accoa_LC | acetyl-CoA in liver | 0 |
| $Malcoa_L$ | Malcoa_L | malonyl-CoA in liver | 0 |
| $Hmgcoa_L$ | Hmgcoa_L | hydroxymethylglutaryl-CoA in liver | 0 |
| $Chol_L$ | Chol_L | cholesterol in liver | 0 |
| $FFA_L$ | FFA_L | free fatty acid in liver | 1 |
| $Glyc_L$ | Glyc_L | glycerol in liver | 1 |
| $Glycp_L$ | Glycp_L | glycerol 3-phosphate in liver | 1 |
| $TG_L$ | TG_L | triglyceride in liver | 2.93 |
| $Atp_L$ | Atp_L | ATP in liver | 2.74 |
| $Adp_L$ | Adp_L | ADP in liver | 1.22 |
| $Amp_L$ | Amp_L | AMP in liver | 0 |

| | | | |
|---|---|---|---|
| $Gtp_L$ | Gtp_L | GTP in liver | 0.29 |
| $Gdp_L$ | Gdp_L | GDP in liver | 0.01 |
| $Utp_L$ | Utp_L | UTP in liver | 0.27 |
| $Udp_L$ | Udp_L | UDP in liver | 0 |
| $Nadh_L$ | Nadh_L | NADH in liver | 0.45 |
| $Nad_L$ | Nad_L | $NAD^+$ in liver | 0.05 |
| $Fadh_L$ | Fadh_L | $FADH_2$ in liver | 0.45 |
| $Fad_L$ | Fad_L | FAD in liver | 0.05 |
| $Nadph_L$ | Nadph_L | NADPH in liver | 0.25 |
| $Nadp_L$ | Nadp_L | $NADP^+$ in liver | 0.25 |
| $Cre_L$ | Cre_L | creatine in liver | 0 |
| $Crep_L$ | Crep_L | creatine phosphate in liver | 0 |
| $Glc_M$ | Glc_M | glucose in skeletal muscle | 4.2 |
| $G6p_M$ | G6p_M | glucose-6-phosphate in skeletal muscle | 8.2 |
| $Glygn_M$ | Glygn_M | glycogen in skeletal muscle | 10 |
| $Gap_M$ | Gap_M | glyceraldehyde 3-phosphate in skeletal muscle | 0.72 |
| $Pyr_M$ | Pyr_M | pyruvate in skeletal muscle | 0.28 |
| $Lac_M$ | Lac_M | lactate in skeletal muscle | 0.72 |
| $Ala_M$ | Ala_M | alanine in skeletal muscle | 0.19 |
| $Accoa_{MM}$ | Accoa_MM | acetyl-CoA in skeletal muscle | 0.0022 |
| $Bhb_M$ | Bhb_M | β-hydroxybutyrate in skeletal muscle | 0 |
| $Accoa_{MC}$ | Accoa_MC | acetyl-CoA in skeletal muscle | 0 |
| $Malcoa_M$ | Malcoa_M | malonyl-CoA in skeletal muscle | 0 |
| $Hmgcoa_M$ | Hmgcoa_M | hydroxymethylglutaryl-CoA in skeletal muscle | 0 |
| $Chol_M$ | Chol_M | cholesterol in skeletal muscle | 0 |
| $FFA_M$ | FFA_M | free fatty acid in skeletal muscle | 0.53 |
| $Glyc_M$ | Glyc_M | glycerol in skeletal muscle | 1 |
| $Glycp_M$ | Glycp_M | glycerol 3-phosphate in skeletal muscle | 1 |
| $TG_M$ | TG_M | triglyceride in skeletal muscle | 4 |
| $Atp_M$ | Atp_M | ATP in skeletal muscle | 6.15 |
| $Adp_M$ | Adp_M | ADP in skeletal muscle | 0.02 |

| | | | |
|---|---|---|---|
| $Amp_M$ | Amp_M | AMP in skeletal muscle | 0 |
| $Gtp_M$ | Gtp_M | GTP in skeletal muscle | 0.29 |
| $Gdp_M$ | Gdp_M | GDP in skeletal muscle | 0 |
| $Utp_M$ | Utp_M | UTP in skeletal muscle | 0.27 |
| $Udp_M$ | Udp_M | UDP in skeletal muscle | 0 |
| $Nadh_M$ | Nadh_M | NADH in skeletal muscle | 0.05 |
| $Nad_M$ | Nad_M | $NAD^+$ in skeletal muscle | 0.45 |
| $Fadh_M$ | Fadh_M | $FADH_2$ in skeletal muscle | 0.3 |
| $Fad_M$ | Fad_M | FAD in skeletal muscle | 0.2 |
| $Nadph_M$ | Nadph_M | NADPH in skeletal muscle | 0.3 |
| $Nadp_M$ | Nadp_M | $NADP^+$ in skeletal muscle | 0.2 |
| $Cre_M$ | Cre_M | creatine in skeletal muscle | 10.45 |
| $Crep_M$ | Crep_M | creatine phosphate in skeletal muscle | 20.1 |
| $Glc_A$ | Glc_A | glucose in adipose tissue | 0.3 |
| $G6p_A$ | G6p_A | glucose-6-phosphate in adipose tissue | 0 |
| $Glygn_A$ | Glygn_A | glycogen in adipose tissue | 10 |
| $Gap_A$ | Gap_A | glyceraldehyde 3-phosphate in adipose tissue | 0.11 |
| $Pyr_A$ | Pyr_A | pyruvate in adipose tissue | 0.37 |
| $Lac_A$ | Lac_A | lactate in adipose tissue | 0.82 |
| $Ala_A$ | Ala_A | alanine in adipose tissue | 0.01 |
| $Accoa_{AM}$ | Accoa_AM | acetyl-CoA in adipose tissue | 0.035 |
| $Bhb_A$ | Bhb_A | β-hydroxybutyrate in adipose tissue | 0 |
| $Accoa_{AC}$ | Accoa_AC | acetyl-CoA in adipose tissue | 0 |
| $Malcoa_A$ | Malcoa_A | malonyl-CoA in adipose tissue | 0 |
| $Hmgcoa_A$ | Hmgcoa_A | hydroxymethylglutaryl-CoA in adipose tissue | 0 |
| $Chol_A$ | Chol_A | cholesterol in adipose tissue | 0 |
| $FFA_A$ | FFA_A | free fatty acid in adipose tissue | 0.57 |
| $Glyc_A$ | Glyc_A | glycerol in adipose tissue | 0.22 |
| $Glycp_A$ | Glycp_A | glycerol 3-phosphate in adipose tissue | 0.24 |
| $TG_A$ | TG_A | triglyceride in adipose tissue | 500 |
| $Atp_A$ | Atp_A | ATP in adipose tissue | 2.74 |

| | | | |
|---|---|---|---|
| $Adp_A$ | Adp_A | ADP in adipose tissue | 1.22 |
| $Amp_A$ | Amp_A | AMP in adipose tissue | 0.16 |
| $Gtp_A$ | Gtp_A | GTP in adipose tissue | 0 |
| $Gdp_A$ | Gdp_A | GDP in adipose tissue | 0 |
| $Utp_A$ | Utp_A | UTP in adipose tissue | 0 |
| $Udp_A$ | Udp_A | UDP in adipose tissue | 0 |
| $Nadh_A$ | Nadh_A | NADH in adipose tissue | 0.05 |
| $Nad_A$ | Nad_A | $NAD^+$ in adipose tissue | 0.45 |
| $Fadh_A$ | Fadh_A | $FADH_2$ in adipose tissue | 0.45 |
| $Fad_A$ | Fad_A | FAD in adipose tissue | 0.05 |
| $Nadph_A$ | Nadph_A | NADPH in adipose tissue | 0.45 |
| $Nadp_A$ | Nadp_A | $NADP^+$ in adipose tissue | 0.05 |
| $Cre_A$ | Cre_A | creatine in adipose tissue | 0 |
| $Crep_A$ | Crep_A | creatine phosphate in adipose tissue | 0 |
| $Glc_G$ | Glc_G | glucose in GI | 1 |
| $G6p_G$ | G6p_G | glucose-6-phosphate in GI | 1 |
| $Glygn_G$ | Glygn_G | glycogen in GI | 10 |
| $Gap_G$ | Gap_G | glyceraldehyde 3-phosphate in GI | 1 |
| $Pyr_G$ | Pyr_G | pyruvate in GI | 1 |
| $Lac_G$ | Lac_G | lactate in GI | 1 |
| $Ala_G$ | Ala_G | alanine in GI | 1 |
| $Accoa_{GM}$ | Accoa_GM | acetyl-CoA in GI | 1 |
| $Bhb_G$ | Bhb_G | β-hydroxybutyrate in GI | 0 |
| $Accoa_{GC}$ | Accoa_GC | acetyl-CoA in GI | 0 |
| $Malcoa_G$ | Malcoa_G | malonyl-CoA in GI | 0 |
| $Hmgcoa_G$ | Hmgcoa_G | hydroxymethylglutaryl-CoA in GI | 0 |
| $Chol_G$ | Chol_G | cholesterol in GI | 0 |
| $FFA_G$ | FFA_G | free fatty acid in GI | 1 |
| $Glyc_G$ | Glyc_G | glycerol in GI | 1 |
| $Glycp_G$ | Glycp_G | glycerol 3-phosphate in GI | 1 |
| $TG_G$ | TG_G | triglyceride in GI | 200 |
| $Atp_G$ | Atp_G | ATP in GI | 3 |

| | | | |
|---|---|---|---|
| $Adp_G$ | Adp_G | ADP in GI | 0.8 |
| $Amp_G$ | Amp_G | AMP in GI | 0 |
| $Gtp_G$ | Gtp_G | GTP in GI | 0.29 |
| $Gdp_G$ | Gdp_G | GDP in GI | 0 |
| $Utp_G$ | Utp_G | UTP in GI | 0.27 |
| $Udp_G$ | Udp_G | UDP in GI | 0 |
| $Nadh_G$ | Nadh_G | NADH in GI | 0.5 |
| $Nad_G$ | Nad_G | $NAD^+$ in GI | 0.05 |
| $Fadh_G$ | Fadh_G | $FADH_2$ in GI | 0.25 |
| $Fad_G$ | Fad_G | FAD in GI | 0.25 |
| $Nadph_G$ | Nadph_G | NADPH in GI | 0.5 |
| $Nadp_G$ | Nadp_G | $NADP^+$ in GI | 0 |
| $Cre_G$ | Cre_G | creatine in GI | 3.5 |
| $Crep_G$ | Crep_G | creatine phosphate in GI | 8.3 |
| $Glc_H$ | Glc_H | glucose in heart | 1 |
| $G6p_H$ | G6p_H | glucose-6-phosphate in heart | 0.17 |
| $Glygn_H$ | Glygn_H | glycogen in heart | 0 |
| $Gap_H$ | Gap_H | glyceraldehyde 3-phosphate in heart | 0.01 |
| $Pyr_H$ | Pyr_H | pyruvate in heart | 0.2 |
| $Lac_H$ | Lac_H | lactate in heart | 3.88 |
| $Ala_H$ | Ala_H | alanine in heart | 0 |
| $Accoa_{HM}$ | Accoa_HM | acetyl-CoA in heart | 0.0012 |
| $Bhb_H$ | Bhb_H | β-hydroxybutyrate in heart | 0 |
| $Accoa_{HC}$ | Accoa_HC | acetyl-CoA in heart | 0 |
| $Malcoa_H$ | Malcoa_H | malonyl-CoA in heart | 0 |
| $Hmgcoa_H$ | Hmgcoa_H | hydroxymethylglutaryl-CoA in heart | 0 |
| $Chol_H$ | Chol_H | cholesterol in heart | 0 |
| $FFA_H$ | FFA_H | free fatty acid in heart | 0.021 |
| $Glyc_H$ | Glyc_H | glycerol in heart | 0.015 |
| $Glycp_H$ | Glycp_H | glycerol 3-phosphate in heart | 0.29 |
| $TG_H$ | TG_H | triglyceride in heart | 3.12 |
| $Atp_H$ | Atp_H | ATP in heart | 3.4 |

| | | | |
|---|---|---|---|
| $Adp_H$ | Adp_H | ADP in heart | 0.02 |
| $Amp_H$ | Amp_H | AMP in heart | 0.16 |
| $Gtp_H$ | Gtp_H | GTP in heart | 0.29 |
| $Gdp_H$ | Gdp_H | GDP in heart | 1 |
| $Utp_H$ | Utp_H | UTP in heart | 0.27 |
| $Udp_H$ | Udp_H | UDP in heart | 1 |
| $Nadh_H$ | Nadh_H | NADH in heart | 0.4 |
| $Nad_H$ | Nad_H | $NAD^+$ in heart | 0.045 |
| $Fadh_H$ | Fadh_H | $FADH_2$ in heart | 0.5 |
| $Fad_H$ | Fad_H | FAD in heart | 0 |
| $Nadph_H$ | Nadph_H | NADPH in heart | 0.5 |
| $Nadp_H$ | Nadp_H | $NADP^+$ in heart | 0 |
| $Cre_H$ | Cre_H | creatine in heart | 10.45 |
| $Crep_H$ | Crep_H | creatine phosphate in heart | 20.1 |
| $Glc_N$ | Glc_N | glucose in brain | 1.12 |
| $G6p_N$ | G6p_N | glucose-6-phosphate in brain | 0.16 |
| $Glygn_N$ | Glygn_N | glycogen in brain | 2 |
| $Gap_N$ | Gap_N | glyceraldehyde 3-phosphate in brain | 0.15 |
| $Pyr_N$ | Pyr_N | pyruvate in brain | 0.15 |
| $Lac_N$ | Lac_N | lactate in brain | 0 |
| $Ala_N$ | Ala_N | alanine in brain | 0 |
| $Accoa_{NM}$ | Accoa_NM | acetyl-CoA in brain | 0.068 |
| $Bhb_N$ | Bhb_N | β-hydroxybutyrate in brain | 0 |
| $Accoa_{NC}$ | Accoa_NC | acetyl-CoA in brain | 0 |
| $Malcoa_N$ | Malcoa_N | malonyl-CoA in brain | 0 |
| $Hmgcoa_N$ | Hmgcoa_N | hydroxymethylglutaryl-CoA in brain | 0 |
| $Chol_N$ | Chol_N | cholesterol in brain | 0 |
| $FFA_N$ | FFA_N | free fatty acid in brain | 0 |
| $Glyc_N$ | Glyc_N | glycerol in brain | 0 |
| $Glycp_N$ | Glycp_N | glycerol 3-phosphate in brain | 0 |
| $TG_N$ | TG_N | triglyceride in brain | 0 |
| $Atp_N$ | Atp_N | ATP in brain | 2.54 |

| | | | |
|---|---|---|---|
| $Adp_N$ | Adp_N | ADP in brain | 0.54 |
| $Amp_N$ | Amp_N | AMP in brain | 0 |
| $Gtp_N$ | Gtp_N | GTP in brain | 0 |
| $Gdp_N$ | Gdp_N | GDP in brain | 0 |
| $Utp_N$ | Utp_N | UTP in brain | 0 |
| $Udp_N$ | Udp_N | UDP in brain | 0.8 |
| $Nadh_N$ | Nadh_N | NADH in brain | 0.026 |
| $Nad_N$ | Nad_N | $NAD^+$ in brain | 0.064 |
| $Fadh_N$ | Fadh_N | $FADH_2$ in brain | 0.45 |
| $Fad_N$ | Fad_N | FAD in brain | 0.05 |
| $Nadph_N$ | Nadph_N | NADPH in brain | 0 |
| $Nadp_N$ | Nadp_N | $NADP^+$ in brain | 0 |
| $Cre_N$ | Cre_N | creatine in brain | 5.6 |
| $Crep_N$ | Crep_N | creatine phosphate in brain | 4.6 |

_X indicates the organ name. B, L, M, A, G, H, and N indicate blood, liver, skeletal muscle, adipose tissue, GI tract, heart and brain, respectively.
_XC and _XM indicate the cytoplasm of organ X and mitochondria of organ X, respectively.
