## Supplemental Table 3 for "Virtual metabolic human dynamic model for pathological analysis and therapy design for diabetes"

**Table S3 Abbreviation**

| Variable name in program | Definition or full name |
|---|---|
| tot_Glc_B | total input glucose |
| t_delay_Glc_B | time delay of glucose release |
| tot_TG_B | total input TG |
| t_delay_TG_B | time delay of TG release |
| k_inssyn_B | insulin base rate constant |
| Vmax_inssyn_Glc_B | insulin synthesis rate constant for plasma glucose |
| Km_inssyn_Glc_B | Michaelis constant for insulin synthesis |
| n_inssyn_Glc_B | Hill coefficient for insulin synthesis |
| k_inssyn_FFA_B | insulin synthesis rate constant for plasma FFA |
| k_insdeg_Ins_B | insulin degradation rate constant |
| k_choluse_Chol_B | cholesterol utilization rate constant in blood |
| alpha_base_L | insulin base factor in liver |
| alpha_band_L | insulin allowable width in liver |
| Km_Ins_B_L | Michaelis constant for insulin in liver |
| n_Ins_B_L | Hill coefficient for insulin in liver |
| Vdif_glut2_Glc_B_L | rate constant of glucose transporter type 2 in liver |
| Kdif_glut2_Glc_B_L | dissociation constant of glucose transporter type 2 for Glc_B in liver |
| Kdif_glut2_Glc_L | dissociation constant of glucose transporter type 2 in liver |
| Vmax_hk_Glc_L | reaction rate constant of hexokinase for Glc in liver |
| Km_hk_Glc_L | Michaelis constant of hexokinase for Glc in liver |
| Ki_hk_G6p_L | inhibition constant of hexokinase for Glc in liver |
| Km_hk_Atp_L | Michaelis constant of hexokinase for Atp in liver |
| Ki_hk_Atp_L | inhibition constant of hexokinase for Atp in liver |
| Vmax_ppp_G6p_L | reaction rate constant of pentose phosphate pathway for G6p in liver |
| Km_ppp_G6p_L | Michaelis constant of pentose phosphate pathway for G6p in liver |
| Km_ppp_Nadp_L | Michaelis constant of pentose phosphate pathway for Nadp in liver |
| Vmax_g6pase_G6p_L | reaction rate constant of G6Pase for G6p in liver |
| Km_g6pase_G6p_L | Michaelis constant of G6Pase for G6p in liver |
| Vmax_gs_G6p_L | reaction rate constant of glycogen synthase for G6p in liver |
| Km_gs_G6p_L | Michaelis constant of glycogen synthase for G6p in liver |
| n_gs_G6p_L | Hill coefficient of glycogen synthase for G6p in liver |
| Km_gs_Utp_L | Michaelis constant of glycogen synthase for G6p in liver |
| Glygn_L_max | maximum glycogen concentration in liver |
| Km_gs_Glygn_L | Michaelis constant of glycogen synthase for glycogen in liver |

| | |
|---|---|
| Vmax_gd_Glygn_L | reaction rate constant of glycogen degradation for glycogen in liver |
| Km_gd_Glygn_L | Michaelis constant of glycogen degradation for glycogen in liver |
| Km_gd_Phos_L | Michaelis constant of glycogen degradation for phosphate in liver |
| Vmax_pfk_G6p_L | reaction rate constant of phosphofructokinase for G6p in liver |
| Km_pfk_G6p_L | Michaelis constant of phosphofructokinase for G6p in liver |
| Km_pfk_Atp_L | Michaelis constant of phosphofructokinase for Atp in liver |
| Ki_pfk_Atp_L | inhibition constant of phosphofructokinase for Atp in liver |
| Km_pfk_Adp_L | Michaelis constant of phosphofructokinase for Adp in liver |
| Ki_pfk_Gap_L | inhibition constant of phosphofructokinase for Gap in liver |
| b_pfk_Gap_L | coefficient of phosphofructokinase for Gap in liver |
| Vmax_fbp_Gap_L | reaction rate constant of fructose bisphosphatase for Gap in liver |
| Km_fbp_Gap_L | Michaelis constant of fructose bisphosphatase for Gap in liver |
| Vmax_pk_Gap_L | reaction rate constant of pyruvate kinase for Gap in liver |
| Km_pk_Gap_L | Michaelis constant of pyruvate kinase for Gap in liver |
| b_pk_Accoa_LM | coefficient of pyruvate kinase for mitochondrial Accoa in liver |
| Ki_pk_Accoa_LM | inhibition constant of pyruvate kinase for mitochondrial Accoa in liver |
| Km_pk_Adp_L | Michaelis constant of pyruvate kinase for Adp in liver |
| Vmax_pepck_Pyr_L | reaction rate constant of phosphoenolpyruvate carboxykinase for Pyr in liver |
| Km_pepck_Pyr_L | Michaelis constant of phosphoenolpyruvate carboxykinase for Pyr in liver |
| Km_pepck_Atp_L | Michaelis constant of phosphoenolpyruvate carboxykinase for Atp in liver |
| Km_pepck_Gtp_L | Michaelis constant of phosphoenolpyruvate carboxykinase for Gtp in liver |
| Vdif_pyrt_Pyr_B_L | rate constant of pyruvate transporter in liver |
| Kdif_pyrt_Pyr_B_L | dissociation constant of pyruvate transporter for Pyr_B in liver |
| Kdif_pyrt_Pyr_L | dissociation constant of pyruvate transporter in liver |
| Vdif_lact_Lac_B_L | rate constant of lactate transporter in liver |
| Kdif_lact_Lac_L | dissociation constant of lactate transporter in liver |
| Kdif_lact_Lac_B_L | dissociation constant of lactate transporter for Lac_B in liver |
| Vmax_ldh_Pyr_L | rate constant of lactate dehydrogenase for Pyr in liver |
| Km_ldh_Pyr_L | Michaelis constant of lactate dehydrogenase for Pyr in liver |
| Km_ldh_Nadh_L | Michaelis constant of lactate dehydrogenase for Nadh in liver |
| Km_ldh_Lac_L | Michaelis constant of lactate dehydrogenase for Lac in liver |
| Km_ldh_Nad_L | Michaelis constant of lactate dehydrogenase for Nad in liver |
| Keq_ldh_Lac_L | equilibrium constant of lactate dehydrogenase for Lac in liver |
| Vdif_alat_Ala_B_L | rate constant of alanine transporter in liver |
| Kdif_alat_Ala_L | dissociation constant of alanine transporter in liver |
| Kdif_alat_Ala_B_L | dissociation constant of alanine transporter for Ala_B in liver |

| | |
|---|---|
| Vmax_alata_Pyr_L | rate constant of alanine transporter for Pyr in liver |
| Km_alata_Pyr_L | Michaelis constant of alanine aminotransferase for Pyr in liver |
| Km_alata_Ala_L | Michaelis constant of alanine aminotransferase for Ala in liver |
| Keq_alata_Ala_L | equilibrium constant of alanine aminotransferase for Ala in liver |
| Vmax_pdh_Pyr_L | rate constant of pyruvate dehydrogenase for Pyr in liver |
| Km_pdh_Pyr_L | Michaelis constant of pyruvate dehydrogenase for Pyr in liver |
| Ki_pdh_Accoa_LM | inhibition constant of pyruvate dehydrogenase for mitochondrial Accoa in liver |
| Km_pdh_Nad_L | Michaelis constant of pyruvate dehydrogenase for Nad in liver |
| Vmax_tca_Accoa_LM | rate constant of TCA cycle for mitochondrial Accoa in liver |
| Km_tca_Accoa_LM | Michaelis constant of TCA cycle for mitochondrial Accoa in liver |
| Km_tca_Phos_L | Michaelis constant of TCA cycle for phosphate in liver |
| Km_tca_Adp_L | Michaelis constant of TCA cycle for Adp in liver |
| Km_tca_Pyr_L | Michaelis constant of TCA cycle for Pyr in liver |
| Km_tca_Nad_L | Michaelis constant of TCA cycle for Nad in liver |
| Km_tca_Fad_L | Michaelis constant of TCA cycle for Fad in liver |
| Vdif_ffat_FFA_B_L | rate constant of FFA transporter in liver |
| Kdif_ffat_FFA_B_L | dissociation constant of FFA transporter for FFA_B in liver |
| Kdif_ffat_FFA_L | dissociation constant of FFA transporter in liver |
| Vmax_ffat_FFA_B_L | rate constant of active FFA transporter in liver |
| Km_ffat_FFA_B_L | Michaelis constant of active FFA transporter in liver |
| Vmax_tgsyn_FFA_L | rate constant of TG synthesis for FFA in liver |
| Km_tgsyn_FFA_L | Michaelis constant of TG synthesis for FFA in liver |
| Km_tgsyn_Glycp_L | Michaelis constant of TG synthesis for Glycp in liver |
| Km_tgsyn_Atp_L | Michaelis constant of TG synthesis for Atp in liver |
| TG_L_max | maximum TG concentration in liver |
| Km_tgsyn_TG_L | Michaelis constant of TG synthesis for TG in liver |
| Vmax_tgdeg_TG_L | rate constant of TG degradation for TG in liver |
| Km_tgdeg_TG_L | Michaelis constant of TG degradation for TG in liver |
| Vdif_glyct_Glyc_B_L | rate constant of glycerol transporter in liver |
| Kdif_glyct_Glyc_B_L | dissociation constant of glycerol transporter for Glyc_B in liver |
| Kdif_glyct_Glyc_L | dissociation constant of glycerol transporter in liver |
| Vmax_glyk_Glyc_L | rate constant of glycerol kinase for Glyc in liver |
| Km_glyk_Glyc_L | Michaelis constant of glycerol kinase for Glyc in liver |
| Km_glyk_Atp_L | Michaelis constant of glycerol kinase for Atp in liver |
| Ki_glyk_Gap_L | inhibition constant of glycerol kinase for Gap in liver |
| Vdif_tgt_TG_B_L | rate constant of TG transporter in liver |

| | |
|---|---|
| Kdif_tgt_TG_B_L | dissociation constant of TG transporter in liver |
| Keq_tgt_TG_L | equilibrium constant of TG transporter in liver |
| Vmax_tgt_TG_B_L | rate constant of active TG transporter in liver |
| Km_tgt_TG_L | Michaelis constant of active TG transporter in liver |
| Vmax_g3pd_Gap_L | rate constant of glyceraldehyde-3-phosphate dehydrogenase for Gap in liver |
| Km_g3pd_Gap_L | Michaelis constant of glyceraldehyde-3-phosphate dehydrogenase for Gap in liver |
| Km_g3pd_Glycp_L | Michaelis constant of glyceraldehyde-3-phosphate dehydrogenase for Glycp in liver |
| Km_g3pd_Nadh_L | Michaelis constant of glyceraldehyde-3-phosphate dehydrogenase for Nadh in liver |
| Km_g3pd_Nad_L | Michaelis constant of glyceraldehyde-3-phosphate dehydrogenase for Nad in liver |
| Keq_g3pd_Glycp_L | equilibrium constant of glyceraldehyde-3-phosphate dehydrogenase for Glycp in liver |
| Vmax_boxid_FFA_L | rate constant of beta-oxidation for FFA in liver |
| Km_boxid_FFA_L | Michaelis constant of beta-oxidation for FFA in liver |
| Km_boxid_Atp_L | Michaelis constant of beta-oxidation for Atp in liver |
| Ki_boxid_Accoa_LM | inhibition constant of beta-oxidation for mitochondrial Accoa in liver |
| Ki_boxid_Malcoa_L | inhibition constant of beta-oxidation for Malcoa in liver |
| Km_boxid_Nad_L | Michaelis constant of beta-oxidation for Nad in liver |
| Km_boxid_Fad_L | Michaelis constant of beta-oxidation for Fad in liver |
| Vmax_accoat_Accoa_LM_LC | rate constant of mitochondrial Accoa transport to cytoplasm in liver |
| Km_accoat_Accoa_LM | Michaelis constant of mitochondrial Accoa transporter to cytoplasm in liver |
| Km_accoat_Atp_L | Michaelis constant of mitochondrial Accoa transporter to cytoplasm for Atp in liver |
| Km_accoat_Pyr_L | activation constant of mitochondrial Accoa transport to cytoplasm for Pyr in liver |
| Vmax_bhbsyn_Accoa_LM | rate constant of Bhb synthesis for mitochondrial Accoa in liver |
| Km_bhbsyn_Accoa_LM | Michaelis constant of Bhb synthesis for mitochondrial Accoa in liver |
| Km_bhbsyn_Nadh_L | Michaelis constant of Bhb synthesis for Nadh in liver |
| n_bhbsyn_Pyr_L | Hill coefficient of Bhb synthesis for Pyr in liver |
| Ki_bhbsyn_Pyr_L | inhibition constant of Bhb synthesis for Pyr in liver |
| Vmax_bhbdeg_Bhb_L | rate constant of Bhb degradation for Bhb in liver |
| Km_bhbdeg_Bhb_L | Michaelis constant of Bhb degradation for Bhb in liver |
| Km_bhbdeg_Atp_L | Michaelis constant of Bhb degradation for Atp in liver |
| Km_bhbdeg_Nad_L | Michaelis constant of Bhb degradation for Nad in liver |
| Vdif_bhbt_Bhb_B_L | rate constant of Bhb transporter in liver |
| Kdif_bhbt_Bhb_B_L | dissociation constant of Bhb transporter for Bhb_B in liver |
| Kdif_bhbt_Bhb_L | dissociation constant of Bhb transporter in liver |
| Vmax_lipog1_Accoa_LC | rate constant of lipogenesis 1 for Accoa in liver |
| Km_lipog1_Accoa_LC | Michaelis constant of lipogenesis 1 for Accoa in liver |

| | |
|---|---|
| Km_lipog1_Atp_L | Michaelis constant of lipogenesis 1 for Atp in liver |
| Vmax_lipog2_Malcoa_L | rate constant of lipogenesis 2 for Malcoa in liver |
| Km_lipog2_Malcoa_L | Michaelis constant of lipogenesis 2 for Malcoa in liver |
| Km_lipog2_Adp_L | Michaelis constant of lipogenesis 2 for Adp in liver |
| Km_lipog2_Nadph_L | Michaelis constant of lipogenesis 2 for Nadph in liver |
| Vmax_cholsyn1_Accoa_LC | rate constant of cholesterol synthesis 1 for Accoa in liver |
| Km_cholsyn1_Accoa_LC | Michaelis constant of cholesterol synthesis 1 for Accoa in liver |
| Vmax_cholsyn2_Hmgcoa_L | rate constant of cholesterol synthesis 2 for Hmgcoa in liver |
| Km_cholsyn2_Hmgcoa_L | Michaelis constant of cholesterol synthesis 2 for Hmgcoa in liver |
| Km_cholsyn2_Atp_L | Michaelis constant of cholesterol synthesis 2 for Atp in liver |
| Km_cholsyn2_Fadh_L | Michaelis constant of cholesterol synthesis 2 for Fadh in liver |
| Km_cholsyn2_Nadph_L | Michaelis constant of cholesterol synthesis 2 for Nadph in liver |
| Vmax_cholt_Chol_L | rate constant of active cholesterol transporter in liver |
| Km_cholt_Chol_L | Michaelis constant of active cholesterol transporter in liver |
| Vmax_atpsynf_Fadh_L | rate constant of Atp synthesis from Fadh in liver |
| Km_atpsynf_Fadh_L | Michaelis constant of Atp synthesis from Fadh for Fadh in liver |
| Km_atpsynf_Adp_L | Michaelis constant of Atp synthesis from Fadh for Adp in liver |
| Vmax_atpsynn_Nadh_L | rate constant of Atp synthesis from Nadh in liver |
| Km_atpsynn_Nadh_L | Michaelis constant of Atp synthesis from Nadh for Nadh in liver |
| Km_atpsynn_Adp_L | Michaelis constant of Atp synthesis from Nadh for Adp in liver |
| Vmax_atpuse_Atp_L | rate constant of Atp utilization for Atp in liver |
| Km_atpuse_Atp_L | Michaelis constant of Atp utilization for Atp in liver |
| Vmax_ampreg_Amp_L | rate constant of Amp regeneration for Amp in liver |
| Km_ampreg_Amp_L | Michaelis constant of Amp regeneration for Amp in liver |
| Km_ampreg_Atp_L | Michaelis constant of Amp regeneration for Atp in liver |
| Km_ampreg_Adp_L | Michaelis constant of Amp regeneration for Adp in liver |
| Vmax_nadhk_Nadh_L | rate constant of Nadh kinase for Nadh in liver |
| Km_nadhk_Nadh_L | Michaelis constant of Nadh kinase for Nadh in liver |
| Km_nadhk_Atp_L | Michaelis constant of Nadh kinase for Atp in liver |
| Vmax_nadhuse_Nadh_L | rate constant of Nadh utilization for Nadh in liver |
| Km_nadhuse_Nadh_L | Michaelis constant of Nadh utilization for Nadh in liver |
| Vmax_gdpreg_Gdp_L | rate constant of Gdp regeneration for Gdp in liver |
| Km_gdpreg_Gdp_L | Michaelis constant of Gdp regeneration for Gdp in liver |
| Km_gdpreg_Atp_L | Michaelis constant of Gdp regeneration for Atp in liver |
| Km_gdpreg_Gtp_L | Michaelis constant of Gdp regeneration for Gtp in liver |
| Km_gdpreg_Adp_L | Michaelis constant of Gdp regeneration for Adp in liver |

| | |
|---|---|
| Keq_gdpreg_Gtp_L | equilibrium constant of Gdp regeneration for Gtp in liver |
| Vmax_gtpuse_Gtp_L | rate constant of Gtp utilization for Gtp in liver |
| Km_gtpuse_Gtp_L | Michaelis constant of Gtp utilization for Gtp in liver |
| Vmax_udpreg_Udp_L | rate constant of Udp regeneration for Udp in liver |
| Km_udpreg_Udp_L | Michaelis constant of Udp regeneration for Udp in liver |
| Km_udpreg_Atp_L | Michaelis constant of Udp regeneration for Atp in liver |
| Km_udpreg_Utp_L | Michaelis constant of Udp regeneration for Utp in liver |
| Km_udpreg_Adp_L | Michaelis constant of Udp regeneration for Adp in liver |
| Keq_udpreg_Utp_L | equilibrium constant of Udp regeneration for Utp in liver |
| Vmax_utpuse_Utp_L | rate constant of Utp utilization for Utp in liver |
| Km_utpuse_Utp_L | Michaelis constant of Utp utilization for Utp in liver |
| Vmax_nadphuse_Nadph_L | rate constant of Nadph utilization for Nadph in liver |
| Km_nadphuse_Nadph_L | Michaelis constant of Nadph utilization for Nadph in liver |
| Vmax_fadhuse_Fadh_L | rate constant of Fadh utilization for Fadh in liver |
| Km_fadhuse_Fadh_L | Michaelis constant of Fadh utilization for Fadh in liver |
| Vmax_ck_Cre_L | rate constant of creatine kinase for Cre in liver |
| Km_ck_Cre_L | Michaelis constant of creatine kinase for Cre in liver |
| Km_ck_Atp_L | Michaelis constant of creatine kinase for Atp in liver |
| Km_ck_Crep_L | Michaelis constant of creatine kinase for Crep in liver |
| Km_ck_Adp_L | Michaelis constant of creatine kinase for Adp in liver |
| Keq_ck_Crep_L | equilibrium constant of creatine kinase for Crep in liver |
| Phos_L | phosphate in liver |
| alpha_base_M | insulin base factor in skeletal muscle |
| alpha_band_M | insulin allowable width in skeletal muscle |
| Km_Ins_B_M | Michaelis constant for insulin in skeletal muscle |
| n_Ins_B_M | Hill coefficient for insulin in skeletal muscle |
| Vdif_glut4_Glc_B_M | rate constant of glucose transporter type 2 in skeletal muscle |
| Kdif_glut4_Glc_B_M | dissociation constant of glucose transporter type 2 for Glc_B in skeletal muscle |
| Kdif_glut4_Glc_M | dissociation constant of glucose transporter type 2 in skeletal muscle |
| Vmax_hk_Glc_M | reaction rate constant of hexokinase for Glc in skeletal muscle |
| Km_hk_Glc_M | Michaelis constant of hexokinase for Glc in skeletal muscle |
| Ki_hk_G6p_M | inhibition constant of hexokinase for Glc in skeletal muscle |
| Km_hk_Atp_M | Michaelis constant of hexokinase for Atp in skeletal muscle |
| Ki_hk_Atp_M | inhibition constant of hexokinase for Atp in skeletal muscle |
| Vmax_ppp_G6p_M | reaction rate constant of pentose phosphate pathway for G6p in skeletal muscle |
| Km_ppp_G6p_M | Michaelis constant of pentose phosphate pathway for G6p in skeletal muscle |

| | |
|---|---|
| Km_ppp_Nadp_M | Michaelis constant of pentose phosphate pathway for Nadp in skeletal muscle |
| Vmax_g6pase_G6p_M | reaction rate constant of G6Pase for G6p in skeletal muscle |
| Km_g6pase_G6p_M | Michaelis constant of G6Pase for G6p in skeletal muscle |
| Vmax_gs_G6p_M | reaction rate constant of glycogen synthase for G6p in skeletal muscle |
| Km_gs_G6p_M | Michaelis constant of glycogen synthase for G6p in skeletal muscle |
| n_gs_G6p_M | Hill coefficient of glycogen synthase for G6p in skeletal muscle |
| Km_gs_Utp_M | Michaelis constant of glycogen synthase for G6p in skeletal muscle |
| Glygn_M_max | maximum glycogen concentration in skeletal muscle |
| Km_gs_Glygn_M | Michaelis constant of glycogen synthase for glycogen in skeletal muscle |
| Vmax_gd_Glygn_M | reaction rate constant of glycogen degradation for glycogen in skeletal muscle |
| Km_gd_Glygn_M | Michaelis constant of glycogen degradation for glycogen in skeletal muscle |
| Km_gd_Phos_M | Michaelis constant of glycogen degradation for phosphate in skeletal muscle |
| Vmax_pfk_G6p_M | reaction rate constant of phosphofructokinase for G6p in skeletal muscle |
| Km_pfk_G6p_M | Michaelis constant of phosphofructokinase for G6p in skeletal muscle |
| Km_pfk_Atp_M | Michaelis constant of phosphofructokinase for Atp in skeletal muscle |
| Ki_pfk_Atp_M | inhibition constant of phosphofructokinase for Atp in skeletal muscle |
| Km_pfk_Adp_M | Michaelis constant of phosphofructokinase for Adp in skeletal muscle |
| Ki_pfk_Gap_M | inhibition constant of phosphofructokinase for Gap in skeletal muscle |
| b_pfk_Gap_M | coefficient of phosphofructokinase for Gap in skeletal muscle |
| Vmax_fbp_Gap_M | reaction rate constant of fructose bisphosphatase for Gap in skeletal muscle |
| Km_fbp_Gap_M | Michaelis constant of fructose bisphosphatase for Gap in skeletal muscle |
| Vmax_pk_Gap_M | reaction rate constant of pyruvate kinase for Gap in skeletal muscle |
| Km_pk_Gap_M | Michaelis constant of pyruvate kinase for Gap in skeletal muscle |
| b_pk_Accoa_MM | coefficient of pyruvate kinase for mitochondrial Accoa in skeletal muscle |
| Ki_pk_Accoa_MM | inhibition constant of pyruvate kinase for mitochondrial Accoa in skeletal muscle |
| Km_pk_Adp_M | Michaelis constant of pyruvate kinase for Adp in skeletal muscle |
| Vmax_pepck_Pyr_M | reaction rate constant of phosphoenolpyruvate carboxykinase for Pyr in skeletal muscle |
| Km_pepck_Pyr_M | Michaelis constant of phosphoenolpyruvate carboxykinase for Pyr in skeletal muscle |
| Km_pepck_Atp_M | Michaelis constant of phosphoenolpyruvate carboxykinase for Atp in skeletal muscle |
| Km_pepck_Gtp_M | Michaelis constant of phosphoenolpyruvate carboxykinase for Gtp in skeletal muscle |
| Vdif_pyrt_Pyr_B_M | rate constant of pyruvate transporter in skeletal muscle |
| Kdif_pyrt_Pyr_B_M | dissociation constant of pyruvate transporter for Pyr_B in skeletal muscle |
| Kdif_pyrt_Pyr_M | dissociation constant of pyruvate transporter in skeletal muscle |

| | |
|---|---|
| Vdif_lact_Lac_B_M | rate constant of lactate transporter in skeletal muscle |
| Kdif_lact_Lac_M | dissociation constant of lactate transporter in skeletal muscle |
| Kdif_lact_Lac_B_M | dissociation constant of lactate transporter for Lac_B in skeletal muscle |
| Vmax_ldh_Pyr_M | rate constant of lactate dehydrogenase for Pyr in skeletal muscle |
| Km_ldh_Pyr_M | Michaelis constant of lactate dehydrogenase for Pyr in skeletal muscle |
| Km_ldh_Nadh_M | Michaelis constant of lactate dehydrogenase for Nadh in skeletal muscle |
| Km_ldh_Lac_M | Michaelis constant of lactate dehydrogenase for Lac in skeletal muscle |
| Km_ldh_Nad_M | Michaelis constant of lactate dehydrogenase for Nad in skeletal muscle |
| Keq_ldh_Lac_M | equilibrium constant of lactate dehydrogenase for Lac in skeletal muscle |
| Vdif_alat_Ala_B_M | rate constant of alanine transporter in skeletal muscle |
| Kdif_alat_Ala_M | dissociation constant of alanine transporter in skeletal muscle |
| Kdif_alat_Ala_B_M | dissociation constant of alanine transporter for Ala_B in skeletal muscle |
| Vmax_alata_Pyr_M | rate constant of alanine transporter for Pyr in skeletal muscle |
| Km_alata_Pyr_M | Michaelis constant of alanine aminotransferase for Pyr in skeletal muscle |
| Km_alata_Ala_M | Michaelis constant of alanine aminotransferase for Ala in skeletal muscle |
| Keq_alata_Ala_M | equilibrium constant of alanine aminotransferase for Ala in skeletal muscle |
| Vmax_pdh_Pyr_M | rate constant of pyruvate dehydrogenase for Pyr in skeletal muscle |
| Km_pdh_Pyr_M | Michaelis constant of pyruvate dehydrogenase for Pyr in skeletal muscle |
| Ki_pdh_Accoa_MM | inhibition constant of pyruvate dehydrogenase for mitochondrial Accoa in skeletal muscle |
| Km_pdh_Nad_M | Michaelis constant of pyruvate dehydrogenase for Nad in skeletal muscle |
| Vmax_tca_Accoa_MM | rate constant of TCA cycle for mitochondrial Accoa in skeletal muscle |
| Km_tca_Accoa_MM | Michaelis constant of TCA cycle for mitochondrial Accoa in skeletal muscle |
| Km_tca_Phos_M | Michaelis constant of TCA cycle for phosphate in skeletal muscle |
| Km_tca_Adp_M | Michaelis constant of TCA cycle for Adp in skeletal muscle |
| Km_tca_Pyr_M | Michaelis constant of TCA cycle for Pyr in skeletal muscle |
| Km_tca_Nad_M | Michaelis constant of TCA cycle for Nad in skeletal muscle |
| Km_tca_Fad_M | Michaelis constant of TCA cycle for Fad in skeletal muscle |
| Vdif_ffat_FFA_B_M | rate constant of FFA transporter in skeletal muscle |
| Kdif_ffat_FFA_B_M | dissociation constant of FFA transporter for FFA_B in skeletal muscle |
| Kdif_ffat_FFA_M | dissociation constant of FFA transporter in skeletal muscle |
| Vmax_ffat_FFA_B_M | rate constant of active FFA transporter in skeletal muscle |
| Km_ffat_FFA_B_M | Michaelis constant of active FFA transporter in skeletal muscle |
| Vmax_tgsyn_FFA_M | rate constant of TG synthesis for FFA in skeletal muscle |
| Km_tgsyn_FFA_M | Michaelis constant of TG synthesis for FFA in skeletal muscle |
| Km_tgsyn_Glycp_M | Michaelis constant of TG synthesis for Glycp in skeletal muscle |

| | |
|---|---|
| Km_tgsyn_Atp_M | Michaelis constant of TG synthesis for Atp in skeletal muscle |
| TG_M_max | maximum TG concentration in skeletal muscle |
| Km_tgsyn_TG_M | Michaelis constant of TG synthesis for TG in skeletal muscle |
| Vmax_tgdeg_TG_M | rate constant of TG degradation for TG in skeletal muscle |
| Km_tgdeg_TG_M | Michaelis constant of TG degradation for TG in skeletal muscle |
| Vdif_glyct_Glyc_B_M | rate constant of glycerol transporter in skeletal muscle |
| Kdif_glyct_Glyc_B_M | dissociation constant of glycerol transporter for Glyc_B in skeletal muscle |
| Kdif_glyct_Glyc_M | dissociation constant of glycerol transporter in skeletal muscle |
| Vmax_glyk_Glyc_M | rate constant of glycerol kinase for Glyc in skeletal muscle |
| Km_glyk_Glyc_M | Michaelis constant of glycerol kinase for Glyc in skeletal muscle |
| Km_glyk_Atp_M | Michaelis constant of glycerol kinase for Atp in skeletal muscle |
| Ki_glyk_Gap_M | inhibition constant of glycerol kinase for Gap in skeletal muscle |
| Vdif_tgt_TG_B_M | rate constant of TG transporter in skeletal muscle |
| Kdif_tgt_TG_B_M | dissociation constant of TG transporter in skeletal muscle |
| Keq_tgt_TG_M | equilibrium constant of TG transporter in skeletal muscle |
| Vmax_tgt_TG_B_M | rate constant of active TG transporter in skeletal muscle |
| Km_tgt_TG_M | Michaelis constant of active TG transporter in skeletal muscle |
| Vmax_g3pd_Gap_M | rate constant of glyceraldehyde-3-phosphate dehydrogenase for Gap in skeletal muscle |
| Km_g3pd_Gap_M | Michaelis constant of glyceraldehyde-3-phosphate dehydrogenase for Gap in skeletal muscle |
| Km_g3pd_Glycp_M | Michaelis constant of glyceraldehyde-3-phosphate dehydrogenase for Glycp in skeletal muscle |
| Km_g3pd_Nadh_M | Michaelis constant of glyceraldehyde-3-phosphate dehydrogenase for Nadh in skeletal muscle |
| Km_g3pd_Nad_M | Michaelis constant of glyceraldehyde-3-phosphate dehydrogenase for Nad in skeletal muscle |
| Keq_g3pd_Glycp_M | equilibrium constant of glyceraldehyde-3-phosphate dehydrogenase for Glycp in skeletal muscle |
| Vmax_boxid_FFA_M | rate constant of beta-oxidation for FFA in skeletal muscle |
| Km_boxid_FFA_M | Michaelis constant of beta-oxidation for FFA in skeletal muscle |
| Km_boxid_Atp_M | Michaelis constant of beta-oxidation for Atp in skeletal muscle |
| Ki_boxid_Accoa_MM | inhibition constant of beta-oxidation for mitochondrial Accoa in skeletal muscle |
| Ki_boxid_Malcoa_M | inhibition constant of beta-oxidation for Malcoa in skeletal muscle |
| Km_boxid_Nad_M | Michaelis constant of beta-oxidation for Nad in skeletal muscle |
| Km_boxid_Fad_M | Michaelis constant of beta-oxidation for Fad in skeletal muscle |

| | |
|---|---|
| Vmax_accoat_Accoa_MM_MC | rate constant of mitochondrial Accoa transport to cytoplasm in skeletal muscle |
| Km_accoat_Accoa_MM | Michaelis constant of mitochondrial Accoa transporter to cytoplasm in skeletal muscle |
| Km_accoat_Atp_M | Michaelis constant of mitochondrial Accoa transporter to cytoplasm for Atp in skeletal muscle |
| Km_accoat_Pyr_M | activation constant of mitochondrial Accoa transport to cytoplasm for Pyr in skeletal muscle |
| Vmax_bhbsyn_Accoa_MM | rate constant of Bhb synthesis for mitochondrial Accoa in skeletal muscle |
| Km_bhbsyn_Accoa_MM | Michaelis constant of Bhb synthesis for mitochondrial Accoa in skeletal muscle |
| Km_bhbsyn_Nadh_M | Michaelis constant of Bhb synthesis for Nadh in skeletal muscle |
| n_bhbsyn_Pyr_M | Hill coefficient of Bhb synthesis for Pyr in skeletal muscle |
| Ki_bhbsyn_Pyr_M | inhibition constant of Bhb synthesis for Pyr in skeletal muscle |
| Vmax_bhbdeg_Bhb_M | rate constant of Bhb degradation for Bhb in skeletal muscle |
| Km_bhbdeg_Bhb_M | Michaelis constant of Bhb degradation for Bhb in skeletal muscle |
| Km_bhbdeg_Atp_M | Michaelis constant of Bhb degradation for Atp in skeletal muscle |
| Km_bhbdeg_Nad_M | Michaelis constant of Bhb degradation for Nad in skeletal muscle |
| Vdif_bhbt_Bhb_B_M | rate constant of Bhb transporter in skeletal muscle |
| Kdif_bhbt_Bhb_B_M | dissociation constant of Bhb transporter for Bhb_B in skeletal muscle |
| Kdif_bhbt_Bhb_M | dissociation constant of Bhb transporter in skeletal muscle |
| Vmax_lipog1_Accoa_MC | rate constant of lipogenesis 1 for Accoa in skeletal muscle |
| Km_lipog1_Accoa_MC | Michaelis constant of lipogenesis 1 for Accoa in skeletal muscle |
| Km_lipog1_Atp_M | Michaelis constant of lipogenesis 1 for Atp in skeletal muscle |
| Vmax_lipog2_Malcoa_M | rate constant of lipogenesis 2 for Malcoa in skeletal muscle |
| Km_lipog2_Malcoa_M | Michaelis constant of lipogenesis 2 for Malcoa in skeletal muscle |
| Km_lipog2_Adp_M | Michaelis constant of lipogenesis 2 for Adp in skeletal muscle |
| Km_lipog2_Nadph_M | Michaelis constant of lipogenesis 2 for Nadph in skeletal muscle |
| Vmax_cholsyn1_Accoa_MC | rate constant of cholesterol synthesis 1 for Accoa in skeletal muscle |
| Km_cholsyn1_Accoa_MC | Michaelis constant of cholesterol synthesis 1 for Accoa in skeletal muscle |
| Vmax_cholsyn2_Hmgcoa_M | rate constant of cholesterol synthesis 2 for Hmgcoa in skeletal muscle |
| Km_cholsyn2_Hmgcoa_M | Michaelis constant of cholesterol synthesis 2 for Hmgcoa in skeletal muscle |
| Km_cholsyn2_Atp_M | Michaelis constant of cholesterol synthesis 2 for Atp in skeletal muscle |
| Km_cholsyn2_Fadh_M | Michaelis constant of cholesterol synthesis 2 for Fadh in skeletal muscle |
| Km_cholsyn2_Nadph_M | Michaelis constant of cholesterol synthesis 2 for Nadph in skeletal muscle |
| Vmax_cholt_Chol_M | rate constant of active cholesterol transporter in skeletal muscle |
| Km_cholt_Chol_M | Michaelis constant of active cholesterol transporter in skeletal muscle |
| Vmax_atpsynf_Fadh_M | rate constant of Atp synthesis from Fadh in skeletal muscle |

| | |
|---|---|
| Km_atpsynf_Fadh_M | Michaelis constant of Atp synthesis from Fadh for Fadh in skeletal muscle |
| Km_atpsynf_Adp_M | Michaelis constant of Atp synthesis from Fadh for Adp in skeletal muscle |
| Vmax_atpsynn_Nadh_M | rate constant of Atp synthesis from Nadh in skeletal muscle |
| Km_atpsynn_Nadh_M | Michaelis constant of Atp synthesis from Nadh for Nadh in skeletal muscle |
| Km_atpsynn_Adp_M | Michaelis constant of Atp synthesis from Nadh for Adp in skeletal muscle |
| Vmax_atpuse_Atp_M | rate constant of Atp utilization for Atp in skeletal muscle |
| Km_atpuse_Atp_M | Michaelis constant of Atp utilization for Atp in skeletal muscle |
| Vmax_ampreg_Amp_M | rate constant of Amp regeneration for Amp in skeletal muscle |
| Km_ampreg_Amp_M | Michaelis constant of Amp regeneration for Amp in skeletal muscle |
| Km_ampreg_Atp_M | Michaelis constant of Amp regeneration for Atp in skeletal muscle |
| Km_ampreg_Adp_M | Michaelis constant of Amp regeneration for Adp in skeletal muscle |
| Vmax_nadhk_Nadh_M | rate constant of Nadh kinase for Nadh in skeletal muscle |
| Km_nadhk_Nadh_M | Michaelis constant of Nadh kinase for Nadh in skeletal muscle |
| Km_nadhk_Atp_M | Michaelis constant of Nadh kinase for Atp in skeletal muscle |
| Vmax_nadhuse_Nadh_M | rate constant of Nadh utilization for Nadh in skeletal muscle |
| Km_nadhuse_Nadh_M | Michaelis constant of Nadh utilization for Nadh in skeletal muscle |
| Vmax_gdpreg_Gdp_M | rate constant of Gdp regeneration for Gdp in skeletal muscle |
| Km_gdpreg_Gdp_M | Michaelis constant of Gdp regeneration for Gdp in skeletal muscle |
| Km_gdpreg_Atp_M | Michaelis constant of Gdp regeneration for Atp in skeletal muscle |
| Km_gdpreg_Gtp_M | Michaelis constant of Gdp regeneration for Gtp in skeletal muscle |
| Km_gdpreg_Adp_M | Michaelis constant of Gdp regeneration for Adp in skeletal muscle |
| Keq_gdpreg_Gtp_M | equilibrium constant of Gdp regeneration for Gtp in skeletal muscle |
| Vmax_gtpuse_Gtp_M | rate constant of Gtp utilization for Gtp in skeletal muscle |
| Km_gtpuse_Gtp_M | Michaelis constant of Gtp utilization for Gtp in skeletal muscle |
| Vmax_udpreg_Udp_M | rate constant of Udp regeneration for Udp in skeletal muscle |
| Km_udpreg_Udp_M | Michaelis constant of Udp regeneration for Udp in skeletal muscle |
| Km_udpreg_Atp_M | Michaelis constant of Udp regeneration for Atp in skeletal muscle |
| Km_udpreg_Utp_M | Michaelis constant of Udp regeneration for Utp in skeletal muscle |
| Km_udpreg_Adp_M | Michaelis constant of Udp regeneration for Adp in skeletal muscle |
| Keq_udpreg_Utp_M | equilibrium constant of Udp regeneration for Utp in skeletal muscle |
| Vmax_utpuse_Utp_M | rate constant of Utp utilization for Utp in skeletal muscle |
| Km_utpuse_Utp_M | Michaelis constant of Utp utilization for Utp in skeletal muscle |
| Vmax_nadphuse_Nadph_M | rate constant of Nadph utilization for Nadph in skeletal muscle |
| Km_nadphuse_Nadph_M | Michaelis constant of Nadph utilization for Nadph in skeletal muscle |
| Vmax_fadhuse_Fadh_M | rate constant of Fadh utilization for Fadh in skeletal muscle |
| Km_fadhuse_Fadh_M | Michaelis constant of Fadh utilization for Fadh in skeletal muscle |

| | |
|---|---|
| Vmax_ck_Cre_M | rate constant of creatine kinase for Cre in skeletal muscle |
| Km_ck_Cre_M | Michaelis constant of creatine kinase for Cre in skeletal muscle |
| Km_ck_Atp_M | Michaelis constant of creatine kinase for Atp in skeletal muscle |
| Km_ck_Crep_M | Michaelis constant of creatine kinase for Crep in skeletal muscle |
| Km_ck_Adp_M | Michaelis constant of creatine kinase for Adp in skeletal muscle |
| Keq_ck_Crep_M | equilibrium constant of creatine kinase for Crep in skeletal muscle |
| Phos_M | phosphate in skeletal muscle |
| alpha_base_A | insulin base factor in adipose tissue |
| alpha_band_A | insulin allowable width in adipose tissue |
| Km_Ins_B_A | Michaelis constant for insulin in adipose tissue |
| n_Ins_B_A | Hill coefficient for insulin in adipose tissue |
| Vdif_glut4_Glc_B_A | rate constant of glucose transporter type 2 in adipose tissue |
| Kdif_glut4_Glc_B_A | dissociation constant of glucose transporter type 2 for Glc_B in adipose tissue |
| Kdif_glut4_Glc_A | dissociation constant of glucose transporter type 2 in adipose tissue |
| Vmax_hk_Glc_A | reaction rate constant of hexokinase for Glc in adipose tissue |
| Km_hk_Glc_A | Michaelis constant of hexokinase for Glc in adipose tissue |
| Ki_hk_G6p_A | inhibition constant of hexokinase for Glc in adipose tissue |
| Km_hk_Atp_A | Michaelis constant of hexokinase for Atp in adipose tissue |
| Ki_hk_Atp_A | inhibition constant of hexokinase for Atp in adipose tissue |
| Vmax_ppp_G6p_A | reaction rate constant of pentose phosphate pathway for G6p in adipose tissue |
| Km_ppp_G6p_A | Michaelis constant of pentose phosphate pathway for G6p in adipose tissue |
| Km_ppp_Nadp_A | Michaelis constant of pentose phosphate pathway for Nadp in adipose tissue |
| Vmax_g6pase_G6p_A | reaction rate constant of G6Pase for G6p in adipose tissue |
| Km_g6pase_G6p_A | Michaelis constant of G6Pase for G6p in adipose tissue |
| Vmax_gs_G6p_A | reaction rate constant of glycogen synthase for G6p in adipose tissue |
| Km_gs_G6p_A | Michaelis constant of glycogen synthase for G6p in adipose tissue |
| n_gs_G6p_A | Hill coefficient of glycogen synthase for G6p in adipose tissue |
| Km_gs_Utp_A | Michaelis constant of glycogen synthase for G6p in adipose tissue |
| Glygn_A_max | maximum glycogen concentration in adipose tissue |
| Km_gs_Glygn_A | Michaelis constant of glycogen synthase for glycogen in adipose tissue |
| Vmax_gd_Glygn_A | reaction rate constant of glycogen degradation for glycogen in adipose tissue |
| Km_gd_Glygn_A | Michaelis constant of glycogen degradation for glycogen in adipose tissue |
| Km_gd_Phos_A | Michaelis constant of glycogen degradation for phosphate in adipose tissue |
| Vmax_pfk_G6p_A | reaction rate constant of phosphofructokinase for G6p in adipose tissue |
| Km_pfk_G6p_A | Michaelis constant of phosphofructokinase for G6p in adipose tissue |
| Km_pfk_Atp_A | Michaelis constant of phosphofructokinase for Atp in adipose tissue |

| | |
|---|---|
| Ki_pfk_Atp_A | inhibition constant of phosphofructokinase for Atp in adipose tissue |
| Km_pfk_Adp_A | Michaelis constant of phosphofructokinase for Adp in adipose tissue |
| Ki_pfk_Gap_A | inhibition constant of phosphofructokinase for Gap in adipose tissue |
| b_pfk_Gap_A | coefficient of phosphofructokinase for Gap in adipose tissue |
| Vmax_fbp_Gap_A | reaction rate constant of fructose bisphosphatase for Gap in adipose tissue |
| Km_fbp_Gap_A | Michaelis constant of fructose bisphosphatase for Gap in adipose tissue |
| Vmax_pk_Gap_A | reaction rate constant of pyruvate kinase for Gap in adipose tissue |
| Km_pk_Gap_A | Michaelis constant of pyruvate kinase for Gap in adipose tissue |
| b_pk_Accoa_AM | coefficient of pyruvate kinase for mitochondrial Accoa in adipose tissue |
| Ki_pk_Accoa_AM | inhibition constant of pyruvate kinase for mitochondrial Accoa in adipose tissue |
| Km_pk_Adp_A | Michaelis constant of pyruvate kinase for Adp in adipose tissue |
| Vmax_pepck_Pyr_A | reaction rate constant of phosphoenolpyruvate carboxykinase for Pyr in adipose tissue |
| Km_pepck_Pyr_A | Michaelis constant of phosphoenolpyruvate carboxykinase for Pyr in adipose tissue |
| Km_pepck_Atp_A | Michaelis constant of phosphoenolpyruvate carboxykinase for Atp in adipose tissue |
| Km_pepck_Gtp_A | Michaelis constant of phosphoenolpyruvate carboxykinase for Gtp in adipose tissue |
| Vdif_pyrt_Pyr_B_A | rate constant of pyruvate transporter in adipose tissue |
| Kdif_pyrt_Pyr_B_A | dissociation constant of pyruvate transporter for Pyr_B in adipose tissue |
| Kdif_pyrt_Pyr_A | dissociation constant of pyruvate transporter in adipose tissue |
| Vdif_lact_Lac_B_A | rate constant of lactate transporter in adipose tissue |
| Kdif_lact_Lac_A | dissociation constant of lactate transporter in adipose tissue |
| Kdif_lact_Lac_B_A | dissociation constant of lactate transporter for Lac_B in adipose tissue |
| Vmax_ldh_Pyr_A | rate constant of lactate dehydrogenase for Pyr in adipose tissue |
| Km_ldh_Pyr_A | Michaelis constant of lactate dehydrogenase for Pyr in adipose tissue |
| Km_ldh_Nadh_A | Michaelis constant of lactate dehydrogenase for Nadh in adipose tissue |
| Km_ldh_Lac_A | Michaelis constant of lactate dehydrogenase for Lac in adipose tissue |
| Km_ldh_Nad_A | Michaelis constant of lactate dehydrogenase for Nad in adipose tissue |
| Keq_ldh_Lac_A | equilibrium constant of lactate dehydrogenase for Lac in adipose tissue |
| Vdif_alat_Ala_B_A | rate constant of alanine transporter in adipose tissue |
| Kdif_alat_Ala_A | dissociation constant of alanine transporter in adipose tissue |
| Kdif_alat_Ala_B_A | dissociation constant of alanine transporter for Ala_B in adipose tissue |
| Vmax_alata_Pyr_A | rate constant of alanine transporter for Pyr in adipose tissue |
| Km_alata_Pyr_A | Michaelis constant of alanine aminotransferase for Pyr in adipose tissue |
| Km_alata_Ala_A | Michaelis constant of alanine aminotransferase for Ala in adipose tissue |
| Keq_alata_Ala_A | equilibrium constant of alanine aminotransferase for Ala in adipose tissue |
| Vmax_pdh_Pyr_A | rate constant of pyruvate dehydrogenase for Pyr in adipose tissue |

| | |
|---|---|
| Km_pdh_Pyr_A | Michaelis constant of pyruvate dehydrogenase for Pyr in adipose tissue |
| Ki_pdh_Accoa_AM | inhibition constant of pyruvate dehydrogenase for mitochondrial Accoa in adipose tissue |
| Km_pdh_Nad_A | Michaelis constant of pyruvate dehydrogenase for Nad in adipose tissue |
| Vmax_tca_Accoa_AM | rate constant of TCA cycle for mitochondrial Accoa in adipose tissue |
| Km_tca_Accoa_AM | Michaelis constant of TCA cycle for mitochondrial Accoa in adipose tissue |
| Km_tca_Phos_A | Michaelis constant of TCA cycle for phosphate in adipose tissue |
| Km_tca_Adp_A | Michaelis constant of TCA cycle for Adp in adipose tissue |
| Km_tca_Pyr_A | Michaelis constant of TCA cycle for Pyr in adipose tissue |
| Km_tca_Nad_A | Michaelis constant of TCA cycle for Nad in adipose tissue |
| Km_tca_Fad_A | Michaelis constant of TCA cycle for Fad in adipose tissue |
| Vdif_ffat_FFA_B_A | rate constant of FFA transporter in adipose tissue |
| Kdif_ffat_FFA_B_A | dissociation constant of FFA transporter for FFA_B in adipose tissue |
| Kdif_ffat_FFA_A | dissociation constant of FFA transporter in adipose tissue |
| Vmax_ffat_FFA_B_A | rate constant of active FFA transporter in adipose tissue |
| Km_ffat_FFA_B_A | Michaelis constant of active FFA transporter in adipose tissue |
| Vmax_tgsyn_FFA_A | rate constant of TG synthesis for FFA in adipose tissue |
| Km_tgsyn_FFA_A | Michaelis constant of TG synthesis for FFA in adipose tissue |
| Km_tgsyn_Glycp_A | Michaelis constant of TG synthesis for Glycp in adipose tissue |
| Km_tgsyn_Atp_A | Michaelis constant of TG synthesis for Atp in adipose tissue |
| TG_A_max | maximum TG concentration in adipose tissue |
| Km_tgsyn_TG_A | Michaelis constant of TG synthesis for TG in adipose tissue |
| Vmax_tgdeg_TG_A | rate constant of TG degradation for TG in adipose tissue |
| Km_tgdeg_TG_A | Michaelis constant of TG degradation for TG in adipose tissue |
| Vdif_glyct_Glyc_B_A | rate constant of glycerol transporter in adipose tissue |
| Kdif_glyct_Glyc_B_A | dissociation constant of glycerol transporter for Glyc_B in adipose tissue |
| Kdif_glyct_Glyc_A | dissociation constant of glycerol transporter in adipose tissue |
| Vmax_glyk_Glyc_A | rate constant of glycerol kinase for Glyc in adipose tissue |
| Km_glyk_Glyc_A | Michaelis constant of glycerol kinase for Glyc in adipose tissue |
| Km_glyk_Atp_A | Michaelis constant of glycerol kinase for Atp in adipose tissue |
| Ki_glyk_Gap_A | inhibition constant of glycerol kinase for Gap in adipose tissue |
| Vdif_tgt_TG_B_A | rate constant of TG transporter in adipose tissue |
| Kdif_tgt_TG_B_A | dissociation constant of TG transporter in adipose tissue |
| Keq_tgt_TG_A | equilibrium constant of TG transporter in adipose tissue |
| Vmax_tgt_TG_B_A | rate constant of active TG transporter in adipose tissue |
| Km_tgt_TG_A | Michaelis constant of active TG transporter in adipose tissue |

| | |
|---|---|
| Vmax_g3pd_Gap_A | rate constant of glyceraldehyde-3-phosphate dehydrogenase for Gap in adipose tissue |
| Km_g3pd_Gap_A | Michaelis constant of glyceraldehyde-3-phosphate dehydrogenase for Gap in adipose tissue |
| Km_g3pd_Glycp_A | Michaelis constant of glyceraldehyde-3-phosphate dehydrogenase for Glycp in adipose tissue |
| Km_g3pd_Nadh_A | Michaelis constant of glyceraldehyde-3-phosphate dehydrogenase for Nadh in adipose tissue |
| Km_g3pd_Nad_A | Michaelis constant of glyceraldehyde-3-phosphate dehydrogenase for Nad in adipose tissue |
| Keq_g3pd_Glycp_A | equilibrium constant of glyceraldehyde-3-phosphate dehydrogenase for Glycp in adipose tissue |
| Vmax_boxid_FFA_A | rate constant of beta-oxidation for FFA in adipose tissue |
| Km_boxid_FFA_A | Michaelis constant of beta-oxidation for FFA in adipose tissue |
| Km_boxid_Atp_A | Michaelis constant of beta-oxidation for Atp in adipose tissue |
| Ki_boxid_Accoa_AM | inhibition constant of beta-oxidation for mitochondrial Accoa in adipose tissue |
| Ki_boxid_Malcoa_A | inhibition constant of beta-oxidation for Malcoa in adipose tissue |
| Km_boxid_Nad_A | Michaelis constant of beta-oxidation for Nad in adipose tissue |
| Km_boxid_Fad_A | Michaelis constant of beta-oxidation for Fad in adipose tissue |
| Vmax_accoat_Accoa_AM_AC | rate constant of mitochondrial Accoa transport to cytoplasm in adipose tissue |
| Km_accoat_Accoa_AM | Michaelis constant of mitochondrial Accoa transporter to cytoplasm in adipose tissue |
| Km_accoat_Atp_A | Michaelis constant of mitochondrial Accoa transporter to cytoplasm for Atp in adipose tissue |
| Km_accoat_Pyr_A | activation constant of mitochondrial Accoa transport to cytoplasm for Pyr in adipose tissue |
| Vmax_bhbsyn_Accoa_AM | rate constant of Bhb synthesis for mitochondrial Accoa in adipose tissue |
| Km_bhbsyn_Accoa_AM | Michaelis constant of Bhb synthesis for mitochondrial Accoa in adipose tissue |
| Km_bhbsyn_Nadh_A | Michaelis constant of Bhb synthesis for Nadh in adipose tissue |
| n_bhbsyn_Pyr_A | Hill coefficient of Bhb synthesis for Pyr in adipose tissue |
| Ki_bhbsyn_Pyr_A | inhibition constant of Bhb synthesis for Pyr in adipose tissue |
| Vmax_bhbdeg_Bhb_A | rate constant of Bhb degradation for Bhb in adipose tissue |
| Km_bhbdeg_Bhb_A | Michaelis constant of Bhb degradation for Bhb in adipose tissue |
| Km_bhbdeg_Atp_A | Michaelis constant of Bhb degradation for Atp in adipose tissue |
| Km_bhbdeg_Nad_A | Michaelis constant of Bhb degradation for Nad in adipose tissue |
| Vdif_bhbt_Bhb_B_A | rate constant of Bhb transporter in adipose tissue |
| Kdif_bhbt_Bhb_B_A | dissociation constant of Bhb transporter for Bhb_B in adipose tissue |
| Kdif_bhbt_Bhb_A | dissociation constant of Bhb transporter in adipose tissue |

| | |
|---|---|
| Vmax_lipog1_Accoa_AC | rate constant of lipogenesis 1 for Accoa in adipose tissue |
| Km_lipog1_Accoa_AC | Michaelis constant of lipogenesis 1 for Accoa in adipose tissue |
| Km_lipog1_Atp_A | Michaelis constant of lipogenesis 1 for Atp in adipose tissue |
| Vmax_lipog2_Malcoa_A | rate constant of lipogenesis 2 for Malcoa in adipose tissue |
| Km_lipog2_Malcoa_A | Michaelis constant of lipogenesis 2 for Malcoa in adipose tissue |
| Km_lipog2_Adp_A | Michaelis constant of lipogenesis 2 for Adp in adipose tissue |
| Km_lipog2_Nadph_A | Michaelis constant of lipogenesis 2 for Nadph in adipose tissue |
| Vmax_cholsyn1_Accoa_AC | rate constant of cholesterol synthesis 1 for Accoa in adipose tissue |
| Km_cholsyn1_Accoa_AC | Michaelis constant of cholesterol synthesis 1 for Accoa in adipose tissue |
| Vmax_cholsyn2_Hmgcoa_A | rate constant of cholesterol synthesis 2 for Hmgcoa in adipose tissue |
| Km_cholsyn2_Hmgcoa_A | Michaelis constant of cholesterol synthesis 2 for Hmgcoa in adipose tissue |
| Km_cholsyn2_Atp_A | Michaelis constant of cholesterol synthesis 2 for Atp in adipose tissue |
| Km_cholsyn2_Fadh_A | Michaelis constant of cholesterol synthesis 2 for Fadh in adipose tissue |
| Km_cholsyn2_Nadph_A | Michaelis constant of cholesterol synthesis 2 for Nadph in adipose tissue |
| Vmax_cholt_Chol_A | rate constant of active cholesterol transporter in adipose tissue |
| Km_cholt_Chol_A | Michaelis constant of active cholesterol transporter in adipose tissue |
| Vmax_atpsynf_Fadh_A | rate constant of Atp synthesis from Fadh in adipose tissue |
| Km_atpsynf_Fadh_A | Michaelis constant of Atp synthesis from Fadh for Fadh in adipose tissue |
| Km_atpsynf_Adp_A | Michaelis constant of Atp synthesis from Fadh for Adp in adipose tissue |
| Vmax_atpsynn_Nadh_A | rate constant of Atp synthesis from Nadh in adipose tissue |
| Km_atpsynn_Nadh_A | Michaelis constant of Atp synthesis from Nadh for Nadh in adipose tissue |
| Km_atpsynn_Adp_A | Michaelis constant of Atp synthesis from Nadh for Adp in adipose tissue |
| Vmax_atpuse_Atp_A | rate constant of Atp utilization for Atp in adipose tissue |
| Km_atpuse_Atp_A | Michaelis constant of Atp utilization for Atp in adipose tissue |
| Vmax_ampreg_Amp_A | rate constant of Amp regeneration for Amp in adipose tissue |
| Km_ampreg_Amp_A | Michaelis constant of Amp regeneration for Amp in adipose tissue |
| Km_ampreg_Atp_A | Michaelis constant of Amp regeneration for Atp in adipose tissue |
| Km_ampreg_Adp_A | Michaelis constant of Amp regeneration for Adp in adipose tissue |
| Vmax_nadhk_Nadh_A | rate constant of Nadh kinase for Nadh in adipose tissue |
| Km_nadhk_Nadh_A | Michaelis constant of Nadh kinase for Nadh in adipose tissue |
| Km_nadhk_Atp_A | Michaelis constant of Nadh kinase for Atp in adipose tissue |
| Vmax_nadhuse_Nadh_A | rate constant of Nadh utilization for Nadh in adipose tissue |
| Km_nadhuse_Nadh_A | Michaelis constant of Nadh utilization for Nadh in adipose tissue |
| Vmax_gdpreg_Gdp_A | rate constant of Gdp regeneration for Gdp in adipose tissue |
| Km_gdpreg_Gdp_A | Michaelis constant of Gdp regeneration for Gdp in adipose tissue |
| Km_gdpreg_Atp_A | Michaelis constant of Gdp regeneration for Atp in adipose tissue |

| | |
|---|---|
| Km_gdpreg_Gtp_A | Michaelis constant of Gdp regeneration for Gtp in adipose tissue |
| Km_gdpreg_Adp_A | Michaelis constant of Gdp regeneration for Adp in adipose tissue |
| Keq_gdpreg_Gtp_A | equilibrium constant of Gdp regeneration for Gtp in adipose tissue |
| Vmax_gtpuse_Gtp_A | rate constant of Gtp utilization for Gtp in adipose tissue |
| Km_gtpuse_Gtp_A | Michaelis constant of Gtp utilization for Gtp in adipose tissue |
| Vmax_udpreg_Udp_A | rate constant of Udp regeneration for Udp in adipose tissue |
| Km_udpreg_Udp_A | Michaelis constant of Udp regeneration for Udp in adipose tissue |
| Km_udpreg_Atp_A | Michaelis constant of Udp regeneration for Atp in adipose tissue |
| Km_udpreg_Utp_A | Michaelis constant of Udp regeneration for Utp in adipose tissue |
| Km_udpreg_Adp_A | Michaelis constant of Udp regeneration for Adp in adipose tissue |
| Keq_udpreg_Utp_A | equilibrium constant of Udp regeneration for Utp in adipose tissue |
| Vmax_utpuse_Utp_A | rate constant of Utp utilization for Utp in adipose tissue |
| Km_utpuse_Utp_A | Michaelis constant of Utp utilization for Utp in adipose tissue |
| Vmax_nadphuse_Nadph_A | rate constant of Nadph utilization for Nadph in adipose tissue |
| Km_nadphuse_Nadph_A | Michaelis constant of Nadph utilization for Nadph in adipose tissue |
| Vmax_fadhuse_Fadh_A | rate constant of Fadh utilization for Fadh in adipose tissue |
| Km_fadhuse_Fadh_A | Michaelis constant of Fadh utilization for Fadh in adipose tissue |
| Vmax_ck_Cre_A | rate constant of creatine kinase for Cre in adipose tissue |
| Km_ck_Cre_A | Michaelis constant of creatine kinase for Cre in adipose tissue |
| Km_ck_Atp_A | Michaelis constant of creatine kinase for Atp in adipose tissue |
| Km_ck_Crep_A | Michaelis constant of creatine kinase for Crep in adipose tissue |
| Km_ck_Adp_A | Michaelis constant of creatine kinase for Adp in adipose tissue |
| Keq_ck_Crep_A | equilibrium constant of creatine kinase for Crep in adipose tissue |
| Phos_A | phosphate in adipose tissue |
| alpha_base_G | insulin base factor in GI |
| alpha_band_G | insulin allowable width in GI |
| Km_Ins_B_G | Michaelis constant for insulin in GI |
| n_Ins_B_G | Hill coefficient for insulin in GI |
| Vdif_glut2_Glc_B_G | rate constant of glucose transporter type 2 in GI |
| Kdif_glut2_Glc_B_G | dissociation constant of glucose transporter type 2 for Glc_B in GI |
| Kdif_glut2_Glc_G | dissociation constant of glucose transporter type 2 in GI |
| Vmax_hk_Glc_G | reaction rate constant of hexokinase for Glc in GI |
| Km_hk_Glc_G | Michaelis constant of hexokinase for Glc in GI |
| Ki_hk_G6p_G | inhibition constant of hexokinase for Glc in GI |
| Km_hk_Atp_G | Michaelis constant of hexokinase for Atp in GI |
| Ki_hk_Atp_G | inhibition constant of hexokinase for Atp in GI |

| | |
|---|---|
| Vmax_ppp_G6p_G | reaction rate constant of pentose phosphate pathway for G6p in GI |
| Km_ppp_G6p_G | Michaelis constant of pentose phosphate pathway for G6p in GI |
| Km_ppp_Nadp_G | Michaelis constant of pentose phosphate pathway for Nadp in GI |
| Vmax_g6pase_G6p_G | reaction rate constant of G6Pase for G6p in GI |
| Km_g6pase_G6p_G | Michaelis constant of G6Pase for G6p in GI |
| Vmax_gs_G6p_G | reaction rate constant of glycogen synthase for G6p in GI |
| Km_gs_G6p_G | Michaelis constant of glycogen synthase for G6p in GI |
| n_gs_G6p_G | Hill coefficient of glycogen synthase for G6p in GI |
| Km_gs_Utp_G | Michaelis constant of glycogen synthase for G6p in GI |
| Glygn_G_max | maximum glycogen concentration in GI |
| Km_gs_Glygn_G | Michaelis constant of glycogen synthase for glycogen in GI |
| Vmax_gd_Glygn_G | reaction rate constant of glycogen degradation for glycogen in GI |
| Km_gd_Glygn_G | Michaelis constant of glycogen degradation for glycogen in GI |
| Km_gd_Phos_G | Michaelis constant of glycogen degradation for phosphate in GI |
| Vmax_pfk_G6p_G | reaction rate constant of phosphofructokinase for G6p in GI |
| Km_pfk_G6p_G | Michaelis constant of phosphofructokinase for G6p in GI |
| Km_pfk_Atp_G | Michaelis constant of phosphofructokinase for Atp in GI |
| Ki_pfk_Atp_G | inhibition constant of phosphofructokinase for Atp in GI |
| Km_pfk_Adp_G | Michaelis constant of phosphofructokinase for Adp in GI |
| Ki_pfk_Gap_G | inhibition constant of phosphofructokinase for Gap in GI |
| b_pfk_Gap_G | coefficient of phosphofructokinase for Gap in GI |
| Vmax_fbp_Gap_G | reaction rate constant of fructose bisphosphatase for Gap in GI |
| Km_fbp_Gap_G | Michaelis constant of fructose bisphosphatase for Gap in GI |
| Vmax_pk_Gap_G | reaction rate constant of pyruvate kinase for Gap in GI |
| Km_pk_Gap_G | Michaelis constant of pyruvate kinase for Gap in GI |
| b_pk_Accoa_GM | coefficient of pyruvate kinase for mitochondrial Accoa in GI |
| Ki_pk_Accoa_GM | inhibition constant of pyruvate kinase for mitochondrial Accoa in GI |
| Km_pk_Adp_G | Michaelis constant of pyruvate kinase for Adp in GI |
| Vmax_pepck_Pyr_G | reaction rate constant of phosphoenolpyruvate carboxykinase for Pyr in GI |
| Km_pepck_Pyr_G | Michaelis constant of phosphoenolpyruvate carboxykinase for Pyr in GI |
| Km_pepck_Atp_G | Michaelis constant of phosphoenolpyruvate carboxykinase for Atp in GI |
| Km_pepck_Gtp_G | Michaelis constant of phosphoenolpyruvate carboxykinase for Gtp in GI |
| Vdif_pyrt_Pyr_B_G | rate constant of pyruvate transporter in GI |
| Kdif_pyrt_Pyr_B_G | dissociation constant of pyruvate transporter for Pyr_B in GI |
| Kdif_pyrt_Pyr_G | dissociation constant of pyruvate transporter in GI |
| Vdif_lact_Lac_B_G | rate constant of lactate transporter in GI |

| | |
|---|---|
| Kdif_lact_Lac_G | dissociation constant of lactate transporter in GI |
| Kdif_lact_Lac_B_G | dissociation constant of lactate transporter for Lac_B in GI |
| Vmax_ldh_Pyr_G | rate constant of lactate dehydrogenase for Pyr in GI |
| Km_ldh_Pyr_G | Michaelis constant of lactate dehydrogenase for Pyr in GI |
| Km_ldh_Nadh_G | Michaelis constant of lactate dehydrogenase for Nadh in GI |
| Km_ldh_Lac_G | Michaelis constant of lactate dehydrogenase for Lac in GI |
| Km_ldh_Nad_G | Michaelis constant of lactate dehydrogenase for Nad in GI |
| Keq_ldh_Lac_G | equilibrium constant of lactate dehydrogenase for Lac in GI |
| Vdif_alat_Ala_B_G | rate constant of alanine transporter in GI |
| Kdif_alat_Ala_G | dissociation constant of alanine transporter in GI |
| Kdif_alat_Ala_B_G | dissociation constant of alanine transporter for Ala_B in GI |
| Vmax_alata_Pyr_G | rate constant of alanine transporter for Pyr in GI |
| Km_alata_Pyr_G | Michaelis constant of alanine aminotransferase for Pyr in GI |
| Km_alata_Ala_G | Michaelis constant of alanine aminotransferase for Ala in GI |
| Keq_alata_Ala_G | equilibrium constant of alanine aminotransferase for Ala in GI |
| Vmax_pdh_Pyr_G | rate constant of pyruvate dehydrogenase for Pyr in GI |
| Km_pdh_Pyr_G | Michaelis constant of pyruvate dehydrogenase for Pyr in GI |
| Ki_pdh_Accoa_GM | inhibition constant of pyruvate dehydrogenase for mitochondrial Accoa in GI |
| Km_pdh_Nad_G | Michaelis constant of pyruvate dehydrogenase for Nad in GI |
| Vmax_tca_Accoa_GM | rate constant of TCA cycle for mitochondrial Accoa in GI |
| Km_tca_Accoa_GM | Michaelis constant of TCA cycle for mitochondrial Accoa in GI |
| Km_tca_Phos_G | Michaelis constant of TCA cycle for phosphate in GI |
| Km_tca_Adp_G | Michaelis constant of TCA cycle for Adp in GI |
| Km_tca_Pyr_G | Michaelis constant of TCA cycle for Pyr in GI |
| Km_tca_Nad_G | Michaelis constant of TCA cycle for Nad in GI |
| Km_tca_Fad_G | Michaelis constant of TCA cycle for Fad in GI |
| Vdif_ffat_FFA_B_G | rate constant of FFA transporter in GI |
| Kdif_ffat_FFA_B_G | dissociation constant of FFA transporter for FFA_B in GI |
| Kdif_ffat_FFA_G | dissociation constant of FFA transporter in GI |
| Vmax_ffat_FFA_B_G | rate constant of active FFA transporter in GI |
| Km_ffat_FFA_B_G | Michaelis constant of active FFA transporter in GI |
| Vmax_tgsyn_FFA_G | rate constant of TG synthesis for FFA in GI |
| Km_tgsyn_FFA_G | Michaelis constant of TG synthesis for FFA in GI |
| Km_tgsyn_Glycp_G | Michaelis constant of TG synthesis for Glycp in GI |
| Km_tgsyn_Atp_G | Michaelis constant of TG synthesis for Atp in GI |
| TG_G_max | maximum TG concentration in GI |

| | |
|---|---|
| Km_tgsyn_TG_G | Michaelis constant of TG synthesis for TG in GI |
| Vmax_tgdeg_TG_G | rate constant of TG degradation for TG in GI |
| Km_tgdeg_TG_G | Michaelis constant of TG degradation for TG in GI |
| Vdif_glyct_Glyc_B_G | rate constant of glycerol transporter in GI |
| Kdif_glyct_Glyc_B_G | dissociation constant of glycerol transporter for Glyc_B in GI |
| Kdif_glyct_Glyc_G | dissociation constant of glycerol transporter in GI |
| Vmax_glyk_Glyc_G | rate constant of glycerol kinase for Glyc in GI |
| Km_glyk_Glyc_G | Michaelis constant of glycerol kinase for Glyc in GI |
| Km_glyk_Atp_G | Michaelis constant of glycerol kinase for Atp in GI |
| Ki_glyk_Gap_G | inhibition constant of glycerol kinase for Gap in GI |
| Vdif_tgt_TG_B_G | rate constant of TG transporter in GI |
| Kdif_tgt_TG_B_G | dissociation constant of TG transporter in GI |
| Keq_tgt_TG_G | equilibrium constant of TG transporter in GI |
| Vmax_tgt_TG_B_G | rate constant of active TG transporter in GI |
| Km_tgt_TG_G | Michaelis constant of active TG transporter in GI |
| Vmax_g3pd_Gap_G | rate constant of glyceraldehyde-3-phosphate dehydrogenase for Gap in GI |
| Km_g3pd_Gap_G | Michaelis constant of glyceraldehyde-3-phosphate dehydrogenase for Gap in GI |
| Km_g3pd_Glycp_G | Michaelis constant of glyceraldehyde-3-phosphate dehydrogenase for Glycp in GI |
| Km_g3pd_Nadh_G | Michaelis constant of glyceraldehyde-3-phosphate dehydrogenase for Nadh in GI |
| Km_g3pd_Nad_G | Michaelis constant of glyceraldehyde-3-phosphate dehydrogenase for Nad in GI |
| Keq_g3pd_Glycp_G | equilibrium constant of glyceraldehyde-3-phosphate dehydrogenase for Glycp in GI |
| Vmax_boxid_FFA_G | rate constant of beta-oxidation for FFA in GI |
| Km_boxid_FFA_G | Michaelis constant of beta-oxidation for FFA in GI |
| Km_boxid_Atp_G | Michaelis constant of beta-oxidation for Atp in GI |
| Ki_boxid_Accoa_GM | inhibition constant of beta-oxidation for mitochondrial Accoa in GI |
| Ki_boxid_Malcoa_G | inhibition constant of beta-oxidation for Malcoa in GI |
| Km_boxid_Nad_G | Michaelis constant of beta-oxidation for Nad in GI |
| Km_boxid_Fad_G | Michaelis constant of beta-oxidation for Fad in GI |
| Vmax_accoat_Accoa_GM_GC | rate constant of mitochondrial Accoa transport to cytoplasm in GI |
| Km_accoat_Accoa_GM | Michaelis constant of mitochondrial Accoa transporter to cytoplasm in GI |
| Km_accoat_Atp_G | Michaelis constant of mitochondrial Accoa transporter to cytoplasm for Atp in GI |
| Km_accoat_Pyr_G | activation constant of mitochondrial Accoa transport to cytoplasm for Pyr in GI |
| Vmax_bhbsyn_Accoa_GM | rate constant of Bhb synthesis for mitochondrial Accoa in GI |
| Km_bhbsyn_Accoa_GM | Michaelis constant of Bhb synthesis for mitochondrial Accoa in GI |
| Km_bhbsyn_Nadh_G | Michaelis constant of Bhb synthesis for Nadh in GI |
| n_bhbsyn_Pyr_G | Hill coefficient of Bhb synthesis for Pyr in GI |

| | |
|---|---|
| Ki_bhbsyn_Pyr_G | inhibition constant of Bhb synthesis for Pyr in GI |
| Vmax_bhbdeg_Bhb_G | rate constant of Bhb degradation for Bhb in GI |
| Km_bhbdeg_Bhb_G | Michaelis constant of Bhb degradation for Bhb in GI |
| Km_bhbdeg_Atp_G | Michaelis constant of Bhb degradation for Atp in GI |
| Km_bhbdeg_Nad_G | Michaelis constant of Bhb degradation for Nad in GI |
| Vdif_bhbt_Bhb_B_G | rate constant of Bhb transporter in GI |
| Kdif_bhbt_Bhb_B_G | dissociation constant of Bhb transporter for Bhb_B in GI |
| Kdif_bhbt_Bhb_G | dissociation constant of Bhb transporter in GI |
| Vmax_lipog1_Accoa_GC | rate constant of lipogenesis 1 for Accoa in GI |
| Km_lipog1_Accoa_GC | Michaelis constant of lipogenesis 1 for Accoa in GI |
| Km_lipog1_Atp_G | Michaelis constant of lipogenesis 1 for Atp in GI |
| Vmax_lipog2_Malcoa_G | rate constant of lipogenesis 2 for Malcoa in GI |
| Km_lipog2_Malcoa_G | Michaelis constant of lipogenesis 2 for Malcoa in GI |
| Km_lipog2_Adp_G | Michaelis constant of lipogenesis 2 for Adp in GI |
| Km_lipog2_Nadph_G | Michaelis constant of lipogenesis 2 for Nadph in GI |
| Vmax_cholsyn1_Accoa_GC | rate constant of cholesterol synthesis 1 for Accoa in GI |
| Km_cholsyn1_Accoa_GC | Michaelis constant of cholesterol synthesis 1 for Accoa in GI |
| Vmax_cholsyn2_Hmgcoa_G | rate constant of cholesterol synthesis 2 for Hmgcoa in GI |
| Km_cholsyn2_Hmgcoa_G | Michaelis constant of cholesterol synthesis 2 for Hmgcoa in GI |
| Km_cholsyn2_Atp_G | Michaelis constant of cholesterol synthesis 2 for Atp in GI |
| Km_cholsyn2_Fadh_G | Michaelis constant of cholesterol synthesis 2 for Fadh in GI |
| Km_cholsyn2_Nadph_G | Michaelis constant of cholesterol synthesis 2 for Nadph in GI |
| Vmax_cholt_Chol_G | rate constant of active cholesterol transporter in GI |
| Km_cholt_Chol_G | Michaelis constant of active cholesterol transporter in GI |
| Vmax_atpsynf_Fadh_G | rate constant of Atp synthesis from Fadh in GI |
| Km_atpsynf_Fadh_G | Michaelis constant of Atp synthesis from Fadh for Fadh in GI |
| Km_atpsynf_Adp_G | Michaelis constant of Atp synthesis from Fadh for Adp in GI |
| Vmax_atpsynn_Nadh_G | rate constant of Atp synthesis from Nadh in GI |
| Km_atpsynn_Nadh_G | Michaelis constant of Atp synthesis from Nadh for Nadh in GI |
| Km_atpsynn_Adp_G | Michaelis constant of Atp synthesis from Nadh for Adp in GI |
| Vmax_atpuse_Atp_G | rate constant of Atp utilization for Atp in GI |
| Km_atpuse_Atp_G | Michaelis constant of Atp utilization for Atp in GI |
| Vmax_ampreg_Amp_G | rate constant of Amp regeneration for Amp in GI |
| Km_ampreg_Amp_G | Michaelis constant of Amp regeneration for Amp in GI |
| Km_ampreg_Atp_G | Michaelis constant of Amp regeneration for Atp in GI |
| Km_ampreg_Adp_G | Michaelis constant of Amp regeneration for Adp in GI |

| | |
|---|---|
| Vmax_nadhk_Nadh_G | rate constant of Nadh kinase for Nadh in GI |
| Km_nadhk_Nadh_G | Michaelis constant of Nadh kinase for Nadh in GI |
| Km_nadhk_Atp_A | Michaelis constant of Nadh kinase for Atp in GI |
| Vmax_nadhuse_Nadh_G | rate constant of Nadh utilization for Nadh in GI |
| Km_nadhuse_Nadh_G | Michaelis constant of Nadh utilization for Nadh in GI |
| Vmax_gdpreg_Gdp_G | rate constant of Gdp regeneration for Gdp in GI |
| Km_gdpreg_Gdp_G | Michaelis constant of Gdp regeneration for Gdp in GI |
| Km_gdpreg_Atp_G | Michaelis constant of Gdp regeneration for Atp in GI |
| Km_gdpreg_Gtp_G | Michaelis constant of Gdp regeneration for Gtp in GI |
| Km_gdpreg_Adp_G | Michaelis constant of Gdp regeneration for Adp in GI |
| Keq_gdpreg_Gtp_G | equilibrium constant of Gdp regeneration for Gtp in GI |
| Vmax_gtpuse_Gtp_G | rate constant of Gtp utilization for Gtp in GI |
| Km_gtpuse_Gtp_G | Michaelis constant of Gtp utilization for Gtp in GI |
| Vmax_udpreg_Udp_G | rate constant of Udp regeneration for Udp in GI |
| Km_udpreg_Udp_G | Michaelis constant of Udp regeneration for Udp in GI |
| Km_udpreg_Atp_G | Michaelis constant of Udp regeneration for Atp in GI |
| Km_udpreg_Utp_G | Michaelis constant of Udp regeneration for Utp in GI |
| Km_udpreg_Adp_G | Michaelis constant of Udp regeneration for Adp in GI |
| Keq_udpreg_Utp_G | equilibrium constant of Udp regeneration for Utp in GI |
| Vmax_utpuse_Utp_G | rate constant of Utp utilization for Utp in GI |
| Km_utpuse_Utp_G | Michaelis constant of Utp utilization for Utp in GI |
| Vmax_nadphuse_Nadph_G | rate constant of Nadph utilization for Nadph in GI |
| Km_nadphuse_Nadph_G | Michaelis constant of Nadph utilization for Nadph in GI |
| Vmax_fadhuse_Fadh_G | rate constant of Fadh utilization for Fadh in GI |
| Km_fadhuse_Fadh_G | Michaelis constant of Fadh utilization for Fadh in GI |
| Vmax_ck_Cre_G | rate constant of creatine kinase for Cre in GI |
| Km_ck_Cre_G | Michaelis constant of creatine kinase for Cre in GI |
| Km_ck_Atp_G | Michaelis constant of creatine kinase for Atp in GI |
| Km_ck_Crep_G | Michaelis constant of creatine kinase for Crep in GI |
| Km_ck_Adp_G | Michaelis constant of creatine kinase for Adp in GI |
| Keq_ck_Crep_G | equilibrium constant of creatine kinase for Crep in GI |
| Phos_G | phosphate in GI |
| alpha_base_H | insulin base factor in heart |
| alpha_band_H | insulin allowable width in heart |
| Km_Ins_B_H | Michaelis constant for insulin in heart |
| n_Ins_B_H | Hill coefficient for insulin in heart |

| | |
|---|---|
| Vdif_glut2_Glc_B_H | rate constant of glucose transporter type 2 in heart |
| Kdif_glut2_Glc_B_H | dissociation constant of glucose transporter type 2 for Glc_B in heart |
| Kdif_glut2_Glc_H | dissociation constant of glucose transporter type 2 in heart |
| Vmax_hk_Glc_H | reaction rate constant of hexokinase for Glc in heart |
| Km_hk_Glc_H | Michaelis constant of hexokinase for Glc in heart |
| Ki_hk_G6p_H | inhibition constant of hexokinase for Glc in heart |
| Km_hk_Atp_H | Michaelis constant of hexokinase for Atp in heart |
| Ki_hk_Atp_H | inhibition constant of hexokinase for Atp in heart |
| Vmax_ppp_G6p_H | reaction rate constant of pentose phosphate pathway for G6p in heart |
| Km_ppp_G6p_H | Michaelis constant of pentose phosphate pathway for G6p in heart |
| Km_ppp_Nadp_H | Michaelis constant of pentose phosphate pathway for Nadp in heart |
| Vmax_g6pase_G6p_H | reaction rate constant of G6Pase for G6p in heart |
| Km_g6pase_G6p_H | Michaelis constant of G6Pase for G6p in heart |
| Vmax_gs_G6p_H | reaction rate constant of glycogen synthase for G6p in heart |
| Km_gs_G6p_H | Michaelis constant of glycogen synthase for G6p in heart |
| n_gs_G6p_H | Hill coefficient of glycogen synthase for G6p in heart |
| Km_gs_Utp_H | Michaelis constant of glycogen synthase for G6p in heart |
| Glygn_H_max | maximum glycogen concentration in heart |
| Km_gs_Glygn_H | Michaelis constant of glycogen synthase for glycogen in heart |
| Vmax_gd_Glygn_H | reaction rate constant of glycogen degradation for glycogen in heart |
| Km_gd_Glygn_H | Michaelis constant of glycogen degradation for glycogen in heart |
| Km_gd_Phos_H | Michaelis constant of glycogen degradation for phosphate in heart |
| Vmax_pfk_G6p_H | reaction rate constant of phosphofructokinase for G6p in heart |
| Km_pfk_G6p_H | Michaelis constant of phosphofructokinase for G6p in heart |
| Km_pfk_Atp_H | Michaelis constant of phosphofructokinase for Atp in heart |
| Ki_pfk_Atp_H | inhibition constant of phosphofructokinase for Atp in heart |
| Km_pfk_Adp_H | Michaelis constant of phosphofructokinase for Adp in heart |
| Ki_pfk_Gap_H | inhibition constant of phosphofructokinase for Gap in heart |
| b_pfk_Gap_H | coefficient of phosphofructokinase for Gap in heart |
| Vmax_fbp_Gap_H | reaction rate constant of fructose bisphosphatase for Gap in heart |
| Km_fbp_Gap_H | Michaelis constant of fructose bisphosphatase for Gap in heart |
| Vmax_pk_Gap_H | reaction rate constant of pyruvate kinase for Gap in heart |
| Km_pk_Gap_H | Michaelis constant of pyruvate kinase for Gap in heart |
| b_pk_Accoa_HM | coefficient of pyruvate kinase for mitochondrial Accoa in heart |
| Ki_pk_Accoa_HM | inhibition constant of pyruvate kinase for mitochondrial Accoa in heart |
| Km_pk_Adp_H | Michaelis constant of pyruvate kinase for Adp in heart |

| | |
|---|---|
| Vmax_pepck_Pyr_H | reaction rate constant of phosphoenolpyruvate carboxykinase for Pyr in heart |
| Km_pepck_Pyr_H | Michaelis constant of phosphoenolpyruvate carboxykinase for Pyr in heart |
| Km_pepck_Atp_H | Michaelis constant of phosphoenolpyruvate carboxykinase for Atp in heart |
| Km_pepck_Gtp_H | Michaelis constant of phosphoenolpyruvate carboxykinase for Gtp in heart |
| Vdif_pyrt_Pyr_B_H | rate constant of pyruvate transporter in heart |
| Kdif_pyrt_Pyr_B_H | dissociation constant of pyruvate transporter for Pyr_B in heart |
| Kdif_pyrt_Pyr_H | dissociation constant of pyruvate transporter in heart |
| Vdif_lact_Lac_B_H | rate constant of lactate transporter in heart |
| Kdif_lact_Lac_H | dissociation constant of lactate transporter in heart |
| Kdif_lact_Lac_B_H | dissociation constant of lactate transporter for Lac_B in heart |
| Vmax_ldh_Pyr_H | rate constant of lactate dehydrogenase for Pyr in heart |
| Km_ldh_Pyr_H | Michaelis constant of lactate dehydrogenase for Pyr in heart |
| Km_ldh_Nadh_H | Michaelis constant of lactate dehydrogenase for Nadh in heart |
| Km_ldh_Lac_H | Michaelis constant of lactate dehydrogenase for Lac in heart |
| Km_ldh_Nad_H | Michaelis constant of lactate dehydrogenase for Nad in heart |
| Keq_ldh_Lac_H | equilibrium constant of lactate dehydrogenase for Lac in heart |
| Vdif_alat_Ala_B_H | rate constant of alanine transporter in heart |
| Kdif_alat_Ala_H | dissociation constant of alanine transporter in heart |
| Kdif_alat_Ala_B_H | dissociation constant of alanine transporter for Ala_B in heart |
| Vmax_alata_Pyr_H | rate constant of alanine transporter for Pyr in heart |
| Km_alata_Pyr_H | Michaelis constant of alanine aminotransferase for Pyr in heart |
| Km_alata_Ala_H | Michaelis constant of alanine aminotransferase for Ala in heart |
| Keq_alata_Ala_H | equilibrium constant of alanine aminotransferase for Ala in heart |
| Vmax_pdh_Pyr_H | rate constant of pyruvate dehydrogenase for Pyr in heart |
| Km_pdh_Pyr_H | Michaelis constant of pyruvate dehydrogenase for Pyr in heart |
| Ki_pdh_Accoa_HM | inhibition constant of pyruvate dehydrogenase for mitochondrial Accoa in heart |
| Km_pdh_Nad_H | Michaelis constant of pyruvate dehydrogenase for Nad in heart |
| Vmax_tca_Accoa_HM | rate constant of TCA cycle for mitochondrial Accoa in heart |
| Km_tca_Accoa_HM | Michaelis constant of TCA cycle for mitochondrial Accoa in heart |
| Km_tca_Phos_H | Michaelis constant of TCA cycle for phosphate in heart |
| Km_tca_Adp_H | Michaelis constant of TCA cycle for Adp in heart |
| Km_tca_Pyr_H | Michaelis constant of TCA cycle for Pyr in heart |
| Km_tca_Nad_H | Michaelis constant of TCA cycle for Nad in heart |
| Km_tca_Fad_H | Michaelis constant of TCA cycle for Fad in heart |
| Vdif_ffat_FFA_B_H | rate constant of FFA transporter in heart |
| Kdif_ffat_FFA_B_H | dissociation constant of FFA transporter for FFA_B in heart |

| | |
|---|---|
| Kdif_ffat_FFA_H | dissociation constant of FFA transporter in heart |
| Vmax_ffat_FFA_B_H | rate constant of active FFA transporter in heart |
| Km_ffat_FFA_B_H | Michaelis constant of active FFA transporter in heart |
| Vmax_tgsyn_FFA_H | rate constant of TG synthesis for FFA in heart |
| Km_tgsyn_FFA_H | Michaelis constant of TG synthesis for FFA in heart |
| Km_tgsyn_Glycp_H | Michaelis constant of TG synthesis for Glycp in heart |
| Km_tgsyn_Atp_H | Michaelis constant of TG synthesis for Atp in heart |
| TG_H_max | maximum TG concentration in heart |
| Km_tgsyn_TG_H | Michaelis constant of TG synthesis for TG in heart |
| Vmax_tgdeg_TG_H | rate constant of TG degradation for TG in heart |
| Km_tgdeg_TG_H | Michaelis constant of TG degradation for TG in heart |
| Vdif_glyct_Glyc_B_H | rate constant of glycerol transporter in heart |
| Kdif_glyct_Glyc_B_H | dissociation constant of glycerol transporter for Glyc_B in heart |
| Kdif_glyct_Glyc_H | dissociation constant of glycerol transporter in heart |
| Vmax_glyk_Glyc_H | rate constant of glycerol kinase for Glyc in heart |
| Km_glyk_Glyc_H | Michaelis constant of glycerol kinase for Glyc in heart |
| Km_glyk_Atp_H | Michaelis constant of glycerol kinase for Atp in heart |
| Ki_glyk_Gap_H | inhibition constant of glycerol kinase for Gap in heart |
| Vdif_tgt_TG_B_H | rate constant of TG transporter in heart |
| Kdif_tgt_TG_B_H | dissociation constant of TG transporter in heart |
| Keq_tgt_TG_H | equilibrium constant of TG transporter in heart |
| Vmax_tgt_TG_B_H | rate constant of active TG transporter in heart |
| Km_tgt_TG_H | Michaelis constant of active TG transporter in heart |
| Vmax_g3pd_Gap_H | rate constant of glyceraldehyde-3-phosphate dehydrogenase for Gap in heart |
| Km_g3pd_Gap_H | Michaelis constant of glyceraldehyde-3-phosphate dehydrogenase for Gap in heart |
| Km_g3pd_Glycp_H | Michaelis constant of glyceraldehyde-3-phosphate dehydrogenase for Glycp in heart |
| Km_g3pd_Nadh_H | Michaelis constant of glyceraldehyde-3-phosphate dehydrogenase for Nadh in heart |
| Km_g3pd_Nad_H | Michaelis constant of glyceraldehyde-3-phosphate dehydrogenase for Nad in heart |
| Keq_g3pd_Glycp_H | equilibrium constant of glyceraldehyde-3-phosphate dehydrogenase for Glycp in heart |
| Vmax_boxid_FFA_H | rate constant of beta-oxidation for FFA in heart |
| Km_boxid_FFA_H | Michaelis constant of beta-oxidation for FFA in heart |
| Km_boxid_Atp_H | Michaelis constant of beta-oxidation for Atp in heart |
| Ki_boxid_Accoa_HM | inhibition constant of beta-oxidation for mitochondrial Accoa in heart |
| Ki_boxid_Malcoa_H | inhibition constant of beta-oxidation for Malcoa in heart |
| Km_boxid_Nad_H | Michaelis constant of beta-oxidation for Nad in heart |

| | |
|---|---|
| Km_boxid_Fad_H | Michaelis constant of beta-oxidation for Fad in heart |
| Vmax_accoat_Accoa_HM_HC | rate constant of mitochondrial Accoa transport to cytoplasm in heart |
| Km_accoat_Accoa_HM | Michaelis constant of mitochondrial Accoa transporter to cytoplasm in heart |
| Km_accoat_Atp_H | Michaelis constant of mitochondrial Accoa transporter to cytoplasm for Atp in heart |
| Km_accoat_Pyr_H | activation constant of mitochondrial Accoa transport to cytoplasm for Pyr in heart |
| Vmax_bhbsyn_Accoa_HM | rate constant of Bhb synthesis for mitochondrial Accoa in heart |
| Km_bhbsyn_Accoa_HM | Michaelis constant of Bhb synthesis for mitochondrial Accoa in heart |
| Km_bhbsyn_Nadh_H | Michaelis constant of Bhb synthesis for Nadh in heart |
| n_bhbsyn_Pyr_H | Hill coefficient of Bhb synthesis for Pyr in heart |
| Ki_bhbsyn_Pyr_H | inhibition constant of Bhb synthesis for Pyr in heart |
| Vmax_bhbdeg_Bhb_H | rate constant of Bhb degradation for Bhb in heart |
| Km_bhbdeg_Bhb_H | Michaelis constant of Bhb degradation for Bhb in heart |
| Km_bhbdeg_Atp_H | Michaelis constant of Bhb degradation for Atp in heart |
| Km_bhbdeg_Nad_H | Michaelis constant of Bhb degradation for Nad in heart |
| Vdif_bhbt_Bhb_B_H | rate constant of Bhb transporter in heart |
| Kdif_bhbt_Bhb_B_H | dissociation constant of Bhb transporter for Bhb_B in heart |
| Kdif_bhbt_Bhb_H | dissociation constant of Bhb transporter in heart |
| Vmax_lipog1_Accoa_HC | rate constant of lipogenesis 1 for Accoa in heart |
| Km_lipog1_Accoa_HC | Michaelis constant of lipogenesis 1 for Accoa in heart |
| Km_lipog1_Atp_H | Michaelis constant of lipogenesis 1 for Atp in heart |
| Vmax_lipog2_Malcoa_H | rate constant of lipogenesis 2 for Malcoa in heart |
| Km_lipog2_Malcoa_H | Michaelis constant of lipogenesis 2 for Malcoa in heart |
| Km_lipog2_Adp_H | Michaelis constant of lipogenesis 2 for Adp in heart |
| Km_lipog2_Nadph_H | Michaelis constant of lipogenesis 2 for Nadph in heart |
| Vmax_cholsyn1_Accoa_HC | rate constant of cholesterol synthesis 1 for Accoa in heart |
| Km_cholsyn1_Accoa_HC | Michaelis constant of cholesterol synthesis 1 for Accoa in heart |
| Vmax_cholsyn2_Hmgcoa_H | rate constant of cholesterol synthesis 2 for Hmgcoa in heart |
| Km_cholsyn2_Hmgcoa_H | Michaelis constant of cholesterol synthesis 2 for Hmgcoa in heart |
| Km_cholsyn2_Atp_H | Michaelis constant of cholesterol synthesis 2 for Atp in heart |
| Km_cholsyn2_Fadh_H | Michaelis constant of cholesterol synthesis 2 for Fadh in heart |
| Km_cholsyn2_Nadph_H | Michaelis constant of cholesterol synthesis 2 for Nadph in heart |
| Vmax_cholt_Chol_H | rate constant of active cholesterol transporter in heart |
| Km_cholt_Chol_H | Michaelis constant of active cholesterol transporter in heart |
| Vmax_atpsynf_Fadh_H | rate constant of Atp synthesis from Fadh in heart |
| Km_atpsynf_Fadh_H | Michaelis constant of Atp synthesis from Fadh for Fadh in heart |
| Km_atpsynf_Adp_H | Michaelis constant of Atp synthesis from Fadh for Adp in heart |

| | |
|---|---|
| Vmax_atpsynn_Nadh_H | rate constant of Atp synthesis from Nadh in heart |
| Km_atpsynn_Nadh_H | Michaelis constant of Atp synthesis from Nadh for Nadh in heart |
| Km_atpsynn_Adp_H | Michaelis constant of Atp synthesis from Nadh for Adp in heart |
| Vmax_atpuse_Atp_H | rate constant of Atp utilization for Atp in heart |
| Km_atpuse_Atp_H | Michaelis constant of Atp utilization for Atp in heart |
| Vmax_ampreg_Amp_H | rate constant of Amp regeneration for Amp in heart |
| Km_ampreg_Amp_H | Michaelis constant of Amp regeneration for Amp in heart |
| Km_ampreg_Atp_H | Michaelis constant of Amp regeneration for Atp in heart |
| Km_ampreg_Adp_H | Michaelis constant of Amp regeneration for Adp in heart |
| Vmax_nadhk_Nadh_H | rate constant of Nadh kinase for Nadh in heart |
| Km_nadhk_Nadh_H | Michaelis constant of Nadh kinase for Nadh in heart |
| Km_nadhk_Atp_H | Michaelis constant of Nadh kinase for Atp in heart |
| Vmax_nadhuse_Nadh_H | rate constant of Nadh utilization for Nadh in heart |
| Km_nadhuse_Nadh_H | Michaelis constant of Nadh utilization for Nadh in heart |
| Vmax_gdpreg_Gdp_H | rate constant of Gdp regeneration for Gdp in heart |
| Km_gdpreg_Gdp_H | Michaelis constant of Gdp regeneration for Gdp in heart |
| Km_gdpreg_Atp_H | Michaelis constant of Gdp regeneration for Atp in heart |
| Km_gdpreg_Gtp_H | Michaelis constant of Gdp regeneration for Gtp in heart |
| Km_gdpreg_Adp_H | Michaelis constant of Gdp regeneration for Adp in heart |
| Keq_gdpreg_Gtp_H | equilibrium constant of Gdp regeneration for Gtp in heart |
| Vmax_gtpuse_Gtp_H | rate constant of Gtp utilization for Gtp in heart |
| Km_gtpuse_Gtp_H | Michaelis constant of Gtp utilization for Gtp in heart |
| Vmax_udpreg_Udp_H | rate constant of Udp regeneration for Udp in heart |
| Km_udpreg_Udp_H | Michaelis constant of Udp regeneration for Udp in heart |
| Km_udpreg_Atp_H | Michaelis constant of Udp regeneration for Atp in heart |
| Km_udpreg_Utp_H | Michaelis constant of Udp regeneration for Utp in heart |
| Km_udpreg_Adp_H | Michaelis constant of Udp regeneration for Adp in heart |
| Keq_udpreg_Utp_H | equilibrium constant of Udp regeneration for Utp in heart |
| Vmax_utpuse_Utp_H | rate constant of Utp utilization for Utp in heart |
| Km_utpuse_Utp_H | Michaelis constant of Utp utilization for Utp in heart |
| Vmax_nadphuse_Nadph_H | rate constant of Nadph utilization for Nadph in heart |
| Km_nadphuse_Nadph_H | Michaelis constant of Nadph utilization for Nadph in heart |
| Vmax_fadhuse_Fadh_H | rate constant of Fadh utilization for Fadh in heart |
| Km_fadhuse_Fadh_H | Michaelis constant of Fadh utilization for Fadh in heart |
| Vmax_ck_Cre_H | rate constant of creatine kinase for Cre in heart |
| Km_ck_Cre_H | Michaelis constant of creatine kinase for Cre in heart |

| | |
|---|---|
| Km_ck_Atp_H | Michaelis constant of creatine kinase for Atp in heart |
| Km_ck_Crep_H | Michaelis constant of creatine kinase for Crep in heart |
| Km_ck_Adp_H | Michaelis constant of creatine kinase for Adp in heart |
| Keq_ck_Crep_H | equilibrium constant of creatine kinase for Crep in heart |
| Phos_H | phosphate in heart |
| alpha_base_N | insulin base factor in brain |
| alpha_band_N | insulin allowable width in brain |
| Km_Ins_B_N | Michaelis constant for insulin in brain |
| n_Ins_B_N | Hill coefficient for insulin in brain |
| Vdif_glut3_Glc_B_N | rate constant of glucose transporter type 2 in brain |
| Kdif_glut3_Glc_B_N | dissociation constant of glucose transporter type 2 for Glc_B in brain |
| Kdif_glut3_Glc_N | dissociation constant of glucose transporter type 2 in brain |
| Vmax_hk_Glc_N | reaction rate constant of hexokinase for Glc in brain |
| Km_hk_Glc_N | Michaelis constant of hexokinase for Glc in brain |
| Ki_hk_G6p_N | inhibition constant of hexokinase for Glc in brain |
| Km_hk_Atp_N | Michaelis constant of hexokinase for Atp in brain |
| Ki_hk_Atp_N | inhibition constant of hexokinase for Atp in brain |
| Vmax_ppp_G6p_N | reaction rate constant of pentose phosphate pathway for G6p in brain |
| Km_ppp_G6p_N | Michaelis constant of pentose phosphate pathway for G6p in brain |
| Km_ppp_Nadp_N | Michaelis constant of pentose phosphate pathway for Nadp in brain |
| Vmax_g6pase_G6p_N | reaction rate constant of G6Pase for G6p in brain |
| Km_g6pase_G6p_N | Michaelis constant of G6Pase for G6p in brain |
| Vmax_gs_G6p_N | reaction rate constant of glycogen synthase for G6p in brain |
| Km_gs_G6p_N | Michaelis constant of glycogen synthase for G6p in brain |
| n_gs_G6p_N | Hill coefficient of glycogen synthase for G6p in brain |
| Km_gs_Utp_N | Michaelis constant of glycogen synthase for G6p in brain |
| Glygn_N_max | maximum glycogen concentration in brain |
| Km_gs_Glygn_N | Michaelis constant of glycogen synthase for glycogen in brain |
| Vmax_gd_Glygn_N | reaction rate constant of glycogen degradation for glycogen in brain |
| Km_gd_Glygn_N | Michaelis constant of glycogen degradation for glycogen in brain |
| Km_gd_Phos_N | Michaelis constant of glycogen degradation for phosphate in brain |
| Vmax_pfk_G6p_N | reaction rate constant of phosphofructokinase for G6p in brain |
| Km_pfk_G6p_N | Michaelis constant of phosphofructokinase for G6p in brain |
| Km_pfk_Atp_N | Michaelis constant of phosphofructokinase for Atp in brain |
| Ki_pfk_Atp_N | inhibition constant of phosphofructokinase for Atp in brain |
| Km_pfk_Adp_N | Michaelis constant of phosphofructokinase for Adp in brain |

| | |
|---|---|
| Ki_pfk_Gap_N | inhibition constant of phosphofructokinase for Gap in brain |
| b_pfk_Gap_N | coefficient of phosphofructokinase for Gap in brain |
| Vmax_fbp_Gap_N | reaction rate constant of fructose bisphosphatase for Gap in brain |
| Km_fbp_Gap_N | Michaelis constant of fructose bisphosphatase for Gap in brain |
| Vmax_pk_Gap_N | reaction rate constant of pyruvate kinase for Gap in brain |
| Km_pk_Gap_N | Michaelis constant of pyruvate kinase for Gap in brain |
| b_pk_Accoa_NM | coefficient of pyruvate kinase for mitochondrial Accoa in brain |
| Ki_pk_Accoa_NM | inhibition constant of pyruvate kinase for mitochondrial Accoa in brain |
| Km_pk_Adp_N | Michaelis constant of pyruvate kinase for Adp in brain |
| Vmax_pepck_Pyr_N | reaction rate constant of phosphoenolpyruvate carboxykinase for Pyr in brain |
| Km_pepck_Pyr_N | Michaelis constant of phosphoenolpyruvate carboxykinase for Pyr in brain |
| Km_pepck_Atp_N | Michaelis constant of phosphoenolpyruvate carboxykinase for Atp in brain |
| Km_pepck_Gtp_N | Michaelis constant of phosphoenolpyruvate carboxykinase for Gtp in brain |
| Vdif_pyrt_Pyr_B_N | rate constant of pyruvate transporter in brain |
| Kdif_pyrt_Pyr_B_N | dissociation constant of pyruvate transporter for Pyr_B in brain |
| Kdif_pyrt_Pyr_N | dissociation constant of pyruvate transporter in brain |
| Vdif_lact_Lac_B_N | rate constant of lactate transporter in brain |
| Kdif_lact_Lac_N | dissociation constant of lactate transporter in brain |
| Kdif_lact_Lac_B_N | dissociation constant of lactate transporter for Lac_B in brain |
| Vmax_ldh_Pyr_N | rate constant of lactate dehydrogenase for Pyr in brain |
| Km_ldh_Pyr_N | Michaelis constant of lactate dehydrogenase for Pyr in brain |
| Km_ldh_Nadh_N | Michaelis constant of lactate dehydrogenase for Nadh in brain |
| Km_ldh_Lac_N | Michaelis constant of lactate dehydrogenase for Lac in brain |
| Km_ldh_Nad_N | Michaelis constant of lactate dehydrogenase for Nad in brain |
| Keq_ldh_Lac_N | equilibrium constant of lactate dehydrogenase for Lac in brain |
| Vdif_alat_Ala_B_N | rate constant of alanine transporter in brain |
| Kdif_alat_Ala_N | dissociation constant of alanine transporter in brain |
| Kdif_alat_Ala_B_N | dissociation constant of alanine transporter for Ala_B in brain |
| Vmax_alata_Pyr_N | rate constant of alanine transporter for Pyr in brain |
| Km_alata_Pyr_N | Michaelis constant of alanine aminotransferase for Pyr in brain |
| Km_alata_Ala_N | Michaelis constant of alanine aminotransferase for Ala in brain |
| Keq_alata_Ala_N | equilibrium constant of alanine aminotransferase for Ala in brain |
| Vmax_pdh_Pyr_N | rate constant of pyruvate dehydrogenase for Pyr in brain |
| Km_pdh_Pyr_N | Michaelis constant of pyruvate dehydrogenase for Pyr in brain |
| Ki_pdh_Accoa_NM | inhibition constant of pyruvate dehydrogenase for mitochondrial Accoa in brain |
| Km_pdh_Nad_N | Michaelis constant of pyruvate dehydrogenase for Nad in brain |

| | |
|---|---|
| Vmax_tca_Accoa_NM | rate constant of TCA cycle for mitochondrial Accoa in brain |
| Km_tca_Accoa_NM | Michaelis constant of TCA cycle for mitochondrial Accoa in brain |
| Km_tca_Phos_N | Michaelis constant of TCA cycle for phosphate in brain |
| Km_tca_Adp_N | Michaelis constant of TCA cycle for Adp in brain |
| Km_tca_Pyr_N | Michaelis constant of TCA cycle for Pyr in brain |
| Km_tca_Nad_N | Michaelis constant of TCA cycle for Nad in brain |
| Km_tca_Fad_N | Michaelis constant of TCA cycle for Fad in brain |
| Vdif_ffat_FFA_B_N | rate constant of FFA transporter in brain |
| Kdif_ffat_FFA_B_N | dissociation constant of FFA transporter for FFA_B in brain |
| Kdif_ffat_FFA_N | dissociation constant of FFA transporter in brain |
| Vmax_ffat_FFA_B_N | rate constant of active FFA transporter in brain |
| Km_ffat_FFA_B_N | Michaelis constant of active FFA transporter in brain |
| Vmax_tgsyn_FFA_N | rate constant of TG synthesis for FFA in brain |
| Km_tgsyn_FFA_N | Michaelis constant of TG synthesis for FFA in brain |
| Km_tgsyn_Glycp_N | Michaelis constant of TG synthesis for Glycp in brain |
| Km_tgsyn_Atp_N | Michaelis constant of TG synthesis for Atp in brain |
| TG_N_max | maximum TG concentration in brain |
| Km_tgsyn_TG_N | Michaelis constant of TG synthesis for TG in brain |
| Vmax_tgdeg_TG_N | rate constant of TG degradation for TG in brain |
| Km_tgdeg_TG_N | Michaelis constant of TG degradation for TG in brain |
| Vdif_glyct_Glyc_B_N | rate constant of glycerol transporter in brain |
| Kdif_glyct_Glyc_B_N | dissociation constant of glycerol transporter for Glyc_B in brain |
| Kdif_glyct_Glyc_N | dissociation constant of glycerol transporter in brain |
| Vmax_glyk_Glyc_N | rate constant of glycerol kinase for Glyc in brain |
| Km_glyk_Glyc_N | Michaelis constant of glycerol kinase for Glyc in brain |
| Km_glyk_Atp_N | Michaelis constant of glycerol kinase for Atp in brain |
| Ki_glyk_Gap_N | inhibition constant of glycerol kinase for Gap in brain |
| Vdif_tgt_TG_B_N | rate constant of TG transporter in brain |
| Kdif_tgt_TG_B_N | dissociation constant of TG transporter in brain |
| Keq_tgt_TG_N | equilibrium constant of TG transporter in brain |
| Vmax_tgt_TG_B_N | rate constant of active TG transporter in brain |
| Km_tgt_TG_N | Michaelis constant of active TG transporter in brain |
| Vmax_g3pd_Gap_N | rate constant of glyceraldehyde-3-phosphate dehydrogenase for Gap in brain |
| Km_g3pd_Gap_N | Michaelis constant of glyceraldehyde-3-phosphate dehydrogenase for Gap in brain |
| Km_g3pd_Glycp_N | Michaelis constant of glyceraldehyde-3-phosphate dehydrogenase for Glycp in brain |
| Km_g3pd_Nadh_N | Michaelis constant of glyceraldehyde-3-phosphate dehydrogenase for Nadh in brain |

| | |
|---|---|
| Km_g3pd_Nad_N | Michaelis constant of glyceraldehyde-3-phosphate dehydrogenase for Nad in brain |
| Keq_g3pd_Glycp_N | equilibrium constant of glyceraldehyde-3-phosphate dehydrogenase for Glycp in brain |
| Vmax_boxid_FFA_N | rate constant of beta-oxidation for FFA in brain |
| Km_boxid_FFA_N | Michaelis constant of beta-oxidation for FFA in brain |
| Km_boxid_Atp_N | Michaelis constant of beta-oxidation for Atp in brain |
| Ki_boxid_Accoa_NM | inhibition constant of beta-oxidation for mitochondrial Accoa in brain |
| Ki_boxid_Malcoa_N | inhibition constant of beta-oxidation for Malcoa in brain |
| Km_boxid_Nad_N | Michaelis constant of beta-oxidation for Nad in brain |
| Km_boxid_Fad_N | Michaelis constant of beta-oxidation for Fad in brain |
| Vmax_accoat_Accoa_NM_NC | rate constant of mitochondrial Accoa transport to cytoplasm in brain |
| Km_accoat_Accoa_NM | Michaelis constant of mitochondrial Accoa transporter to cytoplasm in brain |
| Km_accoat_Atp_N | Michaelis constant of mitochondrial Accoa transporter to cytoplasm for Atp in brain |
| Km_accoat_Pyr_N | activation constant of mitochondrial Accoa transport to cytoplasm for Pyr in brain |
| Vmax_bhbsyn_Accoa_NM | rate constant of Bhb synthesis for mitochondrial Accoa in brain |
| Km_bhbsyn_Accoa_NM | Michaelis constant of Bhb synthesis for mitochondrial Accoa in brain |
| Km_bhbsyn_Nadh_N | Michaelis constant of Bhb synthesis for Nadh in brain |
| n_bhbsyn_Pyr_N | Hill coefficient of Bhb synthesis for Pyr in brain |
| Ki_bhbsyn_Pyr_N | inhibition constant of Bhb synthesis for Pyr in brain |
| Vmax_bhbdeg_Bhb_N | rate constant of Bhb degradation for Bhb in brain |
| Km_bhbdeg_Bhb_N | Michaelis constant of Bhb degradation for Bhb in brain |
| Km_bhbdeg_Atp_N | Michaelis constant of Bhb degradation for Atp in brain |
| Km_bhbdeg_Nad_N | Michaelis constant of Bhb degradation for Nad in brain |
| Vdif_bhbt_Bhb_B_N | rate constant of Bhb transporter in brain |
| Kdif_bhbt_Bhb_B_N | dissociation constant of Bhb transporter for Bhb_B in brain |
| Kdif_bhbt_Bhb_N | dissociation constant of Bhb transporter in brain |
| Vmax_lipog1_Accoa_NC | rate constant of lipogenesis 1 for Accoa in brain |
| Km_lipog1_Accoa_NC | Michaelis constant of lipogenesis 1 for Accoa in brain |
| Km_lipog1_Atp_N | Michaelis constant of lipogenesis 1 for Atp in brain |
| Vmax_lipog2_Malcoa_N | rate constant of lipogenesis 2 for Malcoa in brain |
| Km_lipog2_Malcoa_N | Michaelis constant of lipogenesis 2 for Malcoa in brain |
| Km_lipog2_Adp_N | Michaelis constant of lipogenesis 2 for Adp in brain |
| Km_lipog2_Nadph_N | Michaelis constant of lipogenesis 2 for Nadph in brain |
| Vmax_cholsyn1_Accoa_NC | rate constant of cholesterol synthesis 1 for Accoa in brain |
| Km_cholsyn1_Accoa_NC | Michaelis constant of cholesterol synthesis 1 for Accoa in brain |
| Vmax_cholsyn2_Hmgcoa_N | rate constant of cholesterol synthesis 2 for Hmgcoa in brain |

| | |
|---|---|
| Km_cholsyn2_Hmgcoa_N | Michaelis constant of cholesterol synthesis 2 for Hmgcoa in brain |
| Km_cholsyn2_Atp_N | Michaelis constant of cholesterol synthesis 2 for Atp in brain |
| Km_cholsyn2_Fadh_N | Michaelis constant of cholesterol synthesis 2 for Fadh in brain |
| Km_cholsyn2_Nadph_N | Michaelis constant of cholesterol synthesis 2 for Nadph in brain |
| Vmax_cholt_Chol_N | rate constant of active cholesterol transporter in brain |
| Km_cholt_Chol_N | Michaelis constant of active cholesterol transporter in brain |
| Vmax_atpsynf_Fadh_N | rate constant of Atp synthesis from Fadh in brain |
| Km_atpsynf_Fadh_N | Michaelis constant of Atp synthesis from Fadh for Fadh in brain |
| Km_atpsynf_Adp_N | Michaelis constant of Atp synthesis from Fadh for Adp in brain |
| Vmax_atpsynn_Nadh_N | rate constant of Atp synthesis from Nadh in brain |
| Km_atpsynn_Nadh_N | Michaelis constant of Atp synthesis from Nadh for Nadh in brain |
| Km_atpsynn_Adp_N | Michaelis constant of Atp synthesis from Nadh for Adp in brain |
| Vmax_atpuse_Atp_N | rate constant of Atp utilization for Atp in brain |
| Km_atpuse_Atp_N | Michaelis constant of Atp utilization for Atp in brain |
| Vmax_ampreg_Amp_N | rate constant of Amp regeneration for Amp in brain |
| Km_ampreg_Amp_N | Michaelis constant of Amp regeneration for Amp in brain |
| Km_ampreg_Atp_N | Michaelis constant of Amp regeneration for Atp in brain |
| Km_ampreg_Adp_N | Michaelis constant of Amp regeneration for Adp in brain |
| Vmax_nadhk_Nadh_N | rate constant of Nadh kinase for Nadh in brain |
| Km_nadhk_Nadh_N | Michaelis constant of Nadh kinase for Nadh in brain |
| Km_nadhk_Atp_N | Michaelis constant of Nadh kinase for Atp in brain |
| Vmax_nadhuse_Nadh_N | rate constant of Nadh utilization for Nadh in brain |
| Km_nadhuse_Nadh_N | Michaelis constant of Nadh utilization for Nadh in brain |
| Vmax_gdpreg_Gdp_N | rate constant of Gdp regeneration for Gdp in brain |
| Km_gdpreg_Gdp_N | Michaelis constant of Gdp regeneration for Gdp in brain |
| Km_gdpreg_Atp_N | Michaelis constant of Gdp regeneration for Atp in brain |
| Km_gdpreg_Gtp_N | Michaelis constant of Gdp regeneration for Gtp in brain |
| Km_gdpreg_Adp_N | Michaelis constant of Gdp regeneration for Adp in brain |
| Keq_gdpreg_Gtp_N | equilibrium constant of Gdp regeneration for Gtp in brain |
| Vmax_gtpuse_Gtp_N | rate constant of Gtp utilization for Gtp in brain |
| Km_gtpuse_Gtp_N | Michaelis constant of Gtp utilization for Gtp in brain |
| Vmax_udpreg_Udp_N | rate constant of Udp regeneration for Udp in brain |
| Km_udpreg_Udp_N | Michaelis constant of Udp regeneration for Udp in brain |
| Km_udpreg_Atp_N | Michaelis constant of Udp regeneration for Atp in brain |
| Km_udpreg_Utp_N | Michaelis constant of Udp regeneration for Utp in brain |
| Km_udpreg_Adp_N | Michaelis constant of Udp regeneration for Adp in brain |

| | |
|---|---|
| Keq_udpreg_Utp_N | equilibrium constant of Udp regeneration for Utp in brain |
| Vmax_utpuse_Utp_N | rate constant of Utp utilization for Utp in brain |
| Km_utpuse_Utp_N | Michaelis constant of Utp utilization for Utp in brain |
| Vmax_nadphuse_Nadph_N | rate constant of Nadph utilization for Nadph in brain |
| Km_nadphuse_Nadph_N | Michaelis constant of Nadph utilization for Nadph in brain |
| Vmax_fadhuse_Fadh_N | rate constant of Fadh utilization for Fadh in brain |
| Km_fadhuse_Fadh_N | Michaelis constant of Fadh utilization for Fadh in brain |
| Vmax_ck_Cre_N | rate constant of creatine kinase for Cre in brain |
| Km_ck_Cre_N | Michaelis constant of creatine kinase for Cre in brain |
| Km_ck_Atp_N | Michaelis constant of creatine kinase for Atp in brain |
| Km_ck_Crep_N | Michaelis constant of creatine kinase for Crep in brain |
| Km_ck_Adp_N | Michaelis constant of creatine kinase for Adp in brain |
| Keq_ck_Crep_N | equilibrium constant of creatine kinase for Crep in brain |
| Phos_N | phosphate in brain |
