## Supplemental Figure 1 for "Virtual metabolic human dynamic model for pathological analysis and therapy design for diabetes"

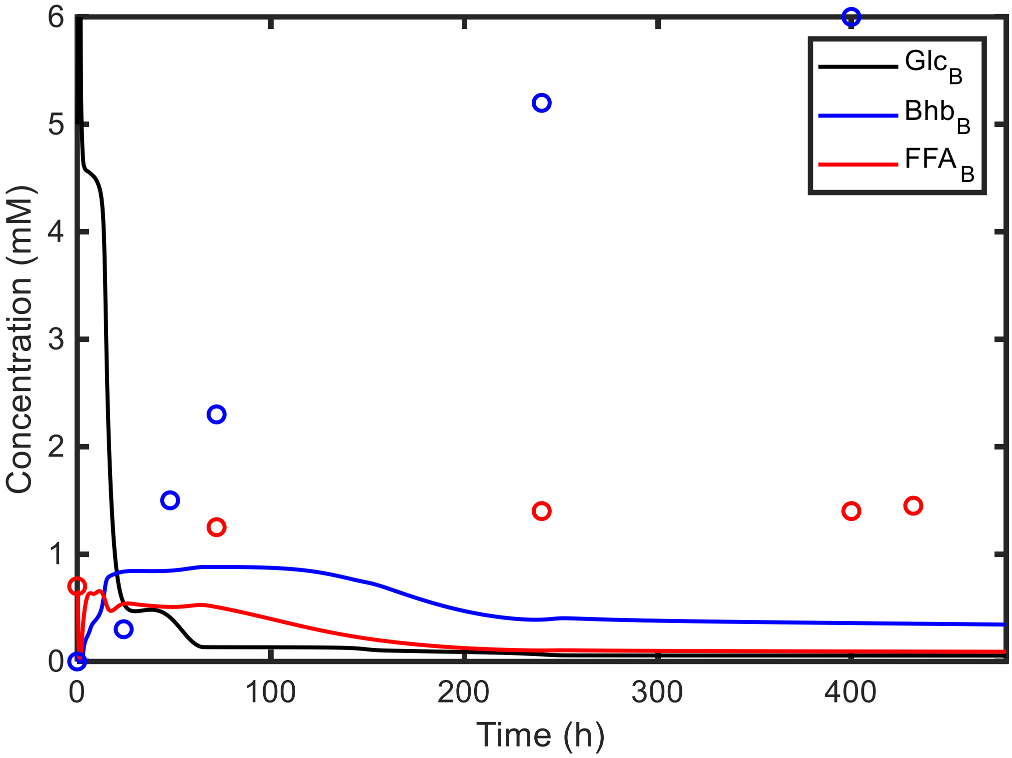


Figure S1 Time-course of key metabolite concentrations under a fasted condition.

The proposed model was simulated during 480 h after an overnight fast and following a single meal of 100 g glucose and 33 g fat. The blue circles and blue lines indicate the experimental and simulated concentrations of plasma Bhb, respectively. The red circles and red lines indicate the experimental and simulated concentrations of plasma FFA, respectively. The black line indicates the plasma glucose concentration.
